# Oncologic therapy shapes the fitness landscape of clonal hematopoiesis

**DOI:** 10.1101/848739

**Authors:** Kelly L Bolton, Ryan N Ptashkin, Teng Gao, Lior Braunstein, Sean M Devlin, Daniel Kelly, Minal Patel, Antonin Berthon, Aijazuddin Syed, Mariko Yabe, Catherine C. Coombs, Nicole M. Caltabellotta, Mike Walsh, Kenneth Offit, Zsofia Stadler, Diana Mandelker, Jessica Schulman, Akshar Patel, John Philip, Elsa Bernard, Gunes Gundem, Juan E Arango, Max Levine, Juan S Medina, Noushin Farnoud, Dominik Glodzik, Sonya Li, Marc E Robson, Choonsik Lee, Paul D P Pharoah, Konrad Stopsack, Barbara Spitzer, Simon Mantha, James Fagin, Laura Boucai, Christopher J Gibson, Benjamin L Ebert, Andrew Young, Todd Druley, Koichi Takahasi, Nancy Gillis, Markus Ball, Eric Padron, David M Hyman, Jose Baselga, Larry Norton, Stuart Gardos, Virginia M Klimek, Howard Scher, Dean Bajorin, Eder Paraiso, Ryma Benayed, Maria E Arcila, Marc Ladanyi, David B Solit, Michael F Berger, Martin Tallman, Montserrat Garcia-Closas, Nilanjan Chatterjee, Luis A Diaz, Ross L Levine, Lindsay M Morton, Ahmet Zehir, Elli Papaemmanuil

## Abstract

Clonal hematopoiesis (CH) is frequent in cancer patients and associated with increased risk of therapy related myeloid neoplasms (tMN). To define the relationship between CH, oncologic therapy, and tMN progression, we studied 24,439 cancer patients. We show that previously treated patients have increased rates of CH, with enrichment of mutations in DNA Damage Response (DDR) genes (*TP53, PPM1D, CHEK2*). Exposure to radiation, platinum and topoisomerase II inhibitors have the strongest association with CH with evidence of dose-dependence and gene-treatment interactions. We validate these associations in serial sampling from 525 patients and show that exposure to cytotoxic and radiation therapy imparts a selective advantage specifically in hematopoietic cells with DDR mutations. In patients who progressed to tMN, the clone at CH demarcated the dominant clone at tMN diagnosis. CH mutational features predict risk of therapy-related myeloid neoplasm in solid tumor patients with clinical implications for early detection and treatment decisions.

## MAIN

Clonal hematopoiesis (CH) is emerging as an important clinical biomarker for early detection and management of individuals at risk of myeloid neoplasms (MN)^1, 2^. CH is characterized by clonal mutations in hematopoietic stem and progenitor cells (HSPCs) in the absence of overt hematologic disease^3–5^. CH is most commonly associated with one mutation or few mutations in genes recognized as early and initiating events in myeloid disease, such as epigenetic modifiers (*DNMT3A, TET2, ASXL1, IDH1, IDH2*), splicing factors (*SF3B1, SRSF2, U2AF1*), *TP53* and *JAK2*^3, 4, 6^. CH is also frequent in patients with solid tumors^7–10^. This is largely driven by a shared association with age. However, recent studies further propose a link between prior exposure to oncologic therapy and CH^7, 11^.

Cancer patients are at a heightened risk for developing myeloid neoplasms including AML and MDS.^12^ When myeloid neoplasms arise following exposure to oncologic therapy, they are referred to as therapy-related (tMN) and represent one of the most aggressive and chemo-resistant malignancies, with a 5-year survival of <10%^13^. With increasing cancer survivorship, the incidence of tMNs is rising. Thus, there is a clear, unmet need to develop a deeper understanding of the pathogenesis of tMN, to inform tMN screening and prevention programs, and to identify novel therapeutic targets for tMN^13^. While tMN was traditionally thought to develop from the mutagenic effects of oncologic therapy^13^, recent studies have shown that tMN-initiating mutations present in hematopoietic cells can predate the receipt of oncologic therapy^14^. Recent studies have linked CH to an increased risk of tMN^7, 15–17^.

Studies of CH offer insights into the first clonal expansion of the multi-step process of carcinogenesis, whereby a single mutation underwrites the transition of a normal cell to one with a considerably stronger fitness advantage. In a small proportion of carriers, CH may lead to overt myeloid disease, but most frequently these clones remain stable throughout life^18^. The ensuing genetic and clonal trajectories are likely shaped by a dynamic interplay of cell intrinsic and extrinsic factors. We sought to investigate the relationships between CH and oncologic therapy exposure amongst other parameters (demographic, clinical, smoking). We study how oncologic therapy shapes CH clonal dynamics in a cohort of prospectively followed patients in which CH status was assessed before and after oncologic therapy. Understanding these associations will offer opportunities for early diagnosis and prevention strategies for cancer patients at high risk for tMN and provide mechanistic insights into tMN pathogenesis with potential therapeutic relevance.

### Molecular presentation of clonal hematopoiesis in cancer patients

We analyzed data from 24,439 cancer patients across a wide range of primary tumor types (N=57) and ages (Extended Data Table 1). CH mutations were identified from targeted, deep coverage next-generation sequencing data (MSK-IMPACT) generated from paired peripheral blood and tumor samples as part of clinical care. We defined CH as a somatic mutation in blood with a minimum variant allele frequency (VAF) of 2%. For further details on CH calling refer to the Methods and Supplementary Notes.

We identified a total of 11,391 unique variants in 7,379 individuals, representing 30% of patients in our cohort. The spectrum of CH mutations in our cohort followed expected patterns of enrichment for truncating variants and hotspot mutations in tumor suppressor and oncogenes respectively (Supplementary Fig. 1). The median VAF of CH mutations was 4.7% but the range was broad (range=2-87%). Among individuals with CH, 68% (n=5,044) had one mutation and 2,335 (32%) had two or more mutations. Consistent with prior literature in healthy individuals^3, 4^, CH mutations were most frequently identified in *DNMT3A* and *TET2*. Mutations in key regulators of DNA Damage Response (DDR) pathway such as *PPM1D*, *TP53* and *CHEK2* were also frequently mutated in our cohort, in line with prior evidence that DDR mutations are enriched in cancer patients exposed to oncologic therapy^7, 11^(Supplementary Fig. 2).

Given that by design we only interrogate bona fide cancer genes, we annotated each mutation on the basis of its putative role in cancer pathogenesis using OncoKB^19^ and recurrence in an in-house dataset of myeloid neoplasms^20–22^(see Methods for more details). Over half of the CH mutations that we detected were classified as putative driver mutations of cancer (CH-PD, 53%, n=6,028). Almost all CH-PD variants (90%, n=5,453) were recurrent mutations in myeloid neoplasms (CH-myeloid PD) (Supplementary Fig. 2). The strong enrichment of myeloid variants highlights the strength of the fitness advantage imparted on HSPCs by mutations in genes implicated in myeloid pathogenesis as compared to oncogenic mutations in other cancer gene drivers.

The prevalence of CH among cancer patients differed by primary tumor type even after adjustment for age (Extended Data Fig. 1). While the overall mutational spectrum of CH was similar across cancer subtypes, mutations in DDR genes were markedly more frequent in patients with ovarian and endometrial cancer. This enrichment was most striking for mutations in *PPM1D*, which were found in 14% of patients with ovarian cancer and 7% of patients with endometrial cancer as compared to <5% in other cancer subgroups (Extended Data Fig. 2). Recent studies show that cell lines with *PPM1D* mutations outcompete normal cells after exposure to cisplatin but not under normal conditions^23^. Thus, the differential enrichment of *PPM1D* mutations by tumor site may, in part, reflect the increased fitness of cells with *PPM1D* mutations under platinum exposure. The observation of tumor-specific CH mutational spectra, including enrichment for DDR mutations in patients with select primary tumor types, points towards the existence of gene-treatment interactions with specific classes of oncologic therapy.

### Clinical associations with clonal hematopoiesis in cancer patients

To determine how CH is influenced by prior cancer therapy, we extracted treatment information on systemic oncologic therapy and external beam radiation therapy including agent class, dosage, drug combination regimens and treatment timing for 10,207 patients who had received their cancer care at Memorial Sloan Kettering (MSK). Data on gender, age at the time of blood draw for sequencing, ethnicity, smoking history and blood count indices within one year from blood draw were also collected (see Supplementary Notes).

A total of 6,240 patients (61%) were exposed to oncologic therapy (including cytotoxic therapy, radiation therapy, targeted therapy and immunotherapy) prior to CH testing (Extended Data Fig. 3), whereas 3,967 (39%) were treatment naive. This cohort provided sufficient statistical power to conduct a detailed evaluation of the relationships between prior oncologic therapy and CH while accounting for demographic factors (gender, age, ethnicity) and smoking history. CH was positively associated with increasing decile of age (OR=1.8, p=<10^-6^), being a current or former smoker (OR=1.1, p=4.1×10^-3^), and prior exposure to oncologic therapy (OR=1.2, p=4.2×10^-6^) and was less common in Asians than in Caucasian patients (OR=0.7, p=9×10^-4^) (Figure 1a, Extended Data Table 2). In both treated and untreated patients, CH mutations in genes associated with myeloid neoplasms (CH-myeloid) and CH-PD mutations showed stronger associations with increasing age (Fig. 1b, Extended Data Table 2) and had higher VAFs compared to non-PD CH mutations (Supplementary Fig. 3a-b, Extended Data Table 3) suggesting the presence of cell-context/mutation specific effects on clonal selection. The VAF of mutations in patients with CH who harbored multiple mutations was higher compared to individuals with one mutation (Extended Data Table 3, Supplementary Fig. 3c). We observed an enrichment for a higher total number of CH mutations with increasing age, prior treatment and smoking exposure (p=<1×10^-6^, p=0.01, and p=0.02, respectively) and increased clonal dominance as estimated by VAF metrics with increasing age and in smokers (p=0.004 and p=1.3×10^-5^, respectively) (Extended Data Table 3–4).

**Figure 1.**
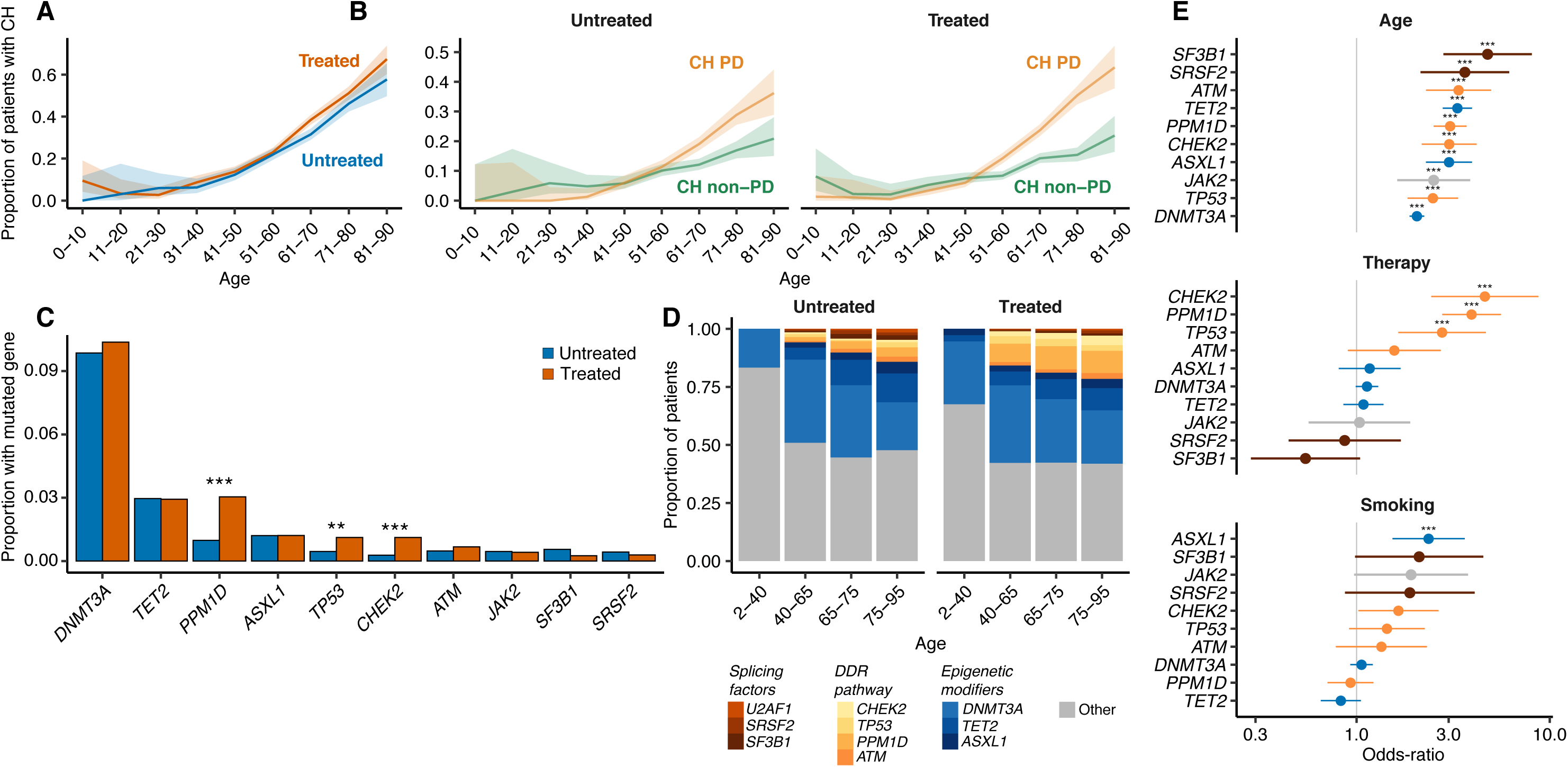
Characteristics of CH by age, prior therapy exposure and smoking history. (A) Proportion of patients with CH among previously treated (received any oncologic therapy prior to blood draw, N=6240) and untreated patients (N=3967) by ten-year age group. (B) Proportion of patients with CH due to a known cancer driver mutation (CH-PD) and those without a known driver (CH non-PD) stratified by prior treatment exposure (C) Proportion of patients with CH mutations in specific genes among treated and untreated patients. Multivariable logistic regression was used to test whether the odds of having a specific gene mutated differed among treated and untreated patients after adjustment for age, gender, smoking and race. (D) Among patients with CH, the proportion with mutations in specific genes, by age group and treatment status. (E) Odds Ratio for CH mutation in ten most commonly mutated genes with 1) increasing age, 2) for patients previously exposed to oncologic therapy compared to those with no exposure prior to blood draw, 3) for current/former smokers compared to non-smokers in multivariable logistic regression models adjusted for therapy, smoking, race, age, gender and time from diagnosis to blood draw. Shown are the q-values (FDR-corrected p-values): * <0.05, ** <0.01,***<0.001. Age is expressed as the mean centered value.

Clonal selection is likely multifactorial, with the realized fitness of cells with specific gene mutations dependent on both cell-intrinsic and environmental parameters. We studied, therefore, how age, race, smoking and prior exposure to oncologic therapy were related to the representation of specific gene mutations through multivariable logistic regression. Mutations in the spliceosome genes *SRSF2* and *SF3B1* were less common than other CH mutations, but showed the strongest enrichment with age (Fig. 1d-e). CH mutations in the DDR genes *TP53, PPM1D* and *CHEK2* were strongly associated with prior exposure to oncologic therapy (OR*_TP53_* = 2.7, q=9.0×10^-4^; OR*_PPM1D_* = 3.6, q= 1.2×10^-5^; OR*_CHEK2_*=4.6, q=<10^-6^, Fig. 1e). Mutations in *ASXL1* were significantly associated with prior smoking history (OR=2.5, q=2.0×10^-4^, Fig. 1e). These associations provide evidence that environmental factors such as oncologic treatment and smoking influence the fitness of specific gene mutations in HSPC’s. While there was an overall higher prevalence of CH in treated patients, the fitness associated with mutations in epigenetic modifiers (*DNMT3A, TET2*) or splicing regulators (*SRSF2, SF3B1, U2AF1*) were not strongly modulated by oncologic therapy (Fig. 1d-e). Overall, the patterns of acquired mutations were similar in treated and untreated patients in regards to mutational consequence and proportion of C to T transitions within a CpG tri-nucleotide context (Supplementary Fig. 4–5). This was true even within DDR-CH genes and *ASXL1* when stratified among smoking and treatment status (Supplementary Fig. 5).

### Associations between clonal hematopoiesis and subclasses of oncologic therapy

Subjects in our study were exposed to a total of 492 different systemic cancer-directed therapies, which we classified according to mechanism of action (Supplementary Notes and Supplementary Table 1). To define the strength of the association between CH-PD and therapy subclass, we performed multivariable logistic regression, adjusted for demographic parameters, smoking and time from solid tumor diagnosis to blood draw. After accounting for exposure to all broad therapy subclasses, CH-PD was positively associated with prior exposure to radionuclide therapy (OR=1.5, p=0.03), external beam radiation therapy (OR=1.4, p=<10^-6^) and cytotoxic chemotherapy (OR=1.2, p=8×10^-4^) but not targeted therapy or immunotherapy (Fig. 2a). With respect to subclasses of cytotoxic therapy, CH-PD was most strongly associated with prior exposure to topoisomerase II inhibitors (OR=1.3, p=0.01) and platinum agents (OR=1.2, p=0.01) (Fig. 2a) after accounting for exposure to major classes of cytotoxic therapy, immunotherapy and external beam radiation therapy. Among platinum agents, CH-PD was significantly associated with prior exposure to carboplatin (OR=1.3, p=0.002) but not cisplatin (OR=1.1, p=0.20) or oxaliplatin (OR=1.1, p=0.63) (Fig. 2a). This is in line with evidence that rates of tMN are highest following exposure to carboplatin^24^.

**Figure 2.**
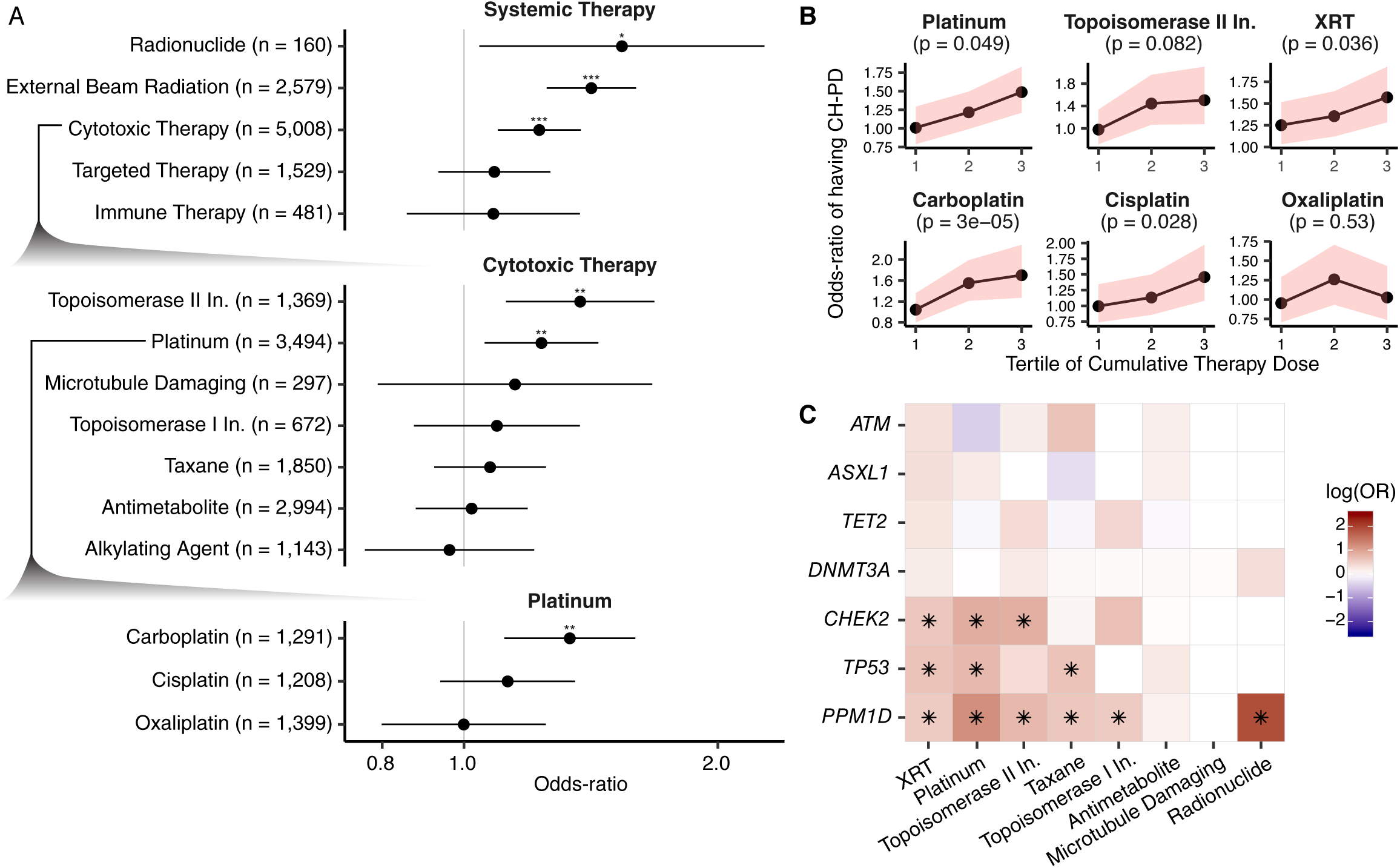
Association between CH and prior exposure to oncologic therapy. (A) Odds ratios (OR) and 95% confidence intervals for CH-PD and specific classes of oncologic therapy in multivariable logistic regression adjusted for each other, smoking, race, gender and time from diagnosis to blood draw. The top panel shows the OR for broad classes of oncologic therapy. The middle panel shows the OR between CH-PD and prior exposure to subclasses of cytotoxic therapy. The bottom panel shows the OR between CH-PD and exposure to specific platinum-based drugs. (B) The OR between prior receipt oncologic therapy and CH-PD stratified by tertile of cumulative exposure for the agent. Multivariable logistic regression was used adjusted as in (A) but with cumulative weight-adjusted dose of systemic therapy classes and cumulative radiation dose (as expressed in EQD_2_. The p-trend was calculated testing for an association between CH and increasing tertiles of cumulative oncologic therapy exposure among those who received the therapy in the multivariable model. (C) Heat-map showing the log(OR) between CH-PD in specific genes and prior exposure to the major classes of cytotoxic therapy and radiation therapy in logistic regression models adjusted for each all therapy subclasses, smoking, race, gender and time from diagnosis to blood draw. Q-values (FDR-corrected p-values) are shown with an asterix: *<0.05, ** <0.01, ***<0.001.

The risk of tMN increases with cumulative exposure to cytotoxic therapy^25, 26^ and ionizing radiation^27^. To evaluate for the presence of dose-response relationships with CH-PD, we calculated each patient’s relative cumulative exposure to specific therapy classes (see Supplementary Notes and Supplementary Fig. 6). After adjustment for cumulative exposure to all major classes of therapy, increasing exposure to external beam radiation therapy and platinum chemotherapy was positively associated with CH-PD (p-trend=0.04 and 0.05 respectively) (Fig. 2b). A similar positive trend was seen for CH-PD and higher cumulative exposure to topoisomerase II inhibitors although the test for trend was not significant (Fig. 2b). Evaluation of dose response relationships with platinum agents showed that CH-PD was associated with higher cumulative doses of carboplatin (p-trend=3×10^-5^) and cisplatin (p-trend=0.03) further supporting the robustness of the association between platinum agents and CH. Considering that chemotherapy is often administered in multi-drug combinations, we explored associations between specific drug regimens and CH-PD in a subset of patients (N=5,594) for which full details on drug regimen was available. Regimens containing carboplatin or cisplatin were most strongly associated with CH (Extended Data Fig. 4).

We next tested for associations between subclasses of oncologic therapy and CH-PD gene mutations, considering gene-treatment combinations with at least 10 individuals. Mutations in *PPM1D*, were strongly associated with prior exposure to radionuclide therapy (OR=6.6, q=2.7×10^-6^) and platinum (OR=3.3, q=<10^-6^) and also showed associations with topoisomerase II inhibitors (OR=2.0, q=0.002), taxanes (OR=1.8, q=0.003), topoisomerase I inhibitors (OR=1.7, q=0.003) and external beam radiation therapy (OR=1.9, q=0.03) (Fig. 2c). Mutations in *TP53* were associated with platinum (OR=2.1, q=0.03), radiation therapy (OR=1.9, q=0.03) and taxanes (OR=1.9, q=0.05) and *CHEK2* was associated with platinum (OR=2.3, q=0.02), topoisomerase II inhibitors (OR=2.3, q=0.02) and external beam radiation therapy (OR=1.7, q=0.05) (Fig. 2c). With larger sample sizes these interactions will be further resolved.

### Characterization of clonal dynamics in response to oncologic therapy

To characterize how treatment shapes the mutational landscape and clonal dynamics of CH, we collected sequential blood samples from 525 cancer patients (median sampling interval time = 23 months, range: 6-53 months) of whom 61% received cytotoxic therapy or external beam radiation therapy and 39% received either targeted/immunotherapy or were untreated (see Methods and Supplementary Figure 7 for more details on patient characteristics). Of these patients, 389 had CH at the time of first sampling and 136 did not. We observed 621 mutations of which the vast majority (95%, N=590) were detected at both timepoints.

We compared the change in VAF of CH clones across treatment modalities and in untreated patients and found evidence of both positive and negative changes in clone size (Fig. 3a). Among mutations detected at both timepoints, 62% (n = 367) of CH mutations remained stable, 28% (n = 164) had evidence of growth, and 10% (n = 59) decreased in clonal size between sampling timepoints based on a binomial test for difference in the VAF between timepoints given the sequencing depth. Among patients with multiple mutations, their mutations exhibited a higher growth rate^1^ as compared to those with one mutation (p = 0.03) irrespective of mutation type, and treatment status (Supplementary Fig. 8). This likely reflects the greater competitive advantage of a subset of clones harboring multiple mutations although this cannot be determined with certainty in the absence of single-cell sequencing. Among patients receiving external beam radiation therapy or cytotoxic therapy, the growth rate was most pronounced for CH with mutations in DDR genes as opposed to mutations in other CH genes such as *DNMT3A, ASXL1 or TET2* (Fig. 3b-c). Increasing cumulative exposure to cytotoxic therapy and external beam radiation therapy resulted in higher growth rates for CH with DDR mutations (Fig. 3d). Among mutations that were detected only at one timepoint (N=21), 6 were detected only at the initial timepoint and not at follow-up and 15 were detected only at the follow-up timepoint. We observed a non-significantly higher proportion of patients with newly detected mutations among those who received interval cytotoxic/radiation therapy (4%, n = 13) as compared to those who did not in a binomial test (1%, n=2, p=0.06) (Supplementary Fig. 9).

**Figure 3.**
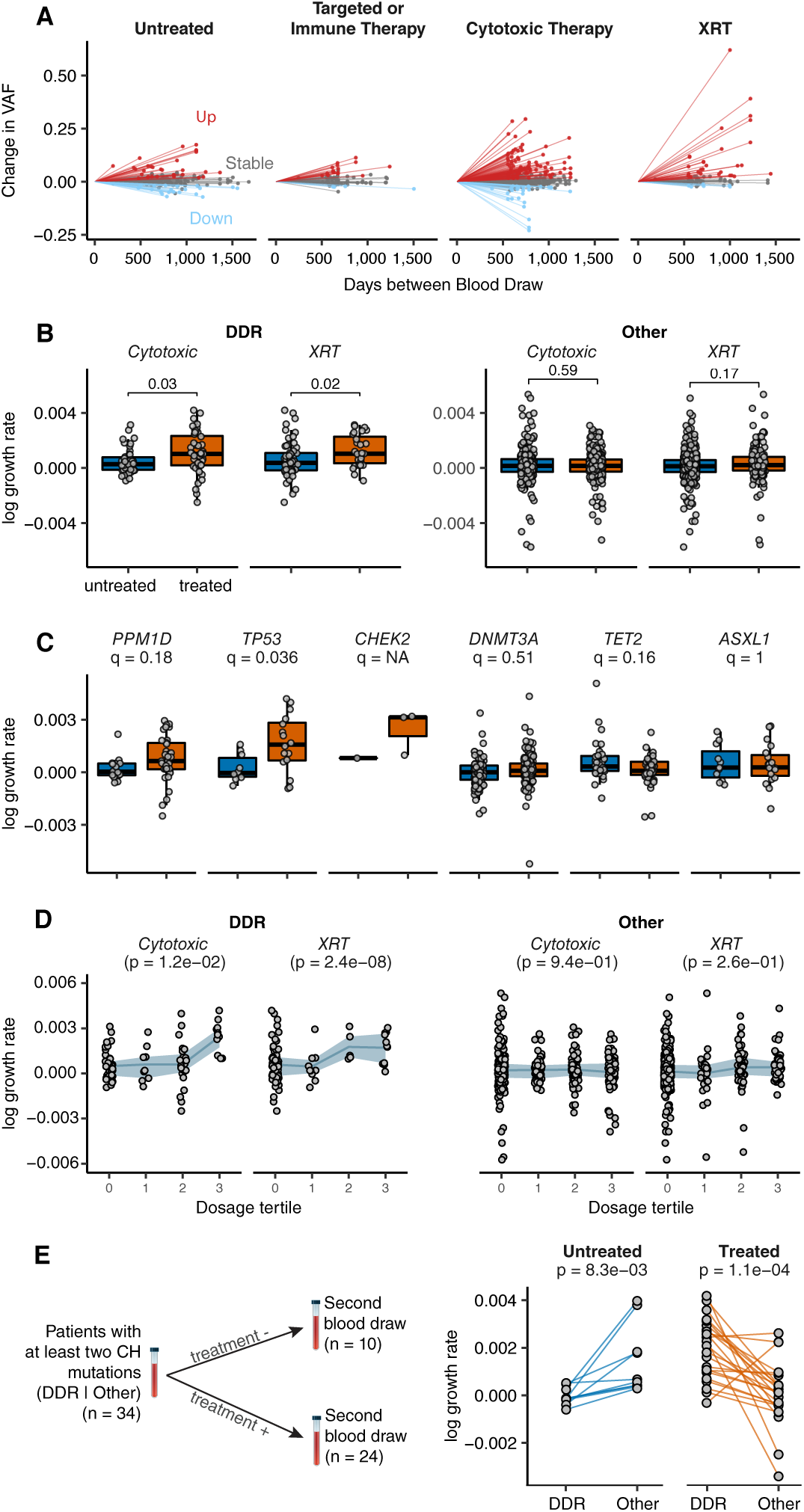
Clonal evolution of CH mutations under the selective pressure of oncologic therapy. (A) Change in VAF for CH mutations from initial sequencing to follow-up sequencing for patients stratified by the type of therapy received during the follow-up period. Shown are those who received cytotoxic therapy, those who received targeted/immune therapy alone, external beam radiation (XRT) alone and those who received no therapy during the follow-up period. (B) Change in the growth rate for DDR CH mutations and non-DDR CH mutations among those who received external beam radiation (XRT) or cytotoxic therapy during the follow-up period. The growth rate for mutations in treated patients are shown in orange and untreated in blue. (C) Change in the growth rate for specific CH mutations stratified by whether patients received cytotoxic therapy/radiation therapy (treated) or whether they received no therapy during the follow-up period. Shown are the FDR-corrected p-values (q-value). (D) Change in the growth rate for DDR CH mutations and CH due to mutations in other genes stratified by tertile of cumulative exposure to cytotoxic therapy and XRT. Shown are the p-values for a trend test for increasing growth rate of CH with increasing tertile of therapy exposure. E) Intra-subject competition between DDR and non-DDR-CH mutations. Among patients with at least one mutation in a DDR CH gene and a non-DDR CH gene we compared the difference in the growth rate between mutations. Connecting lines show the difference in growth rate between DDR vs other genes in patients who received XRT/Cytotoxic therapy (treated) during the follow-up period and in those who did not receive such therapy (untreated) during the follow-up period. A paired t-test was used to test for significance in the difference between growth rates within individuals.

We identified 34 subjects in our prospective serial sampling series with at least two CH mutations in which one mutation was in a DDR gene and one in a non-DDR gene. This offered the opportunity to study competing clonal dynamics for multiple gene mutations within the same patient. In patients receiving interval cytotoxic therapy or radiation therapy, CH clones with DDR mutations had a higher growth rate compared to clones with other CH mutations. However, the reverse was true in patients without interval exposure; clones with mutations in non-DDR CH genes (*e.g*. *DNMT3A*), outcompeted clones with DDR mutations (Fig. 3e). In summary, our serial sampling data provide clear evidence that oncologic therapy strongly selects for clones with mutations in the DDR genes *TP53, PPM1D* and *CHEK2* and that these clones have limited competitive fitness, in the absence of specific environmental factors such as cytotoxic or radiation therapy.

### Risk factors for tMN development in cancer patients

While CH is associated with an increased risk of tMN, the incidence of CH far exceeds that of tMN. To determine whether CH mutation type, number of CH mutations and/or clonal dominance were associated with tMN risk, we performed cause-specific Cox proportional hazards regression on 9,549 cancer patients exposed to oncologic therapy of whom 75 cases developed tMN (median time to transformation=26 months) (Supplementary Table 2). These data were drawn from our extended MSK cohort and from three previously published studies^17, 28, 29^ (see Supplementary Notes). The risk of tMN was positively associated with CH-PD (HR=6.9, p<10^-6^), and increased with the total number of mutations and clone size (Fig. 4a). The strongest associations were observed for mutations in *TP53* and for CH with mutations in spliceosome genes (*SRSF2*, *U2AF1* and *SF3B1*). Lower hemoglobin, lower platelet counts, lower neutrophil counts, higher red cell distribution width (RDW) and higher mean corpuscular volume (MCV) were all positively associated with increased tMN risk. We saw no significant heterogeneity between studies for the strength of the association between CH-PD and tMN. The number of CH mutations, *TP53*, *SF3B1* mutations and peripheral blood count parameters, specifically hemoglobin, platelet count, and RDW retained significance in multivariable model. Given that our estimates were derived from a cohort of cancer patients, we compared our findings to recent studies in healthy individuals^1, 30^ investigating CH as a risk factor for the development of AML. Comparison of HRs for tMN and AML risk showed similar effect sizes (Supplementary Fig. 10). These data suggest that the relative risk of myeloid neoplasms associated with CH and related parameters (gene, VAF and mutation number) are shared among healthy individuals and cancer patients.

**Figure 4.**
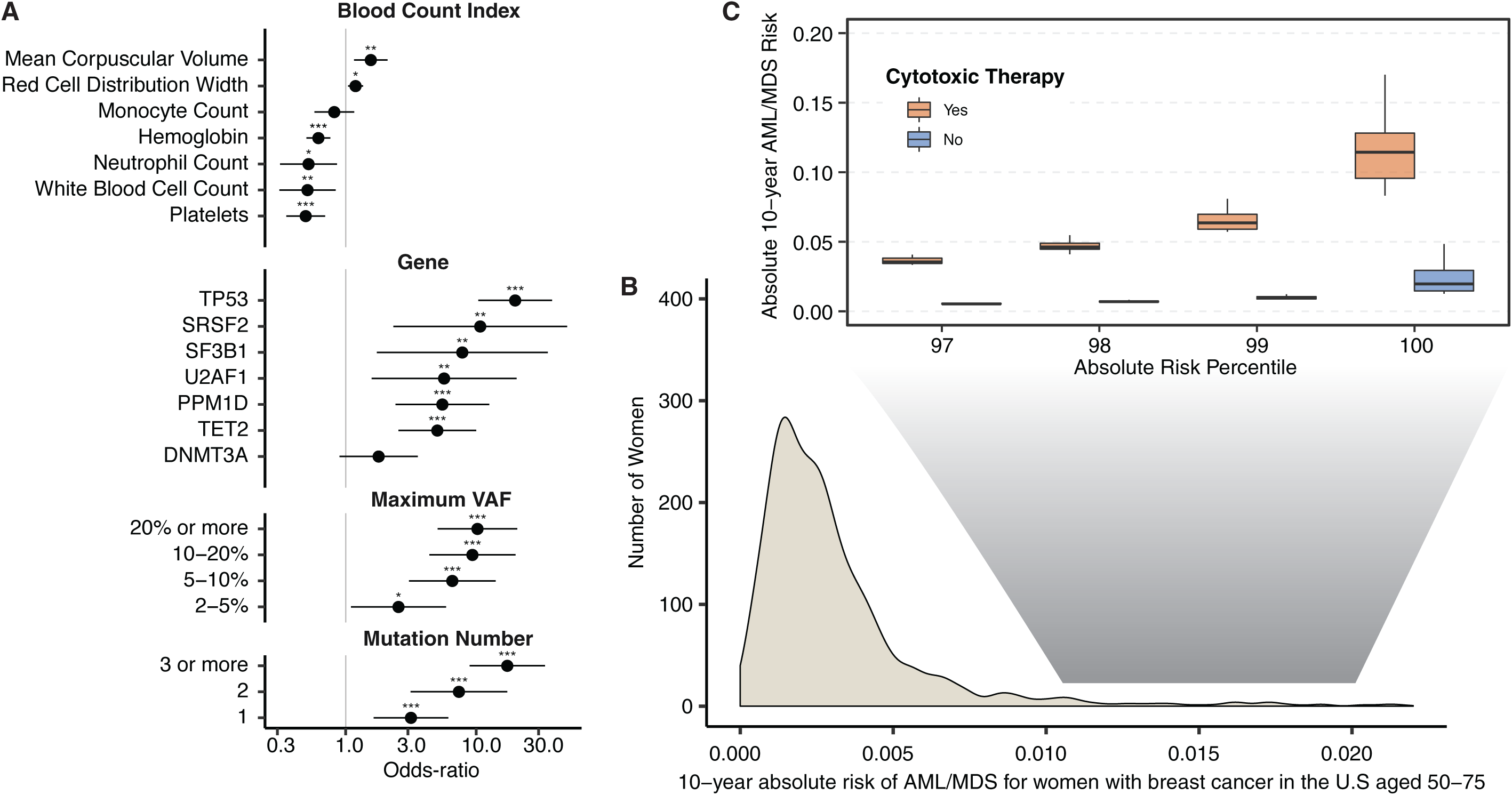
Risk of AML/MDS by clinical and mutational characteristics in solid tumor patients. (A) Hazard ratios from Cox regression for blood count indexes, and CH mutational characteristics for tMN in solid tumor patients. All models were adjusted for age and gender and stratified by study center. (B) Projected distribution of absolute 10-year risk of AML/MDS for women after a breast cancer diagnosis in the United States aged 50-75 at presentation C) Distribution of absolute 10-year risk of AML/MDS for among women at the top percentiles of risk for those who go on to receive adjuvant cytotoxic chemotherapy and those who receive surgery only.

### Relationship of clonal hematopoiesis to tMN development

Findings from our risk model suggest that there is a direct link between CH and tMN, whereby CH likely represents the precursor clone for tMN. To further compare the genetic and clonal relationships between tMN and the proceeding CH, we analyzed 35 cases for which paired samples were available with a median inter-sampling time of 24 months (5-90 months) (Supplementary Table 2). Targeted mutational analysis combined with clinical cytogenetics identified at least one disease defining event at the time of tMN diagnosis in 34 cases (97%). In 19 patients (59%), we found evidence of at least one of these mutations at the time of pre-tMN sequencing and in 13 (41%), we identified two or more in the pre-tMN sample. tMN transformation was frequently associated with acquisition of additional somatic mutations in cases who were CH positive at the time of pre-tMN sampling (Supplementary Fig. 11). In all cases the dominant clone at tMN transformation was defined by a mutation seen at CH (Extended Data Fig. 5).

Nearly half (n = 14, 40%) of the tMN patients had mutations in *TP53* at the time of tMN diagnosis, further validating the relevance of *TP53* mutations in tMN pathogenesis. *TP53* mutations were already present at time of CH in 10/14 patients. At tMN, *TP53* mutations co-occurred with isolated chromosomal aneuploidies or complex karyotype in 12/13 (92%) cases with available karyotype. In agreement with our prospective serial sequencing data, in the four cases with a *TP53* mutation and another non-DDR mutation in the pre-tMN sample that had ongoing exposure to cytotoxic therapy or radiation therapy, the *TP53* clone consistently outcompeted other mutations to attain dominance at the time of tMN transformation (Extended Data Fig. 5). In the cases where we did not identify mutations at time of pre-tMN sequencing (N=16), the tMN sample was commonly defined by chromosomal aneuploidies or MLL fusions common to myeloid neoplasm pathogenesis (Supplementary Fig. 11). Additional pathways to tMN pathogenesis through structural variation, including fusions of the *MLL* gene and aneuploidies of chromosome 5 and 7^7, 13^ would not have been captured by our study.

### Implications of tMN risk stratification in tailoring treatment decisions

Taken together our findings further refine the relevance of CH as a biomarker for tMN risk. We sought to evaluate how CH, in combination with other clinical parameters such as age and peripheral blood counts, might provide clinically meaningful tMN risk stratification. For example, in solid tumor patients undergoing surgical resection, adjuvant oncologic therapy can improve overall survival by reducing cancer recurrence. However, in some situations, the absolute survival benefit of adjuvant therapy is modest and is countered, at least in part, by the risk for subsequent tMN. In the absence of prospective clinical studies, we performed an exploratory analysis using a synthetic model to quantify the absolute risk of AML/MDS following a breast cancer diagnosis. Using previously established methodology^31, 32^ (iCARE), we combined estimates of HR parameters obtained from our multivariable analysis, the distribution of CH mutational features and blood count parameters from untreated patients at MSK aged 50-75 and external data sources for 1) age-specific AML/MDS rates in breast cancer, 2) competing hazards for mortality in breast cancer, 3) the relative risk for tMN conferred by adjuvant chemotherapy compared to surgery alone in breast cancer and 4) the proportion of women with breast cancer who received adjuvant chemotherapy.

Using these parameters we modeled the 10-year cumulative AML/MDS absolute risk distribution for women with breast cancer aged 50-75 in the United States. We estimated how the risk distribution would change with receipt of adjuvant therapy by shifting the population between receiving and not receiving adjuvant chemotherapy. Our estimates showed that the majority (96%) of breast cancer patients have a low 10-year absolute risk (AR < 1%) for tMN (Fig. 4b) and for these patients, deferment of adjuvant chemotherapy would not impact their tMN absolute risk (Fig. 4c). However, for the women at the highest risk of tMN (top 1%), adjuvant chemotherapy was estimated to increase tMN absolute risk by over 9%. This would exceed the predicted absolute benefit in overall survival of chemotherapy in many women with early stage breast cancer ^33^. While not suitable for immediate clinical application, this provides proof-of-concept that incorporating CH mutational and clinical characteristics could achieve adequate tMN risk stratification to influence clinical decision making. However, prospective studies are needed to develop evidence-based guidelines for clinical practice.

## DISCUSSION

Cancer develops as a result of the successive acquisition of genomic alterations that contribute to a cell population’s fitness advantage, selection and malignant potential in a process that is often likened to Darwinian evolution^34,35,36,37^. To this end, estimates of tumor phylogenies are frequently modelled on the basis of the observed genetic and clonal structures at the time of cancer diagnosis ^38–41^. Such approaches do not offer the possibility to study the dynamics of early clones that may never progress into a tumor or allow for an evaluation of the factors that can shape the fitness of specific gene mutations under diverse environmental exposures. We show that the fate of CH mutations is dictated by a complex interplay between the inherent fitness advantage of the mutation(s) in HSPC’s and cell extrinsic features that preferentially select for specific mutations, i.e. oncologic therapy for genes involved in DDR, aging for spliceosome mutations, and smoking for *ASXL1* mutations (Extended Data Fig. 6).

Previously, CH has been shown to demarcate patient subgroups with adverse clinical outcomes^7^. We show that in cancer patients CH is strongly associated with older age, smoking history and exposure to oncologic therapy, highlighting that these factors should be considered in future correlative studies of CH in cancer cohorts. We refine the specificity and strength of the association between oncologic therapy and CH and extend this to characterize the relationship at a gene and treatment-specific resolution. We show that radiation therapy and cytotoxic therapy are associated with CH, with regimens containing platinum and topoisomerase II inhibitors most strongly correlating with CH in DDR pathway genes. The strong dose-response relationships observed in our study provide further supporting evidence of a causal relationship. Serial sampling of patients before and after therapy provide clear, definitive evidence that therapy induces gene-specific clonal expansion, whereby clones with mutations in DDR genes outcompete other clones and attain clonal dominance in the setting of oncologic therapy, but not in its absence. We further validate the relevance of CH as a predictor and precursor of tMN in cancer patients. We show that CH mutations detected prior to tMN diagnosis were consistently part of the dominant clone at tMN diagnosis and demonstrate that oncologic therapy directly promotes clones with mutations in genes associated with chemo-resistant disease such as *TP53*. This demonstrates, for the first time, a causal relationship between genetic subtypes of CH, subsequent oncologic therapy, and clonal expansion of CH to tMN.

The realization of precision medicine is reliant upon the development of evidenced-based guidelines that consider molecular biomarkers alongside standard clinical criteria to inform clinical care. Here we show that prospective clinical sequencing can identify patients with CH, and that this can be used to predict risk of subsequent tMN. Moreover, our observations pave the way to using this data for therapeutic intervention, including the development of therapies aimed to target high risk CH clones and modulating the use of adjuvant systemic cancer therapy in patients at highest risk of subsequent myeloid neoplasm. The decreasing cost of prospective clinical sequencing assays and the high frequency of CH in cancer patients suggest that screening for CH prior to initiation of oncologic therapy may be feasible, and may represent an avenue for molecularly based early detection and interception.

## METHODS

### Sample Ascertainment and clinical data extraction

#### MSK-IMPACT Cohort

The study population included patients with non-hematologic cancers at MSKCC that underwent matched tumor and blood sequencing using the MSK-IMPACT panel on an institutional prospective tumor sequencing protocol <u>ClinicalTrials.gov</u> number, NCT01775072 before July 1st 2018. We extracted data on race, smoking, date of birth and cancer history through the MSK cancer registry. Subjects who had a hematologic malignancy within three years of blood collection for MSK-IMPACT testing or who had an active hematologic malignancy at the time of blood draw were excluded. When unavailable through the cancer registry, we extracted data on race and smoking through structured fields in clinician medical notes if available. Subjects for which age was not available were excluded. Blood indices were taken from clinical labs closest to the date of blood collection for MSK-IMPACT, within one year before or after blood collection (median=0 days).

#### Serial Sampling Cohort

In order to study the growth rate of clonal hematopoiesis mutations over time we collected additional blood samples on patients sequenced using MSK-IMPACT for repeat CH mutation testing. These came from three sources: 1) 372 patients with CH where we obtained second blood sample at least 18 months after initial MSK-IMPACT blood collection, 2) 21 samples from patients with clonal hematopoiesis on MSK-IMPACT who had a blood sample banked at least 12 months prior to MSK-IMPACT testing and 3) 132 samples that were taken for repeat MSK-IMPACT testing for clinical purposes at least six months after the first MSK-IMPACT testing irrespective of clonal hematopoiesis status (Supplementary Fig. 7). For all patients who had sequential sampling data, we manually reviewed their medical records to capture receipt of oncologic therapy received at outside institutions during the follow-up period. If subjects received therapy outside MSK during the follow-up period, we excluded them from analyses of dose-response relationships since cumulative dose of therapy could not be consistently collected from outside records. This study was approved by the MSKCC Institutional Review Board.

#### Targeted Capture-Based Sequencing

Subjects had a tumor and blood sample (as a matched normal) sequenced using MSK-IMPACT, a FDA-authorized hybridization capture-based next-generation sequencing assay encompassing all protein-coding exons of 341, 410, or 468 cancer-associated genes (Supplementary Table 3). MSK-IMPACT is validated and approved for clinical use by New York State Department of Health Clinical Laboratory Evaluation Program and is used to sequence cancer patients at Memorial Sloan Kettering. Genomic DNA is extracted from de-paraffinized formalin fixed paraffin embedded (FFPE) tumor tissue and patient matched blood sample, sheared and DNA fragments were captured using custom probes^44^.

The blood samples in the serial sampling cohort that were obtained for repeat CH testing were sequenced using a comparable capture-based custom panel using 163 genes implicated in myeloid pathogenesis, which included the most commonly mutated genes in our MSK-IMPACT study, with the exception of ATM. The median sequencing depth was 665X (range=111-1987X) which was comparable to that obtained in the blood using MSK-IMPACT. For all subsequent analyses using the serial sampling cohort we only considered mutations that were present in both the initial and follow-up panel.

#### Variant Calling

Pooled libraries were sequenced on an Illumina HiSeq 2500 with 2×100bp paired-end reads. Sequencing reads were aligned to human genome (hg19) using BWA (0.7.5a). Reads were re-aligned around indels using ABRA (0.92), followed by base quality score recalibration with Genome Analysis Toolkit (GATK) (3.3-0). Median coverage in the blood samples was 497x, and median coverage in the tumors was 790x. Variant calling for each blood sample was performed unmatched, using a pooled control sample of DNA from 10 unrelated individuals as a comparator. Single nucleotide variants (SNVs) were called using Mutect and VarDict. Insertions and deletions were called using Somatic Indel Detector (SID) and VarDict. Variants that were called by two callers were retained. Dinucleotide substitution variants (DNVs) were detected by VarDict and retained if any base overlapped a SNV called by Mutect. All called mutations were genotyped in the patient matched tumor sample. Mutations were annotated with VEP(version 86) and OncoKb.

#### Post-Processing Filters for Clonal Hematopoiesis Calling

We applied a series of post-processing filters to further remove false positive variants caused by sequencing artifacts and putative germline polymorphisms. We removed variants that were found (with a VAF of >2% at least once) in a panel of sequencing data from 300 blood samples obtained from persons under 20 years of age and without evidence of clonal hematopoiesis. We further filtered single nucleotide deletions within a homopolymer stretch of (>=3 base repetition) of the same deleted base pair, single nucleotide substitutions completing a stretch of a >= 5bp-long homopolymer (E.g. GGCGG -> GGGGG) in-frame deletions or insertions in a highly repetitive region (DUST^45^ algorithm score of >=5), and variants with unequal proportions of forward/reverse direction supporting reads based on a fisher test. We performed manual review in IGV of recurrent mutations not previously reported in public databases. We required a variant allele fraction of at least 2% and at least 10 supporting reads. All genotypes were calculated using sequencing reads and bases with a quality value of at least 20. Because somatic mutations in the blood would be expected to be detected in the blood but not other tissue compartments, we compared the variant allele fraction (VAF) of mutations in the blood compared to the matched tumor. Variant calls that were present in the blood with a VAF of at least twice that in the tumor or 1.5 times the VAF if the tumor biopsy site was a lymph node were considered somatic. This ratio was chosen based on minimizing sensitivity and specificity of CH calls through simulations of leukocyte contamination in the tumor (see Supplementary Notes and Supplementary Figures 10 and 11). To further filter putative germline polymorphisms that passed the blood/tumor solid tissue ratio due to allelic imbalance in the tumor specimen, we removed any variant reported in any population in the gnomAD database at a frequency greater than 0.005.

#### Validation of Calls

To test the reproducibility of our clonal hematopoiesis mutation calling, we compared the mutational calling results from 1,173 samples, where the same DNA library for a blood sample was sequenced and analyzed twice using MSK-IMPACT. We detected 91% of variants in both samples using our calling criteria with a correlation coefficient of 0.98 for the variant allele fraction between the two calls indicating that the reproducibility of our calls was high. In 10 cases with CH, we obtained a second blood sample and re-sequenced using a custom capture based panel with unique molecular identifiers and found that this independent method confirmed all 18 of our CH calls using MSK-IMPACT.

#### Variant Annotation

Variants were annotated according to evidence for functional relevance in cancer (putative driver or CH-PD) and for relevance to myeloid neoplasms specifically (CH-Myeloid-PD). We annotated variants as oncogenic in myeloid disease (CH-Myeloid-PD) if they fulfilled any of the following criteria:

1. Mutation in a putative myeloid driver gene (Supplementary Table 4)
2. Truncating variants in *NF1, DNMT3A, TET2, IKZF1, RAD21, WT1, KMT2D, SH2B3, TP53, CEBPA, ASXL1, RUNX1, BCOR, KDM6A, STAG2, PHF6, KMT2C, PPM1D, ATM, ARID1A, ARID2, ASXL2, CHEK2, CREBBP, ETV6, EZH2, FBXW7, MGA, MPL, RB1,SETD2, SUZ12, ZRSR2* or in *CALR* exon 9
3. Translation start site mutations in *SH2B3*
4. *TERT* promoter mutations
5. *FLT3*-ITDs
6. In-frame indels in *CALR, CEBPA, CHEK2, ETV6, EZH2*
7. Any variant occurring in the COSMIC “haematopoietic and lymphoid” category greater than or equal to 10 times
8. Any variant noted as potentially oncogenic in an in-house dataset of 7,000 individuals with myeloid neoplasm greater than or equal to 5 times

We annotated variants as oncogenic (CH-PD) if they fulfilled any of the following criteria:

1. Any variant noted as oncogenic or likely oncogenic in OncoKB^19^
2. Any truncating mutations (nonsense, essential splice site or frameshift indel) in known tumor suppressor genes as per the Cancer Gene Census or OncoKB. Genes not listed in the cancer census or OncoKB were reviewed in the literature to determine if they were potentially tumor suppressor genes.
3. Any variant reported as somatic at least 20 times in COSMIC^46^
4. Any variant meeting criteria for CH-Myeloid-PD as above.

All missense variants not meeting the above were individually reviewed for potential oncogenicity as previously described^47^.

#### Statistical Methods

##### dN/dS

We used the dNdScv (https://github.com/im3sanger/dndscv) package to quantify the dN/dS ratios for missense and truncating mutations at the gene level as well as on the panel level. Due to the difference in the gene panel between different MSK-IMPACT panel versions, we excluded all MSK-IMPACT-341 samples and only included genes that were present on both MSK-IMPACT-410 and MSK-IMPACT-468 panels in the analysis. Finally, to generate the overall dN/dS landscape in CH, we only presented genes that reached a significance level of q < 0.1 after multiple testing correction and contained more than 25 variants.

#### Modeling the association between CH and prior exposure to oncologic therapy

We used multivariable logistic regression to evaluate for an association between clonal hematopoiesis (including gene and variant specific factors) and therapy, age, gender and smoking history. In addition to these variables, we also adjusted for time from cancer diagnosis to blood draw for MSK-IMPACT testing because trends in preferred oncologic agents vary over time and CH is known to associate with survival. We did not adjust for primary tumor type since we hypothesized that most of the difference in CH-PD rates reflected differences in oncologic regimens. Indeed, among untreated patients, a global Wald test for differences in CH-PD prevalence by tumor type was not significant (p=0.98). Analyses stratified by the time since start and by completion of external beam radiation and chemotherapy showed no clear evidence of a time-dependence/latency between CH-PD and cumulative exposure to therapy. Thus, the time from start or stop of therapy was not adjusted for. While considering exploratory analyses, we performed multiple hypothesis correction using the false discovery rate (FDR) q-values for gene-specific analyses to control for inflation of type I error. We did not perform multiple hypothesis correction for analyses testing an association between subclasses of oncologic therapy and CH because the association between oncologic therapy and CH is known and our goal was to define the relative strength of these associations with subtypes of therapy rather than hypothesis testing. Heterogeneity p-values to test for differences in the strength of the association between subclasses of CH and clinical variables were calculated through logistic regression models limited to CH-positive individuals testing for a difference in the odds of having CH with the mutational feature of interest (*e.g*. CH-PD) vs. having CH without the mutational feature (*e.g*. non-CH-PD). Generalized estimating equations were used to test for an association between CH VAF and selected clinical and mutational features among CH positive individuals accounting for correlation between the VAF of mutations in the same person. Ordinal logistic regression among CH positive individuals was used to test for an association between clinical characteristics and increasing CH mutation number. A test for trend between increasing cumulative exposure to oncologic therapy and the odds of CH-PD was performed using multivariable logistic regression limited to individuals exposed to the therapy of interest.

#### Modeling the effect of oncologic therapy on mutation growth rate

For each mutation in each individual with sequential sequencing data available, we modeled the growth rate of the mutation between the two time points according to the following formula: α = log (V / V_0_) / (T - T_0_)

Where T and T_0_ indicates the age of the individual (in days) at the two measurement time points and V and V_0_ correspond to the VAF at T and T_0_ respectively. We also classified mutations as having increased, decreased or remained constant during the follow-up period based on a binomial test comparing the two VAFs. Generalized estimating equations were used to test for an association between exposure to cytotoxic therapy and external beam radiation therapy and CH growth rate adjusting for age, gender and smoking status accounting for correlation between the growth rate of mutations in the same person. Among patients with at least one mutation in a DDR CH gene and another non-DDR CH gene (N=34), we calculated the difference in the growth rate between mutations. When patients had more than two mutations in the same gene category, we used the highest growth rate for that category. A paired t-test was used to test for significance in the difference between growth rates of DDR mutations compared to non-DDR mutations within individuals who received cytotoxic therapy and/or external beam radiation therapy and within those who were untreated during the follow-up period.

#### Combined Analysis for AML/MDS Risk

We combined data from three previously published studies, Gillis et al., abbreviated MOF (N=68), Takahasi et al., abbreviated MDA (N=67), Gibson et al., abbreviated DFC (N=401) studying the effect of CH on tMN risk in cancer patients. For all samples, uniform post processing filters were applied to ensure retention of variants in accordance to the QC standards of the MSK cohort including a universal 2% minimum VAF cutoff. We only included mutations within genes that are present on the panel from all centers and on all panel versions from each center. The only exceptions were *SRSF2* which the IMPACT-341 sequencing panel did not cover and *PPM1D* which was not sequenced in IMPACT-341, MDA or MOF. We performed mean imputation of missing clinical data for blood counts. Only mutations that we classified as CH-PD were included in analyses. We performed univariate cause-specific Cox proportional hazards regression for the effect of maximum VAF, total number of CH mutations, CH in specific genes and blood count parameters adjusted for age and gender and stratified by study site. Interaction terms between study and CH were used to test for heterogeneity between studies on the effect of CH on tMN risk. The proportional hazards assumption was tested through visual inspection of residual plots and through the inclusion of time-varying covariates. We performed a multivariable analysis including age, gender and all variables that were significant in the univariate analysis with the exception of the genes not included in all studies to prevent reduction of sample size, *PPM1D* and *SRSF2*.

We also combined data from two studies investigating the effect of CH on AML risk in healthy individuals, Abelson et al., abbreviated PMC (N=969) and Young et al., abbreviated WSU (N=103), with data from MSK and applied uniform processing to mutation data from different centers. As in the solid tumor combined analysis, the same post processing filters used in the main MSK cohort including a universal 2% minimum VAF cutoff were applied to these studies and only mutations that we classified as CH-PD were included in analyses. We performed a multivariable Cox regression adjusted for age and gender including the variables used in the multivariable tMN risk analysis in solid tumor patients.

#### Modeling absolute risk of AML/MDS

We used the iCARE software package^31, 32^ to build a model for absolute risk of AML/MDS in women with breast cancer aged 50-75 in the United States (U.S) by combining 1) the multivariate HR estimates from our study that were significant in the univariate model including maximum VAF of CH, gene specific effects and peripheral blood count indexes (RDW, hemoglobin) 2. Age-specific AML/MDS rates in breast cancer using data provided by the National Comprehensive Cancer Network (NCCN)^49^; 3. Competing hazards for mortality in women with breast cancer in the U.S aged 50-75 as reported in SEER^50^; 4. Previously published HR estimates for chemotherapy on the risk of tMN in women with breast cancer from the NCCN^49^; 5. The distribution of CH VAF, number of mutations, CH gene and peripheral blood count indexes using our cohort of MSK solid tumor cancer patients aged 50-75 who were untreated prior to blood draw and 6. The proportion of women who receive adjuvant chemotherapy for breast cancer in the U.S from SEER^50^. While our IMPACT cohort is not representative of the general breast cancer population in the U.S, since the distribution of CH mutational features is largely driven by age and since we do not see major differences in rates of CH between gender or untreated tumor types, we believe that the distribution of CH mutational features in untreated solid tumor patients sequenced on IMPACT reasonably approximates an age-matched untreated breast cancer population. While blood count indexes are known to differ by sex and we chose to use the distribution of blood counts from the entire treatment-naive IMPACT population (both male and female) to capture the inter-relationship between blood count indexes and CH mutational features. Sensitivity analyses using the distribution of blood count parameters from female IMPACT patients only produced similar results. This risk model assumes an additive association on the log scale of CH mutational features and oncologic therapy for risk of tMN. This assumption is supported by the similarity between risk estimates for CH mutational features between AML in healthy individuals never exposed to therapy and tMN (Supplementary Fig. 10).

All the statistical analyses were performed with the use of the R statistical package (www.r-project.org). The code used in statistical analysis is provided in the Supplementary Appendix.

## Supporting information

R code for results

## ACKNOWLEDGEMENTS

This work was supported by the National Institute of Health (K08CA241318 to K.B., P30 CA008748 to S.D., K12-CA120780 to C.C., P30 CA008748 to M.R., P50 CA172012 to L.B., P50 CA172012 to J.F., UG1-HL069315 to V.K.), American Society of Hematology (K.B. and E.P.), EvansMDS Foundation (K.B.), European Hematology Association (E.P.), Gabrielle’s Angels Foundation (E.P.), V Foundation (E.P.), Geoffrey Beene Foundation (E.P), UNC Oncology Clinical Translational Research Training Program (C.C), Cycle for Survival (V.K.), Starr Cancer Consortium (to R.L., A.Z., M.B, R.P.), and the Cancer Colorectal Cancer Dream Team Translational Research Grant (SU2C-AACR-DT22-17 to L.D.). E.P. is a Josie Robertson Investigator. C.C. is a recipient of the Conquer Cancer Foundation Young Investigator Award and the Prostate Cancer Foundation Young Investigator Award. K.S. is a recipient of the Defense Early Investigator Research Award (W81XWH-18-1-0330), Prostate Cancer Foundation Young Investigator Award and the Prostate Cancer Foundation Challenge Award. C.L, M.G and L.M are supported by funds from the Intramural Research Program of the National Cancer Institute, National Institutes of Health. The University of Cambridge has received salary support in respect of PDPP from the NHS in the East of England through the Clinical Academic Reserve.

## AUTHOR CONTRIBUTIONS

K.B, R.L, A.Z and E.P conceived and designed the study. K.B, L.B, D.K, M.P, A.P and N.C performed collection of clinical data. R.P, A.S, M.B, A.Z, R.B., M.E.A., and M.L. led the generation of IMPACT sequencing data. K.B, M.P, A.P, N.C, D.H, M.T and R.L performed collection of sequential samples. R.P, T.G and K.B performed variant calling and post-processing of sequencing data. K.B, T.G, S.D, M.G, N.C, L.M, A.Z and E.P performed statistical analyses and/or participated in data interpretation. All authors contributed to the writing of the manuscript and approved it for submission. R.P and T.G contributed equally to the work. A.Z. and E.P. are shared senior authors.

## Supplementary Notes for “Oncologic therapy shapes the fitness landscape of clonal hematopoiesis”

### Clinical data ascertainment in MSK solid tumor patients

#### Chemotherapy

Data on chemotherapy were taken from two different sources. Prior to January 1st of 2011, data were extracted from pharmacy dispensing records. These data were accurate for outpatient treatments since 1992 and inpatient treatments since 1994. After January 1st of 2011, these data were taken from completed orders for chemotherapy in the electronic medication administration reconciliation system (EMar). Oral oncologic therapy prescribed through outside pharmacies was not uniformly captured in our cohort. For this reason, we did not attempt to study exposure to hormonal therapy since this is often given orally and filled at outside pharmacies. When chemotherapy is ordered in the EMar at MSK, this is done using a set of orders (orderset) that frequently contain multiple oncologic agents (i.e. the orderset named “FOLFOX” prescribes for Leucovorin, Fluorouracil, Oxaliplatin). Clinicians may choose to drop or add drugs to the order set and this is given as an option when placing chemotherapy orders. The order set used to prescribe chemotherapy was available for orders placed after January 1st of 2011. Among patients who only received chemotherapy after January 1st of 2011, we defined exposure to a drug regimen as the combination of drugs ordered using a given order set for an individual.

Patients in our main MSK cohort received 492 unique cancer-directed systemic therapies. In order to achieve adequate power to characterize the relative strength of associations between oncologic therapy and CH, we grouped systemic therapies according to their primary mechanism of action. “Goodman & Gilman’s The Pharmacologic Basis of Therapeutics”^42^ was primarily used for drug classification, supplemented by literature review for agents not described in this text. We classified agents as “cytotoxic” if they were traditional, non-specific, cytotoxic therapies. Drugs that act primarily through pathway-specific mechanisms (i.e. monoclonal antibodies and protein kinase inhibitors) were classified as “targeted” therapies. Systemically administered radionuclides not conjugated to antibodies (e.g. ^131^I, ^223^Ra, etc.) were classified as “radionuclide therapy” while antibody-conjugated radionuclides were grouped with targeted agents since they were considered to have target-localized effects. Agents that act primarily through modulation of the immune system were classified as “immune therapies”. Drugs that inhibit or modulate hormonal pathways were classified as “hormonal therapies” and were not included in our analysis of cancer-directed therapy (e.g. patients who only received hormonal therapies were considered to be treatment naive). This was done for two reasons; first, because there is a paucity of evidence linking receipt of hormonal therapy with risk of therapy-related leukemia and second, receipt of oral hormonal therapy was not well captured in our study. Cytotoxic drugs were further classified according to mechanism of action including alkylating agents, platinum complexes, antimetabolites, topoisomerase I inhibitors, topoisomerase II inhibitors, microtubule damaging agents, etc. Because microtubule damaging agents encompass a large number of agents, these were further classified into “taxanes” and “other microtubule damaging agents”. Cytotoxic agent groupings that contained fewer than 20 individuals were combined into “other cytotoxic agents”. Supplementary Table 1 shows the final classification of agents in this study.

#### Radiation therapy

Dosimetric parameters for each course of external beam radiation (i.e. dose, fractionation, technique/modality, target) were extracted from the treatment planning system (ARIA; Varian Medical Systems, Palo Alto, CA), from radiotherapeutic prescriptions and from clinical treatment summaries. For patients treated prior to the implementation of contemporary treatment planning (prior to 2003), receipt and timing of radiation therapy was abstracted from medical billing records, although dose and fractionation were not available for these patients and were set as missing.

#### Measurement of cumulative therapy dose

Given the variety of radiotherapy fractionation schemes and prescribed tumor doses, we calculated the cumulative radiation dose received by each patient prior to blood draw in 2-Gy per fraction equivalents (EQD_2_) using an α/β of 3 Gy, considering CH to be a late-responding tissue effect^43^. We calculated tertiles of dose based on the distribution of cumulative EQD_2_ received over the entire cohort and assigned each individual a score based on their tertile of exposure (*e.g*. a patient who did not receive external beam radiation received a score of zero for that particular agent. If the patient’s cumulative radiation dose, as expressed in EQD_2_, was within the first tertile, a score of one was assigned, and so forth).

To derive metrics for cumulative exposure to cytotoxic therapy subclasses, we applied the approach used by the Late Effects Study Group ^25^. For each drug the total dose per kg received prior to blood draw was summed for each patient. The dose distribution for each agent was divided into tertiles and the patient’s dose was assigned a score based on tertile of total exposure. An individual patient’s scores for each drug in a specific drug class were summed. The distribution of the resulting sum across all patients was used to derive tertiles of total exposure to the drug class in the entire cohort (Supplementary Fig. 6).

#### Development of secondary myeloid disease

We used a combination of sources to identify incident hematologic malignancies following blood collection for MSK-IMPACT testing including: 1) the MSK cancer registry for any listed diagnoses of hematologic neoplasms, 2) visits to an oncologist within the leukemia service, 3) pathology reports describing bone marrow biopsies, 4) billing codes relating to hematologic neoplasms, and 5) a free-text search of all EMR documents for leukemia or MDS-related search terms. We reviewed the medical records individual for any patient who was selected as a possible case using the above criteria. If a patient was diagnosed within six months of blood collection for MSK-IMPACT testing, they were considered to have active disease at the time of testing and were excluded. We defined the date of last follow-up as the last visit to MSK, the last date of phone/email contact with the patient, or the date of death as per the Social Security Death Index or death notification from family members or outside institutions.

### Clinical characteristics of previously published studies included in combined analyses

To study the relationships between CH and tMN we aggregated data from 5 previously published studies to include Gillis et al, (MOF) Takahshi et al (MDA), Gibson et al. (DFC), Young et al., (WSU) and Abelson et al., (EPI).

#### MOF

Gillis et al.,^1^ performed a nested case-control study for tMN risk using subjects from an internal biorepository of 123,357 cancer patients who consented to participate in the Total Cancer Care biobanking protocol at Moffitt Cancer Center (Tampa, FL, USA) between Jan 1, 2006, and June 1, 2016. Cases were individuals diagnosed with a primary malignancy, treated with chemotherapy who subsequently developed a therapy-related myeloid neoplasm, and were 70 years or older at either diagnosis. Controls were individuals who were diagnosed with a primary malignancy at age 70 years or older and were treated with chemotherapy but did not develop therapy-related myeloid neoplasms. Controls were matched to cases in at least a 4:1 ratio on the basis of sex, primary tumour type, age at diagnosis, smoking status, chemotherapy drug class, and duration of follow-up. DNA was isolated from peripheral blood collected before therapy-related myeloid neoplasm diagnosis and subjected to Droplet-partitioned, targeted, amplicon-based, next-generation sequencing was used in accordance with the manufacturer’s instructions (RainDance Technologies, Billerica, MA, USA) to identify somatic mutations in 49 myeloid-driver genes (ThunderBolts Myeloid Panel, RainDance, Billerica, MA, USA).

#### MDA

Takahasi et al.^2^ performed a case-control study for cancer patients who developed therapy-related myeloid neoplasms (cases) and lymphoma patients who did not develop therapy-related myeloid neoplasms (controls). Cases were identified using a clinical database at the Department of Leukemia of The University of Texas MD Anderson Cancer Center (Houston, TX, USA) including 40,000 patients who have consented for their data to be used in research. Inclusion criteria were that patients had to have been treated for a primary cancer from June 11, 1997, and subsequently had diagnoses of therapy-related myeloid neoplasms between Jan 1, 2003, and Dec 31, 2015, and had paired samples of diagnostic bone marrow at the time of therapy-related myeloid neoplasm diagnosis and peripheral blood samples obtained at the time of primary cancer diagnosis. An aged-matched control group (using a 3:1 control to case ratio) was identified using a clinical database of patients treated for lymphoma from 2008 to 2015. Eligible patients were those who had a pre-treatment blood sample available, had received a combination chemotherapy regimen including an alkylating agent, had at least 5 years of follow-up with no clinical evidence of therapy-related myeloid neoplasm development, and had no evidence of bone marrow metastasis of lymphoma in a bilateral bone marrow biopsy. Targeted sequencing of 32 myeloid genes was performed using an amplicon-based targeted deep sequencing method, including unique molecular barcodes.

#### DFC

Gibson et al.,^3^ performed a cohort study among 401 adult patients who underwent ASCT for non-Hodgkin lymphoma between 2003 and 2010 (Dana-Farber Cancer Institute, Boston, MA; targeted sequencing cohort) with mobilized stem-cell products available at the time of ASCT. All subjects had been exposed to oncologic therapy prior to stem cell collection. During the follow-up period 18 patients developed tMN. Samples were obtained from mobilized stem-cell products at the time of ASCT. Targeted deep sequencing was performed using 86 known myeloid genes using the Custom SureSelect hybrid capture system (Agilent Technologies, Santa Clara, CA).

#### WSU

Young et al.^4^, utilized a nested case-control design for AML using data from two large cohort studies, the Nurses Health Study (NHS) and the Health Professionals Follow-Up Study. Subjects were drawn from the “blood sub-cohorts” of these two studies which included 32,826 women (NHS) with a blood sample from 1989-90 as well as 18,018 men (HPFS) who provided a whole blood sample from 1993-95. The case definition included all blood sub-cohort participants with confirmed diagnoses of AML occurring after blood draw. Two matched controls were selected per case on cohort (sex), race, birthdate (± 1 year), and blood draw details (date ± 1 year, time ± 4 hours, fasting status <8, 8+ hours). Samples were sequenced using the Illumina TruSight Myeloid Sequencing Panel for targeted capture from 54 leukemia-associated genes Technical replicate libraries were sequenced on different machine runs. Error corrected sequencing analysis of raw sequencing results was performed as described elsewhere^5^.

#### EPI

Abelson et al.^6^, performed a case-control study for AML using samples from EPIC ^7^. We used data from both the discovery and validation sets. The discovery set included individuals who enrolled on the EPIC study between 1993 and 1998 across 17 different centres. The validation cohort included individuals enrolled in the EPIC-Norfolk longitudinal cohort study between 1994 and 2010. Subjects who developed AML during the follow-up period were matched to age- and gender-matched controls without a history of cancer or any hematological conditions. Targeted deep sequencing in the discovery cohort was performed using error-corrected, custom capture based sequencing using the xGen AML Cancer Panel. Targeted sequencing in the validation set was performed using a custom complementary RNA bait set (SureSelect, Agilent, ELID: 0537771) designed complementary to all coding exons of 111 myeloid driver-genes.

### Eliminating germline events and technical artifacts using tumor comparator

Using a synthetic dataset, we profiled the error rates of several methods that use the matched tumor as a comparator to eliminate germline events and false positive calls (artifacts). We simulated pairs of observed variant allele fractions in the blood and the tumor as follows:

Let *f*_b_ be the true variant allele fraction in blood, *f*_t_ be the true variant allele fraction in blood and *c* be the level of blood contamination in the tumor and *r* ∈ { 0,1} be an indicator for whether the variant is real (*r*=1) or artifact (*r* = 0). Let *v_b_* be the observed VAF in the blood, *v_t_* be the observed VAF in tumor and *d* be the sequencing depth in both blood and tumor. For convenience, *d* is fixed to be 500 as per the typical coverage for IMPACT sequencing panels.

A called mutation *m* can be either be true CH (a somatic mutation in the blood), an artifact, or a germline mutation. If *m* is a real CH mutation, then we would expect the tumor VAF to be a product of the amount of blood contamination in the tumor and the true VAF in the blood, *f*_t_ = *cf*_b_. If *m* is an artifact or a germline mutation, we would expect the tumor VAF to be same as the blood VAF, namely *f*_t_ = *f*_b_. *v_t_* is expected to follow a binomial distribution based on the sequencing depth *d* and true VAF in the blood and tumor respectively. Thus, the observed blood VAF can be modeled by *v_b_ ∼ Bin*(*d*,*f*_b_) while the observed tumor VAF can be modeled by *v_t_ ∼ Bin*(*d*,*c*,*f*_b_). We simulated the observed blood and tumor VAFs for real and artifactual mutations under a range of blood contamination levels (*c* = {0.05, 0.1, …, 0.5}) and true blood VAFs (*f*_b_.= {0.02, 0.04, …, 0.2}). Using this synthetic dataset, we evaluated two methods (with different threshold parameters) that aim to differentiate real CH variants from non-CH variants using the observed VAFs:

1. Blood-tumor Ratio: Predict mutation is real if *v_b_ /v_t_* >= C otherwise consider it an artifact. We evaluated a range of cutoffs for C {1, 1.5, 2, 3, 4}.
2. Binomial test: Predict mutation is real if p<0.05 from a binomial test with the null hypothesis of *v_b_ =v_t_*

The predictions by each method were evaluated against the true mutation types that gave rise to the data points, and were classified as true positive (TP), false positive (FP), true negative (TN), and false negative (FN). The overall precision of various methods/cutoffs were calculated as TP/TP+FP and its recall as TP/TP+FN (Supplementary Figures 3 and 4).

Since we expect most CH mutations to have a true variant allele frequency of less than 10% and since we expect most solid tumors to a range of contamination levels with leukocytes (but generally less than 20%), based on our simulations we used *v_b_ /v_t_* cutoff of two. However, in the situation where a lymph node with metastatic disease was chosen as the source for tumor material, a high level (greater than 30%) of leukocyte contamination could be present in some cases.

---

**Extended Data Figure 1.**
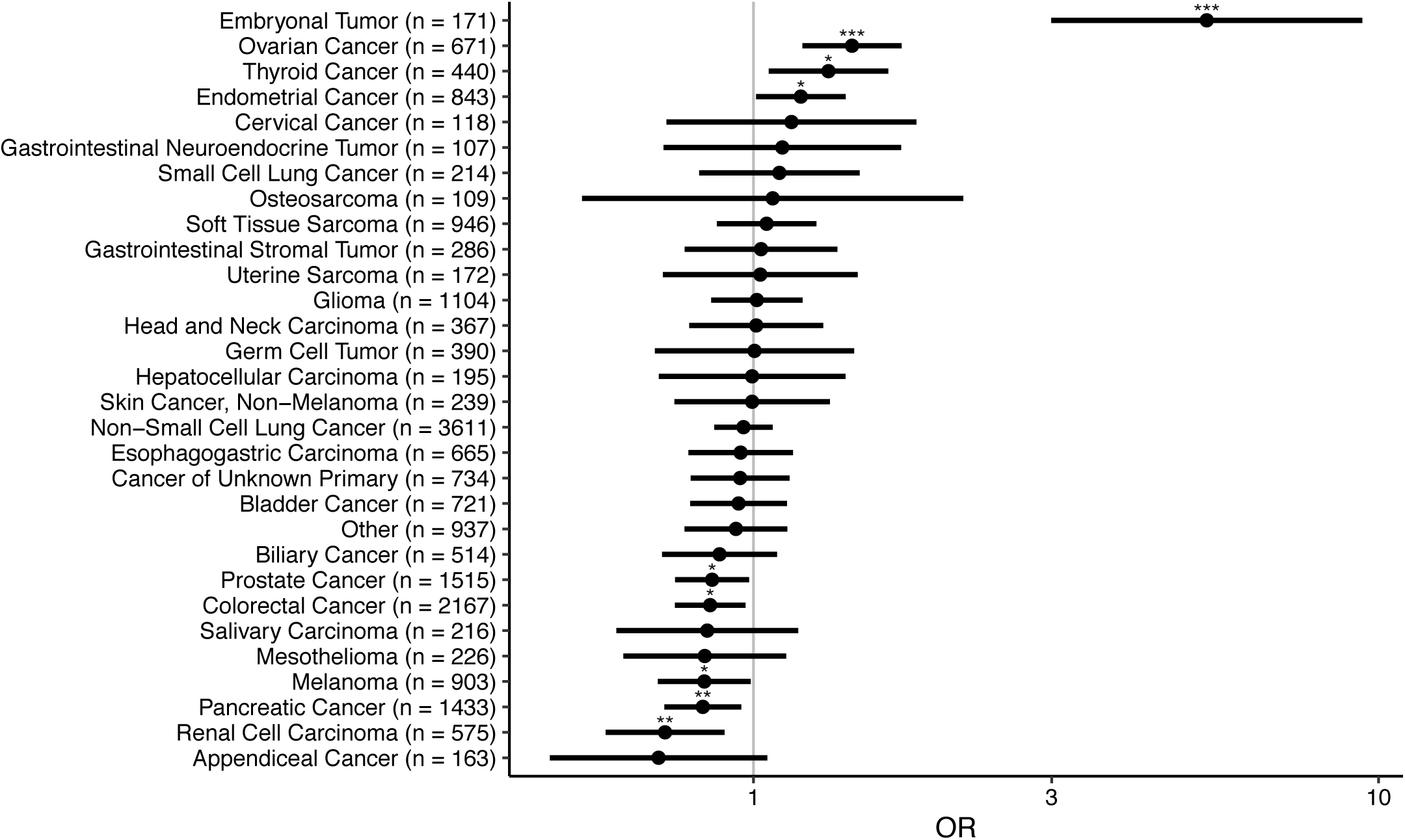
Association between primary tumor site and clonal hematopoiesis. Shown are the odds ratios and 95% confidence intervals for clonal hematopoiesis in selected primary tumor types with at least 100 subjects compared to breast cancer (N=3553) in a logistic regression model adjusted for age. P-values: * <0.05, ** <0.01,***<0.001

**Extended Data Figure 2.**
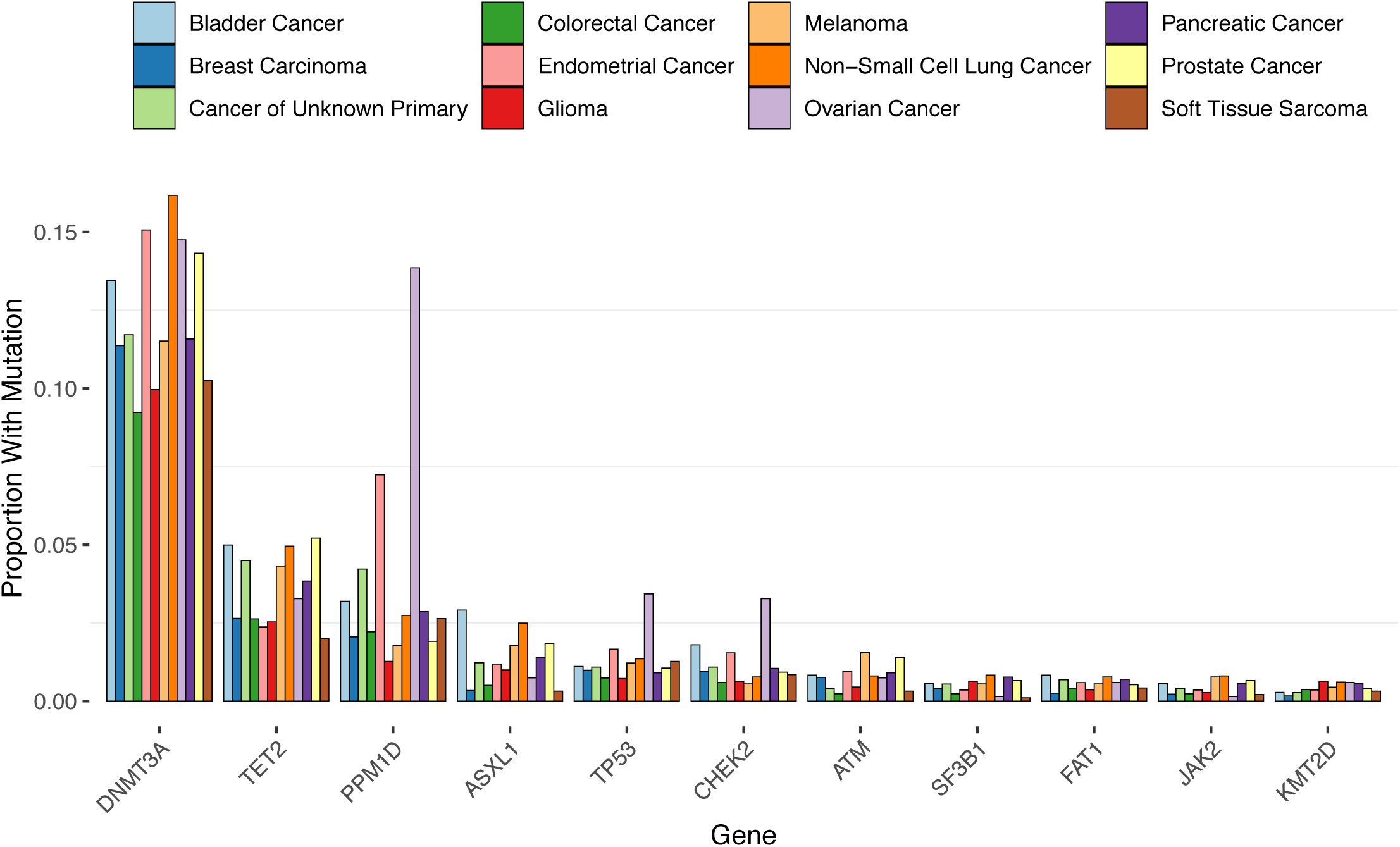
Proportion of patients with common CH mutations by primary tumor sites. Genes mutated in at least 75 individuals and the top 12 primary tumor sites are shown.

**Extended Data Figure 3.**
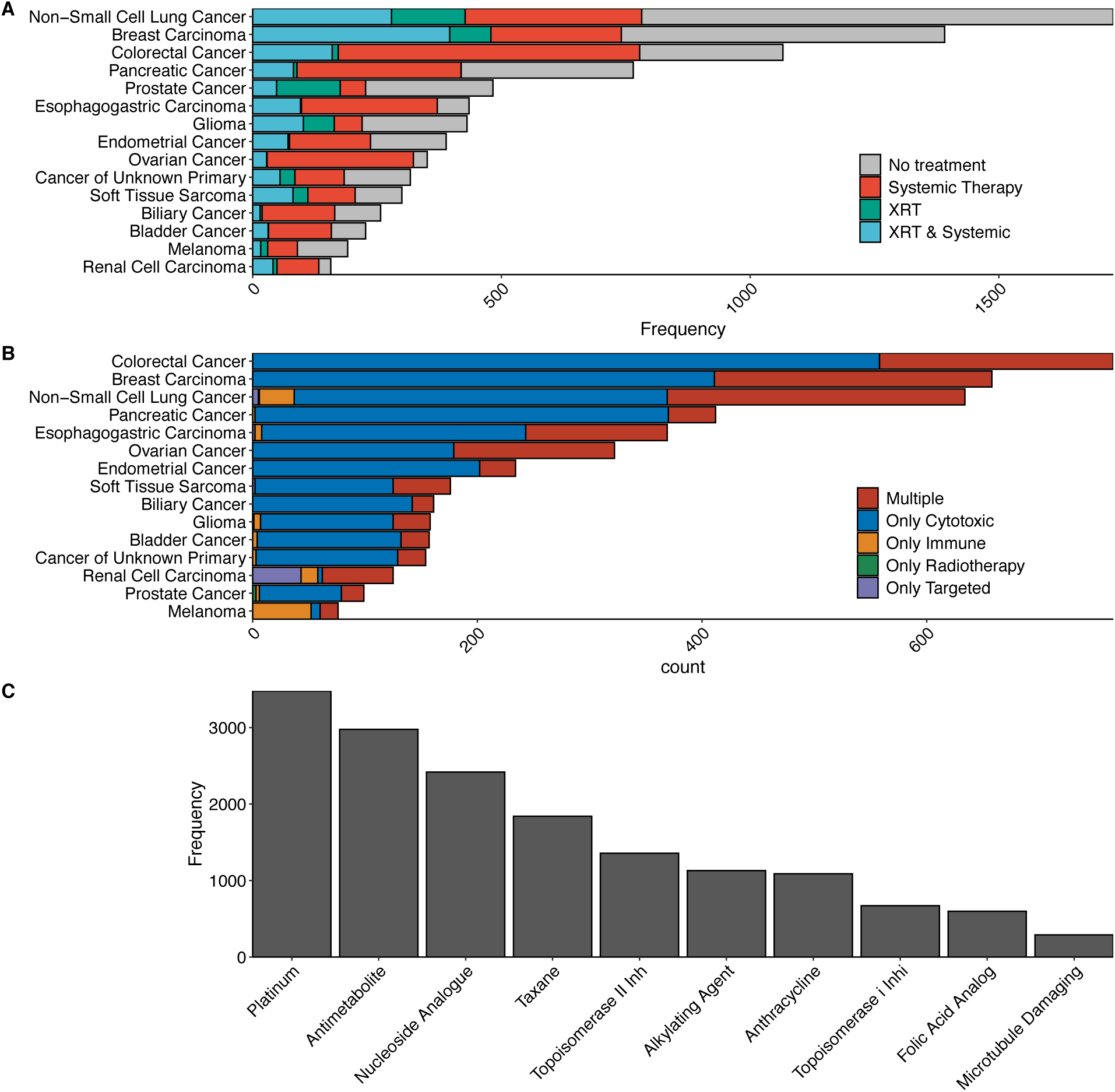
Distribution of oncologic therapy received prior to blood collection for sequencing. A) Frequency of patients receiving systemic therapy or external beam radiation therapy by primary tumor type. B) Frequency of patients receiving specific classes of systemic therapy by primary tumor type. C) Frequency of patients receiving top ten subclasses of cytotoxic therapy. Most patients (91%) who received at least one of these cytotoxic therapy classes received multiple classes.

**Extended Data Figure 4.**
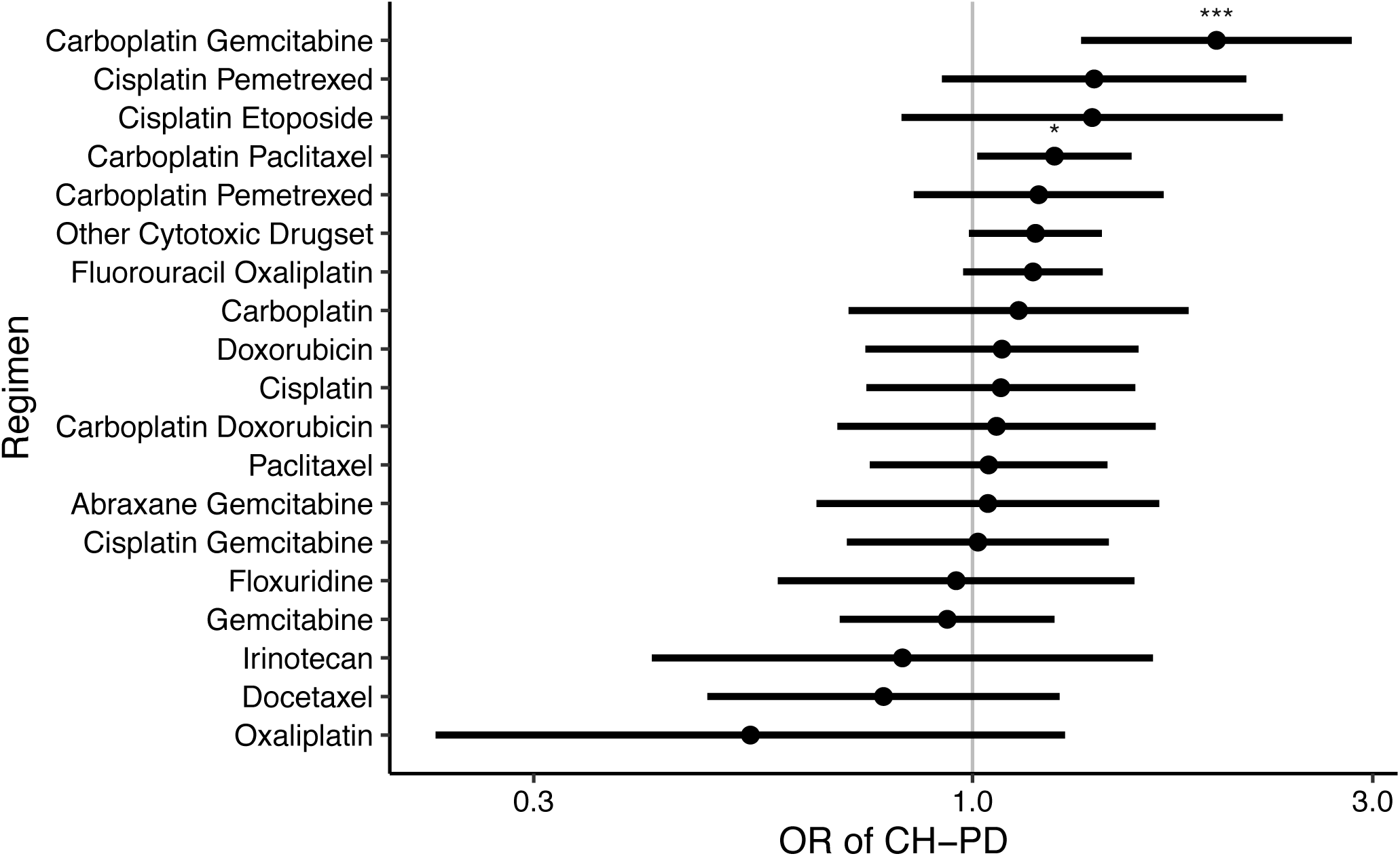
Association between most commonly prescribed drug regimens containing at least one cytotoxic therapy and CH-PD. Shown are the odds ratios of CH-PD given prior exposure to the most commonly prescribed drug regimens adjusted for each other, external beam radiation, smoking, age, race and time from primary tumor diagnosis to blood draw.

**Extended Data Figure 5.**
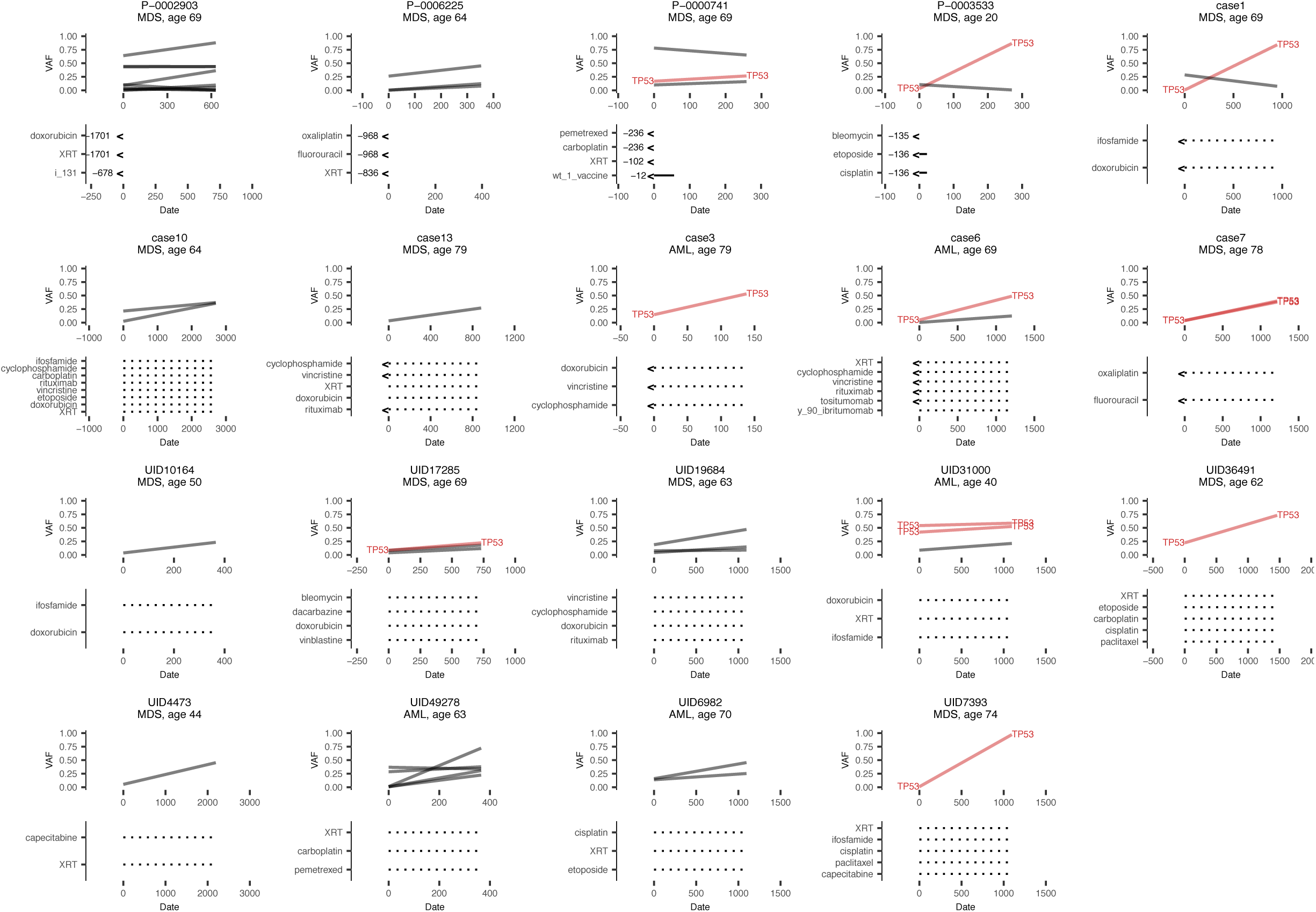
Variant frequencies (VAF) at time of pre-tMN testing and tMN diagnosis. Plots show changes in mutational frequencies in relation to oncologic therapy exposure in 19 CH positive cases. *TP53* mutations are shown in red, other mutations are showing in black. Solid lines denote treatment during the interval period where the exact dates are known and dotted lines denote treatment that was received during the interval period but treatment is unknown. Arrows indicate treatment received prior to pre-tMN testing with the number of days between the end of treatment and the pre-tMN sample.

**Extended Data Figure 6.**
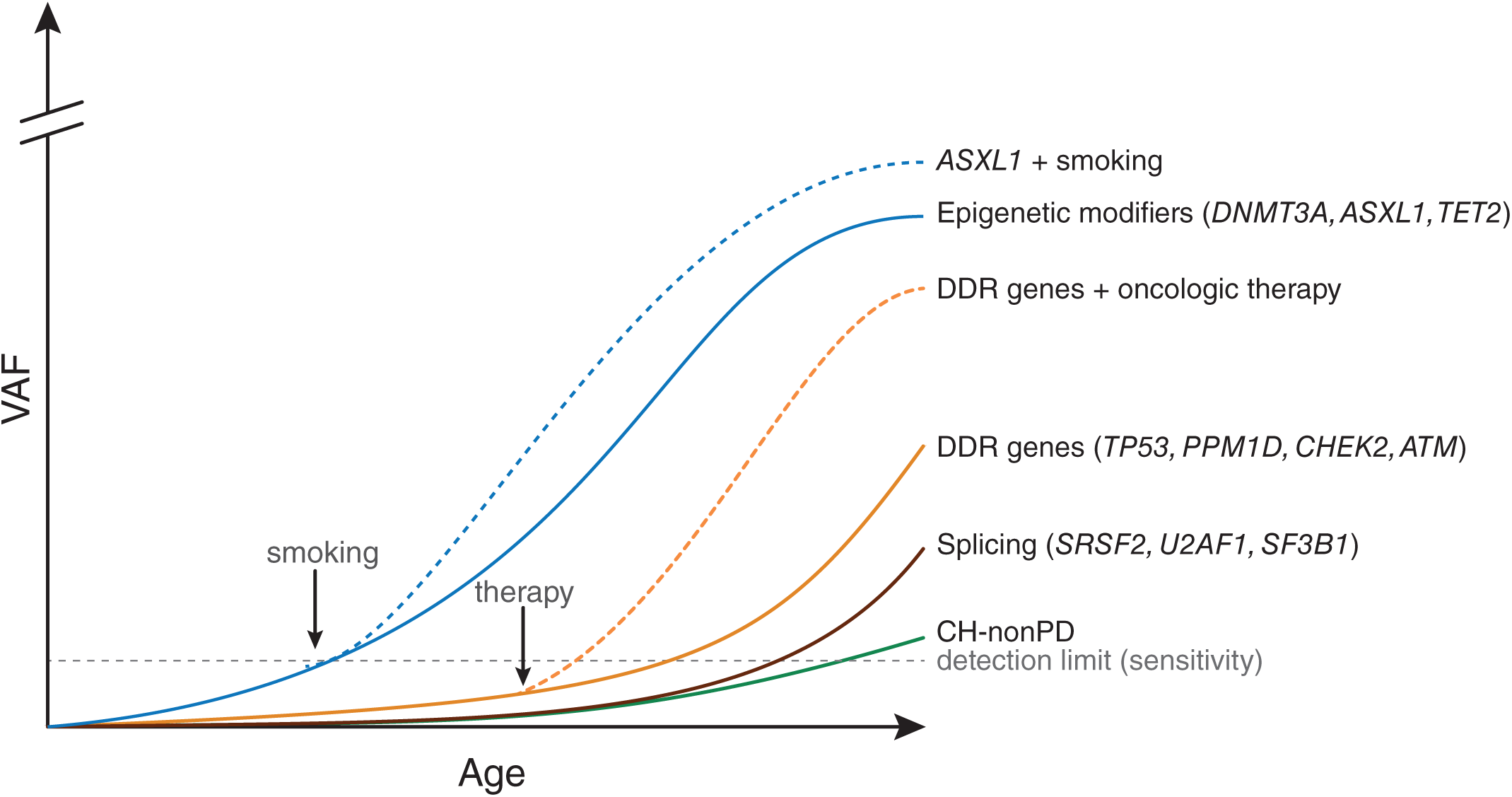
Graphical depiction demonstrating how differences in the fitness effect of CH mutations and the environment shape clonal dominance over an individual’s lifetime. The retrospective associations we observed between smoking and *ASXL1* mutations and oncologic therapy and DDR gene mutations (Figures 1 and 2) largely reflect the result of clonal expansion leading to increased detectability (as shown for treatment and DDR genes in Figure 3.

**Extended Data Table 1.**
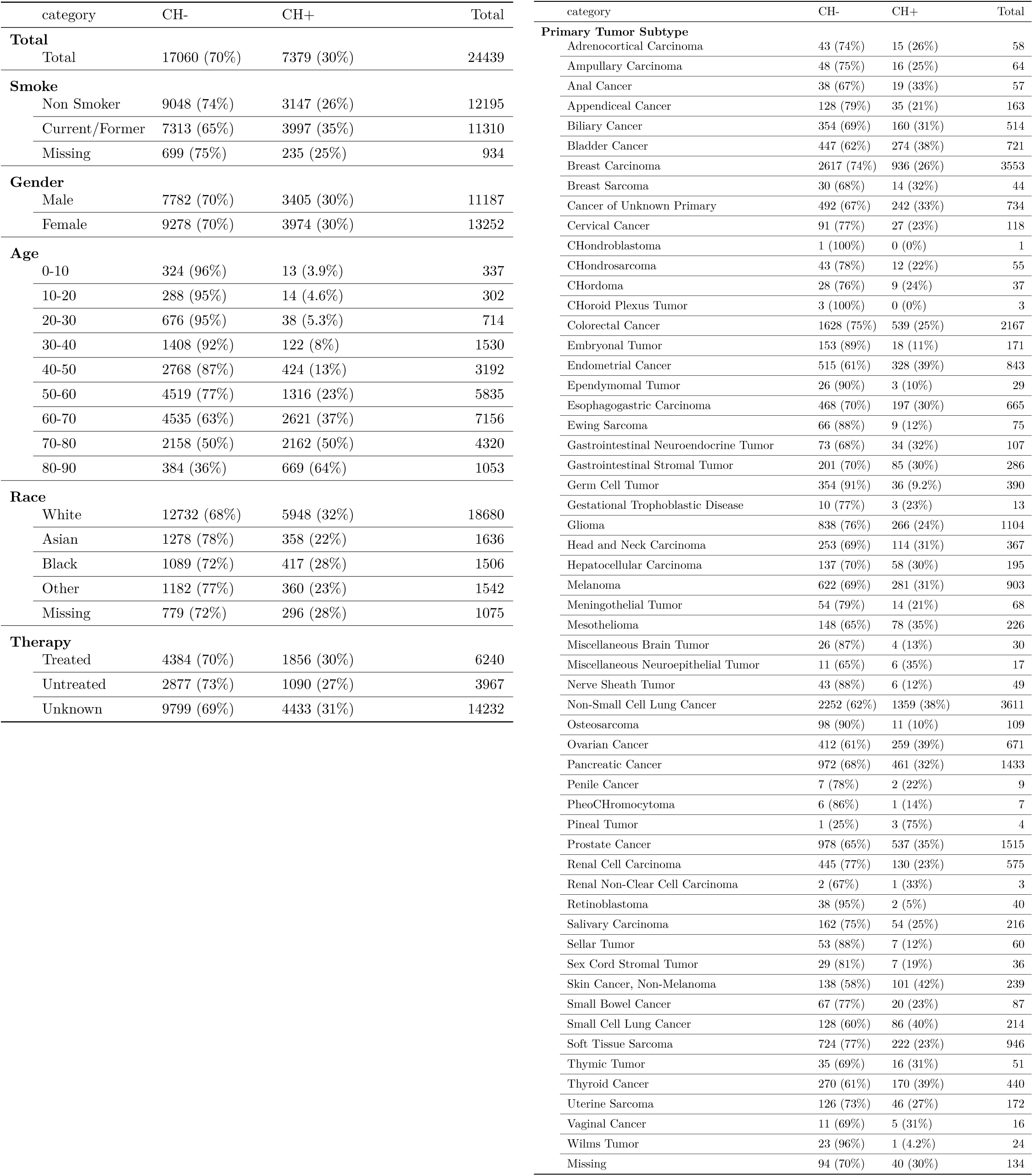
Clinical characteristics of solid tumor patients assessed for CH.

**Extended Data Table 2.**
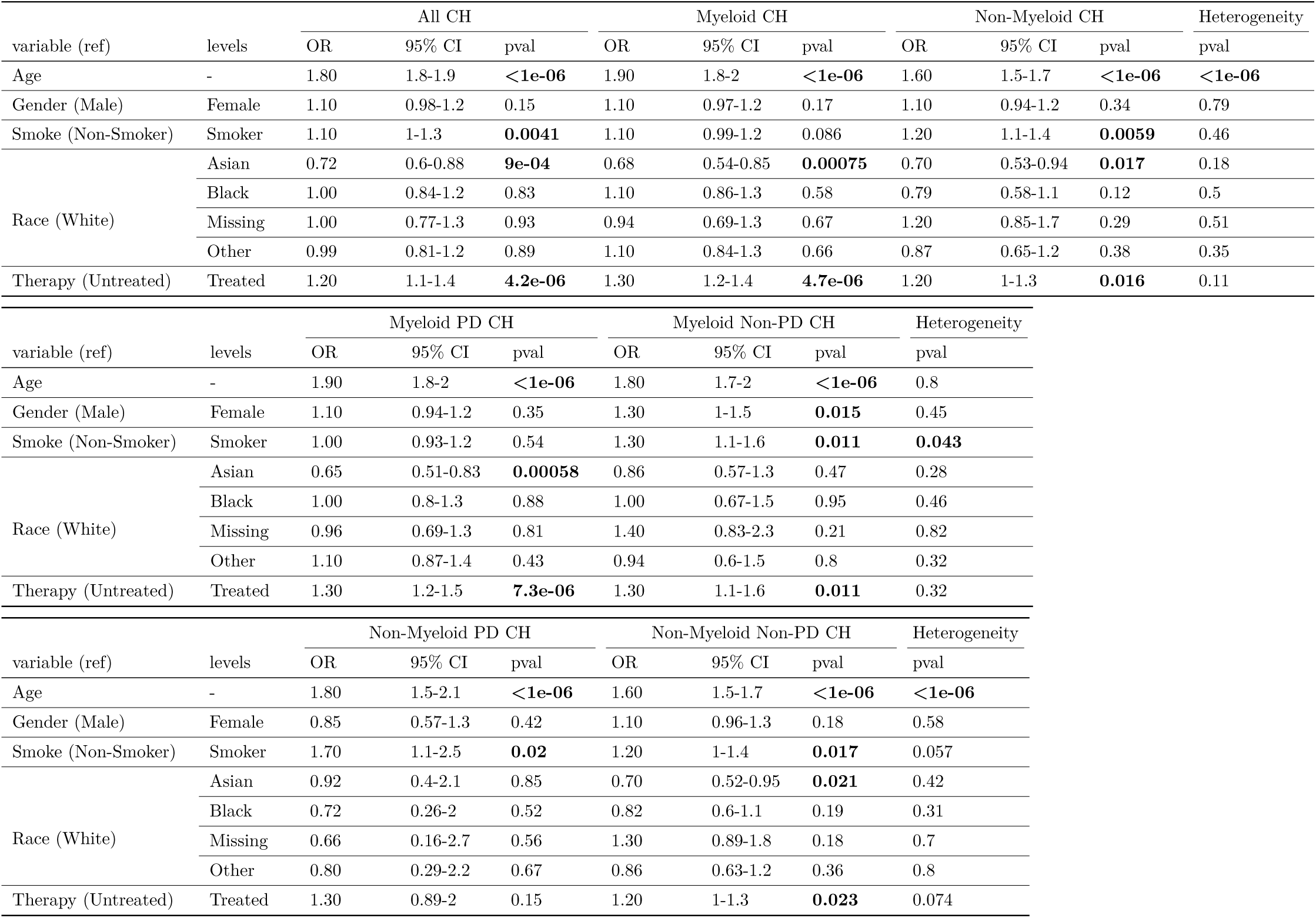
Association between clinical characteristics and CH mutational characteristics. The association between CH and clinical characteristics is evaluated comparing mutations in genes mutated in myeloid neoplasms (Myeloid CH), in genes not linked to myeloid neoplasms (Non-Myeloid CH), synonymous CH variants (silent), non-synonymous CH variants (non-silent), variants known to be myeloid drivers (Myeloid PD CH), mutations that are putative somatic driver mutations in myeloid neoplasms (Myeloid PD CH), mutations within genes linked to myeloid neoplasms but that are not putative drivers (Myeloid Non-PD CH), mutations that are putative somatic driver mutations of cancer in genes not linked to myeloid neoplasms (Non-Myeloid PD CH) and mutations within genes not linked to myeloid neoplasms that are not putative drivers of cancer (Non-Myeloid Non-PD CH) using multivariable logistic regression models. Heterogeneity p-values for clinical variables were calculated using logistic regression testing for a difference in the odds of having at least one CH mutation in myeloid compared to CH only in non-myeloid genes, for silent compared to non-silent mutations, myeloid PD CH compared to myeloid non-PD CH and Non-myeloid PD CH compared to Non-Myeloid Non-PD CH for predictor variables listed in the table. Sensitivity analyses restricted to individuals with only one mutation yielded similar results. Age expressed in decile.

**Extended Data Table 3.**
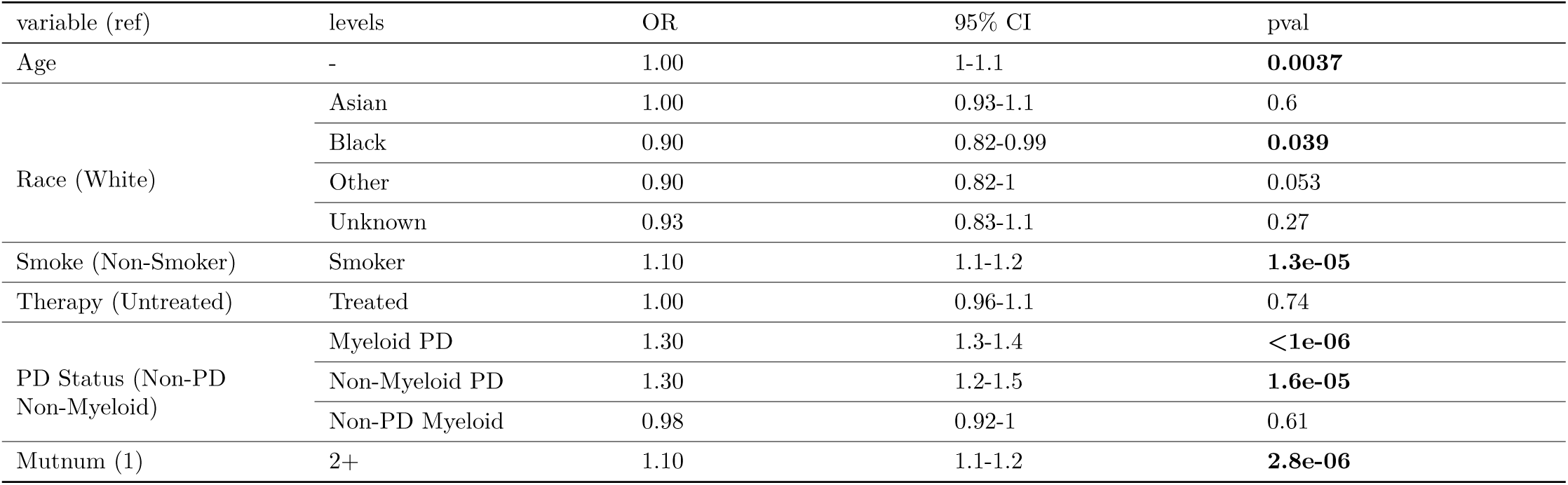
Association between variant allele fraction (VAF) of CH mutations and clinical characteristics. Generalized estimating equations were used to test for an association between VAF of CH among those with a mutation and selected clinical and mutational features accounting for correlation between the VAF of mutations in the same person. Age expressed in decile.

| variable (ref) | levels | OR | 95% CI | pval |
| --- | --- | --- | --- | --- |
| Age | - | 1.00 | 1-1.1 | **0.0037** |
| Race (White) | Asian | 1.00 | 0.93-1.1 | 0.6 |
|  | Black | 0.90 | 0.82-0.99 | **0.039** |
|  | Other | 0.90 | 0.82-1 | 0.053 |
|  | Unknown | 0.93 | 0.83-1.1 | 0.27 |
| Smoke (Non-Smoker) | Smoker | 1.10 | 1.1-1.2 | **1.3e-05** |
| Therapy (Untreated) | Treated | 1.00 | 0.96-1.1 | 0.74 |
| PD Status (Non-PD<br>Non-Myeloid) | Myeloid PD | 1.30 | 1.3-1.4 | **<1e-06** |
|  | Non-Myeloid PD | 1.30 | 1.2-1.5 | **1.6e-05** |
|  | Non-PD Myeloid | 0.98 | 0.92-1 | 0.61 |
| Mutnum (1) | 2+ | 1.10 | 1.1-1.2 | **2.8e-06** |

**Extended Data Table 4.**
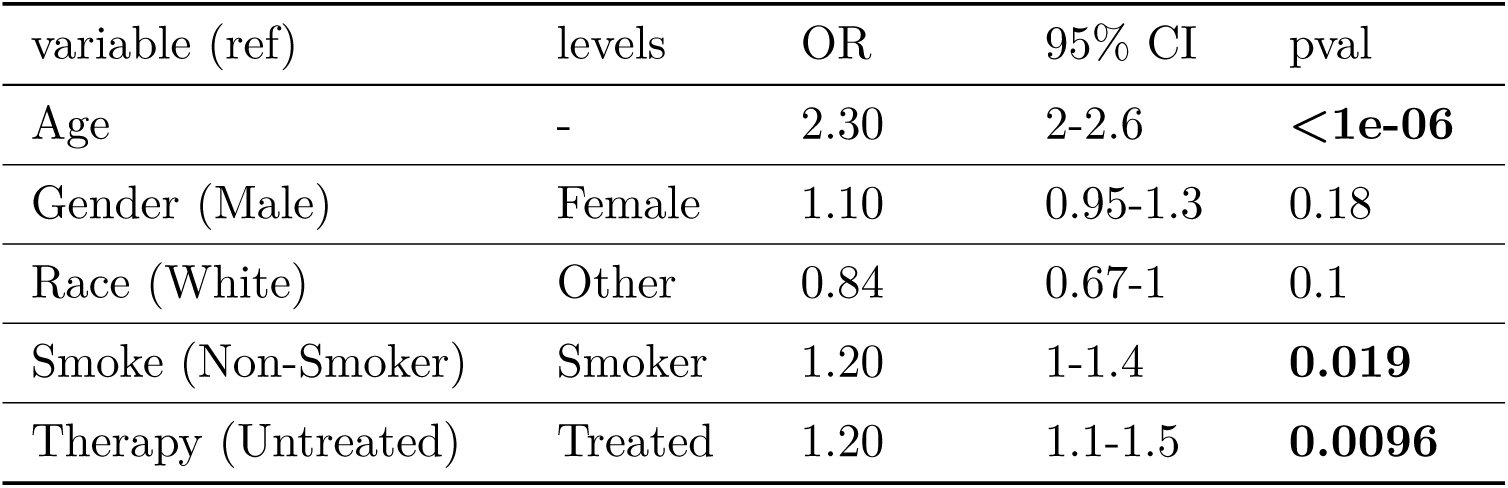
Association between CH mutation number and clinical characteristics. Ordinal logistic regression was used to test for an association between clinical characteristics and mutation number in patients with clonal hematopoiesis in a multivariable model. Age is expressed as the decile.

**Supplementary Figure 1.**
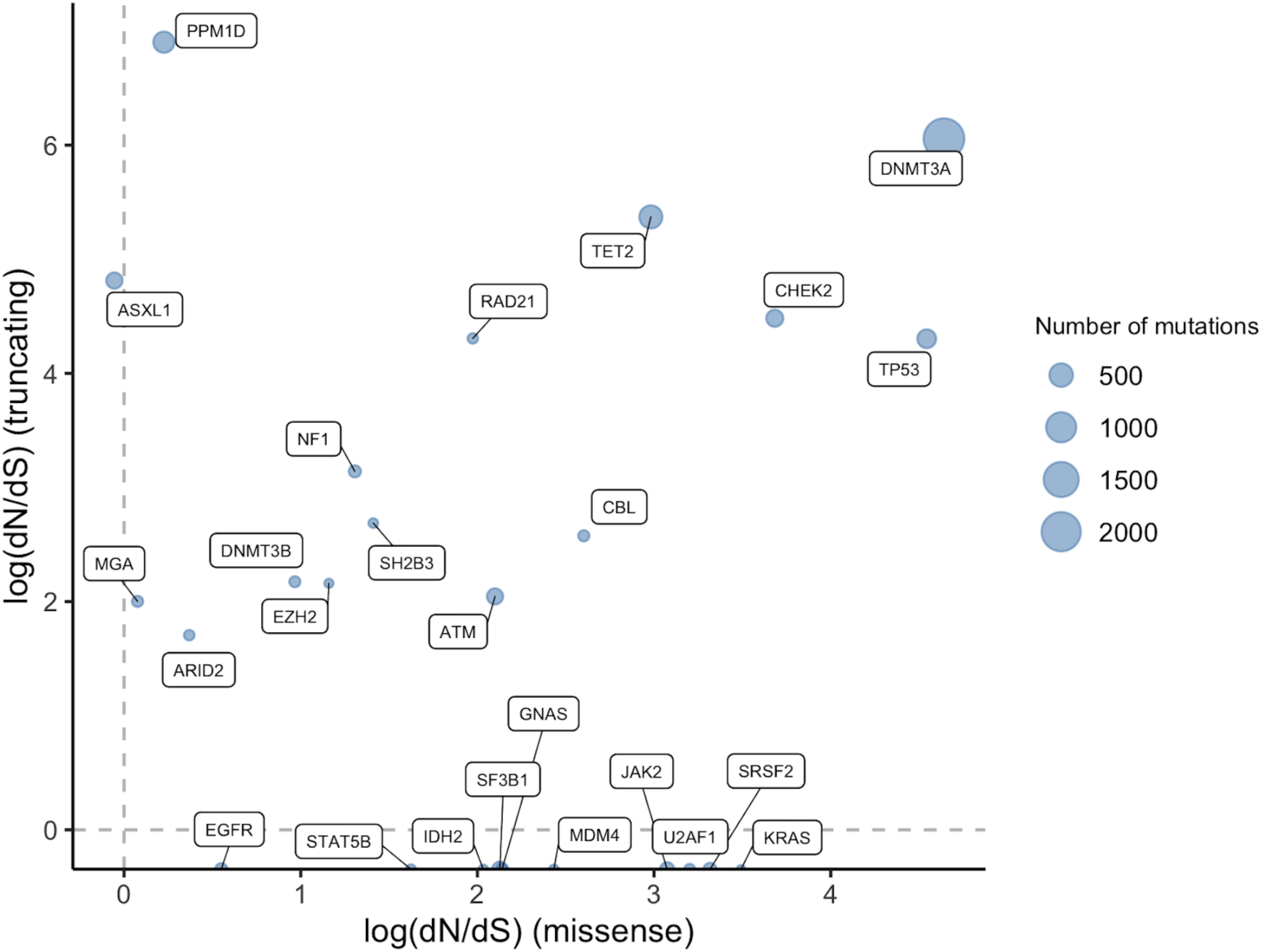
Quantification of the extent of natural selection by gene in clonal hematopoiesis using a dN/dS method. Using the dNdScv method (see Methods) we quantified the dN/dS ratios for missense and nonsense and essential splice mutations (truncating), at the level of individual genes. Shown are the dN/dS ratios for genes mutated at least 25 times showing evidence of significant selection. The log(dN/dS)<0 evidences negative selection and log(dN/dS)>0 evidences positive selection.

**Supplementary Figure 2.**
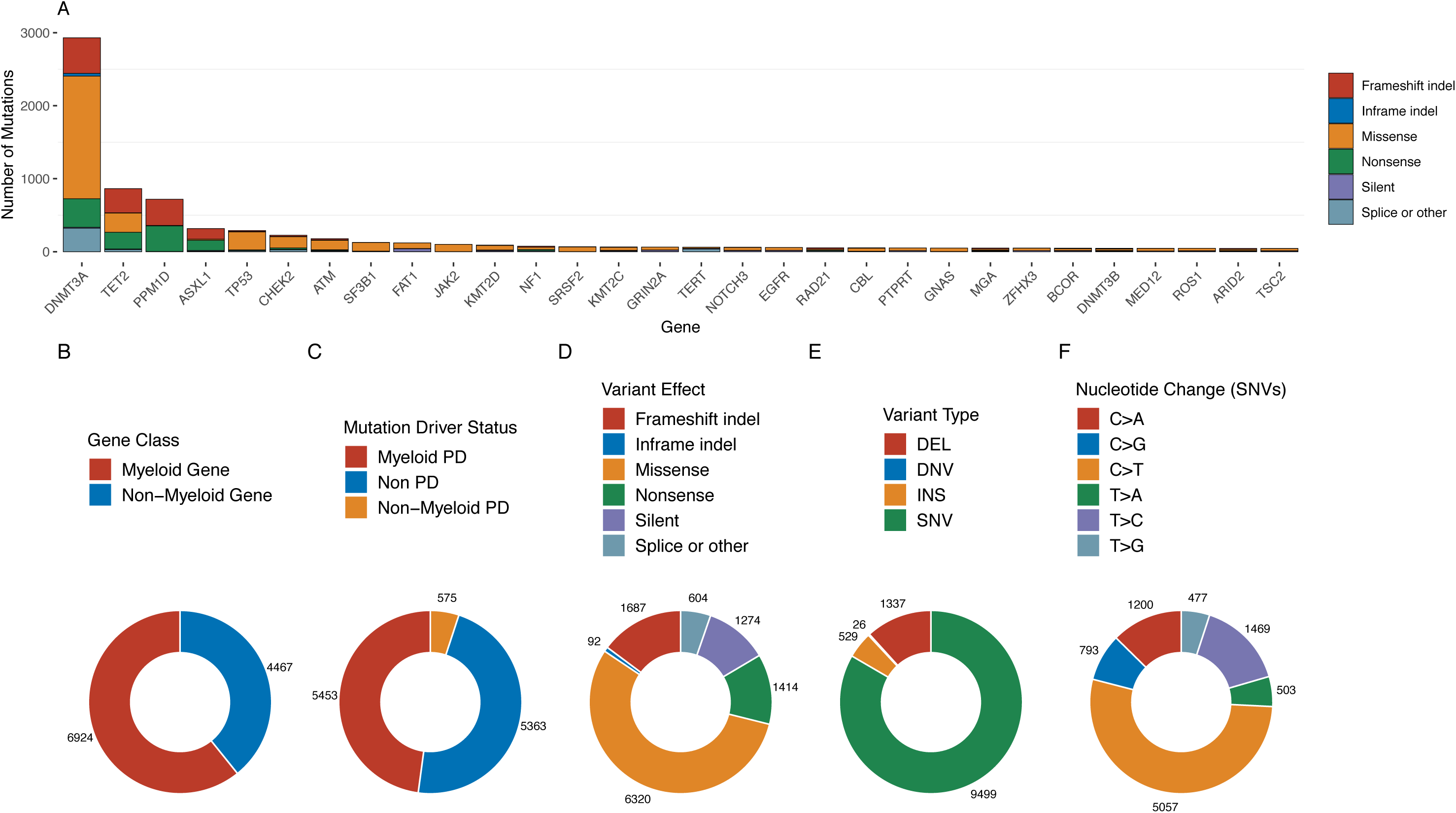
Mutational characteristics of CH in solid tumor patients. (A) Number of mutations observed in the 30 most common genes. (B) Proportion of mutations in a myeloid gene and those not in a myeloid gene. (C) Proportion of mutations considered to be a possible driver of myeloid neoplasms (Myeloid PD), a driver of non-myeloid neoplasms (Non-Myeloid PD) and those not considered to be a possible cancer driver. (D) Proportion of mutations by functional effect. (E). Proportion of deletions (DEL), insertions (INS) or SNVs. (F). Proportion of single nucleotide variants (SNV) with specific nucleotide.

**Supplementary Figure 3.**
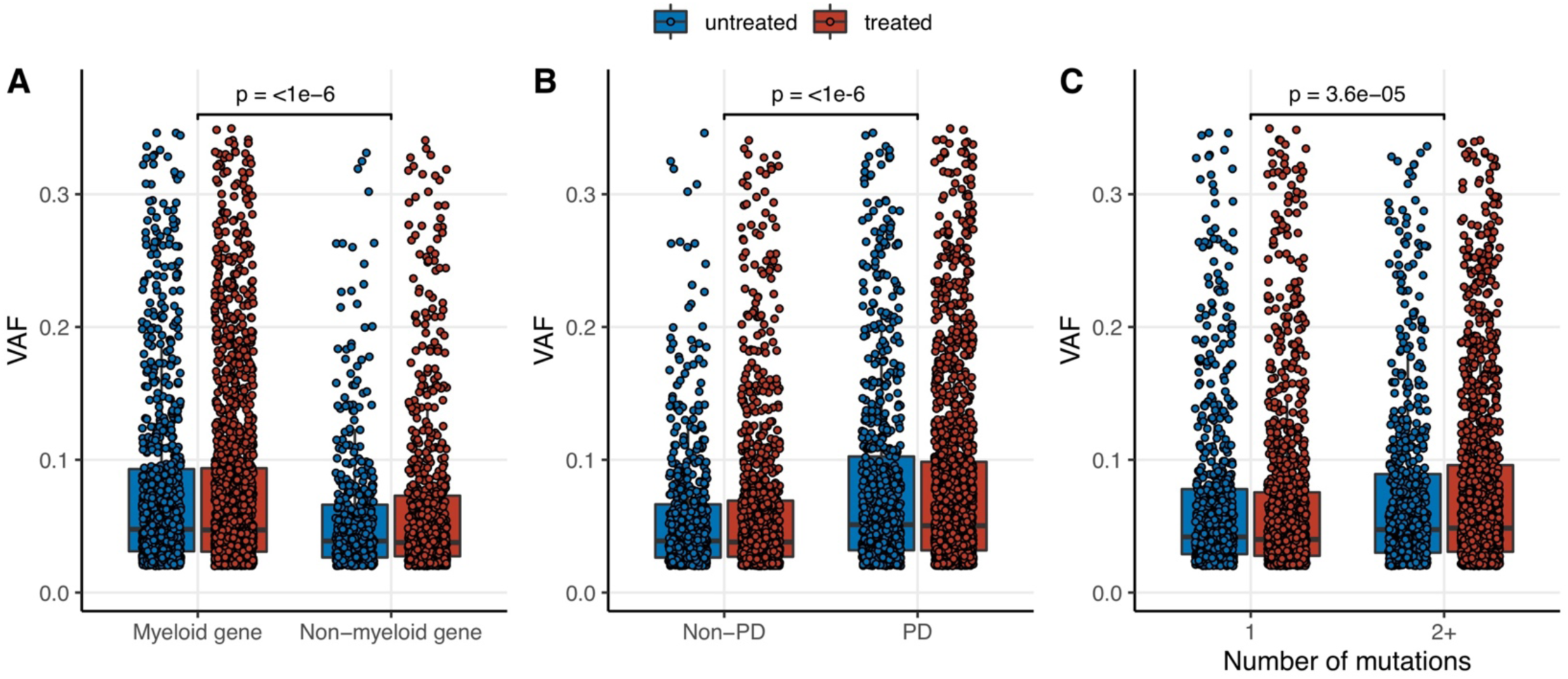
Relationship between variant allele fraction (VAF) and CH mutational features in (A) genes recurrently mutated in myeloid neoplasms (myeloid gene) and those not implicated in myeloid disease (Non-myeloid gene), (B) in variants thought to be putative cancer drivers (PD) and variants not known to be cancer drivers (Non-PD) and (C) by the total number of mutations within the individual. P-values were calculated from generalized estimating equations testing for associations between VAF and mutational features adjusted for age, sex, race, smoking history and exposure to oncologic therapy accounting for the within-subject correlation in VAF.

**Supplementary Figure 4.**
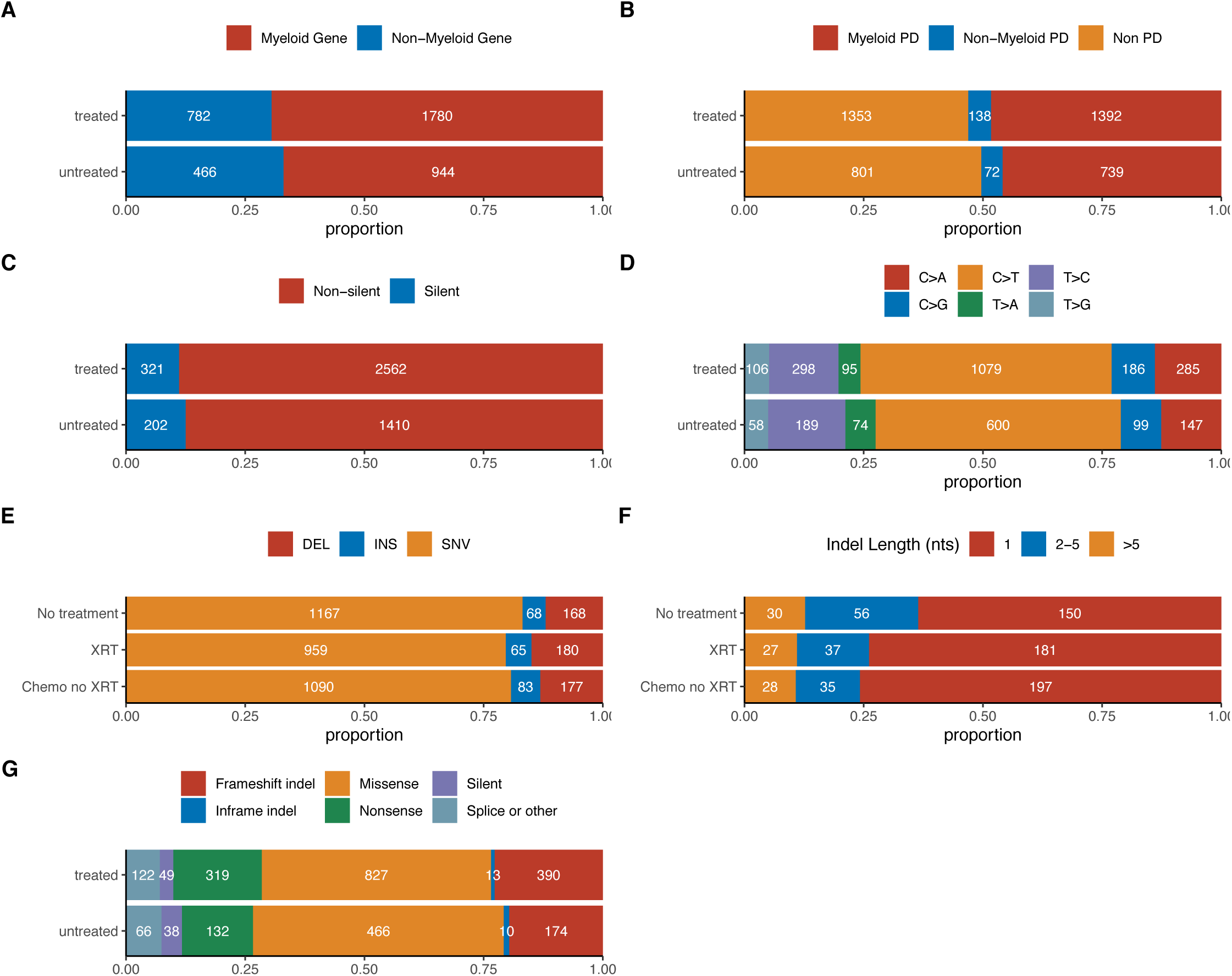
Mutational characteristics of CH in solid tumor patients and prior exposure to oncologic therapy. Among patients who received any oncologic treatment prior to blood draw for mutational testing (treated) and those who did not receive therapy prior to blood draw (untreated) we compared the following. (A) Proportion of mutations in a hypothesized myeloid neoplasm driver gene (myeloid gene) and those in a gene not known to be a driver of myeloid neoplasms (non-myeloid gene). (B) Proportion of mutations considered to be a possible driver of myeloid neoplasms (myeloid PD), a driver of non-myeloid neoplasms (non-myeloid PD) and those not considered to be a possible cancer driver (non-PD). (C) Proportion of non-synonomous (non-silent) and synonymous (silent) mutations. (D). Proportion of mutations within major functional effect categories. (E) Proportion of deletions (DEL), insertions (INS) or SNVs. (F) Proportion of insertions or deletions by the nucleotide length of the alteration. (G) Proportion of single nucleotide variants (SNV) with specific nucleotide changes.

**Supplementary Figure 5.**
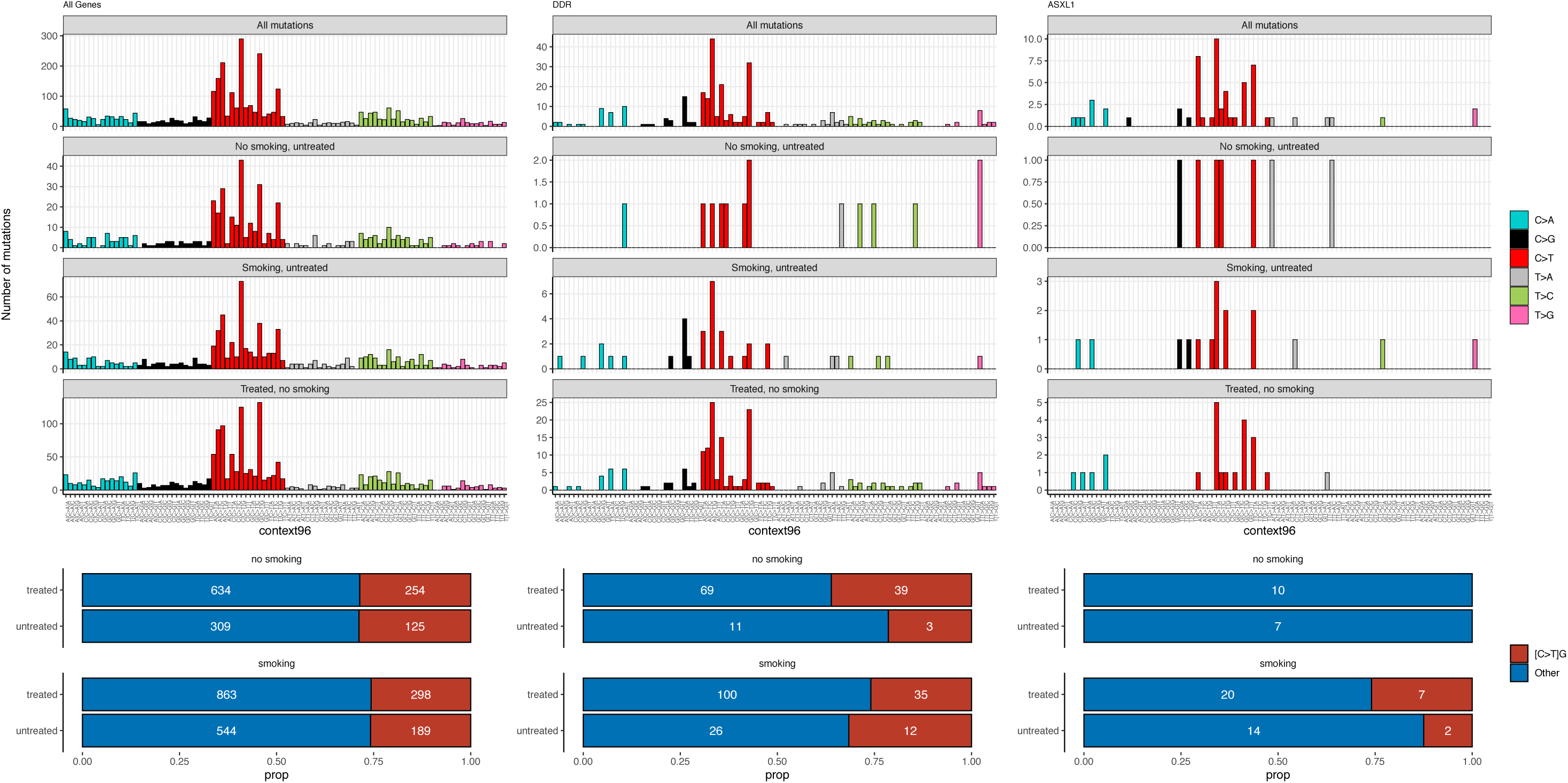
Mutational context of single nucleotide CH mutations and the proportion of SNVs with C>T in CpG context overall (A), in DDR genes (B) and *ASXL1* (C) according to smoking history and prior exposure to oncologic therapy.

**Supplementary Figure 6.**
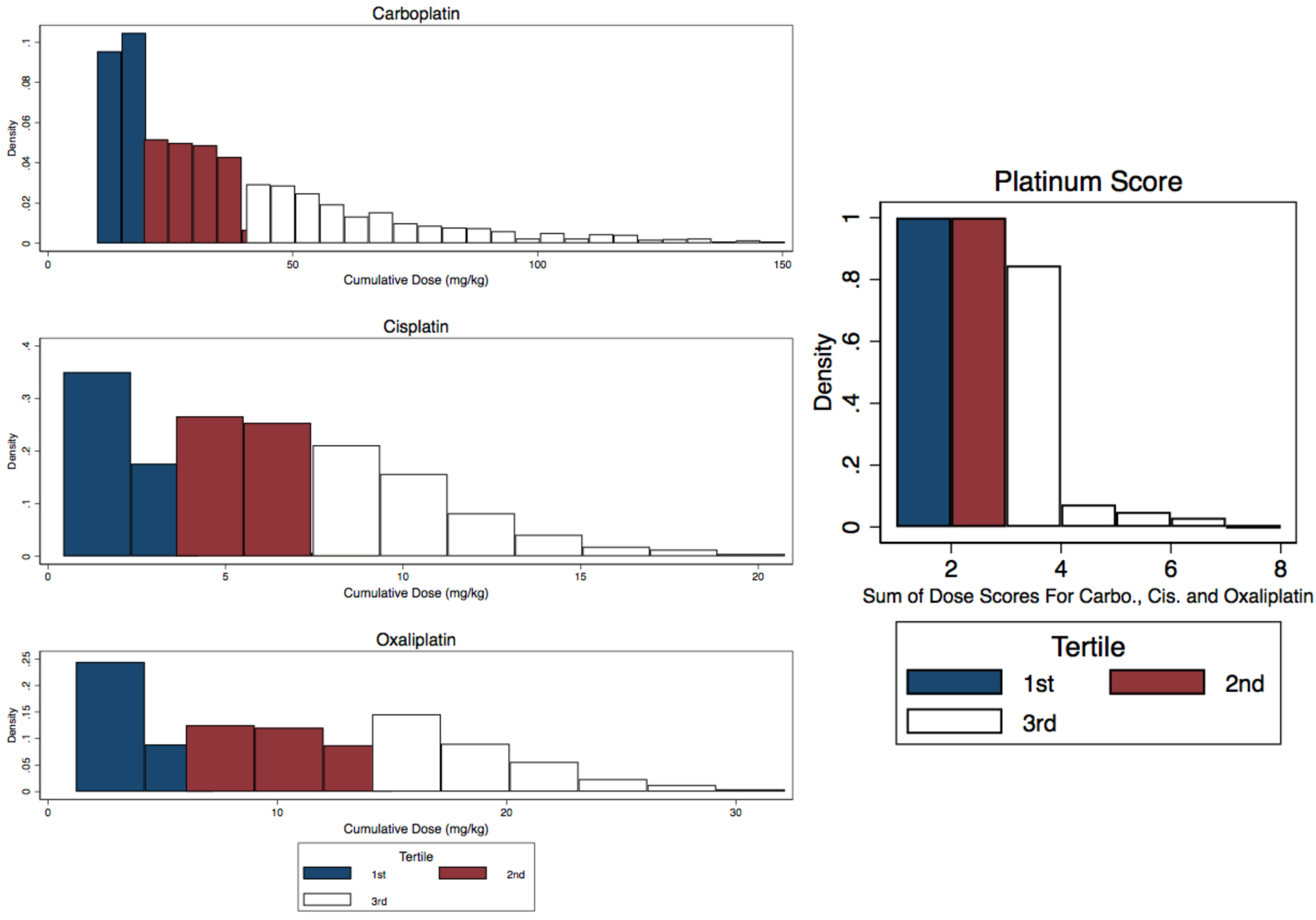
Calculation of cumulative exposure for therapy subclasses. For each drug, the sum of the dose (per kilogram of body weight) received prior to blood draw was summed for each patient. The dose distribution for each agent was divided into tertiles and the patient’s dose was assigned a score based on tertile. The scores for an individual patient were summed among drugs belonging to a specific class. The distribution of the resulting sum was used to derive tertiles of total exposure to the drug class that was again used to assign a score of 0-3 for each individual. For example, if an individual received 100mg/kg of carboplatin, 2mg/kg of cisplatin and no oxaliplatin, their platinum score would be 4 (3 for carboplatin, 1 for cisplatin and 0 for oxaliplatin), they would be assigned to the 3^rd^ tertile of platinum.

**Supplementary Figure 7.**
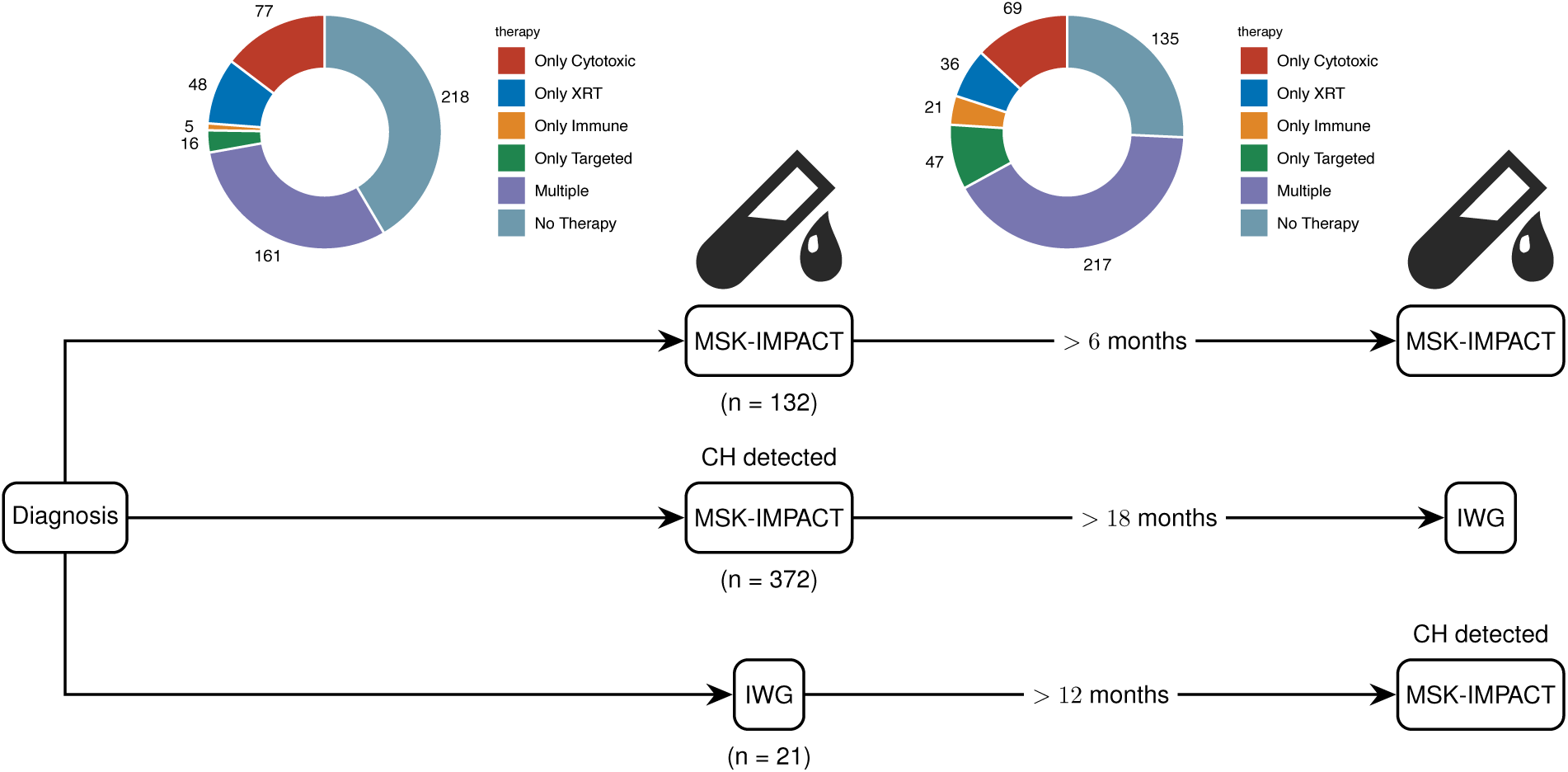
Overview of serial samples included in the study relative to MSK-IMPACT testing. We analyzed sequential blood samples in 525 patients. The majority of these were among patients found to have CH on MSK-IMPACT and a follow-up blood sample was collected at least 18 months following MSK-IMPACT testing (N=132). We obtained 160 samples that had repeat MSK-IMPACT testing performed on the blood at least six months apart for clinical purposes (not selected for CH positivity) and 17 samples that had CH on MSK-IMPACT who also had a previously banked blood sample at least 12 months prior to MSK-IMPACT testing.

**Supplementary Figure 8.**
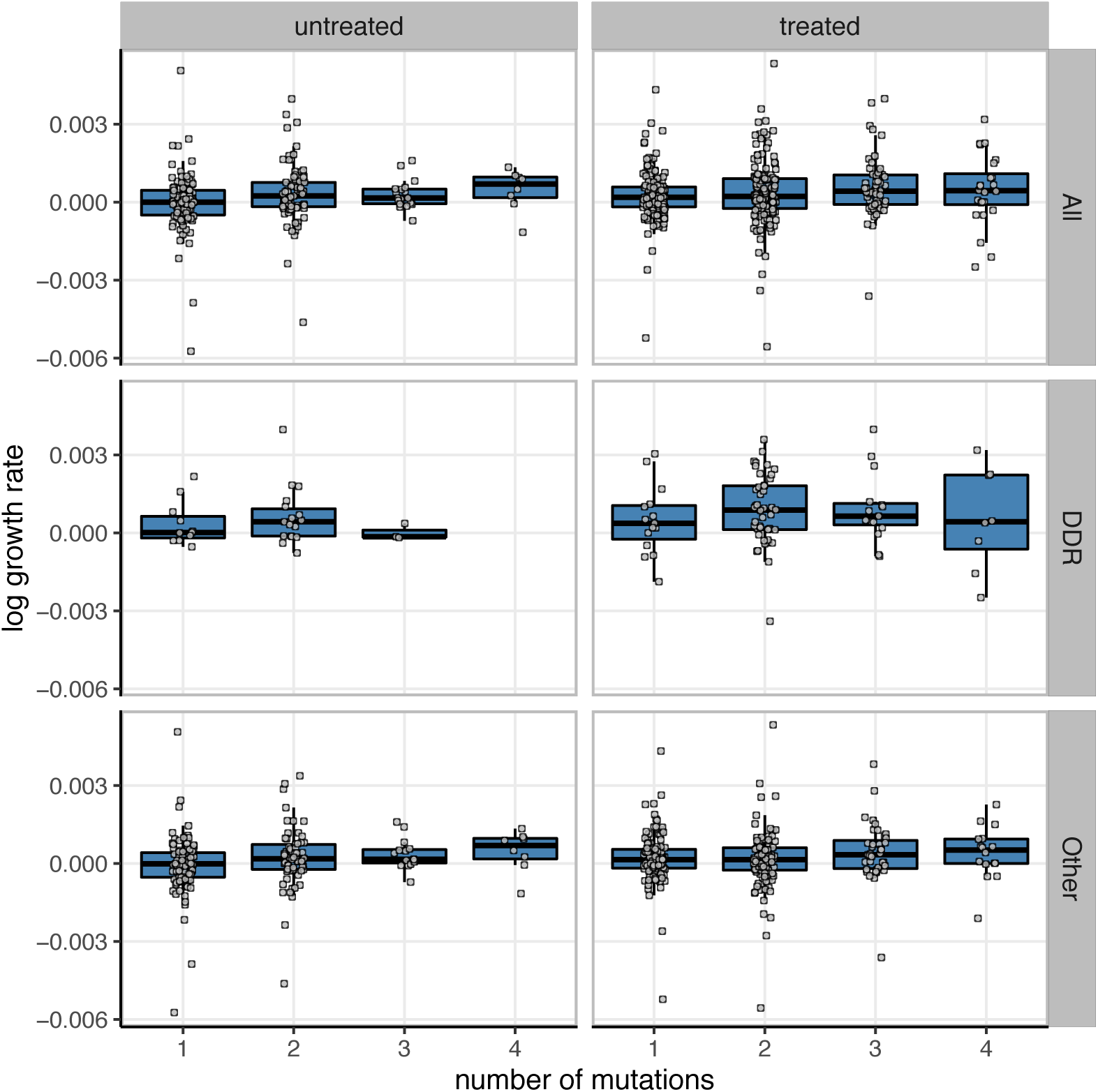
Growth rate of CH mutations and mutation number. Shown are the growth rates for each mutation during follow-up period by the total number of mutations in that individual stratified by therapy during the follow-up period. Regression using generalized estimating equations was used to calculate a test for trend for increasing growth rate with mutation number among subjects with clonal hematopoiesis adjusted for age, gender, treatment and smoking accounting for correlation between the VAF of mutations in the same person. This supported a slight increase in the average growth rate of mutations with increasing total number of mutations (p-trend=0.03).

**Supplementary Figure 9.**
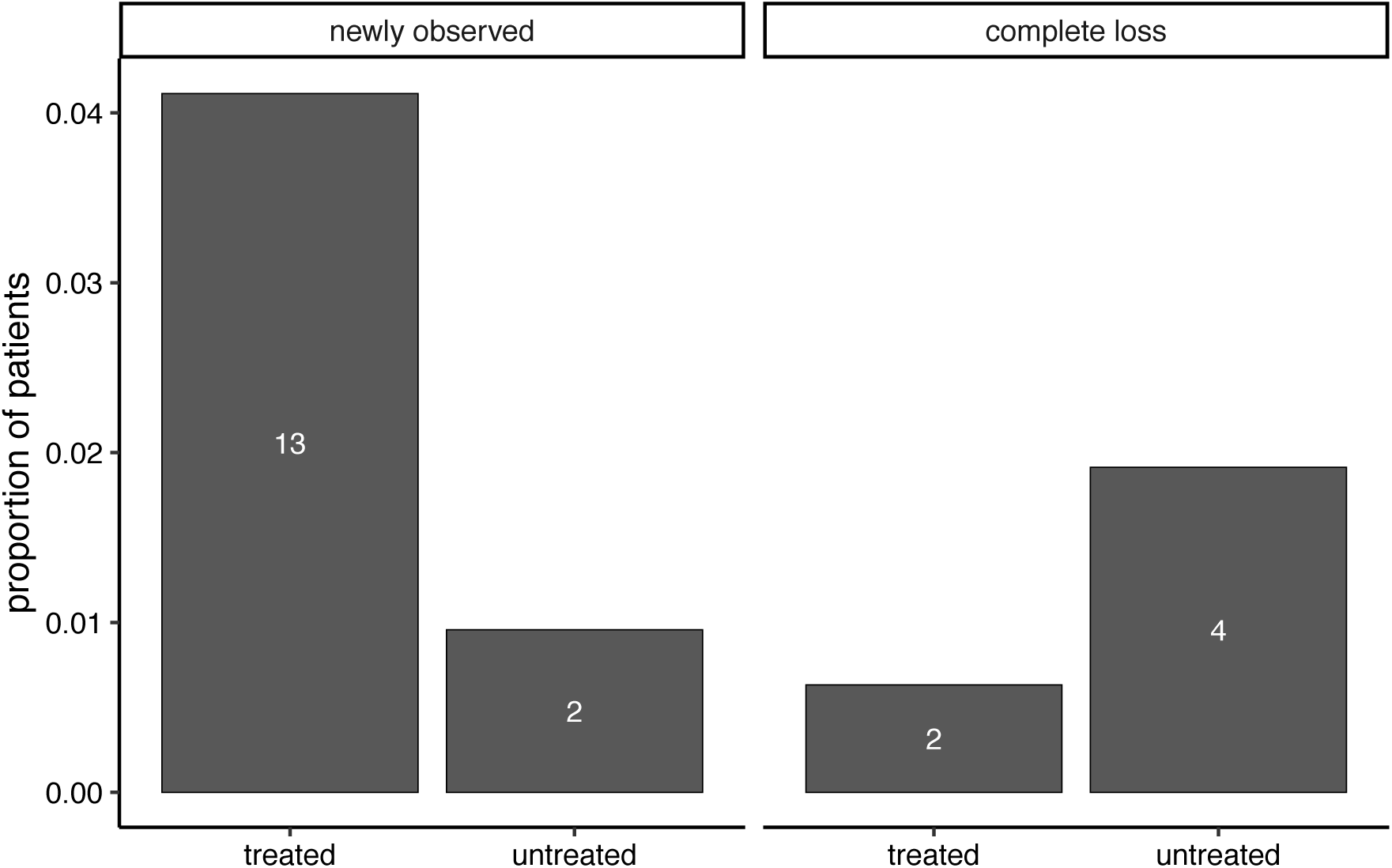
Proportion of patients who gained or lost a mutation during follow-up period stratified by interval therapy. Newly observed: no read at time point 1, VAF >= 2% at time point 2. Complete loss: VAF >= 2% at time point 1, no read detected at time point 2.

**Supplementary Figure 10.**
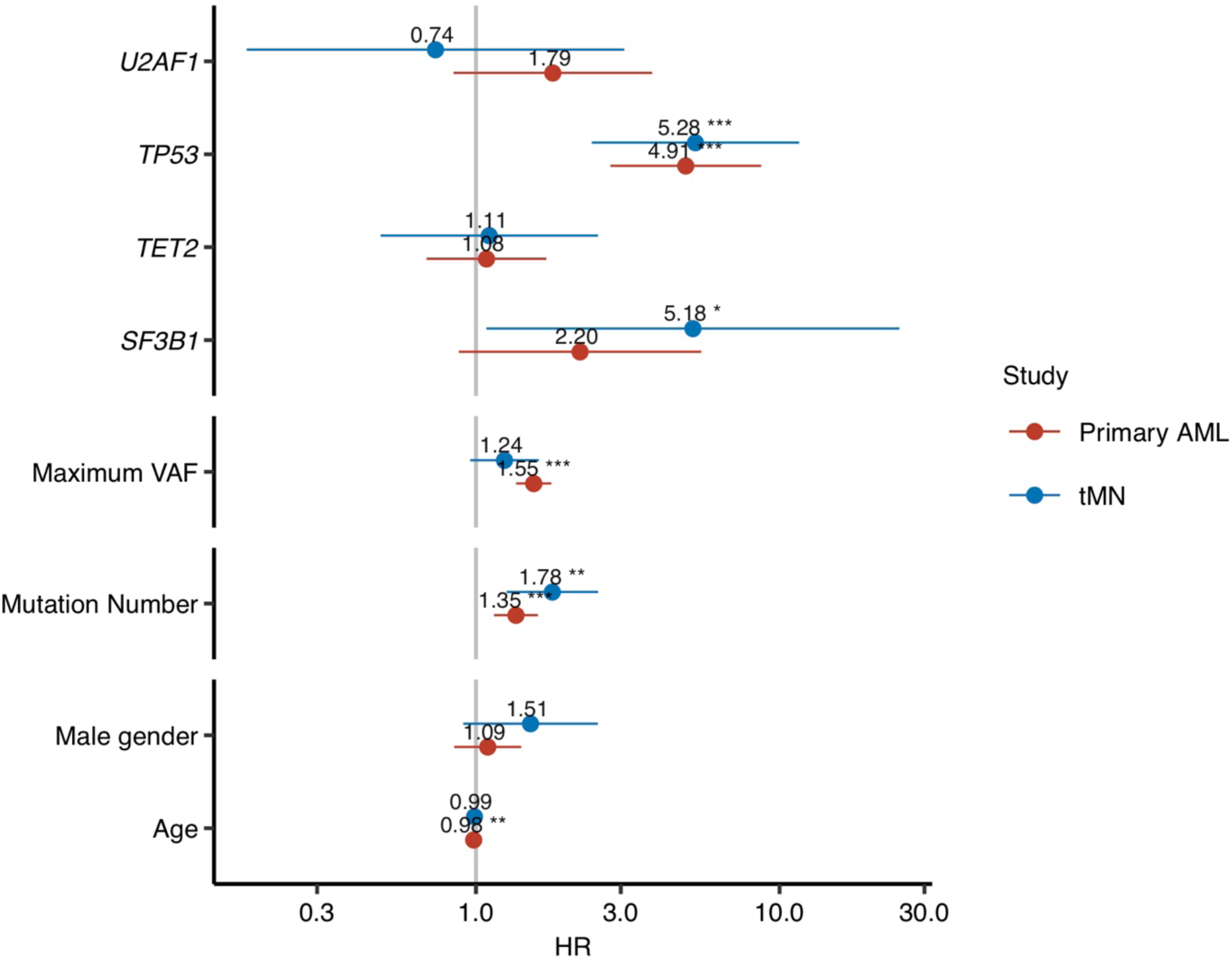
Risk of myeloid malignancy by clinical and mutational characteristics comparing studies for tMN in solid tumor patients and AML risk in healthy individuals. Shown are the hazard ratios from multivariable Cox regression for including CH mutational characteristics. All models were adjusted for age and gender and stratified by study center.

**Supplementary Figure 11.**
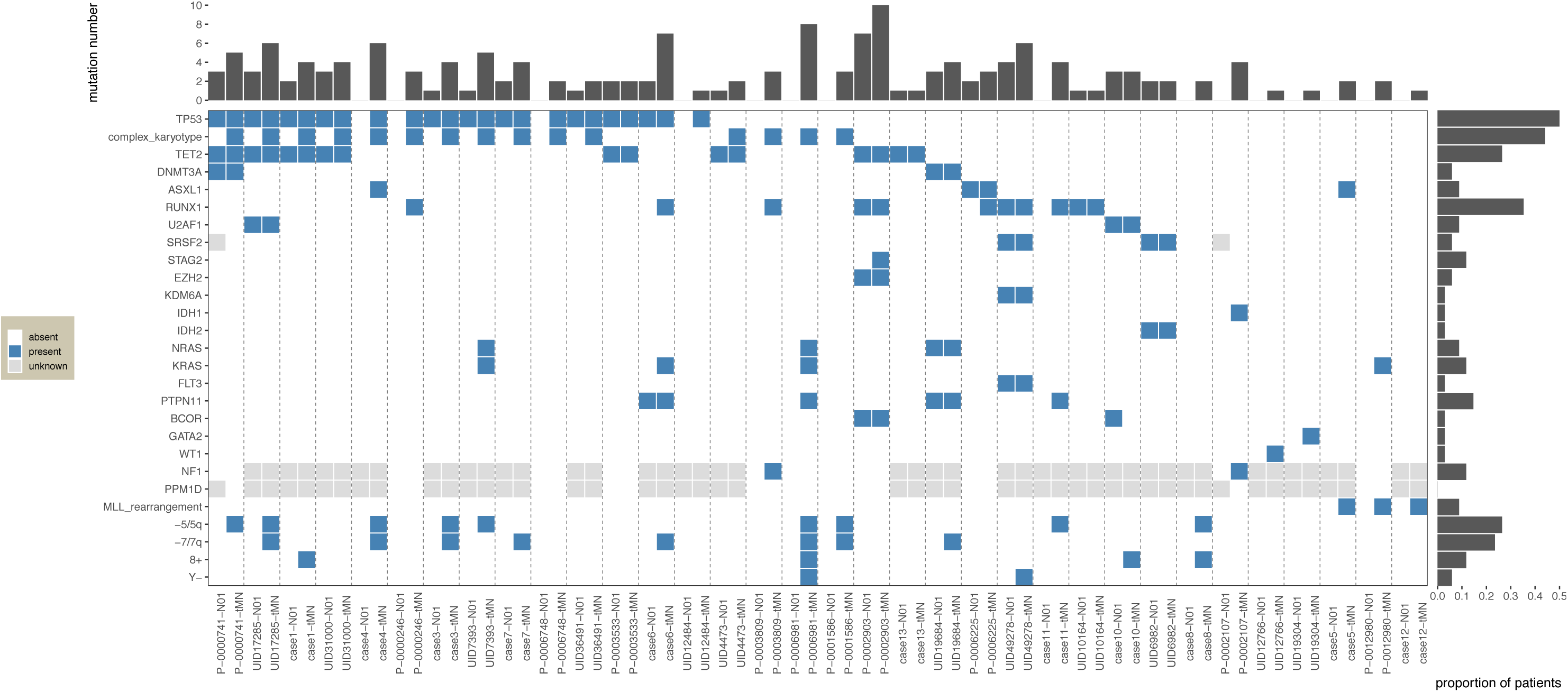
Mutation landscape in 34 tMN cases with at least one genetic alteration present at the time of tMN diagnosis. Pre-tMN sample denoted by “–N01” and sample attained at time of tMN diagnosis denoted by “-tMN”. Grey boxes represent genes not sequenced. Chromosomal abnormalities were not evaluated at the time of pre-tMN testing.

**Supplementary Figure 12.**
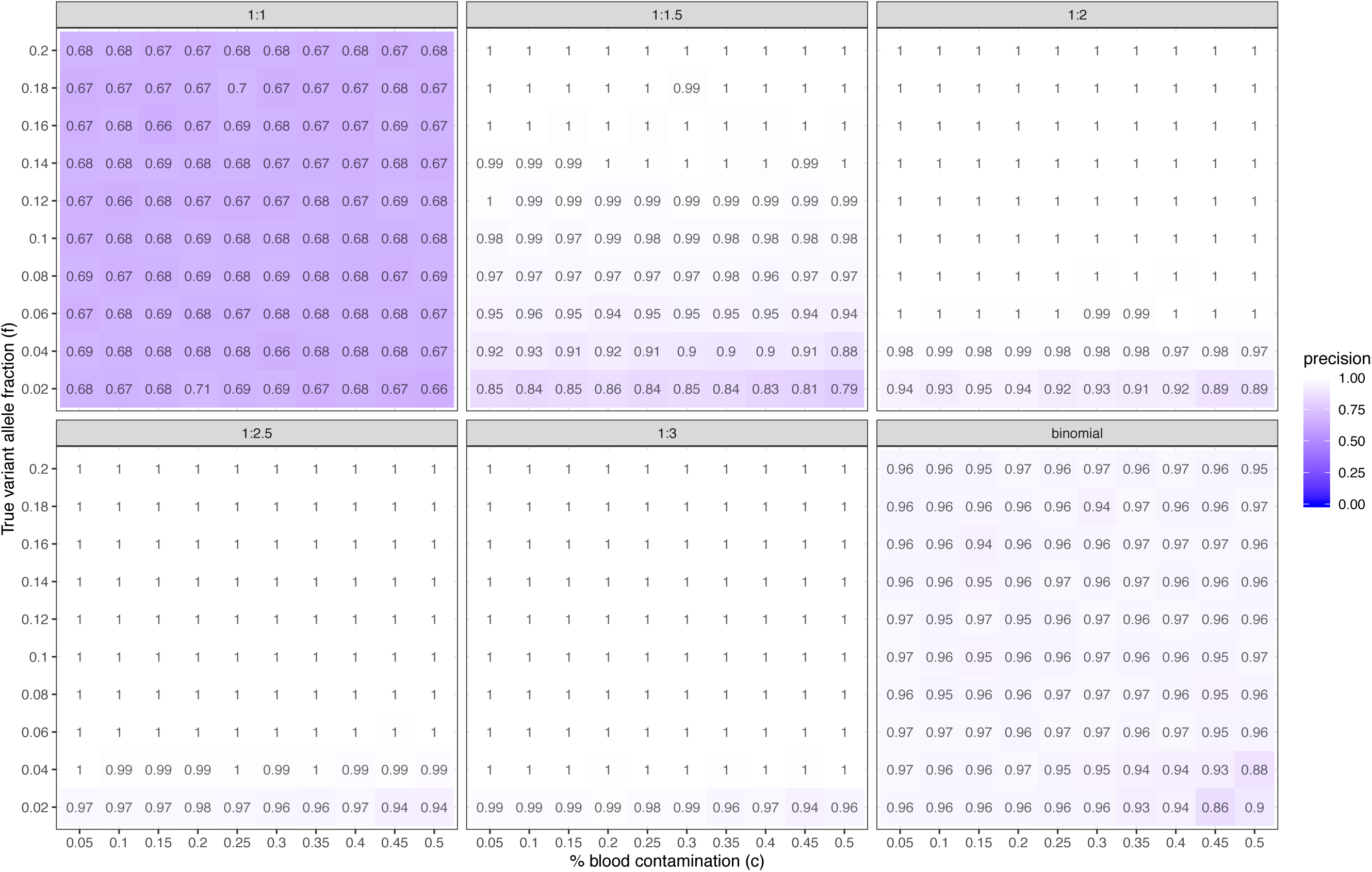
Precision of CH calling by simulation. Precision for discrimination of true CH calls from artifacts using a range of cutoffs (1:1, 1:1.5, 1:2, 1:2.5, 1:3) for the ratio of the VAF in the blood to the VAF in tumor ratio and a binomial test testing the null hypothesis for an equal VAF in the blood and the tumor.

**Supplementary Figure 13.**
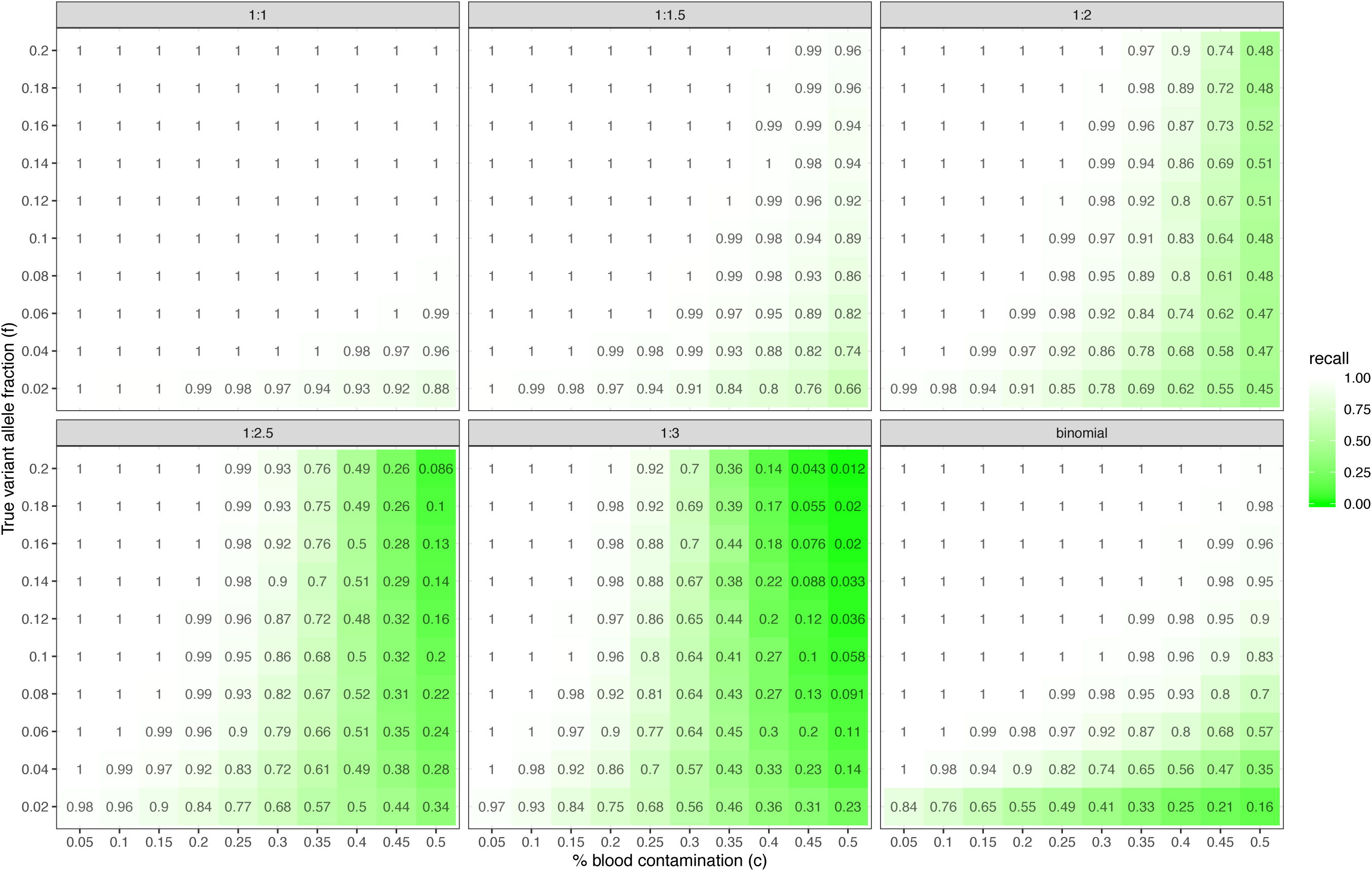
Recall of CH calling by simulation. Recall for discrimination of true CH calls from artifacts using a range of cutoffs (1:1, 1:1.5, 1:2, 1:2.5, 1:3) for the ratio of the VAF in the blood to the VAF in tumor ratio and a binomial test testing the null hypothesis for an equal VAF in the blood and the tumor.

