## Supplementary material for "Oncologic therapy shapes the fitness landscape of clonal hematopoiesis": R code for results

### Oncologic Therapy Shapes the Fitness Landscape of Clonal Hematopoeisis.rmd

#Tabulate main cohort numbers

```
print('All mutations')
```

```
## [1] "All mutations"
```

```
nrow(M_long)
```

```
## [1] 11391
```

```
print('# of Patients')
```

```
## [1] "# of Patients"
```

```
nrow(M_wide_all)
```

```
## [1] 24439
```

```
print('# of positive patients')
```

```
## [1] "# of positive patients"
```

```
unique(M_long$MRN) %>% length()
```

```
## [1] 7379
```

```
print('# of Patients CH positive')
```

```
## [1] "# of Patients CH positive"
```

```
sum(M_wide_all$CH_all)/nrow(M_wide_all)
```

```
## [1] 0.3019354
```

```
print('proportion of Patients CH positive')
```

```
## [1] "proportion of Patients CH positive"
```

```
sum(M_wide_all$CH_all)/nrow(M_wide_all)
```

```
## [1] 0.3019354
```

```
print('# of Patients with treatment available')
```

```
## [1] "# of Patients with treatment available"
```

```
nrow(M_wide)
```

```
## [1] 10207
```

```
print('# of CH-PD mutations')
```

```
## [1] "# of CH-PD mutations"
```

```
sum(M_long$ch_pancan_pd)
```

```
## [1] 6028
```

```
print('# of patients with CH-PD')
```

```
## [1] "# of patients with CH-PD"
```

```
M_wide_all %>% {sum(.$ch_pancan_pd)}
```

```
## [1] 4648
```

```
sum(M_wide_all$ch_pancan_pd)/nrow(M_wide_all)
```

```
## [1] 0.1901878
```

```
print('# of CH-my-PD mutations')
```

```
## [1] "# of CH-my-PD mutations"
```

```
sum(M_long$ch_my_pd)
```

```
## [1] 5453
```

```
print('# of patients with CH-MY-PD')
```

```
## [1] "# of patients with CH-MY-PD"
```

```
M_wide_all %>% {sum(.$ch_my_pd)}
```

```
## [1] 4316
```

```
print('table of mutations')
```

```
## [1] "table of mutations"
```

```
table(M_wide_all$mutnum_all_r)
```

```
##
##     0     1     2
## 17060  5044  2335
```

```
print('# of tumor types')
```

```
## [1] "# of tumor types"
```

```
unique(M_wide_all$generaltumortype) %>% length()
```

```
## [1] 57
```

```
print('# pts with therapy')
```

```
## [1] "# pts with therapy"
```

```
table(M_wide$therapy_binary)
```

```
##
## untreated   treated
##      3967      6240
```

#### Main figures

### Figure 1 - mutational characteristics

```
panel_theme = theme_bw() + theme(
    panel.border = element_blank(),
    legend.position = "none",
    panel.grid.minor = element_blank(),
    plot.subtitle = element_text(hjust = 0.5, size = 8),
    plot.title = element_text(face = 'bold', size = 12, hjust = 0, vjust = -11),
    panel.grid.major = element_blank(),
    strip.background = element_blank(),
    strip.text = element_text(size = 6),
    axis.text.y = element_text(size = 6),
    axis.text.x = element_text(size = 6),
    axis.title = element_text(size = 8),
    axis.line = element_line(),
    plot.margin = unit(c(0,0,0,0), 'pt')
)

age_groups = c("0-10", "11-20", "21-30", "31-40", "41-50", "51-60", "61-70", "71-80", "81-90", "91-100")

get_ch_grouped = function(M_wide, CI = T) {

    CH_by_age_grouped = M_wide %>% select(MRN, age_cat, CH) %>%
        mutate(CH = ifelse(is.na(CH), 0, CH)) %>%
        group_by(age_cat) %>%
        summarise(CH = sum(CH), total = n()) %>%
        filter(!is.na(age_cat)) %>%
        mutate(freq = CH / total)

    if (CI) {
        CH_by_age_grouped = CH_by_age_grouped %>%
        cbind(
            apply(CH_by_age_grouped, 1, function(row) {
                CI = prop.test(row['CH'], row['total'], conf.level=0.95)$conf.int[1:2]
                return(c(lower = CI[1], upper = CI[2]))
            }) %>% t
        )
    }

    return(CH_by_age_grouped)
}

font_size = 8
age_curve_theme =
  theme(
      legend.position = 'top',
      legend.key.size = unit(5, 'mm'),
      legend.title = element_blank(),
      legend.direction = 'horizontal',
      plot.title = element_text(hjust = -0.08),
      axis.text.x = element_text(angle = 45, vjust = 0.5, size = font_size),
      axis.text.y = element_text(size = font_size),
      axis.title = element_text(size = font_size),
      legend.text = element_text(size = font_size)
  )

## Age All
D = rbind(
    get_ch_grouped(M_wide %>% filter(therapy_binary == 'treated') %>% mutate(CH = CH_all)) %>%
        # mutate(therapy = paste0('treated', '\n(n = ', sum(total), ')')),
      mutate(therapy = 'treated'),
    get_ch_grouped(M_wide %>% filter(therapy_binary == 'untreated') %>% mutate(CH = CH_all)) %>%
        # mutate(therapy = paste0('untreated', '\n(n = ', sum(total), ')')) %>%
      mutate(therapy = 'untreated')
)

p_all = ggplot(
      D,
      aes(x = age_cat,
          y = freq,
          ymin = upper,
          ymax = lower,
          color = therapy,
          fill = therapy,
          group = therapy)
  ) +
  geom_ribbon(
      alpha = 0.2,
      colour = NA
  ) +
  geom_line() +
```

```
  scale_x_continuous(
      breaks = c(1, 2, 3, 4, 5, 6, 7, 8, 9, 10),
      labels = age_groups
  ) +
  panel_theme +
  age_curve_theme +
  xlab("Age") +
  ylab("Proportion of patients with CH") +
  theme(plot.title = element_text(hjust = -0.08)) +
  scale_fill_manual(values = therapy_colors) +
  scale_color_manual(values = therapy_colors)

## Age PD
D = rbind(
      get_ch_grouped(M_wide %>% filter(therapy_binary == 'treated') %>% mutate(CH = ch_pancan_pd)) %>%
          mutate(therapy = 'treated', CH = 'CH-PD'),
      get_ch_grouped(M_wide %>% filter(therapy_binary == 'treated') %>% mutate(CH = CH_all & !ch_pancan_pd)) %>%
          mutate(therapy = 'treated', CH = 'CH-non-PD'),
      get_ch_grouped(M_wide %>% filter(therapy_binary == 'untreated') %>% mutate(CH = ch_pancan_pd)) %>%
          mutate(therapy = 'untreated', CH = 'CH-PD'),
      get_ch_grouped(M_wide %>% filter(therapy_binary == 'untreated') %>% mutate(CH = CH_all & !ch_pancan_pd)) %
>%
          mutate(therapy = 'untreated', CH = 'CH-non-PD')
  ) %>%
  mutate_at(c('CH', 'therapy'), format_variable)

p_pd = ggplot(
      D,
      aes(x = age_cat,
          y = freq,
          ymin = upper,
          ymax = lower,
          color = CH,
          fill = CH,
          group = CH)
  ) +
  geom_ribbon(
      alpha = 0.2,
      colour = NA
  ) +
  geom_line() +
  scale_x_continuous(
      breaks = c(1, 2, 3, 4, 5, 6, 7, 8, 9, 10),
      labels = age_groups
  ) +
  xlab("Age") +
  ylab("") +
  panel_theme +
  age_curve_theme +
  theme(strip.text = element_text(size = font_size)) +
  facet_wrap(~therapy) +
  scale_fill_manual(values = c('#20854E99', '#E1872799')) +
  scale_color_manual(values = c('#20854E99', '#E1872799'))

## Histogram by gene frequency
gene_list = M %>% count(Gene) %>% arrange(-n) %>% .$Gene %>% unique %>% .[1:10]

n_treated = M_wide %>% count(therapy_binary) %>% filter(therapy_binary == 'treated') %>% pull(n)
n_untreated = M_wide %>% count(therapy_binary) %>% filter(therapy_binary == 'untreated') %>% pull(n)

# tally
D = M %>%
    filter(CH_nonsilent == 1) %>%
    reshape2::dcast(
        formula = Gene + therapy_binary ~ .,
        value.var = 'MRN',
        fun.aggregate = function(MRNs) {length(unique(MRNs))}
    ) %>%
    dplyr::rename("n_patient" = ".") %>%
    mutate(
        prop_patient = case_when(
            therapy_binary == 'treated' ~ n_patient/n_treated,
            therapy_binary == 'untreated' ~ n_patient/n_untreated
        )
    ) %>%
    filter(Gene %in% gene_list) %>%
    mutate(
        Gene = factor(Gene, gene_list),
        therapy_binary = factor(therapy_binary, c('untreated', 'treated'))
    ) %>%
```

```
    arrange(Gene)

source('../utils/toolbox.R')
# need to test for ch_nonsilent, the gene columns in M_wide are ch_pancan_pd
asterisks = lapply(gene_list,
    function(gene) {
        model = glm(
            formula = paste0(gene, ' ~ age_scaled + smoke_bin + race_b + Gender + therapy_binary'),
            data = M_wide,
            family = "binomial")
        treatment_pval = model %>% summary %$% coefficients %>% .['therapy_binarytreated', 'Pr(>|z|)']
        treatment_qval = p.adjust(treatment_pval, method = 'fdr', n = length(gene_list))
        return(signif.num(treatment_qval, ns = F))
    }
)

p_hist = ggplot(
      D,
      aes(x = Gene, y = prop_patient, fill = therapy_binary)
  ) +
  geom_bar(stat = 'identity', position = "dodge", color = 'black', size = 0.25) +
  panel_theme +
  theme(
      panel.grid.major = element_blank(),
      panel.border = element_blank(),
      axis.line = element_line(colour = "black"),
      legend.title = element_blank(),
      legend.key.size = unit(5, 'mm'),
      legend.position = 'top',
      legend.direction = 'horizontal',
      axis.title = element_text(size = font_size),
      axis.text.x = element_text(angle = 45, hjust = 1, size = font_size),
      legend.text = element_text(size = font_size)
  ) +
  annotate('text', x = gene_list, y = 0.11, label = asterisks, size = 4) +
  ylab("Proportion with mutated Gene") +
  xlab('') +
  scale_fill_manual(values = therapy_colors) +
  scale_color_manual(values = therapy_colors)

## stacked bar
mycols = c(
    brewer.pal(9, "YlOrBr") %>% rev %>% .[1:3] %>% rev,
    brewer.pal(9, "YlOrRd") %>% .[2:5],
    brewer.pal(9, "Blues") %>% rev
)

spliceosome_genes = c('SF3B1', 'SRSF2', 'U2AF1')

D = M %>%
    mutate(
      age_bins = case_when(
        age < 40 ~ 1,
        age < 65 ~ 2,
        age < 75 ~ 3,
        age >= 75 ~ 4)
    ) %>%
    group_by(age_bins) %>%
    mutate(age_bounds = paste0(round(min(age), 0), '-', round(max(age), 0))) %>%
    ungroup() %>%
    mutate(age_bins = age_bounds) %>%
    mutate(Gene = as.character(Gene)) %>%
    group_by(Gene) %>%
    mutate(freq = n()) %>%
    ungroup() %>%
    mutate(
        Gene = ifelse(freq > 50 | Gene %in% spliceosome_genes | Gene %in% tch_genes, Gene, 'Other')
    ) %>%
    arrange(Gene == 'Other', !(Gene %in% spliceosome_genes), !(Gene %in% tch_genes), freq) %>%
    mutate(Gene = factor(Gene, levels = unique(Gene))) %>%
    count(age_bins, therapy_binary, Gene) %>%
    group_by(age_bins, therapy_binary) %>%
    mutate(
        total = sum(n),
        prop = n/sum(n)
    ) %>%
    ungroup()

p_stack = ggplot(
      D,
```

```
      aes(x = age_bins, y = prop, fill = Gene)
  ) +
  geom_bar(stat = 'identity', size = 0) +
  panel_theme +
  theme(
    legend.title = element_blank(),
    panel.grid.major = element_blank(),
    panel.grid.minor = element_blank(),
    panel.border = element_blank(),
    axis.line = element_line(colour = "black"),
    strip.background = element_rect(fill = 'white', color = 'white'),
    axis.text.y = element_text(size = font_size),
    axis.text.x = element_text(size = font_size, angle = 45, hjust = 1),
    legend.text = element_text(size = font_size/1.5),
    legend.key.size = unit(3, "mm"),
    axis.title = element_text(size = font_size),
    legend.position = 'top'
  ) +
  facet_wrap(~therapy_binary) +
  ylab('Proportion of patients') +
  xlab('Age') +
  scale_fill_manual(values = mycols)

## forest plot
DTA = c('DNMT3A', 'TET2', 'ASXL1')
DDR = c('PPM1D', 'TP53', 'CHEK2')
SPL = c('SF3B1', 'SRSF2')
OTH = c('JAK2', 'ATM')

gene_list = c(DDR, DTA, SPL, OTH)

#ALL adjusted for treatment
logit_gene_var = list()

for (gene in gene_list) {
    logit = glm(
        formula = get(gene) ~ age_scaled + smoke_bin + race_b + Gender + therapy_binary,
        data = M_wide,
        family = "binomial")
    logit_data = logit %>% sjPlot::get_model_data() %>% cbind(Gene = gene)
    logit_gene_var = rbind(logit_gene_var, logit_data)
}

# add the interaction for each gene
D = logit_gene_var %>%
    filter(!term %in% c("GenderFemale", "race_b")) %>%
    mutate(
        term = c(
            'therapy_binarytreated' = 'Therapy',
            'smoke_bin1' = 'Smoking',
            'age_scaled' = 'Age'
        )[as.character(term)]
    ) %>%
    mutate(term = factor(term, c("Age", "Therapy", "Smoking"))) %>%
    mutate(p_fdr = p.adjust(p.value, method = "fdr")) %>%
    mutate(termGene = paste0(term, Gene)) %>%
    arrange(estimate, Gene) %>%
    mutate(termGene = factor(termGene, levels = termGene)) %>%
    mutate(gene_cat = case_when(
        Gene %in% DTA ~ 'DTA',
        Gene %in% DDR ~ 'DDR',
        Gene %in% SPL ~ 'Splicing',
        T ~ 'Other'
      )
    ) %>%
    mutate(gene_cat = factor(gene_cat, c('DDR', 'DTA', 'Splicing', 'Other'))) %>%
    mutate(
        q.value = p.adjust(p.value, n = nrow(.), method = 'fdr'),
        q.label = paste0(signif(estimate, 2), signif.num(q.value)),
        q.star = signif.num(q.value)
    )

p_forest = plot_forest(
      D,
      x = "termGene",
      label = 'q.star',
      eb_w = 0,
      eb_s = 0.3,
      ps = 1.5,
      or_s = 2,
```

```
      nudge = -0.3,
      col = 'gene_cat'
  ) +
  facet_wrap(~term, scale = 'free_y', ncol = 1) +
  scale_x_discrete(
      breaks = D$termGene,
      labels = D$Gene,
      expand = c(0.1,0)
  ) +
  xlab('') + ylab('OR') +
  scale_color_nejm() +
  panel_theme +
  theme(
    axis.text = element_text(size = font_size),
    axis.title = element_text(size = font_size),
    strip.text = element_text(size = font_size),
    legend.position = 'top',
    legend.title = element_blank(),
    legend.text = element_text(size = font_size/1.2),
    legend.key.size = unit(3, "mm")
  )

combo = ((((p_all + labs(title = 'A')| p_pd + labs(title = 'B')) + plot_layout(widths = c(1, 2))) / ((p_hist + la
bs(title = 'C') | p_stack + labs(title = 'D')) + plot_layout(widths = c(1.5, 1)))) + plot_layout(heights = c(1,1.
5)) | p_forest + labs(title = 'E')) + plot_layout(widths=c(3, 1))

do_plot(combo, "fig1.png", 10, 6, save_pdf = T)
```

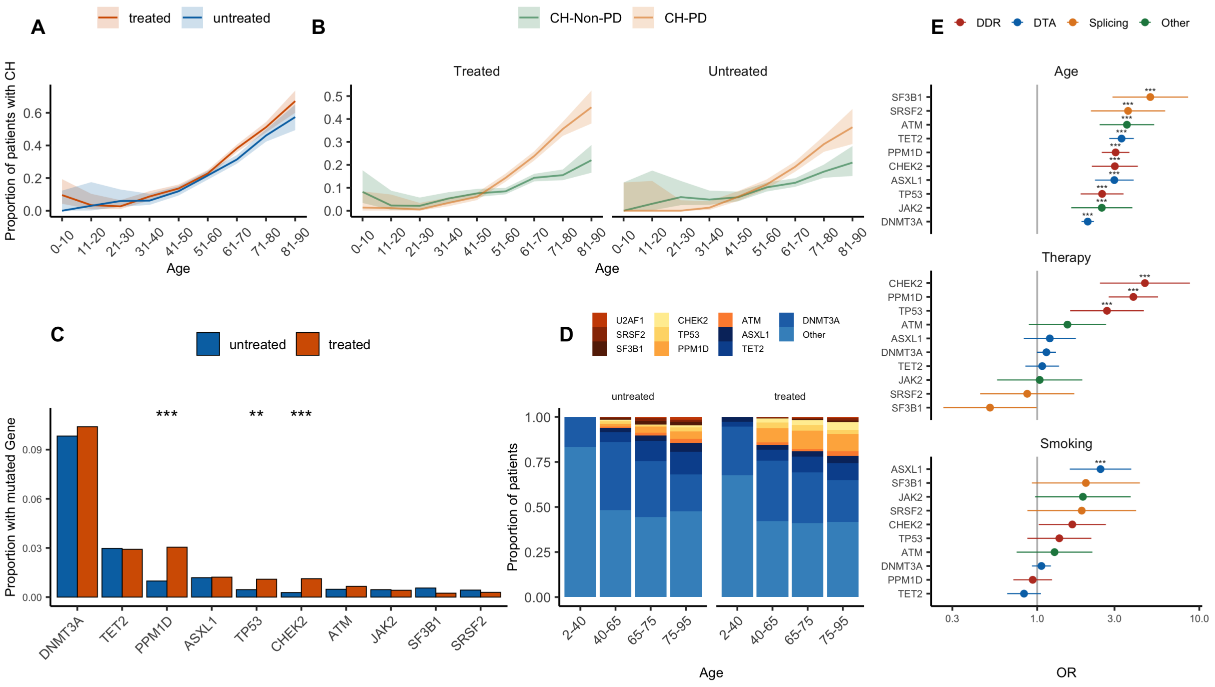


#Figure 2 - therapy and genes

```
#Plotting theme
panel_theme = theme_bw() + theme(
    panel.border = element_blank(),
    legend.position = "none",
    panel.grid.minor = element_blank(),
    plot.subtitle = element_text(hjust = 0.5, size = 8),
    plot.title = element_text(face = 'bold', hjust = 0, vjust = -2, size = 12),
    panel.grid.major = element_blank(),
    strip.background = element_blank(),
    strip.text = element_text(size = 6),
    axis.text.y = element_text(size = 6),
    axis.text.x = element_text(size = 6),
    axis.title = element_text(size = 8),
    axis.line = element_line(),
    plot.margin = unit(c(0,0,0,0), 'pt')
)

interest_drugs = c("XRT", "ind_platinum",
    "ind_topoisomerase_ii_inh", "ind_taxane", "ind_topoisomerase_i_inhi",
    "ind_antimetabolite", "ind_microtubule_damaging", "ind_radiotherapy")

interest_genes = c("PPM1D","TP53","CHEK2","DNMT3A","TET2","ASXL1","ATM")

D = lapply(
        interest_genes,
        function (gene) {

            terms = filter_terms(
                M_wide %>% filter(get(gene) == 1),
                interest_drugs,
                verbose = F,
                threshold = 10
            )

            if (length(terms) == 0) {
                return(NULL)
            }

            formula = paste0(
                gene, ' ~ ',
                paste(terms, collapse = ' + '),
                ' + XRT + age + smoke_bin + race_b + timedx_impact')

            glm(formula = formula, data = M_wide, family = "binomial", na.action = 'na.omit') %>%
                sjPlot::get_model_data() %>% cbind(Gene = gene)
        }
    ) %>%
    Reduce(rbind, .) %>%
    dplyr::filter(term %in% c(interest_drugs) & Gene %in% interest_genes) %>%
    mutate(
        term = factor(format_variable(term), format_variable(interest_drugs)),
        Gene = factor(Gene, interest_genes)
    ) %>%
    mutate(p_fdr = p.adjust(p.value, method = "fdr")) %>%
    rowwise() %>%
    mutate(log_or = log(estimate)) %>%
    ungroup()

p_heatmap = ggplot(
        D,
        aes(x = term, y = Gene, fill = log_or)
    ) +
    geom_tile(color = "lightgrey") +
    scale_fill_gradient2(
        low = "darkblue",
        mid = "white",
        high = "darkred",
        midpoint = 0,
        na.value = "white",
        limits = c(-2.5, 2.5)

    ) +
    geom_point(
        aes(size = ifelse(p_fdr <= 0.05, 1, NA)),
        shape = 8,
        colour = "black",
        show.legend = F
    ) +
    scale_size_area(max_size = 2) +
```

```
    xlab('') +
    ylab('') +
    panel_theme +
    scale_y_discrete(expand = c(0,0)) +
    scale_x_discrete(expand = c(0,0)) +
    theme(
        panel.grid.minor = element_blank(),
        panel.grid.major = element_blank(),
        axis.line = element_blank(),
        legend.position = 'right',
        axis.text.x = element_text(size = 8, angle = 30, hjust = 1),
        plot.margin = unit(c(20, 0, 20, 0), 'pt'),
        legend.text = element_text(size = 4),
        legend.title = element_text(size = 4),
        legend.key.size = unit(10, "pt")
    )

ggsave("p_heatmap.png", plot = p_heatmap, width = 4, height = 4, dpi = 300)
```

```
## Warning: Removed 32 rows containing missing values (geom_point).
```

```
knitr::include_graphics("p_heatmap.png")
```

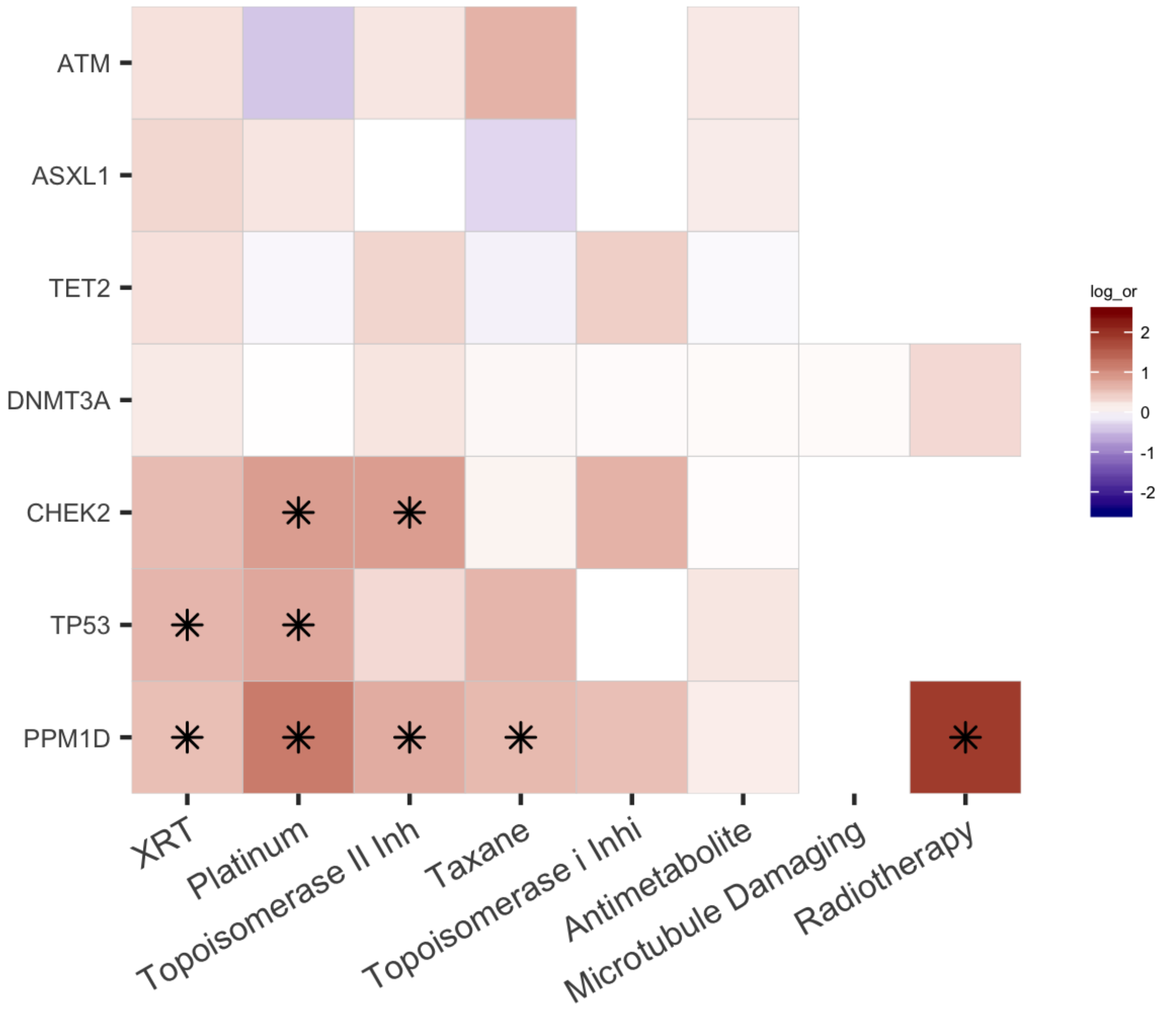

```
## Systemic therapies ##

therapy_labels = c(
    "ind_immune_therapy" = "Immune Therapy",
    "ind_targeted_therapy" = "Targeted Therapy",
    "ind_cytotoxic_therapy" = "Cytotoxic Therapy",
    "XRT" = "External Beam Radiation",
    "ind_radiotherapy" = "Radionuclide")

n_labels = M_wide %>%
    summarise_at(
        names(therapy_labels),
        term_count
    ) %>% t %>%
    as.data.frame %>%
    tibble::rownames_to_column('term') %>%
    dplyr::rename(n = V1)

formula = paste0(
    'ch_pancan_pd', ' ~ ',
    paste(
    'timedx_impact',
    'ind_cytotoxic_therapy',
    'ind_immune_therapy',
    'ind_radiotherapy',
    'ind_targeted_therapy',
    'XRT',
    'age',
    'smoke_bin',
    'race_b', sep = ' + ')
)

summary_systemic = M_wide %>%
    glm(
        formula = formula,
        family = binomial(link="logit"),
        na.action = 'na.omit'
    ) %>%
    sjPlot::get_model_data(model = .) %>%
    left_join(
        n_labels,
        by = 'term'
    ) %>%
    mutate(term = as.character(term)) %>%
    mutate(term = ifelse(term %in% names(therapy_labels), therapy_labels[term], term)) %>%
    filter(!str_detect(term, regex("age|smoke_bin1|race_b|timedx_impact"))) %>%
    mutate(term = paste0(term, ' (n = ', n, ')')) %>%
    mutate(term = format_variable(term)) %>%
    arrange(estimate) %>%
    mutate(term = factor(term, levels = unique(term)))
```

```
## Warning: Column `term` joining factor and character vector, coercing into
## character vector
```

```
## Cytotoxic therapies ##

formula = paste0(
    'ch_pancan_pd ~ ',
    paste(
        'ind_taxane',
        'ind_microtubule_damaging',
        'ind_antimetabolite',
        'ind_alkylating_agent',
        'ind_platinum',
        'ind_topoisomerase_ii_inh',
        'ind_topoisomerase_i_inhi',
        'ind_cytotoxic_therapy_ot',
        'XRT',
        'ind_targeted_therapy',
        'ind_immune_therapy',
        'age',
        'smoke_bin',
        'race_b',
        'timedx_impact',
        sep = ' + '))

drug_list = c(
    "ind_taxane",
    "ind_microtubule_damaging",
    "ind_antimetabolite",
    "ind_alkylating_agent",
    "ind_platinum",
    "ind_topoisomerase_ii_inh",
    "ind_topoisomerase_i_inhi")

n_labels = M_wide %>%
    summarise_at(
        drug_list,
        term_count
    ) %>% t %>%
    as.data.frame %>%
    tibble::rownames_to_column('term') %>%
    dplyr::rename(n = V1)

summary_cytotoxic = M_wide %>%
    glm(formula = formula, family = binomial(link = "logit"), na.action = 'na.omit') %>%
    sjPlot::get_model_data(.) %>%
    left_join(
        n_labels,
        by = 'term'
    ) %>%
    mutate(term = paste0(term, ' (n = ', n, ')')) %>%
    filter(str_detect(term, paste(drug_list, collapse = '|'))) %>%
    mutate(term = format_variable(term)) %>%
    mutate(term = factor(term, levels = unique(term[order(estimate)]))) %>%
    filter(std.error < 100)
```

```
## Warning: Column `term` joining factor and character vector, coercing into
## character vector
```

```
## Platinum ##

formula = paste0(
    'ch_pancan_pd ~ ',
    paste(
        'ind_taxane',
        'ind_microtubule_damaging',
        'ind_antimetabolite',
        'ind_alkylating_agent',
        'ind_carboplatin',
        'ind_cisplatin',
        'ind_oxaliplatin',
        'ind_topoisomerase_ii_inh',
        'ind_topoisomerase_i_inhi',
        'ind_radiotherapy',
        'ind_targeted_therapy',
        'ind_immune_therapy',
        'XRT',
        'age',
        'smoke_bin',
        'race_b',
        'timedx_impact',
        sep = ' + '))

drug_list = c('ind_carboplatin','ind_cisplatin','ind_oxaliplatin')

n_labels = M_wide %>%
    summarise_at(
        drug_list,
        term_count
    ) %>% t %>%
 as.data.frame %>%
   tibble::rownames_to_column('term') %>%
    dplyr::rename(n = V1)

summary_platinum = M_wide %>%
    glm(
        formula = formula,
        family = binomial(link="logit"),
        na.action = 'na.omit'
    ) %>%
    sjPlot::get_model_data(.) %>%
    filter(str_detect(term, regex("carboplatin|cisplatin|oxaliplatin", ignore_case = T))) %>%
    left_join(
        n_labels,
        by = 'term'
    )  %>%   mutate(term = paste0(term, ' (n = ', n, ')')) %>%
    filter(str_detect(term, paste(drug_list, collapse = '|'))) %>%
    mutate(term = as.character(term)) %>%
    mutate(term = format_variable(term)) %>%
    arrange(estimate) %>%
    mutate(term = factor(term, levels = unique(term)))
```

```
## Warning: Column `term` joining factor and character vector, coercing into
## character vector
```

```
summary_platinum %>%
plot_forest(
    x = "term",
    eb_w = 0,
    eb_s = 2,
    ps = 3,
    or_s = 4
) +
xlab('') +
ylab('OR') +
scale_color_manual(values = pal_nejm()(4)[3:4])
```

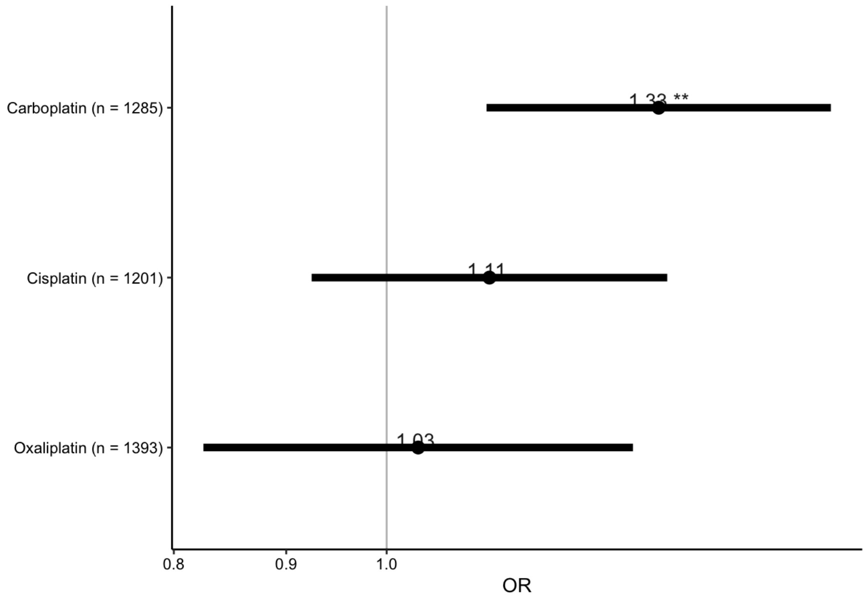


```
## Combined Plot ##

p_forest = rbind(
        summary_systemic %>% mutate(group="Systemic Therapy"),
        summary_cytotoxic %>% mutate(group="Cytotoxic Therapy"),
        summary_platinum %>% mutate(group="Platinum")
    ) %>%
    mutate(group = factor(group, c('Systemic Therapy', 'Cytotoxic Therapy', 'Platinum'))) %>%
    arrange(group, estimate) %>%
    mutate(term = factor(term, levels = unique(term))) %>%
    plot_forest(
        x = "term",
        eb_w = 0,
        eb_s = 1.5,
        label="p.stars",
        ps = 2,
        or_s = 3) +
    ylab('OR') +
    facet_wrap(~group, scales = "free_y", ncol = 1) +
    panel_theme +
    # theme_classic() +
    theme(
        strip.placement = 'bottom',
        strip.text = element_text(size = 8),
        plot.margin = unit(c(0,10,0,0), 'pt'),
        axis.text.y = element_text(size = 6)
    ) +
    scale_fill_nejm() +
    scale_color_nejm() +
    ylab('OR') +
    xlab('') +
    scale_x_discrete(expand = c(0.25,0))

g_forest = ggplot_build(p_forest)
gt = ggplot_gtable(g_forest)
panels = gt$layout$t[grepl("panel", gt$layout$name)]
vtiles = c(5,7,3)
gt$heights[panels] <- unit(vtiles, "null")
p_forest = as.ggplot(gt)
p_forest
```

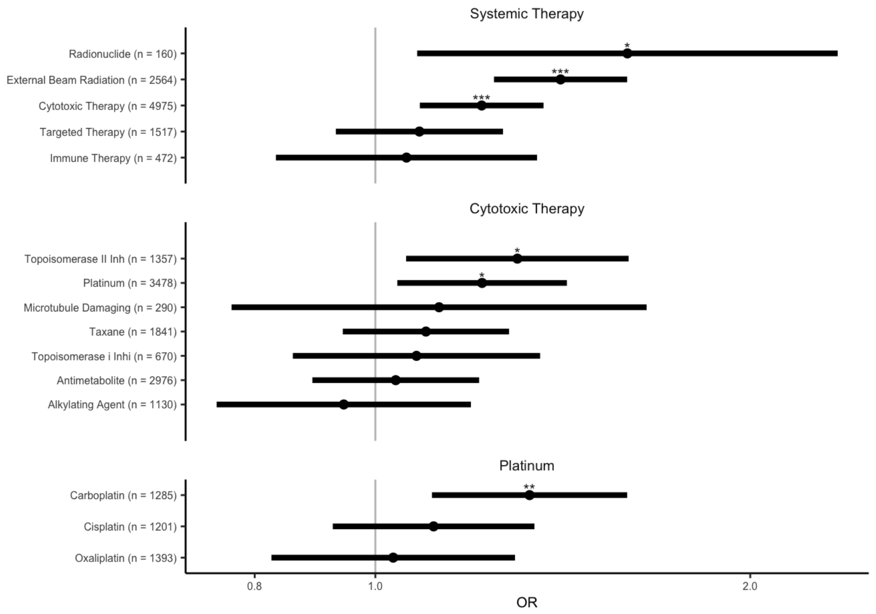


```
ggsave("p_forest.png", plot = p_forest, width = 4, height = 4, dpi = 300)

knitr::include_graphics("p_forest.png")
```

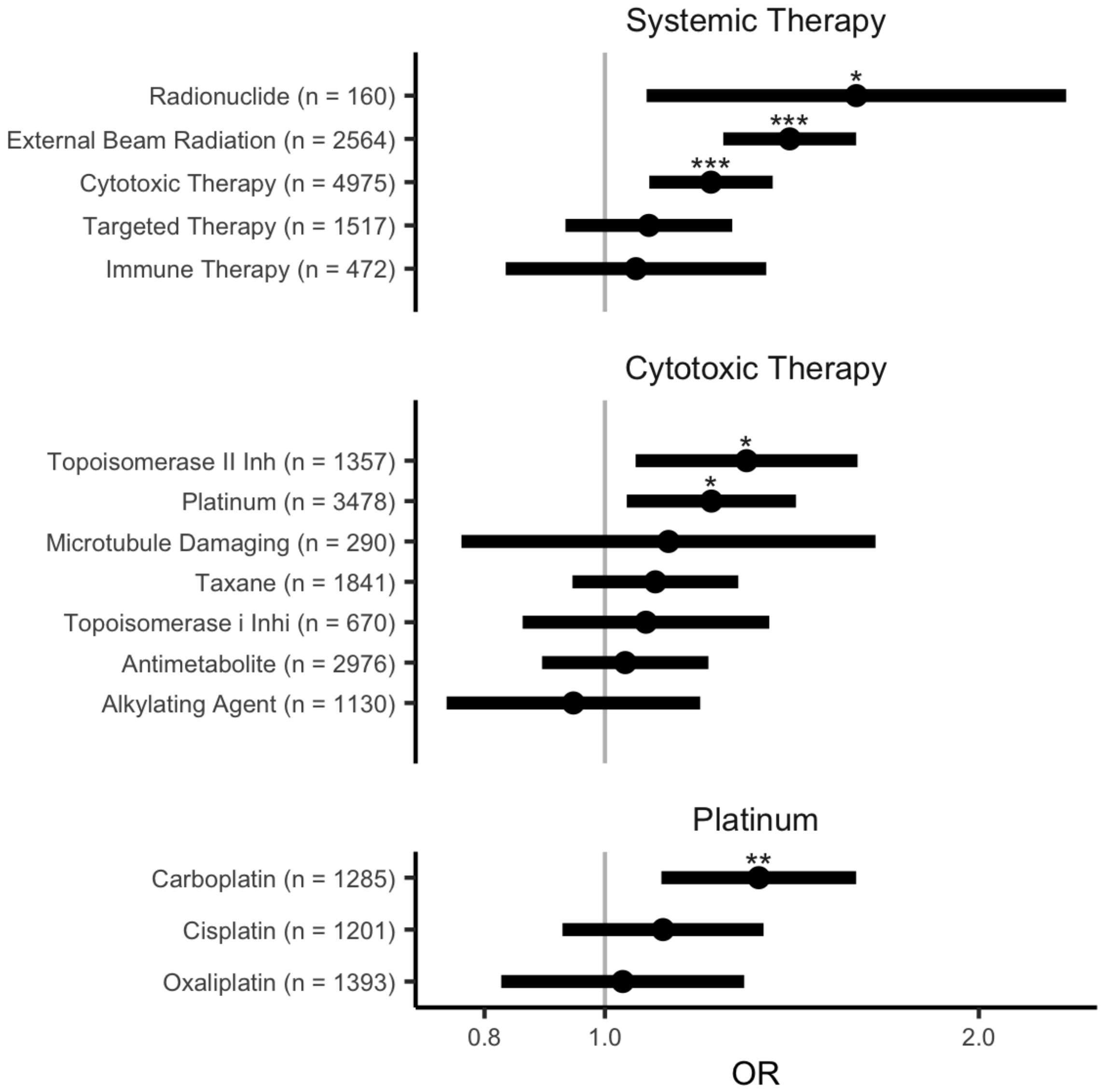
Systemic Therapy
Radionuclide (n = 160)
External Beam Radiation (n = 2564)
Cytotoxic Therapy (n = 4975)
Targeted Therapy (n = 1517)
Immune Therapy (n = 472)
*
***
***
Cytotoxic Therapy
Topoisomerase II Inh (n = 1357)
Platinum (n = 3478)
Microtubule Damaging (n = 290)
Taxane (n = 1841)
Topoisomerase i Inhi (n = 670)
Antimetabolite (n = 2976)
Alkylating Agent (n = 1130)
*
*
Platinum
Carboplatin (n = 1285)
Cisplatin (n = 1201)
Oxaliplatin (n = 1393)
**
0.8
1.0
2.0
OR

```
M_wide = M_wide %>% mutate(ch_pancan_pd = ch_pancan_pd | ch_tch)

formula = paste0(
    'ch_pancan_pd ~ ',
    paste(
        'pct_taxane',
        'pct_microtubule_damaging',
        'pct_antimetabolite',
        'pct_alkylating_agent',
        'pct_platinum',
        'pct_topoisomerase_ii_inh',
        'pct_topoisomerase_i_inhi',
        'pct_cytotoxic_therapy_ot',
        'pct_targeted_therapy',
        'pct_immune_therapy',
        'eqd_3_t',
        'age',
        'smoke_bin',
        'race_b',
        'timedx_impact',
        sep = ' + '))

# p vals for trend
pvals = c(
    'pct_platinum' = M_wide %>%
        filter(pct_platinum >= 1) %>%
        glm(formula = formula,
            family = binomial(link = "logit"),
            na.action = 'na.omit') %>%
        sjPlot::get_model_data(.) %>%
        filter(term == 'pct_platinum') %$%
        p.value,
    'pct_topoisomerase_ii_inh' = M_wide %>%
        filter(pct_topoisomerase_ii_inh >= 1) %>%
        glm(formula = formula, family = binomial(link = "logit"), na.action = 'na.omit') %>%
        sjPlot::get_model_data(.) %>%
        filter(term == 'pct_topoisomerase_ii_inh') %$%
        p.value,
    'eqd_3_t' = M_wide %>%
        filter(eqd_3_t >= 1) %>%
        glm(formula = formula, family = binomial(link = "logit"), na.action = 'na.omit') %>%
        sjPlot::get_model_data(.) %>%
        filter(term == 'eqd_3_t') %$%
        p.value
)

summary_class_b = M_wide %>%
    mutate_at(
        c('pct_taxane', 'pct_microtubule_damaging', 'pct_antimetabolite',
          'pct_alkylating_agent', 'pct_platinum', 'pct_topoisomerase_ii_inh',
          'pct_topoisomerase_i_inhi', 'eqd_3_t'),
        factor
    ) %>%
    glm(formula = formula, family = binomial(link="logit"), na.action = 'na.omit') %>%
    sjPlot::get_model_data(.) %>%
    dplyr::rename(Drug = term) %>%
    filter(str_detect(Drug, 'pct_topoisomerase_ii_inh|pct_platinum|eqd_3_t')) %>%
    mutate(
        Dosage = str_extract(Drug, '[0-9]$'),
        Drug = str_replace(Drug, '[0-9]$', '')
    ) %>%
    mutate(
        Dosage = factor(Dosage, levels = sort(unique(Dosage), decreasing = F))
    ) %>%
    mutate(
        Drug = ifelse(
            Drug %in% names(pvals),
            paste0(Drug, '\n(p = ', signif(pvals[Drug], 2), ')'),
            Drug)
    ) %>%
    mutate(Drug = format_variable(Drug))

## Platinum ##

formula = paste0(
    'ch_pancan_pd ~ ',
    paste(
        'pct_taxane',
        'pct_microtubule_damaging',
```

```
        'pct_antimetabolite',
        'pct_alkylating_agent',
        'pct_carboplatin',
        'pct_cisplatin',
        'pct_oxaliplatin',
        'pct_topoisomerase_ii_inh',
        'pct_topoisomerase_i_inhi',
        'pct_cytotoxic_therapy_ot',
        'pct_targeted_therapy',
        'pct_immune_therapy',
        'eqd_3_t',
        'age',
        'smoke_bin',
        'race_b',
        'timedx_impact',
        sep = ' + '))

drug_list = c('pct_carboplatin', 'pct_cisplatin', 'pct_oxaliplatin')

# p values for trend
pvals = sapply(
    drug_list,
    function(drug) {
        pval = M_wide %>% filter(!!drug > 0) %>%
        glm(formula = formula, family = binomial(link = "logit"), na.action = 'na.omit') %>%
        sjPlot::get_model_data(.) %>%
        filter(term == drug) %$%
        p.value %>%
        signif(2)
    }
)

summary_platinum = M_wide %>%
    mutate_at(
        drug_list,
        factor
    ) %>%
    glm(formula = formula, family = binomial(link = "logit"), na.action = 'na.omit') %>%
    sjPlot::get_model_data(.) %>%
    dplyr::rename(Drug = term) %>%
    mutate(
        Dosage = str_extract(Drug, '[0-9]'),
        Drug = str_replace(Drug, '[0-9]', '')
    ) %>%
    mutate(Dosage = factor(Dosage, levels = sort(unique(Dosage), decreasing = T))) %>%
    filter(Drug %in% drug_list) %>%
    mutate(
        Drug = ifelse(
            Drug %in% names(pvals),
            paste0(Drug, '\n(p = ', signif(pvals[Drug], 2), ')'),
            Drug)
    ) %>%
    mutate(Drug = format_variable(Drug)) %>%
    mutate(Drug = factor(Drug, levels = unique(Drug)))

p_dose = ggplot(
        rbind(summary_class_b, summary_platinum) %>%
            mutate(Drug = factor(Drug, unique(Drug))),
        aes(x = Dosage, y = estimate, group = 1)
    ) +
    geom_point() +
    geom_line() +
    ylab("OR of CH-PD") +
    scale_x_discrete(expand = c(0.1,0)) +
    xlab("Tertile of Cumulative Therapy Dose") +
    geom_ribbon(aes(ymin=conf.low, ymax=conf.high, x=Dosage, fill = ""), alpha = 0.3) +
    facet_wrap(~Drug, nrow = 2, scale = 'free') +
    theme_classic() +
    panel_theme +
    theme(
       legend.position = 'none'
    )

ggsave("p_dose.png", plot = p_dose, width = 4, height = 2, dpi = 300)

knitr::include_graphics("p_dose.png")
```

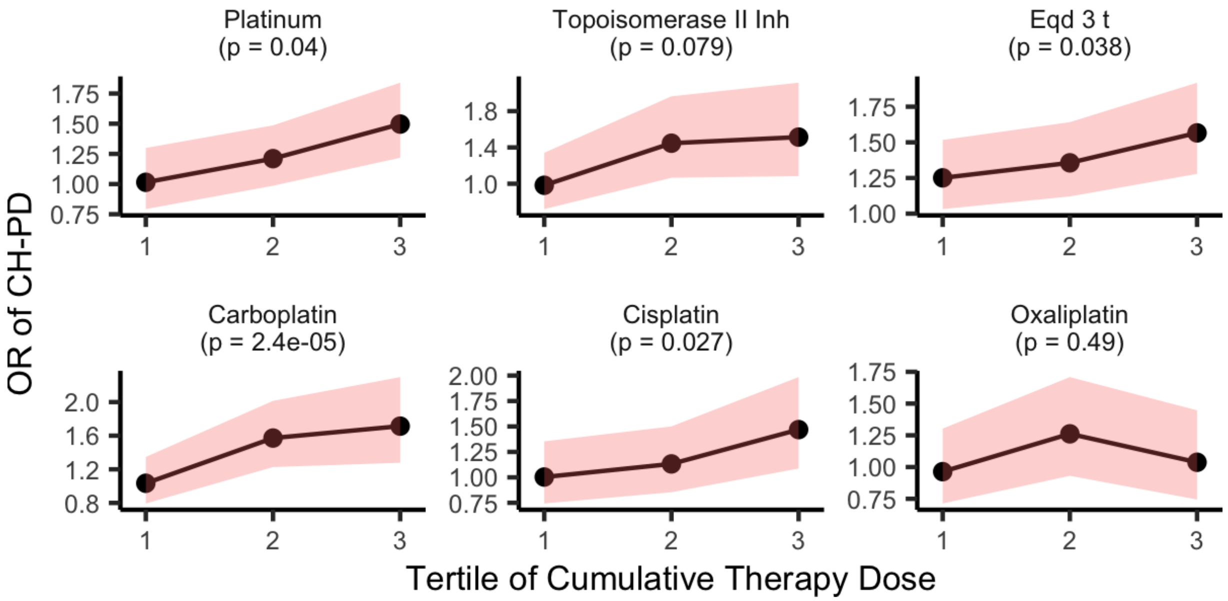


```
# p_forest + labs(title = 'a') + (p_dose + labs(title = 'b') / p_heatmap + labs(title = 'c')) + plot_layout(width
s = c(1.2, 1, 3))
(p_forest + (p_dose / p_heatmap + plot_layout(heights = c(2, 5)))) + plot_layout(widths = c(3, 3))
```

```
## Warning: Removed 32 rows containing missing values (geom_point).
```

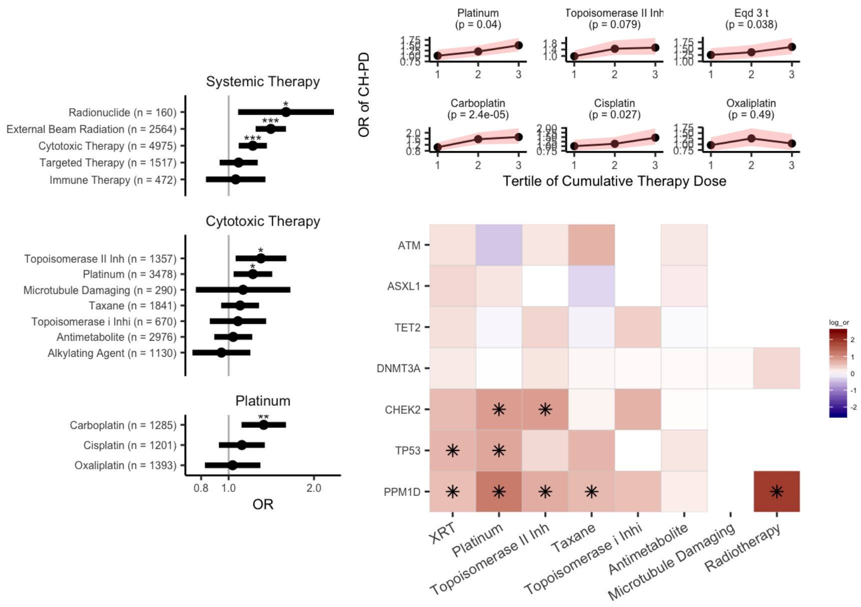


```
# panel = p_forest + labs(title = 'a') + p_dose + labs(title = 'b') + p_heatmap + labs(title = 'c') + plot_layout
(widths = c(1.2, 1, 3))
panel = (p_forest + labs(title = 'a') | ((p_dose + labs(title = 'b'))/ (p_heatmap + labs(title = 'c')) + plot_lay
out(heights = c(3,5)))) + plot_layout(widths = c(3, 2))

# panel
ggsave("figure2.png", plot = panel, width = 8, height = 5, dpi = 300)
```

```
## Warning: Removed 32 rows containing missing values (geom_point).
```

```
knitr::include_graphics("figure2.png")
```

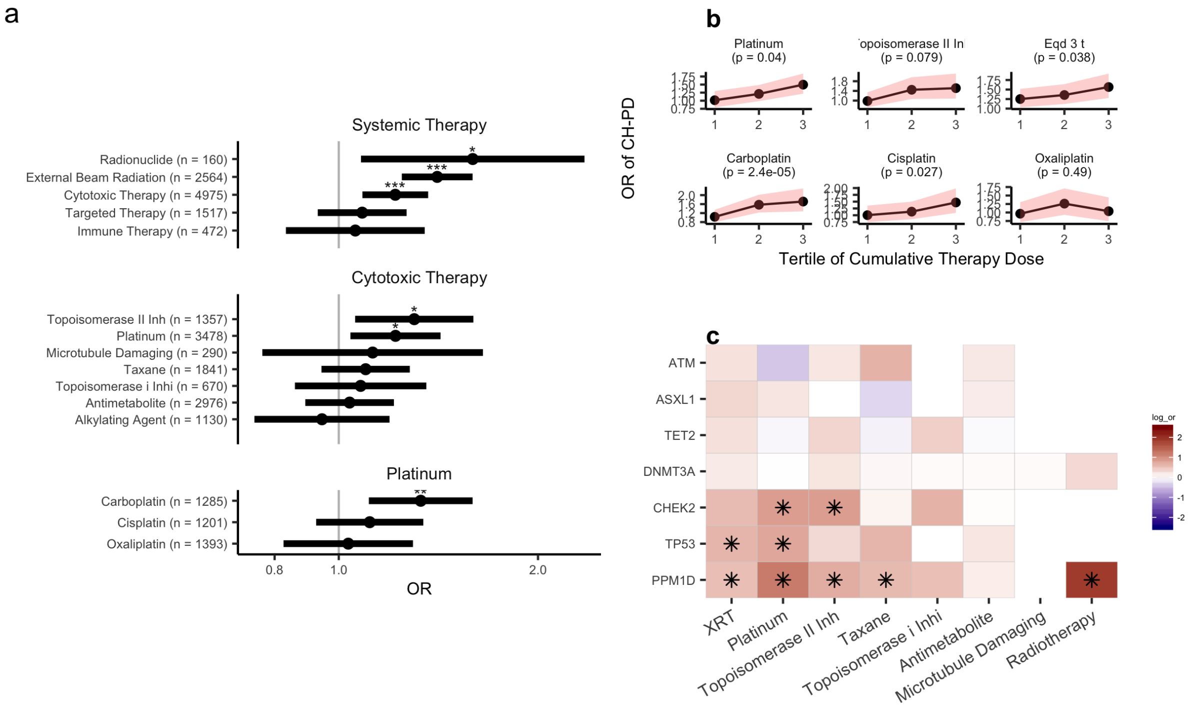


#Figure 3 - clonal dynamics

```
# delta vaf significance
M2['pval_T1_vs_T2'] <-
  apply(M2, 1, function(mut) {

    refs_1 = as.integer(mut['refs_1'])
    alts_1 = as.integer(mut['alts_1'])
    refs_2 = as.integer(mut['refs_2'])
    alts_2 = as.integer(mut['alts_2'])

    # if 0 in Serial, then CI def don't overlap
    if (refs_2 + alts_2 == 0 | refs_1 + alts_1 == 0) {
      p = 0
    } else if (alts_1 < 15 | alts_2 < 15) {
      p = data.frame(
        x = c(alts_1, alts_2),
        n = c(refs_1 + alts_1, refs_2 + alts_2),
        row.names = c('N', 'T')
      ) %>% fisher.test %>% .$p.value
    } else {
      p = prop.test(c(alts_1, alts_2), c(refs_1 + alts_1, refs_2 + alts_2),
                    conf.level = 0.95, correct = FALSE)$p.value
    }

    return(p)
  })

# grouping based on p value
M2 = M2 %>% mutate(direction = ifelse(pval_T1_vs_T2 < 0.05, ifelse(VAF_2 > VAF_1, 'UP', 'DOWN'), 'STABLE'))
```

```
font_size = 8

panel_theme = theme_bw() + theme(
    panel.border = element_blank(),
    legend.position = "none",
    panel.grid.minor = element_blank(),
    plot.subtitle = element_text(hjust = 0.5, size = font_size),
    plot.title = element_text(face = 'bold', size = 12, hjust = 0, vjust = -5),
    axis.title = element_text(size = font_size),
    panel.grid.major = element_blank(),
    strip.background = element_blank(),
    strip.text = element_text(size = font_size),
    axis.text.y = element_text(size = font_size),
    axis.line = element_line(),
    plot.margin = unit(c(0,0,0,0), 'pt')
)

therapy_colors = c('#D55E00', '#0072B2')

## Other vs DDR
D = M2 %>%
    filter(tch) %>%
    melt(
        measure.vars = c('XRT1', 'ind_cytotoxic_therapy1'),
        variable.name = 'treatment_type',
        value.name = 'treatment'
    ) %>%
    mutate(treatment_type = c('XRT1' = 'XRT', 'ind_cytotoxic_therapy1' = 'Cytotoxic')[treatment_type]) %>%
    mutate(treatment = recode(treatment, `1` = 'treated', `0` = 'untreated')) %>%
    mutate(treatment = factor(treatment, levels = c('treated', 'untreated')))

jitter = geom_jitter(
        pch = 21,
        size = 1,
        alpha = 0.8,
        width = 0.1,
        color = 'black',
        fill = 'grey'
        )

p_tch = ggplot(
      D,
      aes(x = treatment, y = log(growth_rate), fill = treatment)) +
  geom_boxplot(outlier.alpha = 0, color = 'black') +
  theme(plot.title = element_text(hjust = 0.5)) +
  xlab("") +
  ylab('log growth rate') +
  jitter +
  facet_wrap(~treatment_type) +
  panel_theme +
  theme(
    axis.text.x = element_blank()
  ) +
  scale_y_continuous(
      expand = c(0.1, 0),
      limits = c(-0.006, 0.006)
  ) +
  stat_compare_means(
    aes(label = paste0("p = ", ..p.format..)),
    method = 't.test',
    comparisons = list(c('treated', 'untreated')),
    size = 2.5
  ) +
  labs(subtitle = 'DDR') +
  scale_fill_manual(values = therapy_colors)
```

```
## Warning: Using `as.character()` on a quosure is deprecated as of rlang 0.3.0.
## Please use `as_label()` or `as_name()` instead.
## This warning is displayed once per session.
```

```
D = M2 %>%
    filter(!tch) %>%
    melt(
        measure.vars = c('XRT1', 'ind_cytotoxic_therapy1'),
        variable.name = 'treatment_type',
        value.name = 'treatment'
    ) %>%
    mutate(treatment_type = c('XRT1' = 'XRT', 'ind_cytotoxic_therapy1' = 'Cytotoxic')[treatment_type]) %>%
    mutate(treatment = recode(treatment, `1` = 'treated', `0` = 'untreated')) %>%
    mutate(treatment = factor(treatment, levels = c('treated', 'untreated')))

p_other = ggplot(
      D,
      aes(x = treatment, y = log(growth_rate), fill = treatment)) +
  geom_boxplot(outlier.alpha = 0, color = 'black') +
  theme(plot.title = element_text(hjust = 0.5)) +
  xlab("") +
  ylab('') +
  jitter +
  facet_wrap(~treatment_type) +
  panel_theme +
  theme(
  axis.text.x = element_blank()
  ) +
  scale_y_continuous(
      expand = c(0.1, 0),
      limits = c(-0.006, 0.006)
  ) +
  stat_compare_means(
    aes(label = paste0("p = ", ..p.format..)),
    method = 't.test',
    comparisons = list(c('treated', 'untreated')),
    size = 2.5
  ) +
  labs(subtitle = 'Other') +
  scale_fill_manual(values = therapy_colors)

# p_other | p_tch

## Clonal competition
D = M2 %>%
    group_by(MRN) %>%
    filter(any(tch) & any(!tch)) %>%
    ungroup() %>%
    group_by(MRN, tch) %>%
    summarise(
        max_log_alpha = log(max(growth_rate)),
        therapy_binary = unique(therapy_binary),
        Gene = ifelse(any(tch), Gene[1], 'other')
    ) %>%
    mutate(therapy_binary = ifelse(therapy_binary == 1, 'treated', 'untreated')) %>%
    mutate(therapy_binary = factor(therapy_binary, c('treated', 'untreated'))) %>%
    mutate(tch = ifelse(tch, 'DDR', 'Other')) %>%
    mutate(tch = factor(tch, c('DDR', 'Other'))) %>%
    ungroup()

p_tr = D %>%
    filter(therapy_binary == 'treated') %>%
    dcast(MRN ~ tch, value.var = 'max_log_alpha') %>%
    {t.test(.[['DDR']], .[['Other']], paired = TRUE, alternatie = 'two.sided')$p.value}

p_untr = D %>%
    filter(therapy_binary == 'untreated') %>%
    dcast(MRN ~ tch, value.var = 'max_log_alpha') %>%
    {t.test(.[['DDR']], .[['Other']], paired = TRUE, alternatie = 'two.sided')$p.value}

D = D %>%
    arrange(therapy_binary) %>%
    mutate(
        therapy_binary = ifelse(
            therapy_binary == 'treated',
            paste0(therapy_binary, ' (n = ', sum(D$therapy_binary == 'treated')/2, ')', '\n', 'p = ', format(sign
if(p_tr, 2), scientific = T)),
            paste0(therapy_binary, ' (n = ', sum(D$therapy_binary == 'untreated')/2, ')', '\n', 'p = ', format(si
gnif(p_untr, 2), scientific = T))
            )
    ) %>%
    mutate(therapy_binary = factor(therapy_binary, unique(therapy_binary)))
```

```
p_comp = ggplot(
        D,
        aes(x = tch, y = max_log_alpha, group = MRN, fill = therapy_binary, color = therapy_binary)
    ) +
    geom_line(alpha = 0.7) +
    geom_point(
        color = 'black', pch = 21, fill = 'grey'
    ) +
    panel_theme +
    xlab('') +
    ylab('log growth rate') +
    scale_fill_manual(values = therapy_colors) +
    scale_color_manual(values = therapy_colors) +
    facet_wrap(~therapy_binary, ncol = 2)

## Single genes
dta_genes = c('DNMT3A', 'TET2', 'ASXL1')
ddr_genes = c('PPM1D', 'TP53', 'CHEK2')
gene_list = c(ddr_genes, dta_genes)

D = M2 %>%
    filter(Gene %in% gene_list) %>%
    mutate(
        therapy_binary = recode(therapy_binary, `1` = 'treated', `0` = 'untreated')
    ) %>%
    mutate(therapy_binary = factor(therapy_binary, c('treated', 'untreated'))) %>%
    mutate(Gene = factor(Gene, gene_list)) %>%
    mutate(gene_cat = ifelse(Gene %in% dta_genes, 'DTA', 'DDR'))

tests = D %>%
    filter(Gene != 'CHEK2') %>%
    group_by(Gene) %>%
    summarise(
        p.val = t.test(
            log(growth_rate[therapy_binary == 'treated']),
            log(growth_rate[therapy_binary == 'untreated'])
        )$p.value
    ) %>%
    rowwise() %>%
    mutate(q.val = p.adjust(p.val, method = 'fdr', n = nrow(.)))

D = D %>% left_join(tests, by = 'Gene') %>%
    arrange(Gene) %>%
    mutate(gene_label = paste0(Gene, '\nq = ', signif(q.val,2))) %>%
    mutate(gene_label = factor(gene_label, unique(gene_label)))

p_genes = ggplot(
      D,
      aes(x = therapy_binary, y = log(growth_rate), fill = therapy_binary)
  ) +
  geom_boxplot(outlier.alpha = 0, color = 'black') +
  facet_wrap(~gene_label, nrow = 1) +
  geom_jitter(
      pch = 21,
      size = 1,
      alpha = 0.8,
      width = 0.1,
      color = 'black',
      fill = 'grey'
  ) +
  ylab('log growth rate') +
  xlab('') +
  panel_theme +
  scale_fill_manual(values = therapy_colors) +
  theme(
    legend.title = element_blank(),
    # axis.text.x = element_text(angle = 15, size = font_size/1.2, hjust = 0.5),
    axis.text.x = element_blank(),
    legend.position = 'right'
  )

## Dosage

# p values
get_p = function(M, indi) {
  M %>%
  filter(get(indi) >= 1) %>%
    filter(!is.na(age_1)) %>%
    geepack::geeglm(
        data = .,
```

```
        formula = as.formula(paste0('log(growth_rate) ~ ', indi, ' + age_1 + Gender + smoke_bin')),
        id = MRN,
        corstr = "exchangeable"
    ) %>%
    summary %>% .$coefficients %>%
    as.data.frame %>%
    tibble::rownames_to_column('factor') %>%
    filter(factor == indi) %>%
    pull('Pr(>|W|)') %>%
    signif(2) %>%
    format(scientific = T)
}

pval_tch_xrt = M2 %>% filter(tch) %>% get_p('eqd_3_t1')
pval_other_xrt = M2 %>% filter(!tch) %>% get_p('eqd_3_t1')

pval_tch_cyto = M2 %>% filter(tch) %>% get_p('pct_cytotoxic_therapy1')
pval_other_cyto = M2 %>% filter(!tch) %>% get_p( 'pct_cytotoxic_therapy1')

pct_continuous <- function(M, title = '') {

    Mr = M %>% group_by(pct, treatment_type) %>%
        summarise(
            mean_log_alpha = mean(log(growth_rate)),
            lower = quantile(log(growth_rate), 0.25),
            upper = quantile(log(growth_rate), 0.75)
        )

    p = ggplot(
            M,
            aes(x = pct, y = log(growth_rate))
        ) +
        geom_ribbon(
            data = Mr,
            inherit.aes = F,
            aes(x = pct,
                ymin = lower,
                ymax = upper,
                group = ''
            ),
            fill = pal_nejm()(8)[6],
            alpha = 0.6
        ) +
        geom_line(
            data = Mr,
            inherit.aes = F,
            aes(x = pct,
                y = mean_log_alpha,
                group = ''
            ),
            color = pal_nejm()(8)[6]
        ) +
        facet_wrap(~treatment_type, nrow = 1) +
        labs(subtitle = title) +
        panel_theme +
        theme(axis.title.x.top = element_text(size = 4)
        ) +
        xlab('dosage tertile') +
        ylab('log growth rate') +
        geom_jitter(
            pch = 21,
            width = 0.1,
            size = 1,
            alpha = 1,
            color = 'black',
            fill = 'grey'
        ) +
        scale_fill_nejm()

    return(p)
}

D = M2 %>%
    melt(
        measure.vars = c('eqd_3_t1', 'pct_cytotoxic_therapy1'),
        variable.name = 'treatment_type',
        value.name = 'pct'
    ) %>%
    filter(!is.na(pct)) %>%
    mutate(pct = factor(as.integer(pct))) %>%
```

```
    mutate(treatment_type = c('eqd_3_t1' = 'XRT', 'pct_cytotoxic_therapy1' = 'Cytotoxic')[treatment_type])

p_dose_ddr = pct_continuous(
    D %>% filter(tch) %>%
      mutate(treatment_type = paste0(treatment_type, '\n(p = ', c('Cytotoxic' = pval_tch_cyto, 'XRT' = pval_tch_x
rt)[treatment_type], ')')),
    'DDR'
) + ylim(-0.006, 0.006)

p_dose_other = pct_continuous(
    D %>% filter(!tch) %>%
      mutate(treatment_type = paste0(treatment_type, '\n(p = ', c('Cytotoxic' = pval_other_cyto, 'XRT' = pval_oth
er_xrt)[treatment_type], ')')),
    'Other'
) + ylab('') + ylim(-0.006, 0.006)

dat = M2
direction_colors = c('firebrick3', 'grey45', 'lightskyblue') %>%
    setNames(c('UP', 'STABLE', 'DOWN')) %>%
    scale_color_manual(name = "status", values = .)
tx_categories = c("Untreated", "Targeted or\nImmune Therapy", "Cytotoxic Therapy", "XRT")
dat$tx_category = ifelse(dat$ind_anychemo1==0 & dat$XRT1==0, tx_categories[1],
                 ifelse((dat$ind_targeted_therapy1==1 | dat$ind_immune_therapy1==1) &
                        dat$ind_cytotoxic_therapy1==0 & dat$XRT1==0,
                        tx_categories[2],
                 ifelse(dat$ind_cytotoxic_therapy1==1, tx_categories[3],
                 ifelse(dat$XRT1 == 1 , tx_categories[4],
                 "OTHER"))))
dat <- dat %>% mutate(changeT2T1 = VAF_2-VAF_1)
dat <- dat %>% mutate(delta_per_year = changeT2T1 / (delta_time/365))
dat_delta_avg <- dat %>% filter(tx_category != "OTHER") %>% group_by(tx_category) %>%
summarize(mean_delta_per_year = mean(delta_per_year),
          mean_delta_percent = round(100 * mean_delta_per_year, 2),
          delta_time = 365)

p_spider = dat %>%
    filter(tx_category != "OTHER") %>%
    mutate(tx_category = factor(tx_category, tx_categories)) %>%
    ggplot(
        aes(x = delta_time, y = changeT2T1, group = paste0(MRN, variant_id), color = direction)
    ) +
    geom_segment(
        data = . %>% filter(direction == 'STABLE'),
        aes(x = delta_time, y = changeT2T1, xend = 0, yend = 0, color = direction),
        size = 0.2,
        alpha = 0.5
    ) +
    geom_point(
        data = . %>% filter(direction == 'STABLE'),
        size = 0.2,
        alpha = 0.8
    ) +
    geom_segment(
        data = . %>% filter(direction != 'STABLE'),
        aes(x = delta_time, y = changeT2T1, xend = 0, yend = 0, color = direction),
        size = 0.2,
        alpha = 0.5
    ) +
    geom_point(
        data = . %>% filter(direction != 'STABLE'),
        size = 0.2,
        alpha = 0.8
    ) +
    theme_classic() +
    labs(
        x = "Days between Blood Draw",
        y = "Change in VAF"
    ) +
    theme(
        axis.text.x = element_text(angle = 45, hjust = 1),
        legend.position="top"
    ) +
  facet_wrap(~tx_category, ncol = 4, nrow = 1) +
  panel_theme +
  theme(
    legend.title = element_blank(),
    legend.position = 'right'
  ) +
  direction_colors
```

```
panel = (p_spider + labs(title = 'A')) /
  ((p_tch + labs(title = 'B'))| p_other) /
  (p_genes + labs(title = 'C')) /
  ((p_dose_ddr + labs(title = 'D'))| p_dose_other) /
  (p_comp + labs(title = 'E'))

do_plot(panel, "figure3.png", 6.5, 11, save_pdf = T)
```

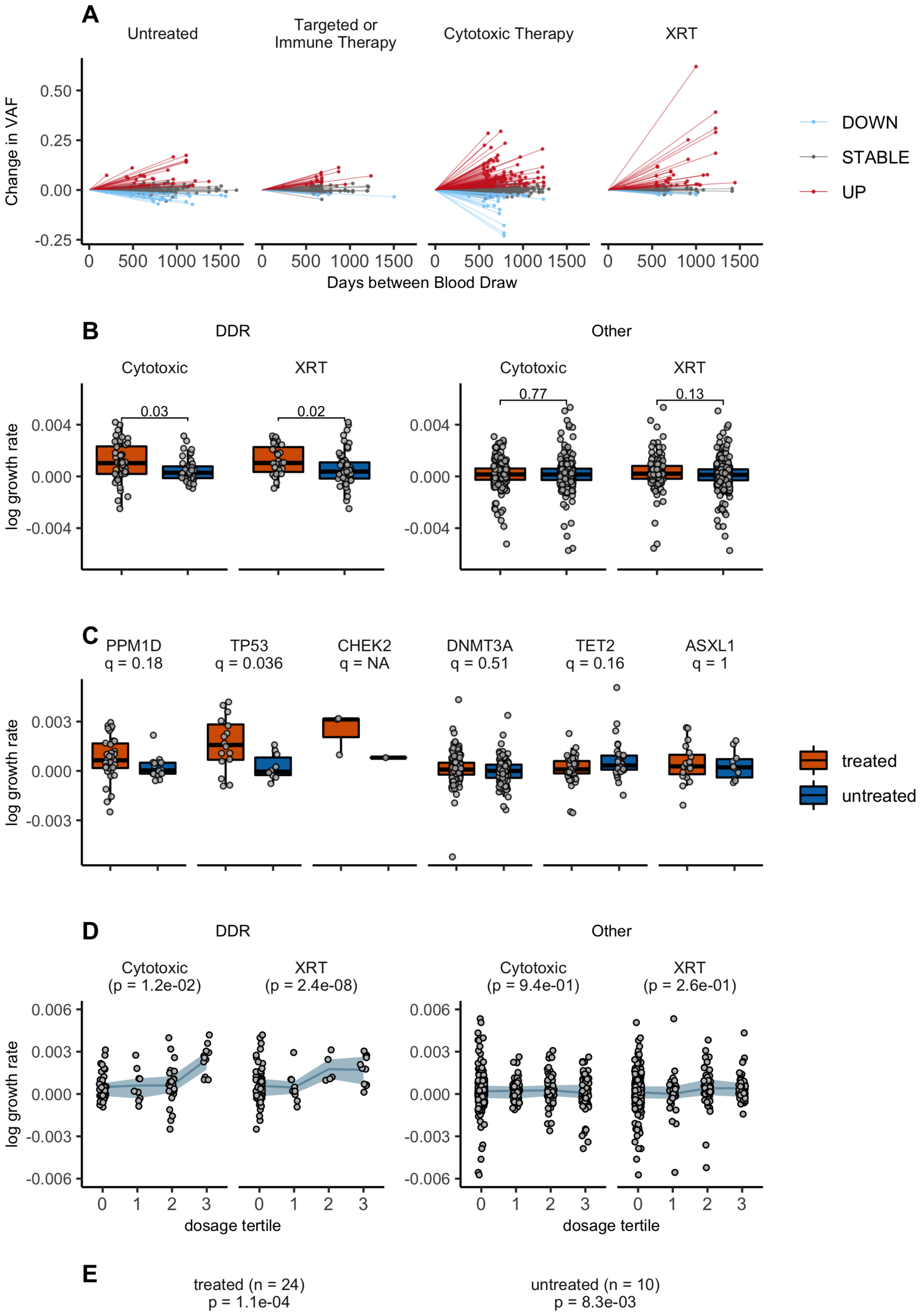

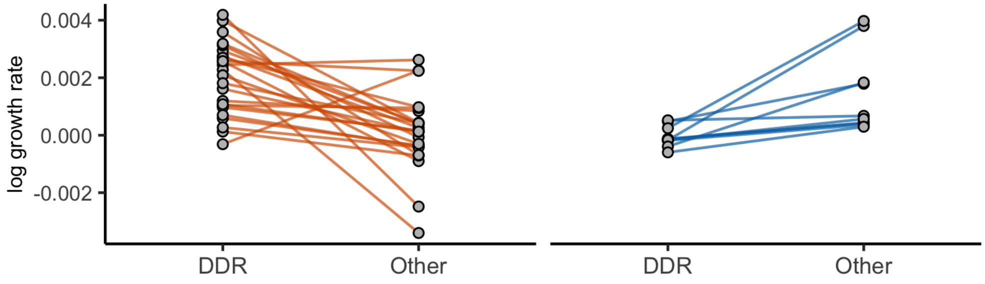


##Figure 4 - CH to tMN

```
#CH overall
M_tmn_wide_st <- M_tmn_wide_st %>% mutate(center =fct_relevel(center, "MSK","MDA","MOF","HCC"))

cox <- coxph(Surv(timelastfu, post_tmn) ~ ch_my_pd + age_d + Gender + strata(center), data= M_tmn_wide_st)
cox
```

```
## Call:
## coxph(formula = Surv(timelastfu, post_tmn) ~ ch_my_pd + age_d +
##     Gender + strata(center), data = M_tmn_wide_st)
##
##              coef exp(coef) se(coef)      z        p
## ch_my_pd  1.76787   5.85837  0.25366  6.969 3.18e-12
## age_d    -0.13097   0.87724  0.09855 -1.329   0.1839
## GenderM   0.54751   1.72894  0.25129  2.179   0.0293
##
## Likelihood ratio test=50.72  on 3 df, p=5.604e-11
## n= 9548, number of events= 75
```

```
cox <- coxph(Surv(timelastfu, post_tmn) ~ ch_my_pd*center + age_d + Gender + strata(center), data= M_tmn_wide_st)
cox
```

```
## Call:
## coxph(formula = Surv(timelastfu, post_tmn) ~ ch_my_pd * center +
##     age_d + Gender + strata(center), data = M_tmn_wide_st)
##
##                        coef exp(coef) se(coef)      z        p
## ch_my_pd             1.9026    6.7035   0.3910  4.866 1.14e-06
## centerMDA                NA        NA   0.0000     NA       NA
## centerMOF                NA        NA   0.0000     NA       NA
## centerHCC                NA        NA   0.0000     NA       NA
## age_d               -0.1297    0.8783   0.1031 -1.259   0.2081
## GenderM              0.5141    1.6722   0.2532  2.031   0.0423
## ch_my_pd:centerMDA  0.6367    1.8902   0.7253  0.878   0.3801
## ch_my_pd:centerMOF -1.1083    0.3301   0.6773 -1.636   0.1018
## ch_my_pd:centerHCC -0.1756    0.8389   0.6186 -0.284   0.7765
##
## Likelihood ratio test=55.51  on 6 df, p=3.654e-10
## n= 9548, number of events= 75
```

```
#Mutation number and max VAF
mut_var <- c("n_mut_r","VAF_nonsilent_r")

cox_data <- list()
cox_mut_var <- list()
  for (var in mut_var) {

    cox <- coxph(Surv(timelastfu, post_tmn) ~ as.factor(get(var))+ age + Gender + strata(center), data= M_tmn_wid
e_st)
    cox_data <- cox %>% get_model_data()
    cox_data <- cox_data %>% cbind(mut_var = var)
    cox_mut_var <- rbind(cox_mut_var, cox_data)
                          }
cox_mut_var_stonly<- cox_mut_var %>% filter(term!="age" & term!="GenderM")

#Gene Effects
top_genes_cases <- M_tmn_long %>%
filter(VAF_1 >= 0.02 & center!="PMC" & center!="WSU" & post_tmn==1) %>%
group_by(Gene) %>%
  summarise(count=n_distinct(center_id)) %>%
  arrange(-count)  %>%
  filter(count>=2) %>%
  pull(Gene) %>%
  as.character()

top_genes_controls <- M_tmn_long %>%
filter(VAF_1 >= 0.02 & center!="PMC" & center!="WSU") %>%
group_by(Gene) %>%
  summarise(count=n_distinct(center_id)) %>%
  arrange(-count)  %>%
  filter(count>=20) %>%
  pull(Gene) %>%
  as.character()

top_genes <- top_genes_cases %>% intersect(top_genes_controls)

cox_gene_var <- list()
  for (gene in top_genes) {
    cox <- coxph(Surv(timelastfu, post_tmn) ~ get(gene) + age + Gender + strata(center), data= M_tmn_wide_st, na.
action=na.omit)
    cox_data <- cox %>% get_model_data()
    cox_data <- cox_data %>% cbind(Gene = gene)
    cox_gene_var <- rbind(cox_gene_var, cox_data)
  }

#CBC parameters
cbc_var <- c("hgb_c","mcv_c","rdw_c","wbc_c","hgb_c","anc_c","amc_c","plt_c")

cox_cbc_var <- list()
  for (var in cbc_var) {
    cox <- coxph(Surv(timelastfu, post_tmn) ~ get(var)+ age + Gender + strata(center), data= M_tmn_wide_st , na.a
ction=na.omit)
    cox_data <- cox %>% get_model_data()
    cox_data <- cox_data %>% cbind(CBC_var = var)
    cox_cbc_var <- rbind(cox_cbc_var, cox_data)
                          }

cbc = cox_cbc_var %>%
    filter(CBC_var %in% cbc_var) %>%
    filter (term=="get(var)") %>%
    mutate(CBC = case_when(
            CBC_var == 'wbc_c' ~ "White Blood Cell Count",
            CBC_var == 'anc_c' ~ "Neutrophil Count",
            CBC_var == 'amc_c' ~ "Monocyte Count",
            CBC_var == 'hgb_c' ~ "Hemoglobin",
            CBC_var == 'mcv_c' ~ "Mean Corpuscular Volume",
            CBC_var == 'rdw_c' ~ "Red Cell Distribution Width",
            CBC_var == 'plt_c' ~ "Platelets"
    ))

#One forest plot for all
cox_gene_var_v2 <- cox_gene_var %>%
    filter(term=="get(gene)") %>%
    mutate(group="Gene") %>%
    mutate(mut_var= factor(Gene, levels=unique(Gene[order(estimate)])))

cbc_v2 <- cbc %>% mutate(mut_var=CBC) %>% mutate(group="Blood Count Index")

cox_gene_var_v2 <- select(cox_gene_var_v2,-c(Gene))
```

```
cbc_v2 <- select(cbc_v2,-c(CBC_var,CBC)) %>% mutate(mut_var= factor(mut_var, levels=unique(mut_var[order(estimat
e)])))

cox_mut_var_stonly_v2 <- cox_mut_var_stonly %>%
            mutate(new_var=case_when(
            term=="as.factor(get(var))1" & mut_var=="n_mut_r" ~ "1",
            term=="as.factor(get(var))2" & mut_var=="n_mut_r" ~ "2",
            term=="as.factor(get(var))3" & mut_var=="n_mut_r" ~ "3 or more"
            ))

cox_mut_var_stonly_v2 <- cox_mut_var_stonly_v2 %>%
            mutate(new_var=case_when(
            term=="as.factor(get(var))1" & mut_var=="VAF_nonsilent_r" ~ "2-5%",
            term=="as.factor(get(var))2" & mut_var=="VAF_nonsilent_r" ~ "5-10%",
            term=="as.factor(get(var))3" & mut_var=="VAF_nonsilent_r" ~ "10-20%",
            term=="as.factor(get(var))4" & mut_var=="VAF_nonsilent_r" ~ "20% or more",
            TRUE ~ new_var
            ))

cox_mut_var_stonly_v2 <- cox_mut_var_stonly_v2 %>%
            mutate(group=case_when(
            mut_var=="n_mut_r" ~ "Mutation Number",
            mut_var=="VAF_nonsilent_r" ~ "Maximum VAF"))

cox_mut_var_stonly_v2 <- cox_mut_var_stonly_v2 %>% select(-c("mut_var"))
cox_mut_var_stonly_v2 <- cox_mut_var_stonly_v2 %>% rename(mut_var=new_var)

combo <- rbind(cox_gene_var_v2, cbc_v2, cox_mut_var_stonly_v2)

uni_plot = plot_forest(
      combo,
      x = "mut_var",
      label = 'p.stars',
      eb_w = 0,
      eb_s = 0.3,
      ps = 1.5,
      or_s = 2,
      nudge = -0.3) +
 #     col = 'group'
  facet_grid(group ~ ., scale = 'free_y', space = "free_y") +
   scale_x_discrete(
       breaks = combo$mut_var,
       labels = combo$mut_var,
       expand = c(0.2,0)
   ) +
  xlab('') + ylab('OR') +
  scale_color_nejm() +
  panel_theme

do_plot(uni_plot, "figure4a.png", w = 3, h = 4)
```

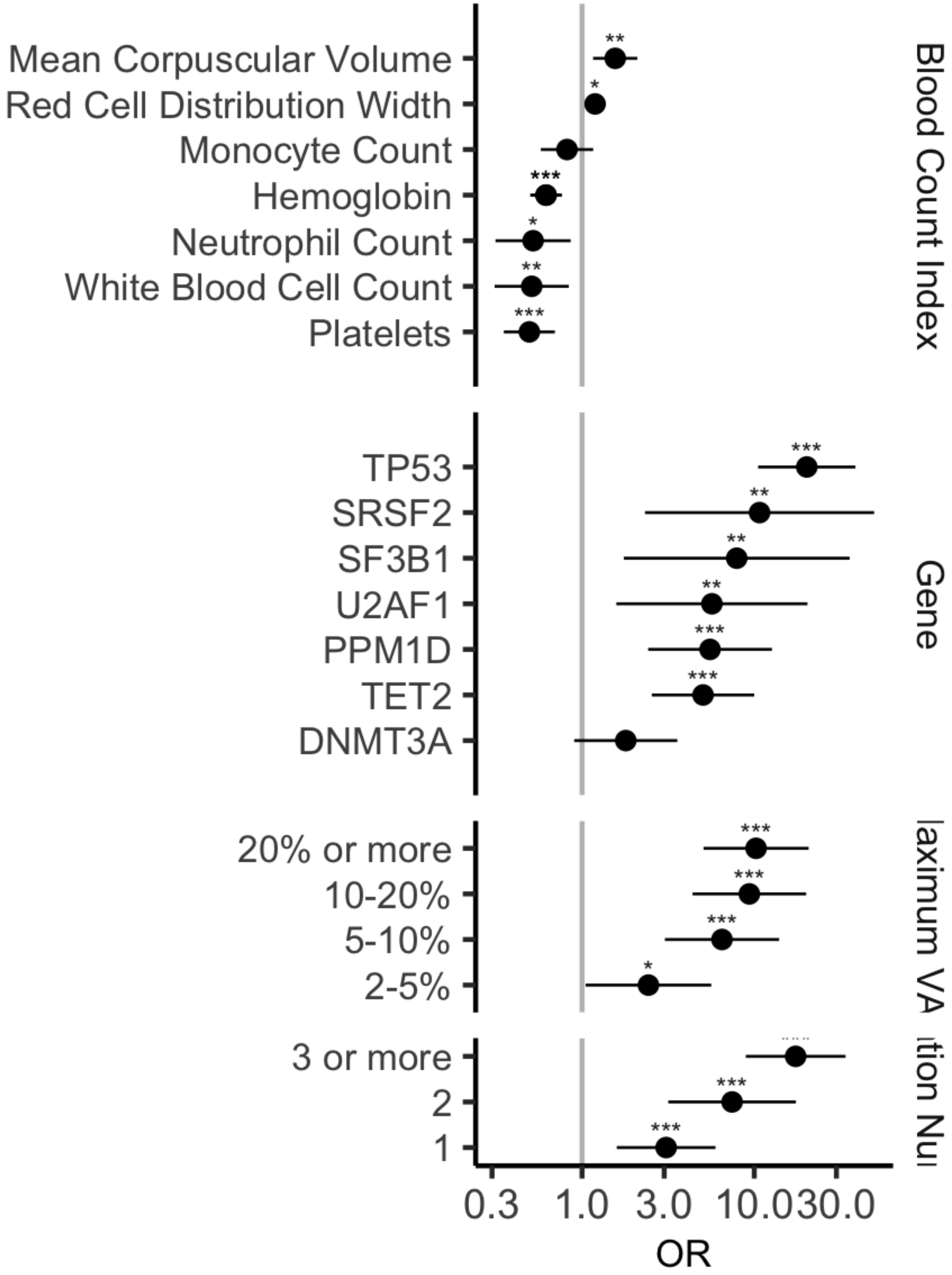


```
#Dont include PPM1D and SRSF2 b/c missing in some
gene_vars = c("TET2","SF3B1","TP53","U2AF1")
mut_vars = c("VAF_nonsilent_r","n_mut_r")
cbc_vars = c("hgb_c","rdw_c", "mcv_c", "wbc_c","anc_c", "plt_c")
demo_vars = c('age', 'Gender','strata(center)')

#include mean imputation for blood counts when missing
D = M_tmn_wide_st %>%
    filter(!is.na(age)) %>%
    filter(center != 'PMC') %>%
    mutate_at(vars(cbc_vars),.funs = funs(ifelse(is.na(.), mean(., na.rm=TRUE), .)))
```

```
## Warning: funs() is soft deprecated as of dplyr 0.8.0
## Please use a list of either functions or lambdas:
##
##   # Simple named list:
##   list(mean = mean, median = median)
##
##   # Auto named with `tibble::lst()`:
##   tibble::lst(mean, median)
##
##   # Using lambdas
##   list(~ mean(., trim = .2), ~ median(., na.rm = TRUE))
## This warning is displayed once per session.
```

```
form = paste0(
    'Surv(timelastfu, post_tmn) ~ ',
    paste(
        c(
            gene_vars,
            mut_vars,
            cbc_vars,
            demo_vars
        ),
        collapse = ' + '
    )
)

model = coxph(formula = as.formula(form), data = D, na.action=na.omit) %>% summary

betas = model %>%
    .$coefficients %>% as.data.frame %>%
    tibble::rownames_to_column('term') %>%
    filter(!(term %in% c(
        'pre_any_therapy',
        'GenderM')
        )
    ) %>% select(term, coef)

# add beta of therapy from external source
betas = betas %>% rbind(data.frame(term = 'chemo', coef = log(6.8)))

# prepare covariate matrix
Z = left_join(
      #add average chemo
      M_tmn_wide_st %>% mutate(
          age_bin = case_when(
              age < 20 ~ '<20',
              age >= 20 & age < 25 ~ '20-24',
              age >= 25 & age < 30 ~ '25-29',
              age >= 30 & age < 35 ~ '30-34',
              age >= 35 & age < 40 ~ '35-39',
              age >= 40 & age < 45 ~ '40-44',
              age >= 45 & age < 50 ~ '45-49',
              age >= 50 & age < 55 ~ '50-54',
              age >= 55 & age < 60 ~ '55-59',
              age >= 60 & age < 65 ~ '60-64',
              age >= 65 & age < 70 ~ '65-69',
              age >= 70 & age < 75 ~ '70-74',
              age >= 75 & age < 80 ~ '75-79',
              age >= 80 & age < 85 ~ '80-84',
              age >= 85 ~ '85+'
              )
      ),
      T_seer %>%
          rename(age_bin = age) %>%
          mutate(age_bin = str_remove(age_bin, ' years')),
      by = 'age_bin'
  ) %>%
  filter(age > 50 & age < 75) %>%
  filter(pre_any_therapy == 0) %>%
  filter(center == 'MSK') %>%
  select(betas$term, post_tmn) %>%
  filter(complete.cases(.))

y = Z %>% pull(post_tmn)
Z = Z %>% select(-post_tmn)

# info for the covariates
Z_info = lapply(betas$term, function(x){list(name = x, type = 'continuous')})

## formula
formula = paste0(
    'observed.outcome ~ ',
    paste(
        betas$term,
        collapse = '+',
        sep = '+'
    )
)

# modified chemoxrt column
Z_tr = Z %>% mutate(chemo = 1)
Z_untr = Z %>% mutate(chemo = 0)
```

```
## build model
res_tr = iCARE::computeAbsoluteRisk(
    model.formula = as.formula(formula),
    model.cov.info = Z_info,
    model.ref.dataset = Z,
    model.log.RR = as.matrix(unname(tibble::column_to_rownames(betas, 'term'))),
    model.disease.incidence.rates = tmn_inc,
    model.competing.incidence.rates = mort_inc,
    apply.age.start = 1,
    apply.age.interval.length = 9,
    apply.cov.profile = Z_tr,
    return.refs.risk = TRUE
)
```

```
##    user  system elapsed
##   0.000   0.000   0.001
```

```
res_untr = iCARE::computeAbsoluteRisk(
    model.formula = as.formula(formula),
    model.cov.info = Z_info,
    model.ref.dataset = Z,
    model.log.RR = as.matrix(unname(tibble::column_to_rownames(betas, 'term'))),
    model.disease.incidence.rates = tmn_inc,
    model.competing.incidence.rates = mort_inc,
    apply.age.start = 1,
    apply.age.interval.length = 9,
    apply.cov.profile = Z_untr,
    return.refs.risk = TRUE
)
```

```
##    user  system elapsed
##   0.000   0.000   0.001
```

```
#Figure 4b
D = data.frame(
        risk_ref = as.vector(res_tr$refs.risk),
        risk_tr = as.vector(res_tr$risk),
        risk_untr = as.vector(res_untr$risk),
        Z
    ) %>%
    mutate(
        risk_untr_pct = ntile(risk_untr, 100)
    )

p_dist <- D %>% ggplot(
    aes(x = risk_ref)
) +
theme_classic() +
geom_density(alpha = 0.5,color = "black", fill = "gray") +
xlim(0.0,0.022) +
ylim(0.0, 400) +
theme_classic() +
geom_density(alpha = 0.5) +
xlim(0,0.022) +
scale_fill_nejm() +
ylab('Number of Women') +
xlab("10-year absolute risk of AML/MDS for women with breast cancer in the U.S aged 50-75")

do_plot(p_dist, "figure4b.svg", w = 6, h = 2.5)
```

```
## Warning: package 'gdtools' was built under R version 3.5.2
```

```
## Warning: Removed 25 rows containing non-finite values (stat_density).

## Warning: Removed 25 rows containing non-finite values (stat_density).

## Warning: Removed 25 rows containing non-finite values (stat_density).

## Warning: Removed 25 rows containing non-finite values (stat_density).
```

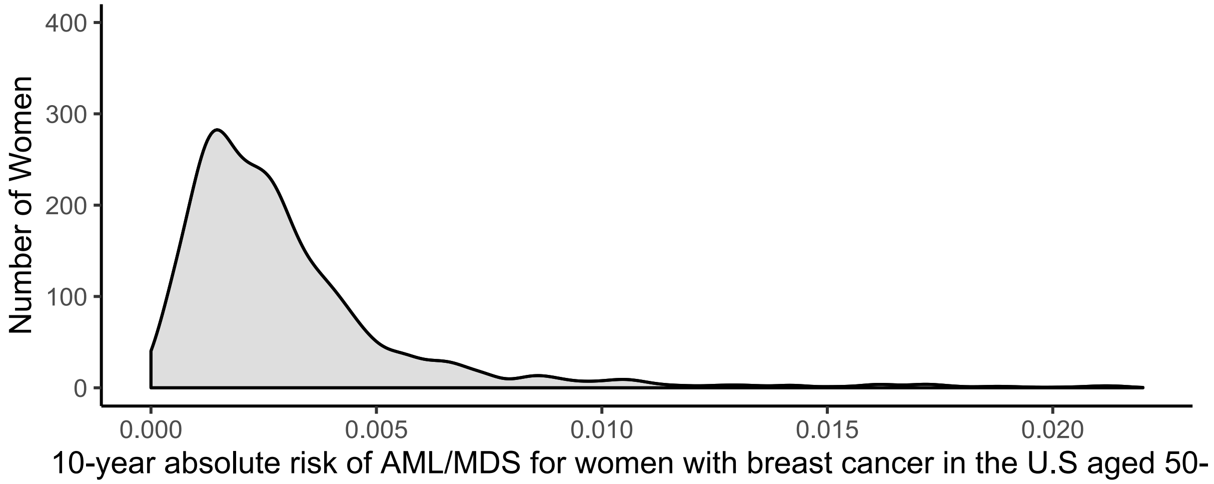


```
#Calculate number of women with absolute risk of more than 1%
morethan1 <- D %>% filter(risk_ref>0.01) %>% count()
total <- D %>% count()
(total-morethan1)/total
```

```
##           n
## 1 0.9551591
```

```
#Figure 4c
D = data.frame(
        risk_ref = as.vector(res_tr$refs.risk),
        risk_tr = as.vector(res_tr$risk),
        risk_untr = as.vector(res_untr$risk)
    ) %>%
    mutate(
        risk_untr_pct = ntile(risk_untr, 100)
    )%>%
    melt(
        measure.vars = c('risk_tr', 'risk_untr', 'risk_ref'),
        value.name = 'risk',
        variable.name = 'treatment'
    )

therapy_colors = c("#D55E00","#0072B2")

p <- ggplot(D %>% filter(treatment %in% c('risk_tr', 'risk_untr')) %>%
        filter(risk_untr_pct >= 97),
    aes(x = as.factor(risk_untr_pct),
        y = risk,
        fill = treatment
    )
) +
theme_bw() +
scale_y_continuous(position = "left", limits = c(0,0.2)) +
geom_boxplot(
    position = 'dodge',
    outlier.alpha = 0,
    alpha = 0.5,
    size = 0.4
) +
scale_fill_nejm() +
theme(
    panel.grid.major.x = element_blank(),
    panel.grid.minor.y = element_blank(),
    panel.grid.major.y = element_line(linetype = 'dashed'),
    legend.position = 'right'
) +
xlab('Absolute Risk Percentile') +
ylab('Absolute 10-year AML/MDS Risk')

p_zoom <- p+labs(fill = "Cytotoxic Therapy") + theme(legend.position = c(0.2, 0.7)) + scale_fill_manual(values =
 therapy_colors,labels = c("Yes", "No")) + scale_color_manual(values = therapy_colors)

do_plot(p_zoom, "figure4c_box.svg", w = 6, h = 3)
```

```
## Warning: Removed 5 rows containing non-finite values (stat_boxplot).
```

```
## Warning: Removed 5 rows containing non-finite values (stat_boxplot).
```

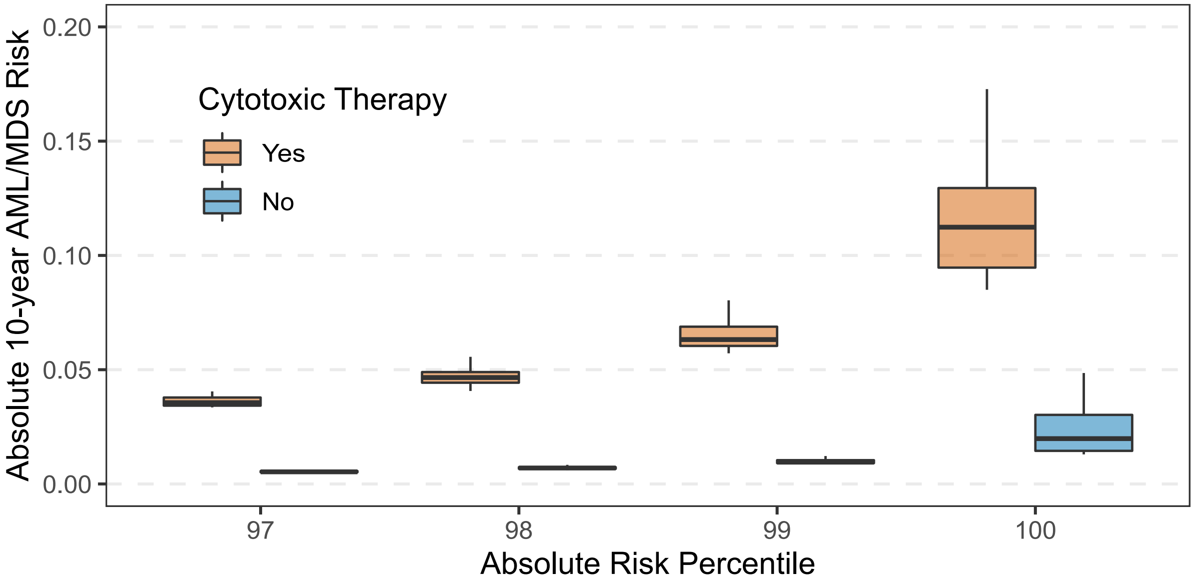


### Extended and Supplemental Figures and Tables

#eTable1 - baseline demographics

```
tally_variable = function(D, baseline_cols, variable) {

    if (variable == 'Total') {

        res = cbind(
            D %>% dcast(. ~ CH_all, fun.aggregate = length, value.var = 'MRN') %>% select(-.),
            D %>% dcast(. ~ therapy_binary, fun.aggregate = length, value.var = 'MRN') %>% select(-.)
        ) %>%
        mutate(Total = rowSums(.[1:2])) %>%
        mutate(variable = 'Total', category = 'Total') %>%
        select(baseline_cols) %>%
        mutate(ref = "")

    } else {

        ref = levels(D[[variable]])[1]

        if (is.null(ref)) {
            ref = ""
        }

        res = cbind(
            D %>% dcast(paste0(variable, ' ~ CH_all'), fun.aggregate = length, value.var = 'MRN') %>%
                mutate(variable = variable) %>% rename(category = !!variable),
            D %>% dcast(paste0(variable, ' ~ therapy_binary'), fun.aggregate = length, value.var = 'MRN'),
            D %>% count(get(variable)) %>% select(n) %>% rename(Total = n)
        ) %>%
        select(baseline_cols) %>%
        arrange(category == 'Missing') %>% # sorting
        mutate(ref = ref)
    }

    res = res %>%
        mutate(`CH-` = paste0(`CH-`, ' (', signif(`CH-`* 100/Total, 2), '%)')) %>%
        mutate(`CH+` = paste0(`CH+`, ' (', signif(`CH+`* 100/Total, 2), '%)'))
    return(res)
}

D = M_wide_all %>%
    mutate(CH_all = ifelse(CH_all, 'CH+', 'CH-')) %>%
    mutate(age_verbose = case_when(
        is.na(age_cat) ~ 'Missing',
        T ~ paste0((age_cat - 1) * 10, '-', age_cat * 10))
    ) %>%
    mutate(smoke_verbose = case_when(
        smoke == 1 ~ "Current/Former",
        smoke == 0 ~ "Non Smoker",
        T ~ 'Missing')
    ) %>%
    mutate(smoke_verbose = relevel(factor(smoke_verbose), ref = 'Non Smoker')) %>%
    mutate(
        therapy_binary = case_when(
          (txMSK | thyroid == 1) & (ind_anychemo == 1 | XRT == 1) ~ "Treated",
          (txMSK | thyroid == 1) ~ "Untreated",
          T ~ "Unknown"
        )
    ) %>%
    mutate(therapy_binary = ifelse(is.na(therapy_binary), 'Unknown', therapy_binary)) %>%
    mutate(therapy_binary = factor(therapy_binary, c("Treated", "Untreated", "Unknown"))) %>%
    mutate(Gender = relevel(factor(Gender), ref = 'Male')) %>%
    mutate(generaltumortype = ifelse(
        generaltumortype == '' | is.na(generaltumortype), 'Missing', generaltumortype)
    ) %>%
    mutate(generaltumortype = format_variable(generaltumortype))

baseline_cols = c('variable', 'category', 'CH-', 'CH+', 'Total')

rbind(
    D %>% tally_variable(baseline_cols = baseline_cols, variable = 'Total'),
    D %>% tally_variable(baseline_cols = baseline_cols, variable = 'smoke_verbose'),
    D %>% tally_variable(baseline_cols = baseline_cols, variable = 'Gender'),
    D %>% tally_variable(baseline_cols = baseline_cols, variable = 'age_verbose'),
    D %>% tally_variable(baseline_cols = baseline_cols, variable = 'race'),
    D %>% tally_variable(baseline_cols = baseline_cols, variable = 'therapy_binary')
) %>%
mutate(variable = ifelse(variable == 'generaltumortype', 'Primary Tumor Subtype', variable)) %>%
mutate(variable = format_variable(variable)) %>%
select(-ref) %>%
kable(format = "latex", booktabs = T) %>%
```

```
kable_styling(
    latex_options = c("hold_position"),
    full_width = T,
    font_size = 10
) %>%
column_spec(1, width = 10) %>%
column_spec(2:5, width = 70) %>%
collapse_rows(columns = 1:2, row_group_label_position = 'stack') %>%
display_kable('table1.1.pdf')
```

| | category | CH- | CH+ | Total |
|---|---|---|---|---|
| **Total** | | | | |
| | Total | 17060 (70%) | 7379 (30%) | 24439 |
| **Smoke** | | | | |
| | Non Smoker | 9048 (74%) | 3147 (26%) | 12195 |
| | Current/Former | 7313 (65%) | 3997 (35%) | 11310 |
| | Missing | 699 (75%) | 235 (25%) | 934 |
| **Gender** | | | | |
| | Male | 7782 (70%) | 3405 (30%) | 11187 |
| | Female | 9278 (70%) | 3974 (30%) | 13252 |
| **Age** | | | | |
| | 0-10 | 324 (96%) | 13 (3.9%) | 337 |
| | 10-20 | 288 (95%) | 14 (4.6%) | 302 |
| | 20-30 | 676 (95%) | 38 (5.3%) | 714 |
| | 30-40 | 1408 (92%) | 122 (8%) | 1530 |
| | 40-50 | 2768 (87%) | 424 (13%) | 3192 |
| | 50-60 | 4519 (77%) | 1316 (23%) | 5835 |
| | 60-70 | 4535 (63%) | 2621 (37%) | 7156 |
| | 70-80 | 2158 (50%) | 2162 (50%) | 4320 |
| | 80-90 | 384 (36%) | 669 (64%) | 1053 |
| **Race** | | | | |
| | White | 12732 (68%) | 5948 (32%) | 18680 |
| | Asian | 1278 (78%) | 358 (22%) | 1636 |
| | Black | 1089 (72%) | 417 (28%) | 1506 |
| | Other | 1182 (77%) | 360 (23%) | 1542 |
| | Missing | 779 (72%) | 296 (28%) | 1075 |
| **Therapy** | | | | |
| | Treated | 4384 (70%) | 1856 (30%) | 6240 |
| | Untreated | 2877 (73%) | 1090 (27%) | 3967 |
| | Unknown | 9799 (69%) | 4433 (31%) | 14232 |

```
D %>% tally_variable(baseline_cols = baseline_cols, variable = 'generaltumortype') %>%
mutate(variable = ifelse(variable == 'generaltumortype', 'Primary Tumor Subtype', variable)) %>%
mutate(variable = format_variable(variable)) %>%
select(-ref) %>%
kable(format = "latex", booktabs = T) %>%
kable_styling(
    latex_options = c("hold_position"),
    full_width = T,
    font_size = 10
) %>%
column_spec(1, width = 10) %>%
column_spec(2, width = 200) %>%
column_spec(3:7, width = 50) %>%
collapse_rows(columns = 1:2, row_group_label_position = 'stack') %>%
display_kable('table1.2.pdf')
```

| category | CH- | CH+ | Total |
|---|---|---|---|
| **Primary Tumor Subtype** | | | |
| Adrenocortical Carcinoma | 43 (74%) | 15 (26%) | 58 |
| Ampullary Carcinoma | 48 (75%) | 16 (25%) | 64 |
| Anal Cancer | 38 (67%) | 19 (33%) | 57 |
| Appendiceal Cancer | 128 (79%) | 35 (21%) | 163 |
| Biliary Cancer | 354 (69%) | 160 (31%) | 514 |
| Bladder Cancer | 447 (62%) | 274 (38%) | 721 |
| Breast Carcinoma | 2617 (74%) | 936 (26%) | 3553 |
| Breast Sarcoma | 30 (68%) | 14 (32%) | 44 |
| Cancer of Unknown Primary | 492 (67%) | 242 (33%) | 734 |
| Cervical Cancer | 91 (77%) | 27 (23%) | 118 |
| CHondroblastoma | 1 (100%) | 0 (0%) | 1 |
| CHondrosarcoma | 43 (78%) | 12 (22%) | 55 |
| CHordoma | 28 (76%) | 9 (24%) | 37 |
| CHoroid Plexus Tumor | 3 (100%) | 0 (0%) | 3 |
| Colorectal Cancer | 1628 (75%) | 539 (25%) | 2167 |
| Embryonal Tumor | 153 (89%) | 18 (11%) | 171 |
| Endometrial Cancer | 515 (61%) | 328 (39%) | 843 |
| Ependymomal Tumor | 26 (90%) | 3 (10%) | 29 |
| Esophagogastric Carcinoma | 468 (70%) | 197 (30%) | 665 |
| Ewing Sarcoma | 66 (88%) | 9 (12%) | 75 |
| Gastrointestinal Neuroendocrine Tumor | 73 (68%) | 34 (32%) | 107 |
| Gastrointestinal Stromal Tumor | 201 (70%) | 85 (30%) | 286 |
| Germ Cell Tumor | 354 (91%) | 36 (9.2%) | 390 |
| Gestational Trophoblastic Disease | 10 (77%) | 3 (23%) | 13 |
| Glioma | 838 (76%) | 266 (24%) | 1104 |
| Head and Neck Carcinoma | 253 (69%) | 114 (31%) | 367 |
| Hepatocellular Carcinoma | 137 (70%) | 58 (30%) | 195 |
| Melanoma | 622 (69%) | 281 (31%) | 903 |
| Meningothelial Tumor | 54 (79%) | 14 (21%) | 68 |
| Mesothelioma | 148 (65%) | 78 (35%) | 226 |

| | | | |
|---|---|---|---|
| Miscellaneous Brain Tumor | 26 (87%) | 4 (13%) | 30 |
| Miscellaneous Neuroepithelial Tumor | 11 (65%) | 6 (35%) | 17 |
| Nerve Sheath Tumor | 43 (88%) | 6 (12%) | 49 |
| Non-Small Cell Lung Cancer | 2252 (62%) | 1359 (38%) | 3611 |
| Osteosarcoma | 98 (90%) | 11 (10%) | 109 |
| Ovarian Cancer | 412 (61%) | 259 (39%) | 671 |
| Pancreatic Cancer | 972 (68%) | 461 (32%) | 1433 |
| Penile Cancer | 7 (78%) | 2 (22%) | 9 |
| PheoCHromocytoma | 6 (86%) | 1 (14%) | 7 |
| Pineal Tumor | 1 (25%) | 3 (75%) | 4 |
| Prostate Cancer | 978 (65%) | 537 (35%) | 1515 |
| Renal Cell Carcinoma | 445 (77%) | 130 (23%) | 575 |
| Renal Non-Clear Cell Carcinoma | 2 (67%) | 1 (33%) | 3 |
| Retinoblastoma | 38 (95%) | 2 (5%) | 40 |
| Salivary Carcinoma | 162 (75%) | 54 (25%) | 216 |
| Sellar Tumor | 53 (88%) | 7 (12%) | 60 |
| Sex Cord Stromal Tumor | 29 (81%) | 7 (19%) | 36 |
| Skin Cancer, Non-Melanoma | 138 (58%) | 101 (42%) | 239 |
| Small Bowel Cancer | 67 (77%) | 20 (23%) | 87 |
| Small Cell Lung Cancer | 128 (60%) | 86 (40%) | 214 |
| Soft Tissue Sarcoma | 724 (77%) | 222 (23%) | 946 |
| Thymic Tumor | 35 (69%) | 16 (31%) | 51 |
| Thyroid Cancer | 270 (61%) | 170 (39%) | 440 |
| Uterine Sarcoma | 126 (73%) | 46 (27%) | 172 |
| Vaginal Cancer | 11 (69%) | 5 (31%) | 16 |
| Wilms Tumor | 23 (96%) | 1 (4.2%) | 24 |
| Missing | 94 (70%) | 40 (30%) | 134 |

#sFigure 2 mutational landscape

```
M_long = M_long %>%
    mutate(myeloid_category = case_when(
      CH_my == 1 ~ "Myeloid Gene",
      CH_my == 0 ~ "Non-Myeloid Gene"
    )) %>%
    mutate(pd_category = case_when(
      ch_my_pd == 1 ~ "Myeloid PD",
      ch_my_pd == 0 & ch_pancan_pd == 1 ~ "Non-Myeloid PD",
      TRUE ~ "Non PD"
    )) %>%
    mutate(
        VariantClass2 = case_when(
            VariantClass == 'Frame_Shift_Del' ~ 'Frameshift indel',
            VariantClass == 'Frame_Shift_Ins' ~ 'Frameshift indel',
            VariantClass == 'In_Frame_Del' ~ 'Inframe indel',
            VariantClass == 'In_Frame_Ins' ~ 'Inframe indel',
            VariantClass == 'Splice_Site' ~ 'Splice or other',
            VariantClass == 'Splice_Region' ~ 'Splice or other',
            VariantClass == "5'Flank" ~ 'Splice or other',
            VariantClass == 'Translation_Start_Site' ~ 'Splice or other',
            VariantClass == 'Nonstop_Mutation' ~ 'Splice or other',
            VariantClass == 'Missense_Mutation' ~ 'Missense',
            VariantClass == 'Nonsense_Mutation' ~ 'Nonsense',
            T ~ VariantClass)
    )

font_size = 8

lab_pos = 'out'

lab_color = 'black'

panel_theme = theme(
    legend.direction = 'vertical',
    legend.position="top",
    legend.text = element_text(size=12),
    plot.margin = margin(t = 0, r = 0, b = 0, l = 0),
    axis.text = element_text(size = font_size)
  )

p1 = ggdonutchart(
    data = M_long %>% filter(myeloid_category != "NA") %>% count(myeloid_category),
    x = "n",
    fill = "myeloid_category",
    color = "white",
    lab.pos = lab_pos,
    palette = pal_nejm('default')(8),
    lab.font = c(lab_color, font_size),
    size = 0.5) +
    labs(fill = "Gene Class") +
    labs(title = "B") +
    panel_theme

p2 = ggdonutchart(
    data = M_long %>% count(pd_category),
    x = "n",
    fill = "pd_category",
    color = "white",
    lab.pos = lab_pos,
    palette = pal_nejm('default')(8),
    lab.font = c(lab_color, font_size),
    size = 0.5) +
    labs(fill = "Mutation Driver Status") +
     labs(title = "C") +
  panel_theme

p3 = ggdonutchart(
    data = M_long %>% count(VariantClass2),
    x = "n",
    fill = "VariantClass2",
    color = "white",
    lab.pos = lab_pos,
    palette = pal_nejm('default')(8),
    lab.font = c(lab_color, font_size),
    size = 0.5) +
    labs(fill = "Variant Effect") +
    labs(title = "D") +
    panel_theme
```

```
p4 = ggdonutchart(
    data = M_long %>% count(variant_type),
    x = "n",
    fill = "variant_type",
    color = "white",
    lab.pos = lab_pos,
    palette = pal_nejm('default')(8),
    lab.font = c(lab_color, font_size),
    size = 0.5) +
    labs(fill = "Variant Type") +
    labs(title = "E") +
    panel_theme

p5 = ggdonutchart(
    data = M_long %>% filter(variant_type == 'SNV') %>% count(sub_nuc),
    x = "n",
    fill = "sub_nuc",
    color = "white",
    lab.pos = lab_pos,
    palette = pal_nejm('default')(8),
    lab.font = c(0.1, "plain", lab_color),
    size = 0.5) +
    labs(fill = "Nucleotide Change (SNVs)") +
    labs(title = "F") +
    panel_theme

panel = (p1 | p2 | p3 | p4 | p5)

#Top Genes

gene_list = M_long %>% count(Gene) %>% arrange(-n) %>% .$Gene %>% unique %>% .[1:30]

genes <- ggplot(
    M_long %>% filter(Gene %in% gene_list) %>%
    mutate(Gene = factor(Gene, gene_list)) %>%
    count(Gene, VariantClass2),
    aes(x = Gene, y = n, fill = VariantClass2)
) +
geom_bar(stat = 'identity', color = 'black', size = 0.2) +
ylab("Number of Mutations") +
theme_bw() +
theme(
    panel.grid.major = element_blank(),
    panel.border = element_blank(),
    axis.line = element_line(colour = "white"),
    legend.title = element_blank()
) +
theme(axis.text.x = element_text(angle = 45, hjust = 1)) +
scale_fill_nejm() + scale_color_nejm() + labs(title = 'A')

combo = (genes / panel) + plot_layout(ncol = 1, heights = c(1, 1))

do_plot(p = combo, f = "sfig2.png", w = 14, h = 8, save_pdf = T)
```

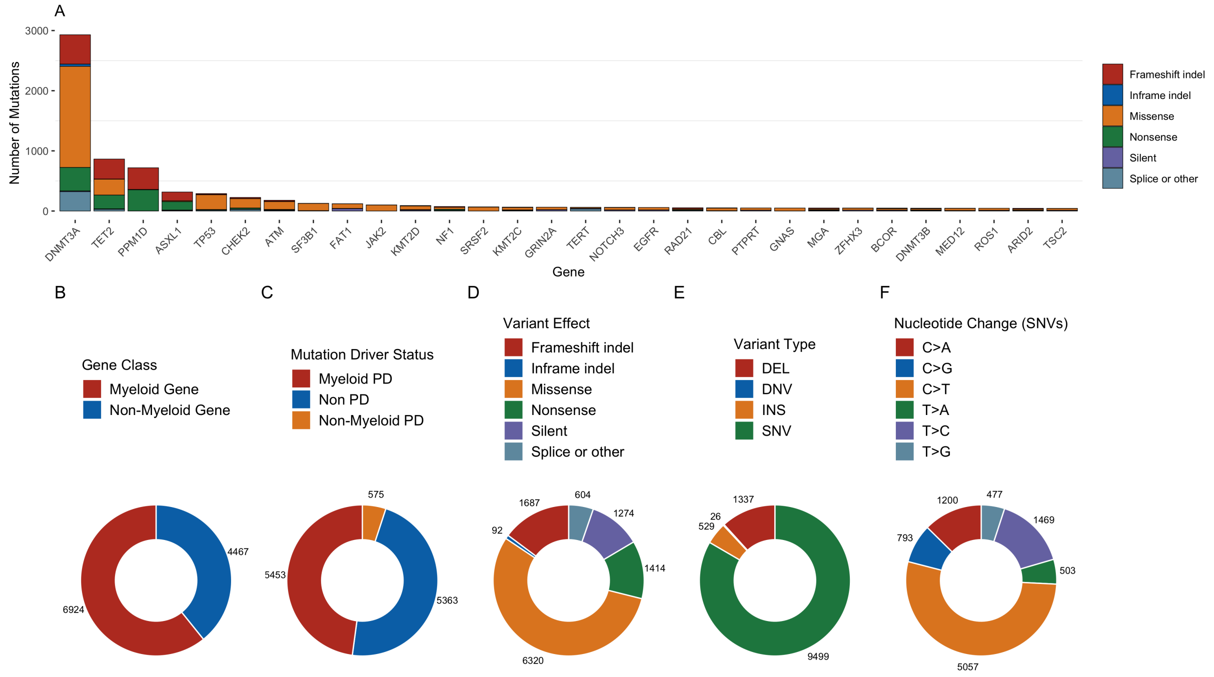


#eFigure 1 - tumor types and CH

```
rare_tumors <- M_wide_all %>%
group_by(generaltumortype) %>%
  summarise(count=n_distinct(MRN)) %>%
  arrange(-count)  %>%
  filter(count<100) %>%
  pull(generaltumortype) %>%
  as.character()

D <- M_wide_all %>% mutate(generaltumortype_r=ifelse(generaltumortype %in% rare_tumors,"Other",as.character(gener
altumortype)))
D <-  D %>% mutate(generaltumortype_r = relevel(factor(generaltumortype_r), "Breast Carcinoma"))

N <- as.data.frame(table(D$generaltumortype_r)) %>% dplyr::rename(term = Var1)

logit <- D %>%
  glm(formula = CH_all ~ age + generaltumortype_r, family = binomial(link="logit"), na.action = 'na.omit')

results <- logit %>% get_model_data()


results <- results %>% filter(term!="age") %>%
        mutate(term=str_remove(term, "generaltumortype_r")) %>%
        arrange(estimate) %>%

        mutate(
        q.value = p.adjust(p.value, n = nrow(.)),
        q.label = paste0(signif(estimate, 2), signif.num(q.value)))

results_bind <- left_join(results,N)
```

```
## Warning: Column `term` joining character vector and factor, coercing into
## character vector
```

```
results_bind <- results_bind %>% mutate(term = paste0(term, ' (n = ', Freq, ')'))
results_bind <- results_bind %>% mutate(term = factor(term, levels = term))

p = plot_forest(
      results_bind,
      x = "term",
      eb_w = 0,
      eb_s = 1,
      ps = 2,
      or_s = 3,
      label = 'p.stars'
  ) +
  xlab('') + ylab('OR') +
  scale_fill_nejm() +
  scale_color_nejm() +
  scale_x_discrete(expand = c(0.025, 0))

do_plot(p, "efig1.png", 8, 5, save_pdf = T)
```

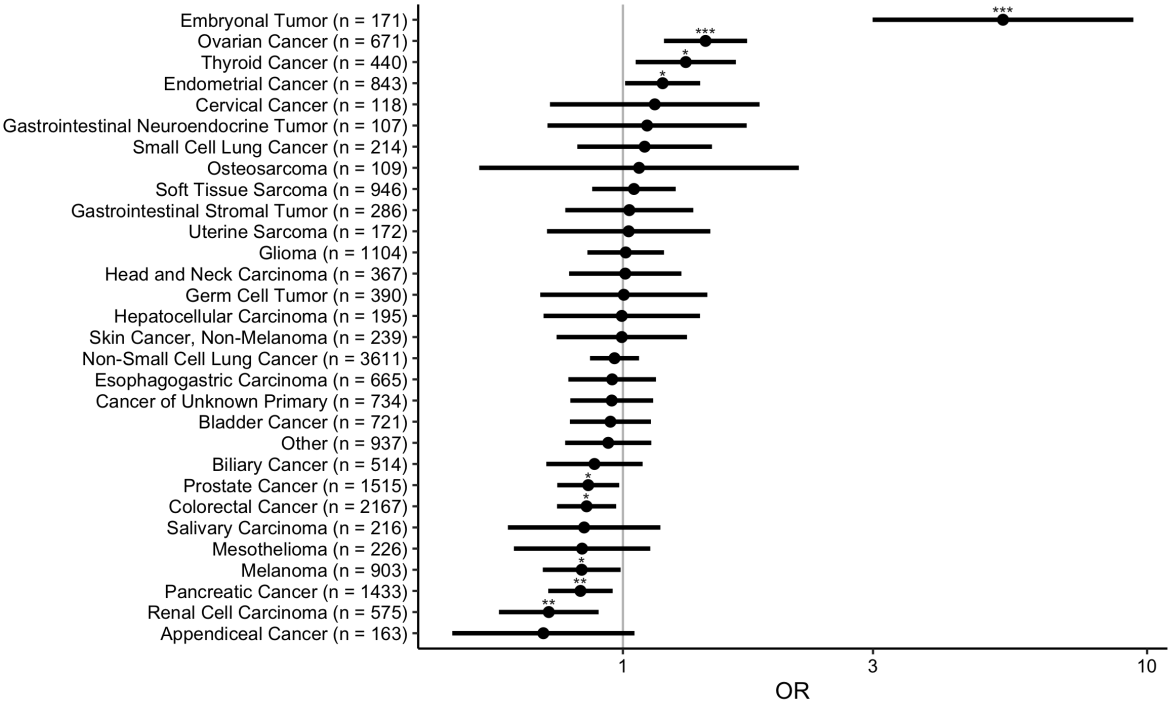


#eFigure 2 - prop CH by tumor type

```
options(repr.plot.width = 10, repr.plot.height = 6, repr.plot.res = 300)

library(beyonce)

n_dict = M_wide_all %>% count(generaltumortype) %>%
    arrange(-n) %>% {setNames(.$n, .$generaltumortype)}

top_genes <- M_long %>%
group_by(Gene) %>%
  summarise(count=n_distinct(MRN)) %>%
  arrange(-count)  %>%
  filter(count>75) %>%
  pull(Gene) %>%
  as.character()

gene_list <- M_long %>%
group_by(Gene) %>%
  summarise(count=n_distinct(MRN)) %>%
  arrange(-count)  %>%
  filter(count>75) %>%
  pull(Gene)

tumor_list = M_wide_all %>% count(generaltumortype) %>%
    arrange(-n) %>%
    top_n(12) %>%
    pull(generaltumortype) %>%
    as.character()

D <- M_long %>%
filter(Gene %in% top_genes) %>%
count(generaltumortype, Gene) %>%
filter(!is.na(generaltumortype)) %>%
group_by(generaltumortype) %>%
mutate(prop = n/n_dict[generaltumortype]) %>%
filter(generaltumortype %in% tumor_list) %>%
ungroup() %>%
mutate(Gene = factor(Gene, top_genes))


D_wide <- M_wide_all %>% filter(generaltumortype %in% tumor_list)

p <- D %>%
ggplot(
    aes(x = Gene, y = prop, fill = generaltumortype)
) +
geom_bar(stat = 'identity', color = 'black', size = 0.2, position = position_dodge()) +
theme_bw() +
theme(
    legend.position = "top",
    panel.grid.major = element_blank(),
    panel.border = element_blank(),
    axis.line = element_line(colour = "white"),
    legend.title = element_blank()
) +
theme(axis.text.x = element_text(angle = 45, hjust = 1)) +
scale_fill_brewer(palette = "Paired") +
ylab('Proportion With Mutation')

do_plot(p, "efig2.png", 8, 5, save_pdf = T)
```

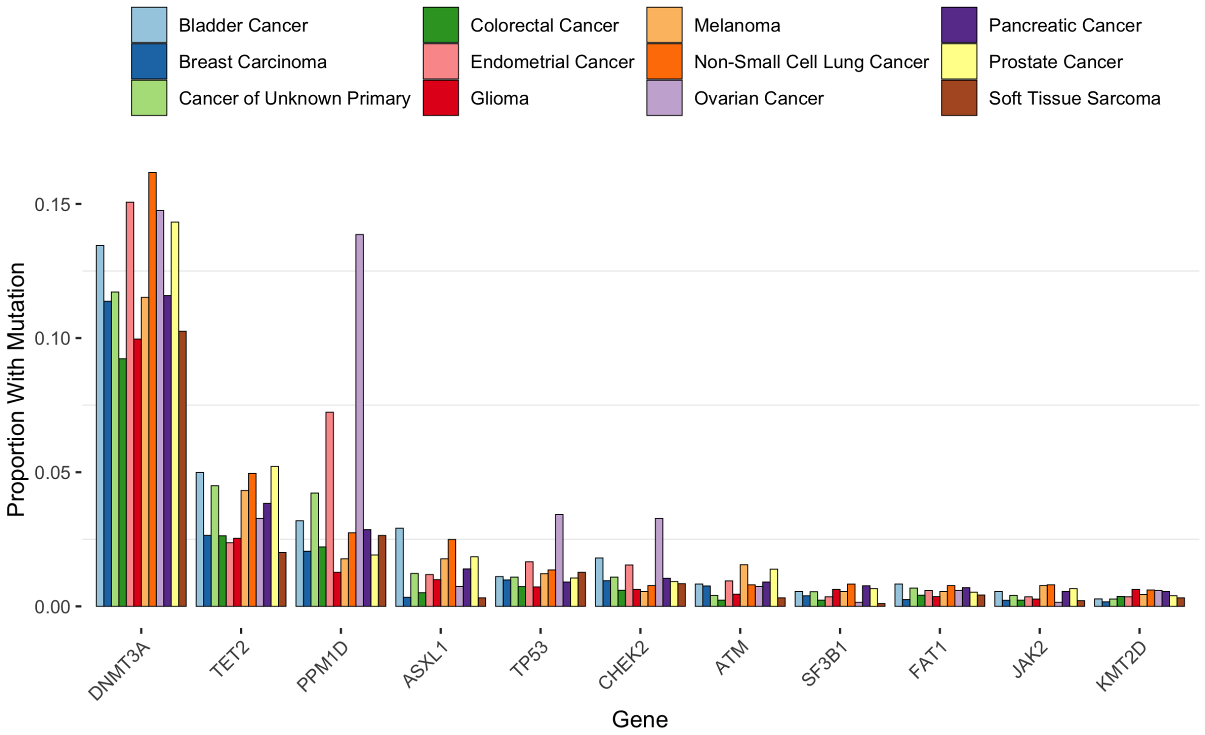


#eFigure 3 - therapy distribution

```
font_size = 13

tumor_list = M_wide_all %>% count(generaltumortype) %>%
    arrange(-n) %>%
    top_n(15) %>%
    pull(generaltumortype) %>%
    as.character()

D = M_wide %>%
    filter(generaltumortype %in% tumor_list) %>%
    mutate(treatment_type = case_when(
        XRT == 1 & ind_anychemo == 1 ~ "XRT & Systemic",
        XRT == 1 ~ "XRT",
        ind_anychemo == 1 ~ "Systemic Therapy",
        T ~ "No treatment")) %>%
    mutate(
        generaltumortype = factor(
            generaltumortype,
            levels = names(sort(table(generaltumortype), decreasing = F)))
    )

p1 = ggplot(D) +
  geom_bar(aes(x = generaltumortype, fill = treatment_type), color = 'black') +
  theme(
      axis.text.x = element_text(angle = 45, vjust = 1, hjust = 1, size = font_size),
      axis.text.y = element_text(size = font_size),
      legend.position = c(0.8, 0.3),
      plot.title = element_text(hjust = .5, size = 16),
      axis.title=element_text(size = 14),
      legend.text = element_text(size = font_size),
      legend.title = element_blank()
  ) +
  ylab("Frequency") +
  xlab("") +
  scale_x_discrete(expand = c(0, 0)) +
  scale_y_continuous(expand = c(0, 0)) +
  coord_flip()+
  scale_fill_manual(values = c(
      "No treatment" = 'gray',
      'Systemic Therapy' = pal_npg()(4)[1],
      'XRT & Systemic' = pal_npg()(4)[2],
      'XRT' = pal_npg()(4)[3]
    )
  ) +
  scale_color_nejm()

D = M_wide %>% mutate(
        Treatment = case_when(
            ind_cytotoxic_therapy == 1 & ind_immune_therapy == 0 &
               ind_targeted_therapy == 0 & ind_radiotherapy == 0 ~ "Only Cytotoxic",
            ind_cytotoxic_therapy == 0 & ind_immune_therapy == 1 &
               ind_targeted_therapy == 0 & ind_radiotherapy == 0 ~ "Only Immune",
            ind_cytotoxic_therapy == 0 & ind_immune_therapy == 1 &
               ind_targeted_therapy == 1 & ind_radiotherapy == 0 ~ "Only Targeted",
            ind_cytotoxic_therapy == 0 & ind_immune_therapy == 0 &
               ind_targeted_therapy == 0 & ind_radiotherapy == 1 ~ "Only Radiotherapy",
            ind_anychemo == 0 ~ "NA",
            T ~ "Multiple"
    )) %>%
    filter(Treatment != "NA") %>%
    filter(generaltumortype %in% tumor_list) %>%
    mutate(
        generaltumortype = factor(
            generaltumortype,
            levels = names(sort(table(generaltumortype), decreasing = F)))
    )

p2 = ggplot(D) +
  geom_bar(aes(x = generaltumortype, fill = Treatment), color = 'black') +
  xlab("") +
  theme(
      axis.text.x = element_text(angle = 45, vjust = 1, hjust = 1, size = font_size),
      axis.text.y = element_text(size = font_size),
      legend.position = c(0.8,0.3),
      plot.title = element_text(hjust = .5, size = 16),
      axis.title=element_text(size = 14),
      legend.text = element_text(size = 13),
      legend.title = element_blank()
  ) +
```

```
  scale_x_discrete(expand = c(0, 0)) +
  scale_y_continuous(expand = c(0, 0)) +
  coord_flip() +
  scale_fill_nejm() +
  scale_color_nejm()

options(repr.plot.width = 10, repr.plot.height = 5, repr.plot.res = 300)

exclude_list = c(
    "ind_carboplatin", "ind_oxaliplatin", "ind_cisplatin",
    "ind_gnrh_antagonist",  "ind_anychemo", "ind_cytotoxic_therapy",
    "ind_hormonal_therapy", "ind_antiestrogen", "ind_targeted_therapy",
    "ind_nitrogen_mustard", "ind_vegf_inhibitor", "ind_aromatase_inhibitor",
    "ind_immune_checkpoint_in", "ind_anti_pd1_antibody", "ind_carbo_pacli",
    "ind_antiandrogen", "ind_camptothecins", "ind_epipodophyllotoxins",
    "ind_vinca_alkaloids", 'ind_protein_kinase_inhib', 'ind_vascular_targeted_th',
    'ind_immune_therapy'
)

D = M_wide %>%
    select(str_subset(colnames(.), '^ind')) %>%
    replace(., is.na(.), 0) %>%
    colSums() %>%
    as.data.frame %>%
    setNames('freq') %>%
    tibble::rownames_to_column('drug')  %>%
    arrange(-freq) %>%
    filter(!(drug %in% exclude_list)) %>%
    filter(!str_detect(drug, regex('_ds_|post'))) %>%
    mutate(drug = format_variable(drug)) %>%
    mutate(drug = factor(drug, drug))

p3 = ggplot(
      D[1:10,]
  ) +
  geom_col(
      aes(x = drug, y = freq), color = 'black'
  ) +
  theme(
      plot.title = element_text(hjust = .5, size = 16),
      axis.text.x = element_text(angle = 45, vjust = 1, hjust = 1, size = font_size),
      axis.title = element_text(size=14),
      axis.text.y = element_text(size=13)
  ) +
  xlab("") +
  ylab("Frequency")  +
  scale_fill_nejm() +
  scale_color_nejm() +
  scale_x_discrete(expand = c(0, 0)) +
  scale_y_continuous(expand = c(0, 0))

p <- egg::ggarrange(
    p1, p2, p3,
    labels = c('A', 'B', 'C'),
    label.args = list(gp = grid::gpar(fontface = 'bold', fontsize = 15)),
    draw = F
)

do_plot(p, "efig3.png", w = 12, h = 12, save_pdf = T)
```

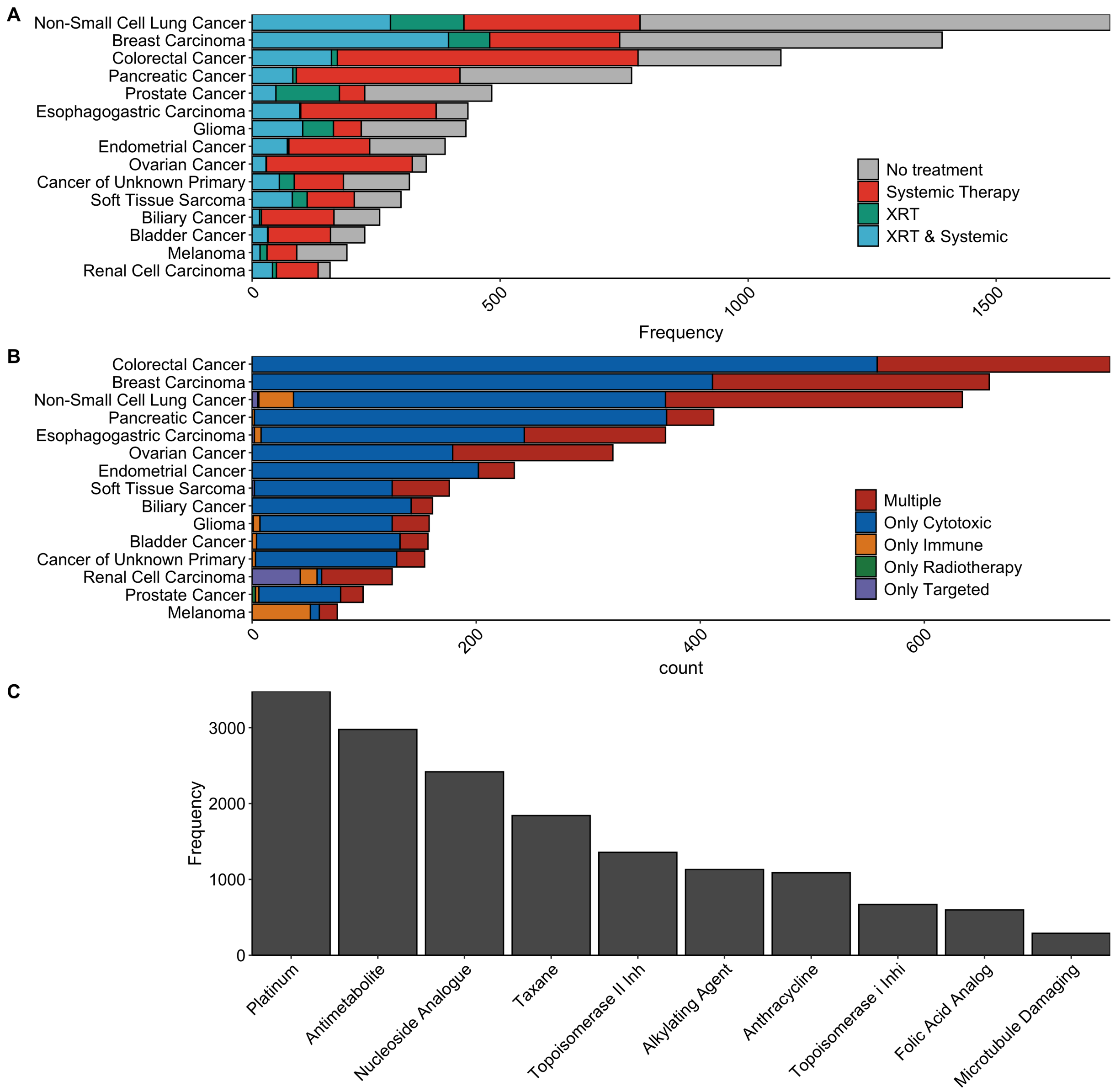


```
#number of pts recieving multiple classes of cytotoxic therapy
c <- M_wide %>% mutate(sum=ind_platinum+ind_antimetabolite+ind_nucleoside_analogue+ind_taxane+ind_topoisomerase_i
i_inh+ind_alkylating_agent+ind_anthracycline+ind_topoisomerase_i_inhi+ind_folic_acid_analog+ind_microtubule_damag
ing)
c <- c %>% mutate(mult=ifelse(sum>1,2,sum))
table(c$mult)
```

```
##
##    0    1    2
## 5238  443 4526
```

#eTable 2/3 - associations with CH

```
source('../utils/toolbox.R')
#All
D = M_wide %>%
    mutate(smoke_verbose = ifelse(smoke_bin == 1, 'Smoker', 'Non-smoker')) %>%
    mutate(smoke_verbose = factor(smoke_verbose, c('Non-smoker', 'Smoker'))) %>%
    mutate(age = age_d)

summary_all = D %>%
    glm(formula = CH_all ~ age + Gender + smoke_verbose + race + therapy_binary,
        family = binomial(link = "logit"), na.action = 'na.omit') %>%
    summar(CI = T, truncate_p = T)
```

```
## Warning in ifelse(as.numeric(pval) < 0.05 | pval == "<1e-06", T, F): NAs
## introduced by coercion
```

```
#My and Nonmy
summary_my = D %>%
    glm(formula = CH_my ~ age + Gender + smoke_verbose + race + therapy_binary,
        family = binomial(link = "logit"), na.action = 'na.omit') %>%
    summar(CI = T, truncate_p = T)
```

```
## Warning in ifelse(as.numeric(pval) < 0.05 | pval == "<1e-06", T, F): NAs
## introduced by coercion
```

```
summary_nonmy = D %>%
    glm(formula = ch_nonmy ~ age + Gender + smoke_verbose + race + therapy_binary,
        family = binomial(link = "logit"), na.action = 'na.omit') %>%
    summar(CI = T, truncate_p = T)
```

```
## Warning in ifelse(as.numeric(pval) < 0.05 | pval == "<1e-06", T, F): NAs
## introduced by coercion
```

```
#Silent and Nonsilent
summary_silent = D %>%
    glm(formula = CH_silent ~ age + Gender + smoke_verbose + race + therapy_binary,
        family = binomial(link = "logit"), na.action = 'na.omit') %>%
    summar(CI = T, truncate_p = T)
```

```
## Warning in ifelse(as.numeric(pval) < 0.05 | pval == "<1e-06", T, F): NAs
## introduced by coercion
```

```
summary_nonsilent = D %>%
    glm(formula = CH_nonsilent ~ age + Gender + smoke_verbose + race + therapy_binary,
        family = binomial(link="logit"), na.action = 'na.omit') %>%
    summar(CI = T, truncate_p = T)
```

```
## Warning in ifelse(as.numeric(pval) < 0.05 | pval == "<1e-06", T, F): NAs
## introduced by coercion
```

```
#My PD and My nonPD
summary_mypd = D %>%
    glm(formula = ch_my_pd ~ age + Gender + smoke_verbose + race + therapy_binary,
        family = binomial(link="logit"), na.action = 'na.omit') %>%
    summar(CI = T, truncate_p = T)
```

```
## Warning in ifelse(as.numeric(pval) < 0.05 | pval == "<1e-06", T, F): NAs
## introduced by coercion
```

```
summary_mynonpd = D %>%
    glm(formula = ch_my_nonpd ~ age + Gender + smoke_verbose + race + therapy_binary,
        family = binomial(link="logit"), na.action = 'na.omit') %>%
    summar(CI = T, truncate_p = T)
```

```
## Warning in ifelse(as.numeric(pval) < 0.05 | pval == "<1e-06", T, F): NAs
## introduced by coercion
```

```
#NonMy PD and NonMY nonPD
summary_nonmypd = D %>%
    glm(formula = ch_nonmy_pd ~ age + Gender + smoke_verbose + race + therapy_binary,
        family = binomial(link="logit"), na.action = 'na.omit') %>%
    summar(CI = T, truncate_p = T)
```

```
## Warning in ifelse(as.numeric(pval) < 0.05 | pval == "<1e-06", T, F): NAs
## introduced by coercion
```

```
summary_nonmynonpd = D %>%
    glm(formula = ch_nonmy_nonpd ~ age + Gender + smoke_verbose + race + therapy_binary,
        family = binomial(link="logit"), na.action = 'na.omit') %>%
    summar(CI = T, truncate_p = T)
```

```
## Warning in ifelse(as.numeric(pval) < 0.05 | pval == "<1e-06", T, F): NAs
## introduced by coercion
```

```
# Myeloid vs non myeloid
summary_my_het = D %>% filter(CH_all == 1) %>%
    glm(formula = CH_my ~ age + Gender + smoke_verbose + race + therapy_binary,
        family = binomial(link="logit"), na.action = 'na.omit') %>%
    summar(CI = T, truncate_p = T) %>%
    select(variable, levels, pval)
```

```
## Warning in ifelse(as.numeric(pval) < 0.05 | pval == "<1e-06", T, F): NAs
## introduced by coercion
```

```
# Silent vs nonsilent
summary_si_het = D %>% filter(CH_all == 1) %>%
    glm(formula = CH_nonsilent ~ age + Gender + smoke_verbose + race + therapy_binary,
        family = binomial(link = "logit"), na.action = 'na.omit') %>%
    summar(CI = T, truncate_p = T) %>%
    select(variable, levels, pval)
```

```
## Warning in ifelse(as.numeric(pval) < 0.05 | pval == "<1e-06", T, F): NAs
## introduced by coercion
```

```
# Myeloid PD vs Myeloid nonPD
summary_mypd_het = D %>% filter(CH_my == 1) %>%
    glm(formula = ch_my_pd ~ age + Gender + smoke_verbose + race + therapy_binary,
        family = binomial(link="logit"), na.action = 'na.omit') %>%
    summar(CI = T, truncate_p = T) %>%
    select(variable, levels, pval)

# Non_Myeloid PD
summary_nonmypd_het = D %>% filter(CH_my == 0) %>%
    glm(formula = ch_nonmy_pd ~ age + Gender + smoke_verbose + race + therapy_binary,
        family = binomial(link="logit"), na.action = 'na.omit') %>%
    summar(CI = T, truncate_p = T) %>%
    select(variable, levels, pval)
```

```
## Warning in ifelse(as.numeric(pval) < 0.05 | pval == "<1e-06", T, F): NAs
## introduced by coercion
```

```
## all, meyloid
summary = Reduce(
        f = function(x, y) {
            full_join(x, y, by = c('variable', 'levels'))
        },
        x = list(
            summary_all,
            summary_my, summary_nonmy, summary_my_het)
    ) %>%
    mutate(levels = ifelse(levels == 'Age', '-', levels)) %>%
    rename(`variable (ref)` = variable) %>%
    rename_all(
        function(x){str_remove(x, '(.x)+|(.y)+')}
    )

summary %>%
kable(format = "latex", booktabs = T, align = 'l', escape = F) %>%
kable_styling(
    latex_options = c("hold_position"),
    full_width = T,
    font_size = 10
) %>%
column_spec(1, width = 100) %>%
column_spec(2:50, width = 45) %>%
add_header_above(
    c(" " = 2, "All CH" = 3, "Myeloid CH" = 3, "Non-Myeloid CH" = 3,
      "Heterogeneity" = 1)
) %>%
collapse_rows(columns = 1:2, row_group_label_position = 'identity') %>%
display_kable('etable2.1.pdf')
```

| | | All CH | | | Myeloid CH | | | Non-Myeloid CH | | | Heterogeneity |
|---|---|---|---|---|---|---|---|---|---|---|---|
| variable (ref) | levels | OR | 95% CI | pval | OR | 95% CI | pval | OR | 95% CI | pval | pval |
| Age | - | 1.80 | 1.8-1.9 | **<1e-06** | 1.90 | 1.8-2 | **<1e-06** | 1.60 | 1.5-1.7 | **<1e-06** | **<1e-06** |
| Gender (Male) | Female | 1.10 | 0.98-1.2 | 0.15 | 1.10 | 0.97-1.2 | 0.17 | 1.10 | 0.94-1.2 | 0.34 | 0.79 |
| Smoke (Non-Smoker) | Smoker | 1.10 | 1-1.3 | **0.0041** | 1.10 | 0.99-1.2 | 0.086 | 1.20 | 1.1-1.4 | **0.0059** | 0.46 |
| Race (White) | Asian | 0.72 | 0.6-0.88 | **9e-04** | 0.68 | 0.54-0.85 | **0.00075** | 0.70 | 0.53-0.94 | **0.017** | 0.18 |
| | Black | 1.00 | 0.84-1.2 | 0.83 | 1.10 | 0.86-1.3 | 0.58 | 0.79 | 0.58-1.1 | 0.12 | 0.5 |
| | Missing | 1.00 | 0.77-1.3 | 0.93 | 0.94 | 0.69-1.3 | 0.67 | 1.20 | 0.85-1.7 | 0.29 | 0.51 |
| | Other | 0.99 | 0.81-1.2 | 0.89 | 1.10 | 0.84-1.3 | 0.66 | 0.87 | 0.65-1.2 | 0.38 | 0.35 |
| Therapy (Untreated) | Treated | 1.20 | 1.1-1.4 | **4.2e-06** | 1.30 | 1.2-1.4 | **4.7e-06** | 1.20 | 1-1.3 | **0.016** | 0.11 |

```
## silent
summary = Reduce(
        f = function(x, y) {
            full_join(x, y, by = c('variable', 'levels'))
        },
        x = list(
            summary_silent, summary_nonsilent, summary_si_het)
    ) %>%
    mutate(levels = ifelse(levels == 'Age', '-', levels)) %>%
    rename(`variable (ref)` = variable) %>%
    rename_all(
        function(x){str_remove(x, '(.x)+|(.y)+')}
    )

summary %>%
kable(format = "latex", booktabs = T, align = 'l', escape = F) %>%
kable_styling(
    latex_options = c("hold_position"),
    full_width = T,
    font_size = 10
) %>%
column_spec(1, width = 100) %>%
column_spec(2:50, width = 45) %>%
add_header_above(
    c(" " = 2, "Silent CH" = 3, "Non-Silent CH" = 3,
      "Heterogeneity" = 1)
) %>%
collapse_rows(columns = 1:2, row_group_label_position = 'identity') %>%
display_kable('etable2.2.pdf')
```

| | | Silent CH | | | Non-Silent CH | | | Heterogeneity |
|---|---|---|---|---|---|---|---|---|
| variable (ref) | levels | OR | 95% CI | pval | OR | 95% CI | pval | pval |
| Age | - | 1.60 | 1.5-1.8 | **<1e-06** | 1.90 | 1.8-2 | **<1e-06** | **<1e-06** |
| Gender (Male) | Female | 1.20 | 0.96-1.4 | 0.13 | 1.10 | 0.99-1.2 | 0.072 | 0.24 |
| Smoke (Non-Smoker) | Smoker | 1.30 | 1.1-1.6 | **0.0084** | 1.20 | 1-1.3 | **0.0044** | 0.65 |
| Race (White) | Asian | 0.78 | 0.52-1.2 | 0.24 | 0.72 | 0.59-0.87 | **0.00092** | 0.61 |
| | Black | 0.93 | 0.62-1.4 | 0.72 | 1.00 | 0.83-1.2 | 0.95 | 0.76 |
| | Missing | 1.00 | 0.59-1.8 | 0.95 | 0.99 | 0.75-1.3 | 0.97 | 0.74 |
| | Other | 0.91 | 0.59-1.4 | 0.68 | 1.00 | 0.82-1.2 | 0.95 | 0.6 |
| Therapy (Untreated) | Treated | 1.10 | 0.91-1.3 | 0.33 | 1.30 | 1.1-1.4 | **2.8e-06** | 0.32 |

```
## My PD
summary = Reduce(
        f = function(x, y) {
            full_join(x, y, by = c('variable', 'levels'))
        },
        x = list(summary_mypd, summary_mynonpd, summary_mypd_het)
    ) %>%
    mutate(levels = ifelse(levels == 'Age', '-', levels)) %>%
    rename(`variable (ref)` = variable) %>%
    rename_all(
        function(x){str_remove(x, '(.x)+|(.y)+')}
    )

summary %>%
kable(format = "latex", booktabs = T, align = 'l', escape = F) %>%
kable_styling(
    latex_options = c("hold_position"),
    full_width = T,
    font_size = 10
) %>%
column_spec(1, width = 100) %>%
column_spec(2:50, width = 45) %>%
add_header_above(
    c(" " = 2, "Myeloid PD CH" = 3, "Myeloid Non-PD CH" = 3,
      "Heterogeneity" = 1)
) %>%
collapse_rows(columns = 1:2, row_group_label_position = 'identity') %>%
display_kable('etable2.2.pdf')
```

| | | Myeloid PD CH | | | Myeloid Non-PD CH | | | Heterogeneity |
|---|---|---|---|---|---|---|---|---|
| variable (ref) | levels | OR | 95% CI | pval | OR | 95% CI | pval | pval |
| Age | - | 1.90 | 1.8-2 | **<1e-06** | 1.80 | 1.7-2 | **<1e-06** | 0.8 |
| Gender (Male) | Female | 1.10 | 0.94-1.2 | 0.35 | 1.30 | 1-1.5 | **0.015** | 0.45 |
| Smoke (Non-Smoker) | Smoker | 1.00 | 0.93-1.2 | 0.54 | 1.30 | 1.1-1.6 | **0.011** | **0.043** |
| Race (White) | Asian | 0.65 | 0.51-0.83 | **0.00058** | 0.86 | 0.57-1.3 | 0.47 | 0.28 |
| | Black | 1.00 | 0.8-1.3 | 0.88 | 1.00 | 0.67-1.5 | 0.95 | 0.46 |
| | Missing | 0.96 | 0.69-1.3 | 0.81 | 1.40 | 0.83-2.3 | 0.21 | 0.82 |
| | Other | 1.10 | 0.87-1.4 | 0.43 | 0.94 | 0.6-1.5 | 0.8 | 0.32 |
| Therapy (Untreated) | Treated | 1.30 | 1.2-1.5 | **7.3e-06** | 1.30 | 1.1-1.6 | **0.011** | 0.32 |

```
## Non Myeloid PD
summary = Reduce(
        f = function(x, y) {
            full_join(x, y, by = c('variable', 'levels'))
        },
        x = list(summary_nonmypd, summary_nonmynonpd, summary_nonmypd_het)
    ) %>%
    mutate(levels = ifelse(levels == 'Age', '-', levels)) %>%
    rename(`variable (ref)` = variable) %>%
    rename_all(
        function(x){str_remove(x, '(.x)+|(.y)+')}
    )

summary %>%
kable(format = "latex", booktabs = T, align = 'l', escape = F) %>%
kable_styling(
    latex_options = c("hold_position"),
    full_width = T,
    font_size = 10
) %>%
column_spec(1, width = 100) %>%
column_spec(2:50, width = 45) %>%
add_header_above(
    c(" " = 2, "Non-Myeloid PD CH" = 3, "Non-Myeloid Non-PD CH" = 3,
      "Heterogeneity" = 1)
) %>%
collapse_rows(columns = 1:2, row_group_label_position = 'identity') %>%
display_kable('etable2.3.pdf')
```

| | | Non-Myeloid PD CH | | | Non-Myeloid Non-PD CH | | | Heterogeneity |
|---|---|---|---|---|---|---|---|---|
| variable (ref) | levels | OR | 95% CI | pval | OR | 95% CI | pval | pval |
| Age | - | 1.80 | 1.5-2.1 | **<1e-06** | 1.60 | 1.5-1.7 | **<1e-06** | **<1e-06** |
| Gender (Male) | Female | 0.85 | 0.57-1.3 | 0.42 | 1.10 | 0.96-1.3 | 0.18 | 0.58 |
| Smoke (Non-Smoker) | Smoker | 1.70 | 1.1-2.5 | **0.02** | 1.20 | 1-1.4 | **0.017** | 0.057 |
| Race (White) | Asian | 0.92 | 0.4-2.1 | 0.85 | 0.70 | 0.52-0.95 | **0.021** | 0.42 |
| | Black | 0.72 | 0.26-2 | 0.52 | 0.82 | 0.6-1.1 | 0.19 | 0.31 |
| | Missing | 0.66 | 0.16-2.7 | 0.56 | 1.30 | 0.89-1.8 | 0.18 | 0.7 |
| | Other | 0.80 | 0.29-2.2 | 0.67 | 0.86 | 0.63-1.2 | 0.36 | 0.8 |
| Therapy (Untreated) | Treated | 1.30 | 0.89-2 | 0.15 | 1.20 | 1-1.3 | **0.023** | 0.074 |

#Supp Figure 3 - CH mutational features and VAF

```
treatment_palette = c(pal_nejm()(2)[2], pal_nejm()(2)[1])

D = M %>% filter(VAF_N < 0.35) %>%
    filter(CH_nonsilent == 1) %>%
    mutate(CH_my = ifelse(CH_my == 1, 'Myeloid gene', 'Non-myeloid gene')) %>%
    mutate(pd_status = ifelse(ch_pancan_pd == 1, 'PD', 'Non-PD')) %>%
    mutate(pd_status = factor(pd_status))

D = D %>% group_by(MRN) %>% dplyr::mutate(mutnum=n())

D = D %>% mutate(mutnum = ifelse(mutnum > 1, '2+', as.character(mutnum)))

panel_theme = theme_bw() + theme(
    panel.grid.minor = element_blank(),
    panel.border = element_blank(),
    axis.line = element_line(colour = "black"),
    legend.title = element_blank(),
    plot.title = element_text(size = 8)
)

model_my = geeglm(
    data = D,
    formula = log(VAF_N) ~ age_scaled + race + smoke_bin + therapy_binary + CH_my,
    family = gaussian,
    corstr = "exchangeable",
    id = MRN)

pval_my = model_my %>% summary %>% coefficients %>% .['CH_myNon-myeloid gene', 'Pr(>|W|)']

p_my = ggplot(
        D,
        aes(y = VAF_N, x = CH_my, fill = therapy_binary)
    ) +
    geom_boxplot(outlier.alpha = 0) +
    geom_point(position = position_jitterdodge(), pch = 21, color = 'black', size = 1) +
    panel_theme + xlab("") + ylab('VAF') +
    scale_fill_manual(values = treatment_palette) +
    geom_signif(
      comparisons = list(levels(as.factor(D$CH_my))),
      y_position = 0.36,
      vjust = -0.5,
      tip_length = 0.01,
      annotation = paste0('p = ', scientific(pval_my, digits = 2)),
      map_signif_level = TRUE,
      textsize = 3
    ) +
    scale_y_continuous(expand = c(0.1, 0))

model_pd = geeglm(
    data = D,
    formula = log(VAF_N) ~ age_scaled + race + smoke_bin + therapy_binary + ch_pancan_pd,
    family = gaussian,
    corstr = "exchangeable",
    id = MRN)

pval_pd = model_pd %>% summary %>% coefficients %>% .['ch_pancan_pd', 'Pr(>|W|)']

p_pd = ggplot(
      D,
      aes(y = VAF_N, x = pd_status, fill = therapy_binary)
  ) +
  geom_boxplot(outlier.alpha = 0) +
  geom_point(position = position_jitterdodge(), pch = 21, color = 'black', size = 1) +
  panel_theme + xlab("") + ylab('VAF') +
  scale_fill_manual(values = treatment_palette) +
  geom_signif(
    comparisons = list(levels(D$pd_status)),
    y_position = 0.36,
    vjust = -0.5,
    tip_length = 0.01,
    annotation = paste0('p = ', scientific(pval_pd, digits = 2)),
    map_signif_level = TRUE,
    textsize = 3
  ) +
  scale_y_continuous(expand = c(0.1, 0))

model_mutnum = geeglm(
    data = D,
    formula = log(VAF_N) ~ age_scaled + race + smoke_bin + therapy_binary + mutnum,
```

```
    family = gaussian,
    corstr = "exchangeable",
    id = MRN)

pval_mutnum = model_mutnum %>% summary %>% coefficients %>% .['mutnum2+', 'Pr(>|W|)']

p_mutnum = ggplot(
      D,
      aes(x = factor(mutnum), y = VAF_N, fill = therapy_binary)
  ) +
  geom_boxplot(outlier.alpha = 0) +
  theme_bw() +
  theme(
      panel.grid.minor = element_blank(),
      panel.border = element_blank(),
      axis.line = element_line(colour = "black"),
      strip.background = element_rect(fill = "white", color = 'white'),
      legend.title = element_blank()
  ) +
  ylab('VAF') +
  geom_point(position = position_jitterdodge(), pch = 21, color = 'black', size = 1) +
  scale_fill_manual(values = treatment_palette) +
  xlab('Number of mutations') +
  geom_signif(
    comparisons = list(levels(factor(D$mutnum))),
    y_position = 0.36,
    vjust = -0.5,
    tip_length = 0.01,
    annotation = paste0('p = ', scientific(pval_mutnum, digits = 2)),
    map_signif_level = TRUE,
    textsize = 3
  ) +
  scale_y_continuous(expand = c(0.1, 0))

panel <-  ggarrange(
        p_my, p_pd, p_mutnum,
        ncol = 3,
        nrow = 1,
        common.legend = TRUE,
        legend = "top", align = 'hv',
        labels = c('A', 'B', 'C')
    )

do_plot(panel, "SuppFig3.png", w = 9, h = 4, save_pdf = T)
```

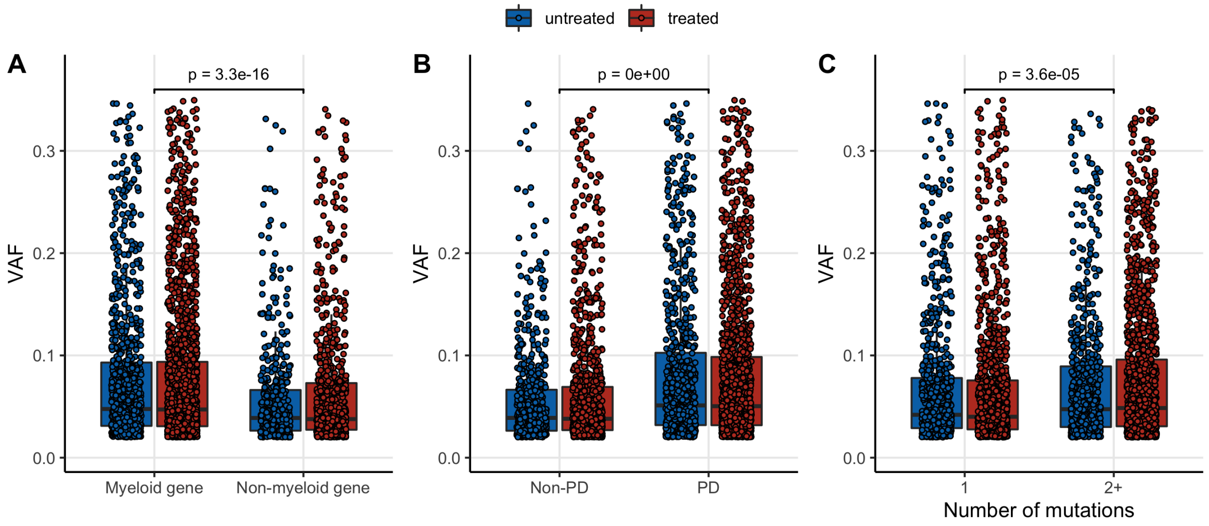


#eTable 3 - associations with VAF

```
D = M_all %>%
    filter(!is.na(smoke_bin)) %>%
    mutate(
        smoke_bin = factor(ifelse(smoke_bin == 1, 'Smoker', 'Non-smoker'), c('Non-smoker', 'Smoker'))
    ) %>%
    mutate(
        pd_status = case_when(
            ch_my_pd==1  ~ "Myeloid PD",
            ch_nonmy_pd==1 ~ "Non-Myeloid PD",
            ch_my_pd==0 & ch_nonmy_pd==0 & myeloid_gene==1  ~ "Non-PD Myeloid",
            ch_my_pd==0 & ch_nonmy_pd==0 & myeloid_gene==0  ~ "Non-PD Non-Myeloid"
        )
    ) %>%
    mutate(pd_status = relevel(factor(pd_status), ref = 'Non-PD Non-Myeloid')) %>%
    group_by(MRN) %>%
    mutate(mutnum = as.character(min(n(), 2))) %>%
    mutate(age = age_d) %>%
    ungroup()

summary_multi = geeglm(
    data = D,
    formula = log(VAF_N) ~ age + race + smoke_bin + therapy_binary + pd_status + mutnum,
    family = gaussian,
    corstr = "exchangeable",
    id = MRN
) %>% summar(truncate_p = T)
```

```
## Warning in ifelse(as.numeric(pval) < 0.05 | pval == "<1e-06", T, F): NAs
## introduced by coercion
```

```
summary_multi %>%
mutate(levels = ifelse(levels == '2', '2+', levels)) %>%
mutate(levels = ifelse(levels == 'Age', '-', levels)) %>%
rename(`variable (ref)` = variable) %>%
kable(
    format = "latex", booktabs = T, align = 'l', escape = F, linesep = ''
) %>%
kable_styling(
    latex_options = c("basic", "hold_position"),
    full_width = T,
    font_size = 10
) %>%
column_spec(1:10, width = 120) %>%
collapse_rows(columns = 1:2, row_group_label_position = 'identity') %>%
display_kable('etable3.pdf')
```

| variable (ref) | levels | OR | 95% CI | pval |
|---|---|---|---|---|
| Age | - | 1.00 | 1-1.1 | **0.0037** |
| Race (White) | Asian | 1.00 | 0.93-1.1 | 0.6 |
| | Black | 0.90 | 0.82-0.99 | **0.039** |
| | Other | 0.90 | 0.82-1 | 0.053 |
| | Unknown | 0.93 | 0.83-1.1 | 0.27 |
| Smoke (Non-Smoker) | Smoker | 1.10 | 1.1-1.2 | **1.3e-05** |
| Therapy (Untreated) | Treated | 1.00 | 0.96-1.1 | 0.74 |
| PD Status (Non-PD Non-Myeloid) | Myeloid PD | 1.30 | 1.3-1.4 | **<1e-06** |
| | Non-Myeloid PD | 1.30 | 1.2-1.5 | **1.6e-05** |
| | Non-PD Myeloid | 0.98 | 0.92-1 | 0.61 |
| Mutnum (1) | 2+ | 1.10 | 1.1-1.2 | **2.8e-06** |

#eTable 4 - associations with mutation number

```
D = M_wide %>%
    mutate(
        race_b = factor(ifelse(race_b == 1, 'White', 'Other'), c('White', 'Other')),
        smoke_bin = factor(ifelse(smoke_bin == 1, 'Smoker', 'Non-smoker'), c('Non-smoker', 'Smoker')),
        age = age_d
    )

summary_all = D %>%
    filter(mutnum_all > 0) %>%
    MASS::polr(
        data = .,
        formula = factor(mutnum_all) ~ age_scaled + Gender + race_b + smoke_bin + therapy_binary,
        Hess = TRUE,
        method = "logistic") %>%
    summar(truncate_p = T)
```

```
## Warning in ifelse(as.numeric(pval) < 0.05 | pval == "<1e-06", T, F): NAs
## introduced by coercion
```

```
summary_all %>%
rename(`variable (ref)` = variable) %>%
mutate(levels = ifelse(levels == 'Age', '-', levels)) %>%
kable(format = "latex", booktabs = T, align = 'l', escape = F) %>%
kable_styling(
    latex_options = c("hold_position"),
    full_width = T,
    font_size = 10
) %>%
column_spec(1, width = 100) %>%
column_spec(2:50, width = 40) %>%
collapse_rows(columns = 1:2, row_group_label_position = 'identity') %>%
display_kable('etable4.pdf')
```

| variable (ref) | levels | OR | 95% CI | pval |
|---|---|---|---|---|
| Age | - | 2.30 | 2-2.6 | **<1e-06** |
| Gender (Male) | Female | 1.10 | 0.95-1.3 | 0.18 |
| Race (White) | Other | 0.84 | 0.67-1 | 0.1 |
| Smoke (Non-Smoker) | Smoker | 1.20 | 1-1.4 | **0.019** |
| Therapy (Untreated) | Treated | 1.20 | 1.1-1.5 | **0.0096** |

#Supplementary Fig. 4 - mutational characteristics

```
# Proportion of myeloid mutations
C = M %>%
    mutate(myeloid_category = case_when(
      CH_my == 1 & CH_nonsilent == 1 ~ "Myeloid Gene",
      CH_my == 0 & CH_nonsilent == 1 ~ "Non-Myeloid Gene",
      TRUE ~ "NA"
    )) %>%
    count(myeloid_category, therapy_binary) %>%
    filter(myeloid_category != "NA") %>%
    group_by(therapy_binary) %>%
    mutate(total = sum(n)) %>%
    mutate(proportion = n/total) %>%
    mutate(myeloid_category = factor(myeloid_category, levels = c('Myeloid Gene', 'Non-Myeloid Gene')))

label_size = 3

p_my = ggplot(
    C,
      aes(x = therapy_binary, y = proportion, fill = myeloid_category, label = n)
  ) +
  geom_bar(stat = 'identity') + theme_bw() +
  theme(
      panel.grid.major = element_blank(),
      panel.grid.minor = element_blank(),
      panel.border = element_blank(),
      axis.line = element_line(colour = "black"),
      legend.title = element_blank(),
  ) + xlab('') + coord_flip() +
  geom_text(size = label_size, position = position_stack(vjust = 0.5), color = 'white') +
  theme(strip.background = element_rect(fill = "white", color = 'white')) +
  scale_fill_nejm() + scale_color_nejm() +
  scale_y_continuous(expand = c(0,0)) +
  scale_x_discrete(expand = c(0,0))

#(b) Proportion of putative drivers

C = M_all %>%
    mutate(pd_category = case_when(
      ch_my_pd == 1 ~ "Myeloid PD",
      ch_my_pd == 0 & ch_pancan_pd == 1 ~ "Non-Myeloid PD",
      TRUE ~ "Non PD"
    )) %>%
    count(pd_category, therapy_binary) %>%
    filter(pd_category != "NA") %>%
    group_by(therapy_binary) %>%
    mutate(total = sum(n)) %>%
    mutate(proportion = n/total) %>%
    mutate(pd_category = factor(pd_category, levels = c('Myeloid PD', 'Non-Myeloid PD', 'Non PD')))

p_pd = ggplot(
      C,
      aes(x = therapy_binary, y = proportion, fill = pd_category, label = n)
  ) +
  geom_bar(stat = 'identity') + theme_bw() +
  theme(
      panel.grid.major = element_blank(),
      panel.grid.minor = element_blank(),
      panel.border = element_blank(),
      axis.line = element_line(colour = "black"),
      legend.title = element_blank()
  ) + xlab('') + coord_flip() +
  geom_text(size = label_size, position = position_stack(vjust = 0.5), color = 'white') +
  theme(strip.background = element_rect(fill = "white", color = 'white')) +
  scale_fill_nejm() + scale_color_nejm() +
  scale_y_continuous(expand = c(0,0)) +
  scale_x_discrete(expand = c(0,0))

#(c) Proportion of silent mutations
C = M_all %>%
    count(CH_silent, therapy_binary) %>%
    group_by(therapy_binary) %>%
    mutate(total = sum(n)) %>%
    mutate(proportion = n/total) %>%
    mutate(CH_silent = ifelse(CH_silent == 1, 'Silent', 'Non-silent'))

p_silent = ggplot(
      C,
      aes(x = therapy_binary, y = proportion, fill = CH_silent, label = n)
  ) +
```

```
  geom_bar(stat = 'identity') + theme_bw() +
  theme(
      panel.grid.major = element_blank(),
      panel.grid.minor = element_blank(),
      panel.border = element_blank(),
      axis.line = element_line(colour = "black"),
      legend.title = element_blank()
  ) + xlab('') + ylab('') + coord_flip() +
  geom_text(size = label_size, position = position_stack(vjust = 0.5), color = 'white') +
  theme(strip.background = element_rect(fill = "white", color = 'white')) +
  scale_fill_nejm() + scale_color_nejm() +
  scale_y_continuous(expand = c(0,0)) +
  scale_x_discrete(expand = c(0,0))

#(d) Variant class
# tally
D = M_all %>% mutate(therapy_binary = factor(therapy_binary, c('untreated', 'treated'))) %>%
    mutate(
        VariantClass = case_when(
            VariantClass == 'Frame_Shift_Del' ~ 'Frameshift indel',
            VariantClass == 'Frame_Shift_Ins' ~ 'Frameshift indel',
            VariantClass == 'In_Frame_Del' ~ 'Inframe indel',
            VariantClass == 'In_Frame_Ins' ~ 'Inframe indel',
            VariantClass == 'Splice_Site' ~ 'Splice or other',
            VariantClass == 'Splice_Region' ~ 'Splice or other',
            VariantClass == "5'Flank" ~ 'Splice or other',
            VariantClass == 'Translation_Start_Site' ~ 'Splice or other',
            VariantClass == 'Nonstop_Mutation' ~ 'Splice or other',
            VariantClass == 'Missense_Mutation' ~ 'Missense',
            VariantClass == 'Nonsense_Mutation' ~ 'Nonsense',
            T ~ VariantClass)
    ) %>%
    group_by(Gene) %>% mutate(n_gene = n()) %>% ungroup() %>%
    filter(n_gene > 20) %>%
    arrange(-n_gene) %>% mutate(Gene = factor(Gene, unique(Gene))) %>%
    count(VariantClass, therapy_binary) %>%
    group_by(therapy_binary) %>%
        mutate(total = sum(n)) %>%
        mutate(proportion = n/total) %>%
    ungroup()

# plot
p_vc = ggplot(
      D,
      aes(x = therapy_binary, y = proportion, fill = VariantClass, label = n)
  ) +
  geom_bar(stat = 'identity') +
  theme(axis.text.x = element_text(angle = 45, hjust = 1)) +
  coord_flip() +
  theme_bw() +
  theme(
      panel.grid.major = element_blank(),
      panel.grid.minor = element_blank(),
      panel.border = element_blank(),
      axis.line = element_line(colour = "black"),
      legend.title = element_blank()
  ) +
  scale_fill_nejm() + scale_color_nejm() +
  scale_y_continuous(expand = c(0,0)) +
  scale_x_discrete(expand = c(0,0)) +
  xlab('') + ylab('') +
  geom_text(size = label_size, position = position_stack(vjust = 0.5), color = 'white')

#(e) Nucleotide substitution
C = M %>% filter(variant_type == 'SNV') %>%
    count(sub_nuc, therapy_binary) %>%
    group_by(therapy_binary) %>%
    mutate(total = sum(n)) %>%
    mutate(proportion = n/total)

p_nuc = ggplot(
      C,
      aes(x = therapy_binary, y = proportion, fill = sub_nuc, label = n)
  ) +
  geom_bar(stat = 'identity') + theme_bw() +
  theme(
      panel.grid.major = element_blank(),
      panel.grid.minor = element_blank(),
      panel.border = element_blank(),
      axis.line = element_line(colour = "black"),
```

```
      legend.title = element_blank()
  ) + xlab('') + coord_flip() +
  geom_text(size = label_size, position = position_stack(vjust = 0.5), color = 'white') +
  scale_fill_nejm() + scale_color_nejm() +
  scale_y_continuous(expand = c(0,0)) +
  scale_x_discrete(expand = c(0,0))
# grid.arrange(p_nuc, p_type, ncol = 1)

#(f) Mutation type vs treatment type
C = M %>% filter(variant_type %in% c('INS', 'DEL', 'SNV')) %>%
    mutate(
        treatment_type = case_when(
            XRT == 0 & ind_anychemo == 1 ~ "Chemo no XRT",
            XRT == 1 ~ "XRT",
            XRT == 0 & ind_anychemo == 0 ~ "No treatment",
            T ~ "NA")
    ) %>%
    filter((!is.na(treatment_type)) & (treatment_type != 'NA')) %>%
    mutate(treatment_type = factor(treatment_type, levels = c('Chemo no XRT', 'XRT', 'No treatment'))) %>%
    count(variant_type, treatment_type) %>%
    group_by(treatment_type) %>%
    mutate(total = sum(n)) %>%
    mutate(proportion = n/total)

p_indel = ggplot(
      C,
      aes(x = treatment_type, y = proportion, fill = variant_type, label = n)
  ) +
  geom_bar(stat = 'identity') + theme_bw() +
  theme(
      panel.grid.major = element_blank(),
      panel.grid.minor = element_blank(),
      panel.border = element_blank(),
      axis.line = element_line(colour = "black"),
      legend.title = element_blank()
  ) + xlab('') + coord_flip() +
  geom_text(size = label_size, position = position_stack(vjust = 0.5), color = 'white') +
  scale_fill_nejm() + scale_color_nejm() +
  scale_y_continuous(expand = c(0,0)) +
  scale_x_discrete(expand = c(0,0))

#(g) Indel length
C = M %>% filter(variant_type %in% c('DEL', 'INS')) %>%
    mutate(
        treatment_type = case_when(
            XRT == 0 & ind_anychemo == 1 ~ "Chemo no XRT",
            XRT == 1 ~ "XRT",
            XRT == 0 & ind_anychemo == 0 ~ "No treatment",
            T ~ "NA")
    ) %>%
    mutate(treatment_type = factor(treatment_type, levels = c('Chemo no XRT', 'XRT', 'No treatment'))) %>%
    mutate(indel_length = as.numeric(End) - as.numeric(Start)) %>%
    mutate(
        indel_length_bin = cut(
            x = indel_length,
            breaks = c(-Inf, 1, 5, Inf),
            labels = c('1', '2-5', '>5'))
    ) %>%
    count(indel_length_bin, treatment_type) %>%
    group_by(treatment_type) %>%
    mutate(total = sum(n)) %>%
    mutate(proportion = n/total)

p_len = ggplot(
      C,
      aes(x = treatment_type, y = proportion, fill = indel_length_bin, label = n)
  ) +
  geom_bar(stat = 'identity') + theme_bw() +
  theme(
      panel.grid.major = element_blank(),
      panel.grid.minor = element_blank(),
      panel.border = element_blank(),
      axis.line = element_line(colour = "black")
  ) + xlab('') + coord_flip() +
  geom_text(size = label_size, position = position_stack(vjust = 0.5), color = 'white') +
  guides(fill = guide_legend(title = 'Indel Length (nts)')) +
  scale_fill_nejm() + scale_color_nejm() +
  scale_y_continuous(expand = c(0,0)) +
  scale_x_discrete(expand = c(0,0))
```

```
panel = ggarrange(p_my, p_pd, p_silent, p_nuc, p_indel, p_len, p_vc,
          ncol = 2, nrow = 4, common.legend = F, legend = "top", align = 'hv', labels = 'AUTO')

do_plot(panel, 'supp_fig4.png', w = 11, h = 9, save_pdf = T)
```

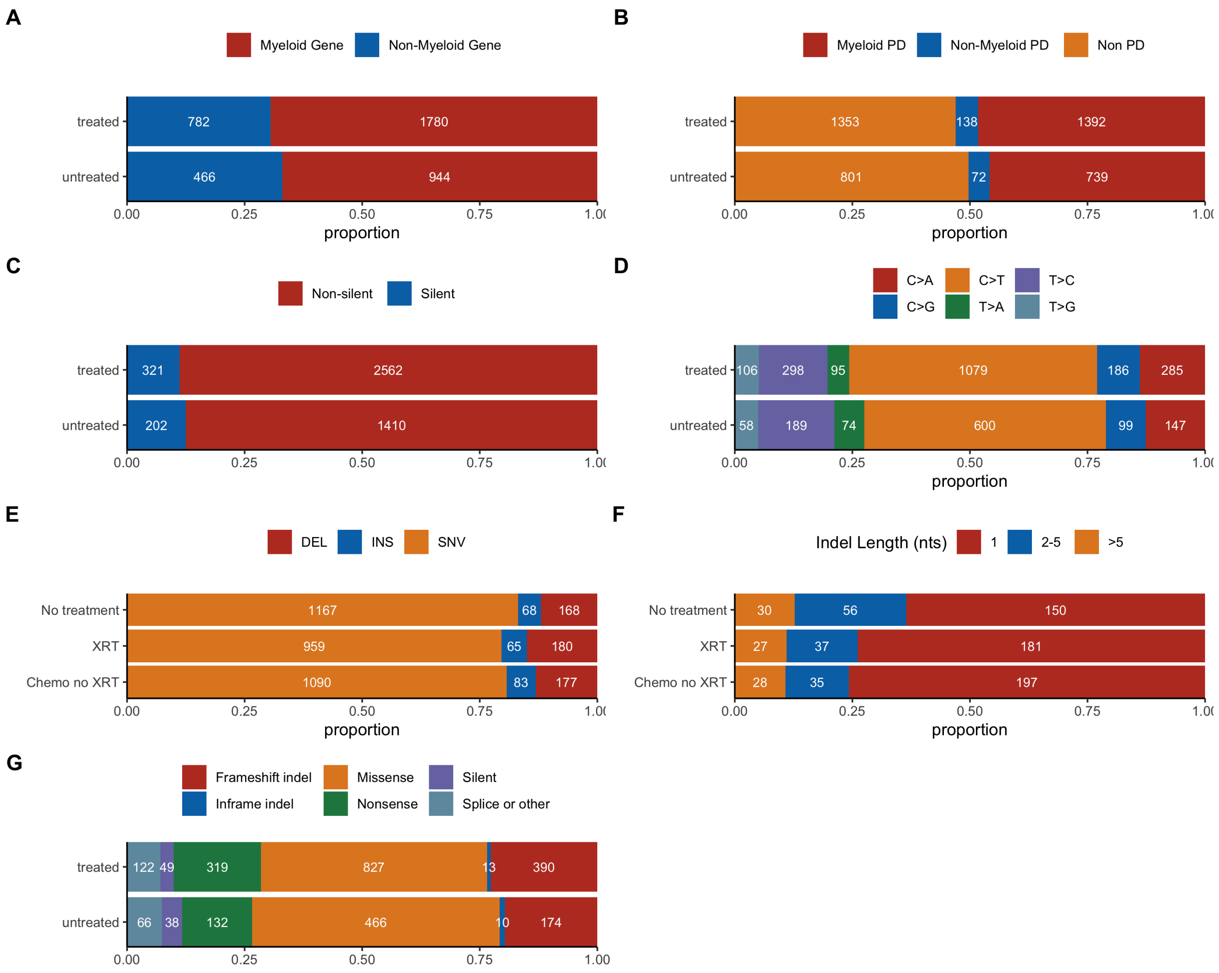


#Supplementary Fig. 5 - mutational context

```
source('../mutational-descriptive-characteristics/plotSignature.R')
preBase = rep(c(rep('A', 4), rep('C', 4), rep('G', 4), rep('T', 4)),6)
refBase = c(rep('C', 48 ), rep('T', 48))
altBase = c(rep('A', 16 ), rep('G', 16), rep('T', 16),rep('A', 16 ), rep('C', 16), rep('G', 16))
postBase = rep(c('A', 'C', 'G', 'T'), 96/4)
mut.order = paste0(preBase, '[',refBase, '>', altBase, ']', postBase )

M_snv = M %>% filter(variant_type == 'SNV') %>%
    mutate(
        triplets = paste0(
            substr(context_5, nchar(context_5), nchar(context_5)),
            Ref,
            substr(context_3, 1, 1)
        )
    ) %>% get_context()

font_size = 5

p_sig_all = M_snv %>%
  {rbind(
    get_context_table(., mut.order) %>%
      mutate(bin = 'All mutations'),
    filter(., XRT == 0 & ind_anychemo == 0 & smoke_bin == 0) %>%
      get_context_table(mut.order) %>%
      mutate(bin = 'No smoking, untreated'),
    filter(., XRT == 1 | ind_anychemo == 1 & smoke_bin == 0) %>%
      get_context_table(mut.order) %>%
      mutate(bin = 'Treated, no smoking'),
    filter(., XRT == 0 & ind_anychemo == 0 & smoke_bin == 1) %>%
      get_context_table(mut.order) %>%
      mutate(bin = 'Smoking, untreated')
  )} %>%
  plot_signature() +
  facet_wrap(~bin, ncol = 1, scale = 'free_y') +
  ggtitle('') +
  ylab('Number of mutations') +
  panel_theme +
  theme(
    legend.position = 'none',
    axis.text.x = element_text(angle = 90, size = font_size)
  )

p_sig_ddr = M_snv %>%
  filter(tCH) %>%
  {rbind(
    get_context_table(., mut.order) %>%
      mutate(bin = 'All mutations'),
    filter(., XRT == 0 & ind_anychemo == 0 & smoke_bin == 0) %>%
      get_context_table(mut.order) %>%
      mutate(bin = 'No smoking, untreated'),
    filter(., XRT == 1 | ind_anychemo == 1 & smoke_bin == 0) %>%
      get_context_table(mut.order) %>%
      mutate(bin = 'Treated, no smoking'),
    filter(., XRT == 0 & ind_anychemo == 0 & smoke_bin == 1) %>%
      get_context_table(mut.order) %>%
      mutate(bin = 'Smoking, untreated')
  )} %>%
  plot_signature() +
  facet_wrap(~bin, ncol = 1, scale = 'free_y') +
  ggtitle('') +
  ylab('') +
  panel_theme +
  theme(
    legend.position = 'none',
    axis.text.x = element_text(angle = 90, size = font_size)
  )

p_sig_asxl1 = M_snv %>%
  filter(Gene == 'ASXL1') %>%
  {rbind(
    get_context_table(., mut.order) %>%
      mutate(bin = 'All mutations'),
    filter(., XRT == 0 & ind_anychemo == 0 & smoke_bin == 0) %>%
      get_context_table(mut.order) %>%
      mutate(bin = 'No smoking, untreated'),
    filter(., XRT == 1 | ind_anychemo == 1 & smoke_bin == 0) %>%
      get_context_table(mut.order) %>%
      mutate(bin = 'Treated, no smoking'),
    filter(., XRT == 0 & ind_anychemo == 0 & smoke_bin == 1) %>%
```

```
      get_context_table(mut.order) %>%
      mutate(bin = 'Smoking, untreated')
  )} %>%
  plot_signature() +
  facet_wrap(~bin, ncol = 1, scale = 'free_y') +
  ggtitle('')  +
  ylab('') +
  panel_theme +
  theme(
    legend.position = 'right',
    axis.text.x = element_text(angle = 90, size = font_size)
  )

## stacked bars
stack_theme = theme_bw() +
  theme(
      panel.grid.minor = element_blank(),
      panel.grid.major = element_blank(),
      panel.border = element_blank(),
      axis.line.x = element_blank(),
      axis.line = element_line(colour = "black"),
      strip.background = element_rect(fill = "white", color = 'white'),
      legend.title = element_blank(),
      legend.position = 'none'
  )


#All
D = M_snv %>%
    filter(!is.na(smoke_bin)) %>%
    count(context96, therapy_binary, smoke_bin) %>%
    mutate(ctpg = str_detect(context96, paste0('C>T', "]", 'G'))) %>%
    group_by(ctpg, therapy_binary, smoke_bin) %>% summarise(n = sum(n)) %>%
    group_by(therapy_binary, smoke_bin) %>%
    mutate(total = sum(n), prop = n/sum(n)) %>%
    mutate(smoking = ifelse(smoke_bin == 1, 'smoking', 'no smoking')) %>%
    mutate(ctpg = ifelse(ctpg, '[C>T]G', 'Other'))

p_stack_all <- ggplot(
      D,
      aes(x = therapy_binary, y = prop, fill = ctpg, group = therapy_binary)
  ) +
  geom_bar(stat = 'identity', color = 'black') + coord_flip() + xlab("") +
  geom_text(aes(label = n), size = 4, position = position_stack(vjust = 0.5), color = 'white') +
  stack_theme +
  facet_wrap(~smoking, ncol = 1) +
  scale_fill_nejm() + scale_color_nejm()

#TCH
D = M_snv %>%
    filter(tCH) %>%
    filter(!is.na(smoke_bin)) %>%
    count(context96, therapy_binary, smoke_bin) %>%
    mutate(ctpg = str_detect(context96, paste0('C>T', "]", 'G'))) %>%
    group_by(ctpg, therapy_binary, smoke_bin) %>% summarise(n = sum(n)) %>%
    group_by(therapy_binary, smoke_bin) %>%
    mutate(total = sum(n), prop = n/sum(n)) %>%
    mutate(smoking = ifelse(smoke_bin == 1, 'smoking', 'no smoking')) %>%
    mutate(ctpg = ifelse(ctpg, '[C>T]G', 'Other'))

p_stack_ddr <- ggplot(
      D,
      aes(x = therapy_binary, y = prop, fill = ctpg, group = therapy_binary)
  ) +
  geom_bar(stat = 'identity', color = 'black') + coord_flip() + xlab("") +
  geom_text(aes(label = n), size = 4, position = position_stack(vjust = 0.5), color = 'white') +
  stack_theme +
  facet_wrap(~smoking, ncol = 1) +
  scale_fill_nejm() + scale_color_nejm()

#ASXL1
D = M_snv %>%
    filter(Gene=="ASXL1") %>%
    filter(!is.na(smoke_bin)) %>%
    count(context96, therapy_binary, smoke_bin) %>%
    mutate(ctpg = str_detect(context96, paste0('C>T', "]", 'G'))) %>%
    group_by(ctpg, therapy_binary, smoke_bin) %>% summarise(n = sum(n)) %>%
    group_by(therapy_binary, smoke_bin) %>%
    mutate(total = sum(n), prop = n/sum(n)) %>%
    mutate(smoking = ifelse(smoke_bin == 1, 'smoking', 'no smoking')) %>%
```

```
    mutate(ctpg = ifelse(ctpg, '[C>T]G', 'Other'))

p_stack_asxl1 <- ggplot(
      D,
      aes(x = therapy_binary, y = prop, fill = ctpg, group = therapy_binary)
  ) +
  geom_bar(stat = 'identity', color = 'black') + coord_flip() + xlab("") +
  geom_text(aes(label = n), size = 4, position = position_stack(vjust = 0.5), color = 'white') +
  stack_theme +
  theme(legend.position = 'right') +
  facet_wrap(~smoking, ncol = 1) +
  scale_fill_nejm() + scale_color_nejm()

panel = ((p_sig_all + ggtitle('All Genes') | p_sig_ddr + ggtitle('DDR') | p_sig_asxl1 + ggtitle('ASXL1') ) / (p_s
tack_all | p_stack_ddr | p_stack_asxl1)) + plot_layout(heights = c(5, 2))

do_plot(panel, 'supp_fig_5.png', w = 20, h = 10, save_pdf = T)
```

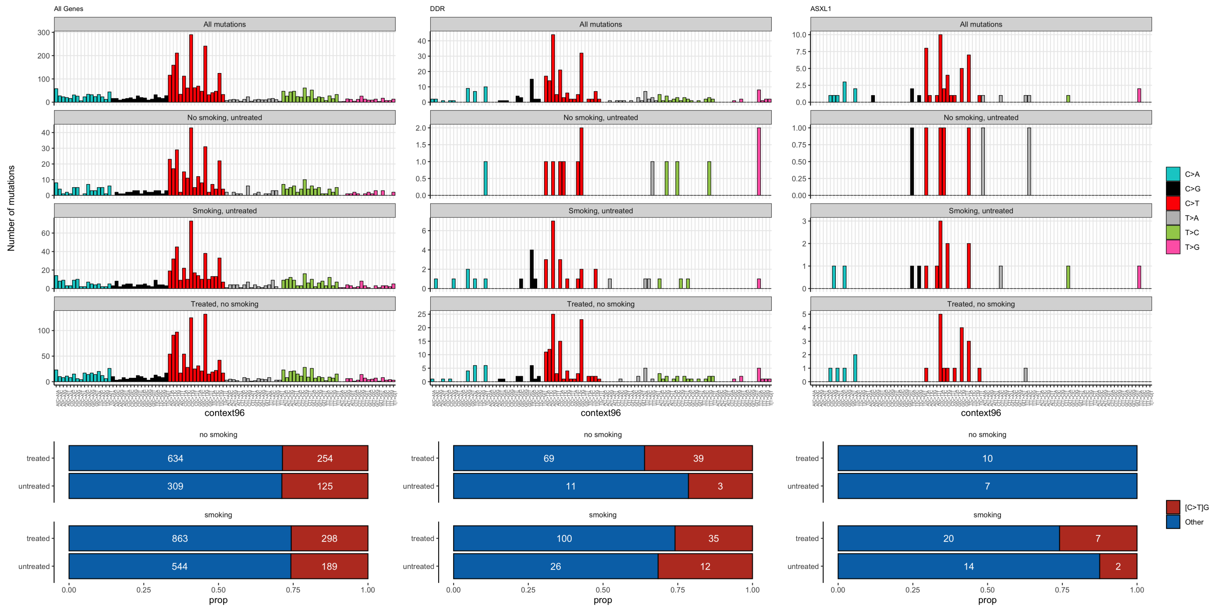


#eFigure 4 - drugset and CH

```
drug_sets =
    get_chemo_column(
        class_dict,
        drugsets_dict,
        class = 'sets',
        prefix = 'ind'
    ) %>%
    {.[. %in% colnames(M_wide)]} %>%
    paste(collapse = ' + ')

M_wide = M_wide %>% mutate(sumSets = rowSums(select(., contains("ind_ds"))))

#ind multivariate
formula = paste0('ch_pancan_pd ~ ', drug_sets, ' + XRT + age + smoke_bin + race_b + timedx_impact')

D = M_wide %>% glm(formula = as.formula(formula), family = binomial(link="logit"), na.action = 'na.omit') %>%
    sjPlot::get_model_data(.) %>%
    cbind(Group = "multi") %>%
    mutate(Drug = term) %>%
    filter(!(term %in% c("age", "smoke_bin1", "race_b", "timedx_impact", "XRT"))) %>%
    mutate(Drug = case_when(
        Drug == "ind_ds_carboplatin" ~ "Carboplatin",
        Drug == "ind_ds_irino_oxali" ~ "Irinotecan Oxaliplatin",
        Drug == "ind_ds_docetaxel" ~ "Docetaxel",
        Drug == "ind_ds_irinotecan" ~ "Irinotecan",
        Drug == "ind_ds_cis_gem" ~ "Cisplatin Gemcitabine",
        Drug == "ind_ds_gemcitabine" ~ "Gemcitabine",
        Drug == "ind_ds_cisplatin" ~ "Cisplatin",
        Drug == "ind_ds_floxuridine" ~ "Floxuridine",
        Drug == "ind_ds_doxorubicin" ~ "Doxorubicin",
        Drug == "ind_ds_cyclo_doxo" ~ "Cyclophosphamide Doxorubicin",
        Drug == "ind_ds_oxaliplatin" ~ "Oxaliplatin",
        Drug == "ind_ds_abr_gem" ~ "Abraxane Gemcitabine",
        Drug == "ind_ds_paclitaxel" ~ "Paclitaxel",
        Drug == "ind_ds_fluo_oxali" ~ "Fluorouracil Oxaliplatin",
        Drug == "ind_ds_carbo_doxo" ~ "Carboplatin Doxorubicin",
        Drug == "ind_ds_carbo_pem" ~ "Carboplatin Pemetrexed",
        Drug == "ind_ds_carbo_pacli" ~ "Carboplatin Paclitaxel",
        Drug == "ind_ds_cis_pem" ~ "Cisplatin Pemetrexed",
        Drug == "ind_ds_carbo_gem" ~ "Carboplatin Gemcitabine",
        Drug == "ind_ds_cis_etop" ~ "Cisplatin Etoposide",
        Drug == "ind_ds_cytotoxic_other" ~ "Other Cytotoxic Drugset")
    ) %>%
    mutate(
        Drug = factor(Drug, levels=unique(Drug[order(estimate)]))
    )

drug_sets_list=
    get_chemo_column(
        class_dict,
        drugsets_dict,
        class = 'sets',
        prefix = 'ind')

p = D %>%
  plot_forest(
      x = "Drug",
      label = "p.stars",
      eb_w = 0,
      eb_s = 1,
      ps = 2,
      or_s = 3.1,
      nudge = -0.5
  ) +
  scale_x_discrete(expand = c(0,1)) +
  ylab("OR of CH-PD") +
  xlab("Regimen") +
  scale_fill_nejm() +
  scale_color_nejm()

do_plot(p, 'efig4.png', w = 6, h = 4, save_pdf = T)
```

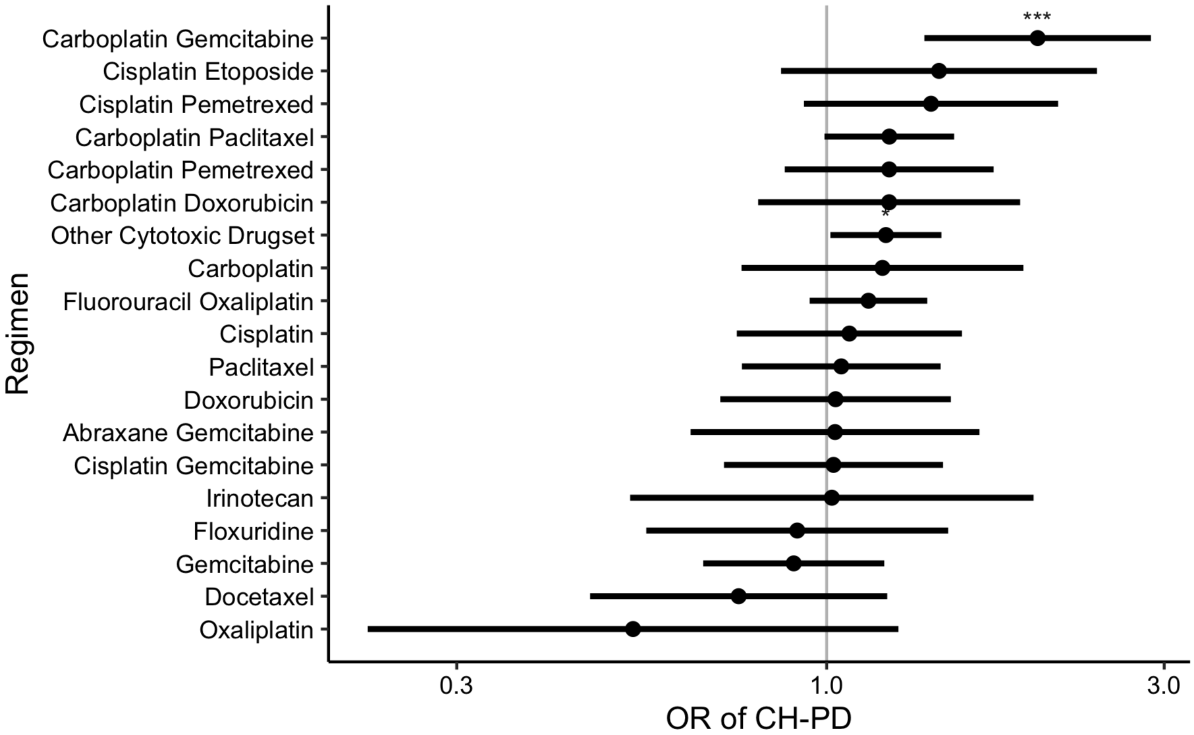


### Tabulate serial

```
#Tabulate serial
print('All patients')
```

```
## [1] "All patients"
```

```
nrow(P_serial)
```

```
## [1] 525
```

```
print('All mutations')
```

```
## [1] "All mutations"
```

```
nrow(M2_all)
```

```
## [1] 621
```

```
print('Therapy Binary')
```

```
## [1] "Therapy Binary"
```

```
table(P_serial$therapy_binary)
```

```
##
##   0   1
## 209 316
```

```
print('# of Mutation Positive Patients')
```

```
## [1] "# of Mutation Positive Patients"
```

```
unique(M2_all$MRN) %>% length()
```

```
## [1] 394
```

```
print('Describe FU')
```

```
## [1] "Describe FU"
```

```
psych::describe(P_serial$delta_time)
```

```
##    vars   n   mean     sd median trimmed    mad min  max range skew
## X1    1 525 733.76 235.92    679  718.31 169.02 186 1602  1416  0.7
##    kurtosis   se
## X1     0.75 10.3
```

```
print('# of Patients who got treatment')
```

```
## [1] "# of Patients who got treatment"
```

```
sum(P_serial$therapy_binary)
```

```
## [1] 316
```

```
print('# of Patients who got no cyto/xrt')
```

```
## [1] "# of Patients who got no cyto/xrt"
```

```
filter(P_serial,therapy_binary==0) %>% nrow()
```

```
## [1] 209
```

```
print('Number of Patients with CH at inital timepoint')
```

```
## [1] "Number of Patients with CH at inital timepoint"
```

```
det <- filter(M2_all,VAF_1>0)
unique(det$MRN) %>% length()
```

```
## [1] 389
```

```
print('Number of mutations detected at both timepoints')
```

```
## [1] "Number of mutations detected at both timepoints"
```

```
filter(M2_all,VAF_1>0 & VAF_2>0) %>% nrow()
```

```
## [1] 590
```

```
print('Number of mutations detected at only one timepoint')
```

```
## [1] "Number of mutations detected at only one timepoint"
```

```
nrow(filter(M2_all,VAF_1==0 | VAF_2==0))
```

```
## [1] 31
```

```
print('Number of mutations detected at first timepoint only')
```

```
## [1] "Number of mutations detected at first timepoint only"
```

```
nrow(filter(M2_all,VAF_1>0 & VAF_2==0))
```

```
## [1] 10
```

```
print('Number of mutations detected at second timepoint only')
```

```
## [1] "Number of mutations detected at second timepoint only"
```

```
nrow(filter(M2_all,VAF_1==0 & VAF_2>0))
```

```
## [1] 21
```

```
print('Number of newly detected mutations after tx')
```

```
## [1] "Number of newly detected mutations after tx"
```

```
new_tx <- filter(M2_all,VAF_1==0 & VAF_2>=0.02 & therapy_binary==1)
unique(new_tx$MRN) %>% length()
```

```
## [1] 13
```

```
print('Number of newly detected mutations after no tx')
```

```
## [1] "Number of newly detected mutations after no tx"
```

```
new_tx <- filter(M2_all,VAF_1==0 & VAF_2>0 & therapy_binary==0)
unique(new_tx$MRN) %>% length()
```

```
## [1] 2
```

```
print('Table of direction')
```

```
## [1] "Table of direction"
```

```
table(M2$direction)
```

```
##
##   DOWN STABLE     UP
##     59    367    164
```

#Supplementary Figure 8 - GR and mutnum

```
p = ggplot(
      M2 %>%
          group_by(MRN) %>%
          mutate(n_mut = n()) %>%
          mutate(any_tch = ifelse(any(tch), 'DDR', 'Other')) %>%
          ungroup() %>%
          filter(n_mut <= 4) %>%
          mutate(therapy = recode(therapy_binary, `1` = 'treated', `0` = 'untreated')) %>%
          mutate(therapy = factor(therapy, levels = c('untreated', 'treated'))) %>%
      {rbind(
        .,
        mutate(., any_tch = 'All')
      )},
      aes(x = n_mut, y = log(growth_rate), group = factor(n_mut))) +
  geom_boxplot(outlier.alpha = 0, color = 'black', fill = 'steelblue') +
  geom_jitter(
      pch = 21,
      size = 1,
      alpha = 0.8,
      width = 0.1,
      color = 'black',
      fill = 'gray'
  ) +
  panel_theme +
  theme(
    legend.position = "none",
    strip.background = element_rect(fill = 'grey', size = 0),
    panel.border = element_rect(color = "grey", fill = NA, size = 1),
    strip.text = element_text(size = 10)

  ) +
  facet_grid(vars(any_tch), vars(therapy)) +
  # scale_fill_nejm() + scale_color_nejm() +
  xlab('number of mutations') +
  ylab('log growth rate')

do_plot(p, 'supp8.png', w = 6, h = 6, save_pdf = T)
```

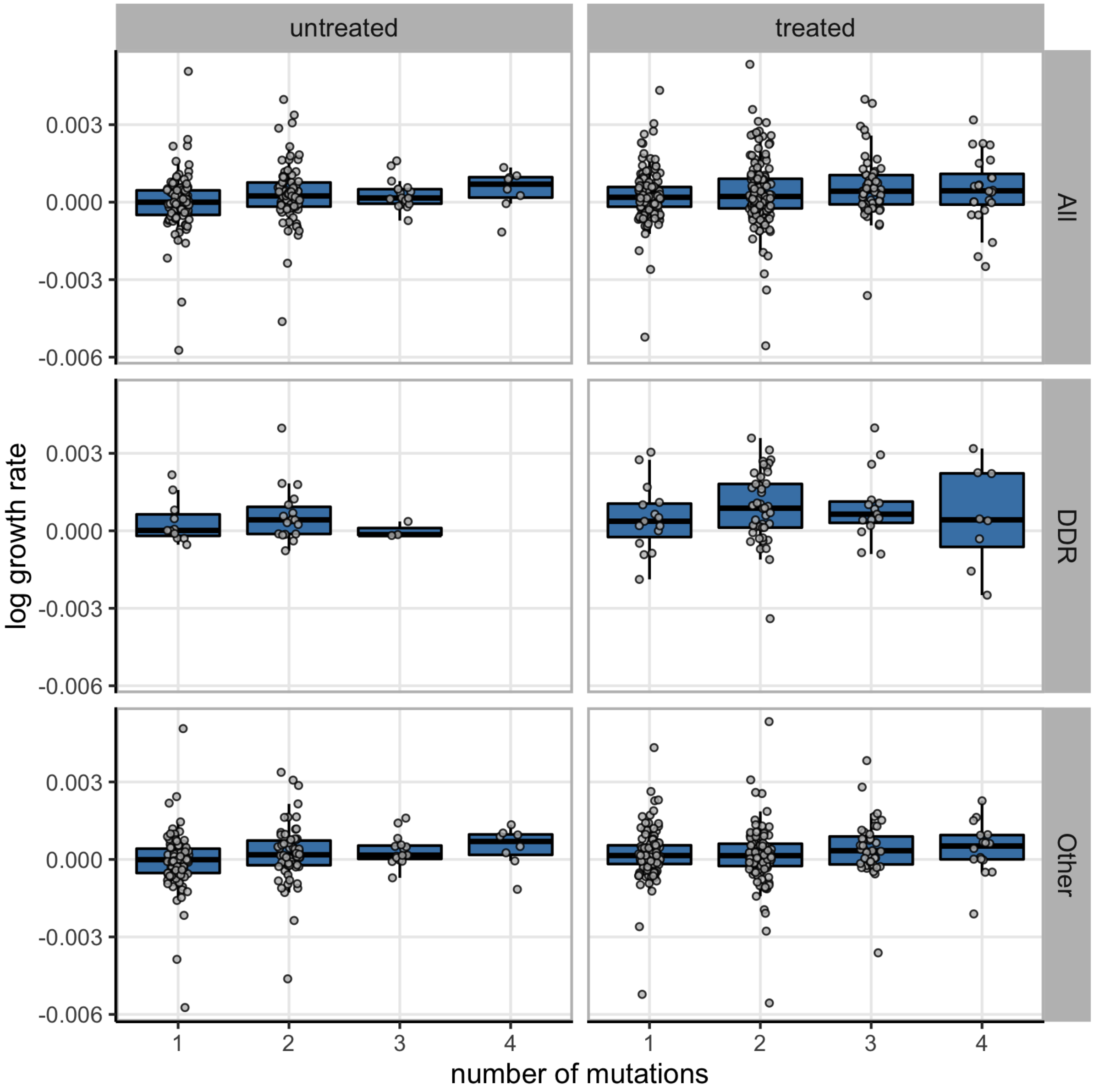


#Supplementary Figure 9 - bar chart singletons

```
P_serial_1 = P_serial %>% left_join(
  M2_all %>% group_by(MRN) %>%
    summarise(
      ch_t1 = any(VAF_1 >= 0.02),
      ch_t2 = any(VAF_2 >= 0.02),
      tch_t1 = any(tch & VAF_1 >= 0.02),
      tch_t2 = any(tch & VAF_2 >= 0.02),
      non_tch_t1 = any(!tch & VAF_1 >= 0.02),
      non_tch_t2 = any(!tch & VAF_2 >= 0.02),
      tch_gain = any(tch & VAF_1 != 0 & VAF_1 < 0.02 & VAF_2 >= 0.02),
      other_gain = any(!tch & VAF_1 != 0 & VAF_1 < 0.02 & VAF_2 >= 0.02),
      tch_loss = any(tch & VAF_1 >= 0.02 & VAF_2 < 0.02),
      tch_loss_complete = any(tch & VAF_1 >= 0.02 & VAF_2 == 0),
      any_loss_complete = any(VAF_1 >= 0.02 & VAF_2 == 0),
      other_loss = any(!tch & VAF_1 >= 0.02 & VAF_2 < 0.02),
      other_loss_complete = any(!tch & VAF_1 >= 0.02 & VAF_2 == 0),
      tch_denovo = any(tch & VAF_1 == 0 & VAF_2 >= 0.02),
      other_denovo = any(!tch & VAF_1 == 0 & VAF_2 >= 0.02),
      any_denovo = any(VAF_1 == 0 & VAF_2 >= 0.02)
    ),
  by = 'MRN',
) %>%
  mutate_at(
    c('ch_t1', 'ch_t2', 'tch_t1', 'tch_t2',
      'non_tch_t1', 'non_tch_t2',
      'tch_gain', 'other_gain', 'tch_loss', 'other_loss',
      'tch_denovo', 'other_denovo'),
    function(x){tidyr::replace_na(x, FALSE)}
  )

D_loss = P_serial_1 %>%
    mutate(
        tch = case_when(
            tch_loss_complete ~ 'complete loss',
            # tch_loss ~ 'loss',
            T ~ 'none'
        ),
        other = case_when(
            other_loss_complete ~ 'complete loss',
            # other_loss ~ 'loss',
            T ~ 'none'
        ),
        any = case_when(
            any_loss_complete ~ 'complete loss',
            # other_loss ~ 'loss',
            T ~ 'none'
        )
    ) %>%
    melt(measure.vars = c('tch', 'other', 'any'),
        variable.name = 'gene',
        value.name = 'event_type') %>%
    count(therapy_binary, gene, event_type) %>%
    group_by(therapy_binary, gene) %>%
    mutate(total = sum(n), prop = n/total)

D_gain = P_serial_1 %>% mutate(
        tch = case_when(
            tch_denovo ~ 'newly observed',
            T ~ 'none'
        ),
        other = case_when(
            other_denovo ~ 'newly observed',
            T ~ 'none'
        ),
        any = case_when(
            any_denovo ~ 'newly observed',
            T ~ 'none'
        )
    ) %>%
    melt(measure.vars = c('tch', 'other', 'any'),
         variable.name = 'gene',
         value.name = 'event_type') %>%
    count(therapy_binary, gene, event_type) %>%
    group_by(therapy_binary, gene) %>%
    mutate(total = sum(n), prop = n/total)

D = rbind(
        D_gain %>% mutate(event = 'gain'),
        D_loss %>% mutate(event = 'loss')
```

```
    ) %>%
    filter(event_type != 'none') %>%
    mutate(event_type = factor(event_type, c('gain', 'newly observed', 'loss', 'complete loss'))) %>%
    ungroup() %>%
    mutate(therapy_binary = ifelse(therapy_binary == 1, 'treated', 'untreated')) %>%
    mutate(gene = case_when(gene == 'tch' ~ 'DDR',
                            gene == 'other' ~ 'Other',
                            gene == 'any' ~ 'Any'))
D <- filter(D,gene=="Any")

p = ggplot(
   D,
    aes(x = therapy_binary,
        y = prop,
        label = n)
) +
geom_bar(stat = 'identity', color = 'black', size = 0.2) +
geom_text(
    size = 3,
    color = 'white',
    position = position_stack(vjust = 0.5)
) +
facet_grid(~event_type) +
theme(
    # strip.background = element_rect(fill = 'white', color = 'black'),
    legend.title = element_blank()
) +
xlab('') +
ylab('proportion of patients') +
scale_fill_nejm()

do_plot(p, "supp_fig9.png", w = 6, h = 4, save_pdf = T)
```

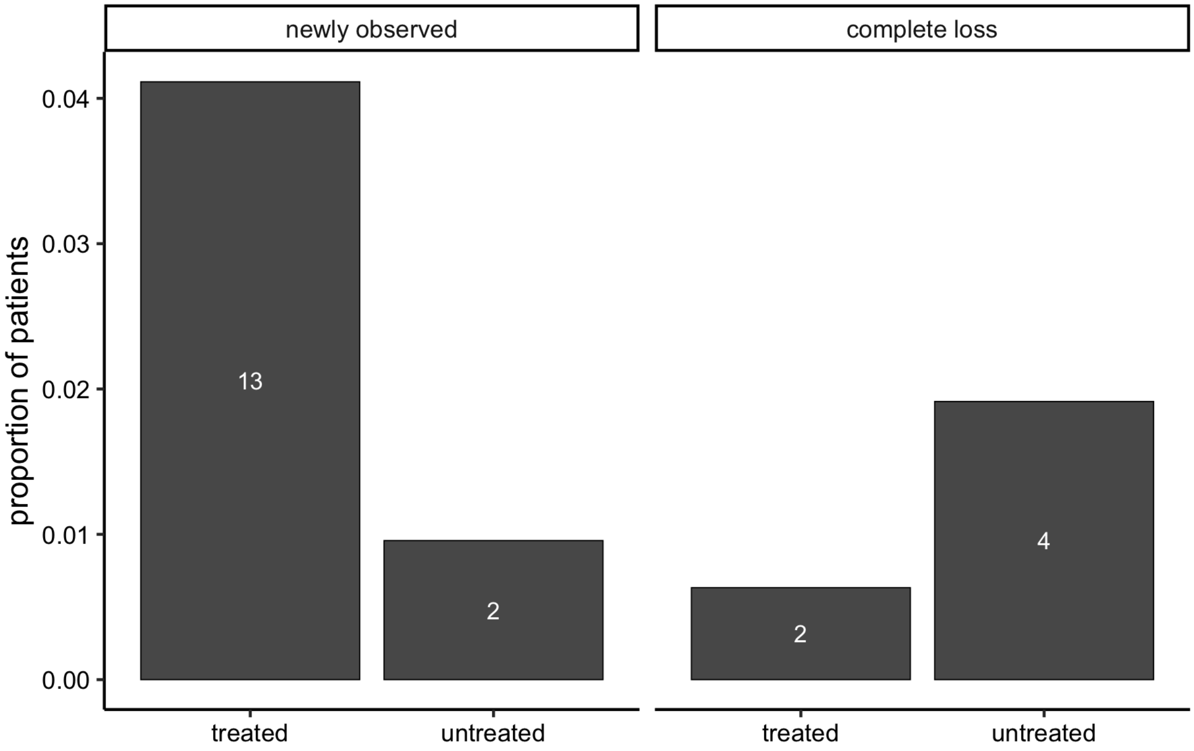


```
D %>% filter(event_type=="newly observed") %>%
group_by(event_type, therapy_binary) %>%
summarise(n = sum(n), total = sum(unique(total))) %>%
  {prop.test(.$n, .$total)}
```

```
##
##  2-sample test for equality of proportions with continuity
##  correction
##
## data:  .$n out of .$total
## X-squared = 3.4514, df = 1, p-value = 0.0632
## alternative hypothesis: two.sided
## 95 percent confidence interval:
##  0.00202691 0.06111282
## sample estimates:
##      prop 1      prop 2
## 0.041139241 0.009569378
```

```
D %>% filter(event_type=="complete loss") %>%
group_by(event_type, therapy_binary) %>%
summarise(n = sum(n), total = sum(unique(total))) %>%
{prop.test(.$n, .$total)}
```

```
## Warning in prop.test(.$n, .$total): Chi-squared approximation may be
## incorrect
```

```
##
##  2-sample test for equality of proportions with continuity
##  correction
##
## data:  .$n out of .$total
## X-squared = 0.86914, df = 1, p-value = 0.3512
## alternative hypothesis: two.sided
## 95 percent confidence interval:
##  -0.03731459  0.01169531
## sample estimates:
##      prop 1      prop 2
## 0.006329114 0.019138756
```

#Supp Table 2 tMN cohort descriptive table

```
d <- M_tmn_wide_st %>% mutate(
  diagnosis=ifelse(post_tmn==1,"tMN","No tMN"))

# Factor the basic variables that
# we're interested in
d$diagnosis <-
  factor(d$diagnosis,
         levels=c("No tMN", "tMN"))

d$Gender <- factor(d$Gender,
                    levels=c("F", "M"),
                   labels=c("Female","Male"))

d$VAF_nonsilent_r <-
  factor(d$VAF_nonsilent_r,
         labels=c("Negative", # Reference
                  "2-5%",
                  "5-10%",
                  "10-20%",
                  ">20%"))

label(d$VAF_nonsilent_r) <- "Maximum VAF CH-myeloid PD"

d$n_mut_r <-
  factor(d$n_mut_r,
         labels=c("Negative", # Reference
                  "1",
                  "2",
                  "3 or more"))

label(d$n_mut_r) <- "Number of CH-myeloid PD mutations"

d$center <-
  factor(d$center,
         levels=c("MSK", "MOF", "MDA", "HCC"),
         labels=c("MSK", # Reference
                  "MOF",
                  "MDA",
                  "DFC"))

label(d$age) <- "Age"
label(d$yearslastfu) <- "Years of follow-up"
label(d$hgb) <- "Hemoglobin"
label(d$wbc) <- "White Blood Cell Count"
label(d$plt) <- "Platelet"
label(d$anc) <- "Absolute Neutrophil Count"
label(d$mcv) <- "Mean Corpuscular Volume"
label(d$rdw) <- "Red Cell Distribution Width"

d <- d %>% mutate(SystemTumorType=ifelse(SystemTumorType=="","Other",SystemTumorType))
label(d$SystemTumorType) <- "Primary Tumor Subtype"

table1 = table1(~ Gender + age + SystemTumorType + yearslastfu + hgb + rdw + mcv + wbc + anc + plt + n_mut_r + VA
F_nonsilent_r  | center*diagnosis, data=d, overall=F, output="markdown", export="ktab")

table1
```

| | MSK | | MOF | | MDA | | |
|---|---|---|---|---|---|---|---|
| | **No tMN (n=8982)** | **tMN (n=30)** | **No tMN (n=54)** | **tMN (n=13)** | **No tMN (n=54)** | **tMN (n=14)** | |
| **Gender** | | | | | | | |
| Female | 4963 (55.3%) | 12 (40.0%) | 23 (42.6%) | 6 (46.2%) | 25 (46.3%) | 3 (21.4%) | |
| Male | 4019 (44.7%) | 18 (60.0%) | 31 (57.4%) | 7 (53.8%) | 29 (53.7%) | 11 (78.6%) | |
| **Age** | | | | | | | |
| Mean (SD) | 58.5 (15.0) | 54.2 (22.4) | 72.5 (5.77) | 72.4 (4.61) | 56.4 (9.91) | 56.4 (13.6) | |
| Median [Min, Max] | 60.6 [0.249, 98.7] | 63.2 [9.82, 80.7] | 73.0 [61.0, 87.0] | 73.0 [64.0, 79.0] | 57.8 [40.1, 79.2] | 62.5 [25.0, 74.0] | 5[illegible] |
| **Primary Tumor Subtype** | | | | | | | |
| Brain | 436 (4.9%) | 1 (3.3%) | 0 (0%) | 0 (0%) | 0 (0%) | 0 (0%) | |
| Breast | 1262 (14.1%) | 3 (10.0%) | 0 (0%) | 0 (0%) | 0 (0%) | 0 (0%) | |
| Cancer of Unknown Primary | 292 (3.3%) | 0 (0%) | 0 (0%) | 0 (0%) | 0 (0%) | 0 (0%) | |
| Colorectal | 1180 (13.1%) | 2 (6.7%) | 4 (7.4%) | 2 (15.4%) | 0 (0%) | 1 (7.1%) | |
| GE | 436 (4.9%) | 2 (6.7%) | 2 (3.7%) | 1 (7.7%) | 0 (0%) | 1 (7.1%) | |
| Germ Cell | 241 (2.7%) | 6 (20.0%) | 0 (0%) | 0 (0%) | 0 (0%) | 0 (0%) | |

| | MSK | | MOF | | MDA | | |
|---|---|---|---|---|---|---|---|
| | No tMN (n=8982) | tMN (n=30) | No tMN (n=54) | tMN (n=13) | No tMN (n=54) | tMN (n=14) | |
| GU | 196 (2.2%) | 2 (6.7%) | 0 (0%) | 0 (0%) | 0 (0%) | 0 (0%) | |
| Gyn | 789 (8.8%) | 2 (6.7%) | 6 (11.1%) | 0 (0%) | 0 (0%) | 0 (0%) | |
| Head and Neck | 230 (2.6%) | 0 (0%) | 0 (0%) | 0 (0%) | 0 (0%) | 0 (0%) | |
| Liver | 32 (0.4%) | 0 (0%) | 0 (0%) | 0 (0%) | 0 (0%) | 0 (0%) | |
| Lung | 1718 (19.1%) | 4 (13.3%) | 8 (14.8%) | 1 (7.7%) | 0 (0%) | 4 (28.6%) | |
| Other | 71 (0.8%) | 2 (6.7%) | 5 (9.3%) | 2 (15.4%) | 0 (0%) | 3 (21.4%) | |
| Pancreatic | 988 (11.0%) | 3 (10.0%) | 0 (0%) | 0 (0%) | 0 (0%) | 0 (0%) | |
| Prostate | 372 (4.1%) | 1 (3.3%) | 0 (0%) | 0 (0%) | 0 (0%) | 0 (0%) | |
| Renal | 87 (1.0%) | 0 (0%) | 0 (0%) | 0 (0%) | 0 (0%) | 0 (0%) | |
| Sarcoma | 424 (4.7%) | 0 (0%) | 3 (5.6%) | 1 (7.7%) | 0 (0%) | 2 (14.3%) | |
| Skin | 138 (1.5%) | 1 (3.3%) | 0 (0%) | 0 (0%) | 0 (0%) | 0 (0%) | |
| Thyroid | 90 (1.0%) | 1 (3.3%) | 0 (0%) | 0 (0%) | 0 (0%) | 0 (0%) | |
| Lymphoma | 0 (0%) | 0 (0%) | 18 (33.3%) | 5 (38.5%) | 54 (100%) | 3 (21.4%) | |
| Multiple myeloma | 0 (0%) | 0 (0%) | 8 (14.8%) | 1 (7.7%) | 0 (0%) | 0 (0%) | |
| **Years of follow-up** | | | | | | | |
| Mean (SD) | 1.39 (0.969) | 1.42 (0.843) | 3.60 (2.80) | 2.51 (2.04) | 5.00 (0.00) | 3.57 (2.10) | |
| Median [Min, Max] | 1.17 [0.00, 4.49] | 1.34 [0.287, 3.31] | 2.80 [0.164, 10.0] | 1.66 [0.378, 7.40] | 5.00 [5.00, 5.00] | 3.00 [1.00, 8.00] | 5.5[illegible] |
| **Hemoglobin** | | | | | | | |
| Mean (SD) | 12.3 (1.87) | 11.4 (1.51) | 12.7 (1.52) | 12.2 (2.28) | 12.8 (2.07) | 12.2 (1.81) | |
| Median [Min, Max] | 12.4 [5.30, 19.0] | 11.6 [7.30, 15.0] | 12.7 [9.60, 16.0] | 12.3 [8.20, 17.4] | 13.1 [4.10, 16.3] | 12.0 [9.00, 15.3] | 1[illegible] |
| Missing | 18 (0.2%) | 2 (6.7%) | 2 (3.7%) | 0 (0%) | 0 (0%) | 0 (0%) | |
| **Red Cell Distribution Width** | | | | | | | |
| Mean (SD) | 14.5 (2.28) | 15.6 (2.62) | 47.8 (9.34) | 52.3 (10.7) | NA (NA) | NA (NA) | |
| Median [Min, Max] | 13.8 [10.7, 30.6] | 14.9 [12.3, 23.4] | 47.3 [12.9, 66.4] | 48.5 [41.6, 75.1] | NA [NA, NA] | NA [NA, NA] | 1[illegible] |
| Missing | 20 (0.2%) | 2 (6.7%) | 2 (3.7%) | 0 (0%) | 54 (100%) | 14 (100%) | |
| **Mean Corpuscular Volume** | | | | | | | |
| Mean (SD) | 89.3 (6.69) | 93.4 (9.45) | 93.4 (4.68) | 93.5 (5.39) | NA (NA) | NA (NA) | |
| Median [Min, Max] | 90.0 [9.00, 123] | 95.0 [77.0, 109] | 93.1 [84.3, 105] | 93.3 [83.8, 103] | NA [NA, NA] | NA [NA, NA] | 9[illegible] |
| Missing | 15 (0.2%) | 2 (6.7%) | 2 (3.7%) | 0 (0%) | 54 (100%) | 14 (100%) | |
| **White Blood Cell Count** | | | | | | | |
| Mean (SD) | 7.59 (4.41) | 5.63 (2.51) | 6.55 (2.19) | 6.05 (2.60) | 7.14 (2.81) | 5.94 (1.81) | |
| Median [Min, Max] | 6.80 [0.100, 171] | 5.10 [2.10, 12.0] | 6.16 [2.68, 14.8] | 5.21 [3.01, 12.8] | 6.70 [3.20, 19.5] | 5.50 [3.60, 9.40] | 5.[illegible] |
| Missing | 19 (0.2%) | 2 (6.7%) | 2 (3.7%) | 0 (0%) | 0 (0%) | 0 (0%) | |
| **Absolute Neutrophil Count** | | | | | | | |
| Mean (SD) | 5.29 (3.68) | 3.85 (2.18) | 6.22 (11.0) | 3.31 (1.14) | 5.06 (2.63) | 3.72 (1.57) | |
| Median [Min, Max] | 4.50 [0.00, 77.1] | 3.30 [0.800, 9.10] | 4.16 [1.67, 77.5] | 3.12 [1.40, 5.95] | 4.51 [1.85, 16.6] | 3.20 [1.80, 6.64] | 3.[illegible] |
| Missing | 57 (0.6%) | 2 (6.7%) | 8 (14.8%) | 0 (0%) | 0 (0%) | 0 (0%) | |
| **Platelet** | | | | | | | |
| Mean (SD) | 261 (109) | 209 (99.4) | 175 (64.2) | 166 (56.8) | 275 (95.6) | 208 (51.6) | |
| Median [Min, Max] | 244 [8.00, 1240] | 193 [10.0, 402] | 170 [22.0, 320] | 150 [103, 281] | 256 [105, 559] | 214 [96.0, 303] | 2[illegible] |
| Missing | 30 (0.3%) | 2 (6.7%) | 2 (3.7%) | 0 (0%) | 0 (0%) | 0 (0%) | |
| **Number of CH-myeloid PD mutations** | | | | | | | |
| Negative | 7595 (84.6%) | 16 (53.3%) | 41 (75.9%) | 7 (53.8%) | 54 (100%) | 5 (35.7%) | |
| 1 | 1133 (12.6%) | 6 (20.0%) | 9 (16.7%) | 3 (23.1%) | 0 (0%) | 5 (35.7%) | |
| 2 | 196 (2.2%) | 5 (16.7%) | 4 (7.4%) | 0 (0%) | 0 (0%) | 1 (7.1%) | |
| 3 or more | 58 (0.6%) | 3 (10.0%) | 0 (0%) | 3 (23.1%) | 0 (0%) | 3 (21.4%) | |
| **Maximum VAF CH-myeloid PD** | | | | | | | |
| Negative | 7595 (84.6%) | 16 (53.3%) | 41 (75.9%) | 7 (53.8%) | 54 (100%) | 5 (35.7%) | |
| 2-5% | 624 (6.9%) | 1 (3.3%) | 6 (11.1%) | 1 (7.7%) | 0 (0%) | 1 (7.1%) | |
| 5-10% | 353 (3.9%) | 2 (6.7%) | 2 (3.7%) | 2 (15.4%) | 0 (0%) | 2 (14.3%) | |
| 10-20% | 231 (2.6%) | 5 (16.7%) | 3 (5.6%) | 1 (7.7%) | 0 (0%) | 3 (21.4%) | |
| >20% | 179 (2.0%) | 6 (20.0%) | 2 (3.7%) | 2 (15.4%) | 0 (0%) | 3 (21.4%) | |

#Tabulate tMN analysis

```
#Tabulate tMN analysis
print('All tmn risk set samples')
```

```
## [1] "All tmn risk set samples"
```

```
M_tmn_wide_st<- filter(M_tmn_wide,(center=="MSK" | center=="MDA" | center=="HCC" | center=="MOF"))
M_tmn_wide_st2 <- M_tmn_wide_st %>% filter(!(any_therapy==0 & center=="MSK"))
table(M_tmn_wide_st2$post_tmn)
```

```
##
##    0    1
## 9474   75
```

```
print('Time to transformation')
```

```
## [1] "Time to transformation"
```

```
case <- filter(M_tmn_wide_st,post_tmn==1)
psych::describe(case$timelastfu)
```

```
##    vars  n    mean     sd median trimmed    mad min     max   range skew
## X1    1 75 1103.75 917.34    791  991.15 631.22 105 4422.03 4317.03 1.26
##    kurtosis     se
## X1     1.14 105.92
```

```
#Tabulate paired tMN analysis
print('All paired samples')
```

```
## [1] "All paired samples"
```

```
filter(M_tmn_wide,has_serial == 1) %>% nrow()
```

```
## [1] 35
```

```
print('Time to trans')
```

```
## [1] "Time to trans"
```

```
case <- filter(M_tmn_wide_st,has_serial==1)
psych::describe(case$timelastfu)
```

```
##    vars  n   mean     sd median trimmed    mad min  max range skew
## X1    1 35 864.39 598.77  730.5  786.39 541.52 138 2702  2564 1.14
##    kurtosis     se
## X1     1.08 101.21
```

```
print("Mutation positive at time of tMN")
```

```
## [1] "Mutation positive at time of tMN"
```

```
M_tmn_long %>% filter(has_serial == 1 & VAF_2>0) %>% pull(center_id) %>% unique() %>% length()
```

```
## [1] 33
```

```
print("Mutation positive at time of CH")
```

```
## [1] "Mutation positive at time of CH"
```

```
M_tmn_long %>% filter(has_serial == 1 & VAF_1>0) %>% pull(center_id) %>% unique() %>% length()
```

```
## [1] 19
```

```
print("Multiple mutations at time of CH")
```

```
## [1] "Multiple mutations at time of CH"
```

```
testing<- M_tmn_long %>% filter(has_serial == 1 & VAF_1>0) %>% group_by(center_id) %>% summarise(mutnum_ch = leng
th(VAF_1>0))
filter(testing,mutnum_ch>1) %>% nrow()
```

```
## [1] 13
```

```
print("More mutations at time of tMN")
```

```
## [1] "More mutations at time of tMN"
```

```
M_tmn_long %>% filter(has_serial == 1 & VAF_1==0) %>% pull(center_id) %>% unique() %>% length()
```

```
## [1] 19
```

```
print("Has a TP53 mutation at tMN")
```

```
## [1] "Has a TP53 mutation at tMN"
```

```
filter(M_tmn_long,has_serial == 1 & Gene=="TP53" & VAF_2>0) %>% pull(center_id) %>% unique() %>% length()
```

```
## [1] 14
```

```
print("Has a TP53 mutation and is detectable at CH")
```

```
## [1] "Has a TP53 mutation and is detectable at CH"
```

```
filter(M_tmn_long,has_serial == 1 & Gene=="TP53" & VAF_1>0) %>% pull(center_id) %>% unique() %>% length()
```

```
## [1] 10
```

```
#Karotype not missing and TP53 positive
M_tmn_long %>% filter(has_serial == 1 & Gene=="TP53" & is.na(karyotype_classification)) %>% pull(center_id) %>% u
nique() %>% length()
```

```
## [1] 1
```

```
print("Among mutation positive, has a TP53 mutation and karyotype abnormal")
```

```
## [1] "Among mutation positive, has a TP53 mutation and karyotype abnormal"
```

```
filter(M_tmn_long,has_serial == 1 & Gene=="TP53" & M_tmn_long$karyotype_classification=="abnormal") %>% pull(cent
er_id) %>% unique() %>% length()
```

```
## [1] 12
```

```
print("Has multiple CH-PD mutations")
```

```
## [1] "Has multiple CH-PD mutations"
```

```
filter(M_tmn_wide,has_serial == 1 & n_mut>1 ) %>% nrow()
```

```
## [1] 8
```

```
#Karotype normal
M_tmn_wide %>% filter(has_serial == 1 & karyotype_classification =="normal") %>% pull(center_id) %>% unique() %>%
length()
```

```
## [1] 6
```

```
#Karotype not missing
M_tmn_wide %>% filter(has_serial == 1 & is.na(karyotype_classification)) %>% pull(center_id) %>% unique() %>% len
gth()
```

```
## [1] 3
```

```
#Mutation negative at time of CH
mut_pos_ids = M_tmn_long %>% filter(has_serial == 1 & VAF_1>0) %>% pull(center_id) %>% unique()
all_ids = M_tmn_wide %>% filter(has_serial == 1) %>% pull(center_id) %>% unique()
neg_ids = setdiff(all_ids,mut_pos_ids)
#Karyotype abnormal and mut neg at time CH
ka_negch <- filter(M_tmn_wide, has_serial==1 & (karyotype_classification=="abnormal" |karyotype_classification==
"complex") & center_id %in% neg_ids)  %>% pull(center_id)
#MLL and mut neg at time CH
mll_negch <- filter(M_tmn_wide, has_serial==1 & MLL_rearrangement==1 & (center_id %in% neg_ids))  %>% pull(center
_id)
#Mut neg at CH but not tMN
M_tmn_long %>% filter(has_serial == 1 & VAF_2>0 & VAF_1==0 & (center_id %in% neg_ids)) %>% pull(center_id) %>% un
ique() %>% length()
```

```
## [1] 14
```

#Supp Figure 10 tMN regression combined with healthy

```
#Full CH model only including genes present in all studies
gene_vars = c("TET2","TP53","U2AF1","SF3B1")
mut_vars = c("VAF_nonsilent_r","n_mut_r")
demo_vars = c('age', 'Gender','strata(center)')

D = M_tmn_wide_st

form = paste0(
    'Surv(timelastfu, post_tmn) ~ ',
    paste(
        c(
            gene_vars,
            mut_vars,
            demo_vars
        ),
        collapse = ' + '
    )
)

cox <- coxph(formula = as.formula(form), data = D, na.action = 'na.omit')
multi <- cox %>% get_model_data()
multi2 <- multi %>% mutate(group=case_when(
  term %in% gene_vars ~ "Gene",
  term=="n_mut_r" ~ "Mutation Number",
  term=="VAF_nonsilent_r" ~ "Maximum VAF"
))
multi2 <- multi2 %>% rename(mut_var=term) %>% cbind(study="tMN")

#Healthy
M_tmn_h <- M_tmn_wide

#Main effect of CH
M_tmn_h <- M_tmn_h %>% mutate(center =fct_relevel(center, "PMC", "WSU"))

cox <- coxph(Surv(timelastfu, post_tmn) ~ ch_my_pd + age + Gender + strata(center), data= M_tmn_h, na.action = 'n
a.omit')
cox
```

```
## Call:
## coxph(formula = Surv(timelastfu, post_tmn) ~ ch_my_pd + age +
##     Gender + strata(center), data = M_tmn_h, na.action = "na.omit")
##
##              coef exp(coef) se(coef)      z       p
## ch_my_pd  1.87281   6.50654  0.13034 14.369 < 2e-16
## age      -0.01727   0.98288  0.00650 -2.657 0.00788
## GenderM   0.14912   1.16082  0.12701  1.174 0.24034
##
## Likelihood ratio test=183.8  on 3 df, p=< 2.2e-16
## n= 11807, number of events= 261
##    (3 observations deleted due to missingness)
```

```
cox <- coxph(Surv(timelastfu, post_tmn) ~ ch_my_pd*center + age + Gender + strata(center), data=M_tmn_h, na.actio
n = 'na.omit')
cox
```

```
## Call:
## coxph(formula = Surv(timelastfu, post_tmn) ~ ch_my_pd * center +
##     age + Gender + strata(center), data = M_tmn_h, na.action = "na.omit")
##
##                         coef exp(coef)  se(coef)      z       p
## ch_my_pd            2.018767  7.529039  0.164318 12.286 < 2e-16
## centerWSU                 NA        NA  0.000000     NA      NA
## centerHCC                 NA        NA  0.000000     NA      NA
## centerMDA                 NA        NA  0.000000     NA      NA
## centerMOF                 NA        NA  0.000000     NA      NA
## centerMSK                 NA        NA  0.000000     NA      NA
## age                -0.017822  0.982336  0.006687 -2.665 0.00769
## GenderM             0.171787  1.187425  0.128637  1.335 0.18173
## ch_my_pd:centerWSU -0.766425  0.464671  0.437716 -1.751 0.07995
## ch_my_pd:centerHCC -0.258140  0.772487  0.520391 -0.496 0.61986
## ch_my_pd:centerMDA  0.551610  1.736046  0.628595  0.878 0.38020
## ch_my_pd:centerMOF -1.225652  0.293566  0.581380 -2.108 0.03502
## ch_my_pd:centerMSK -0.058823  0.942873  0.406607 -0.145 0.88497
##
## Likelihood ratio test=192.3  on 8 df, p=< 2.2e-16
## n= 11807, number of events= 261
##    (3 observations deleted due to missingness)
```

```
#Full model with max VAF and mutation number modeled as continuous

D = M_tmn_h

form = paste0(
    'Surv(timelastfu, post_tmn) ~ ',
    paste(
        c(
            gene_vars,
            mut_vars,
            demo_vars
        ),
        collapse = ' + '
    )
)

cox <- coxph(formula = as.formula(form), data = D, na.action = 'na.omit')
multi <- cox %>% get_model_data()
multi2_h <- multi %>% mutate(group=case_when(
  term %in% gene_vars ~ "Gene",
  term=="n_mut_r" ~ "Mutation Number",
  term=="VAF_nonsilent_r" ~ "Maximum VAF"
))
multi2_h <- multi2_h %>% rename(mut_var=term) %>% cbind(study="Primary AML")

combo <- rbind(multi2_h, multi2)

combo <- combo %>% mutate(mut_var2=case_when(mut_var=="VAF_nonsilent_r" ~ "Maximum VAF",
                                            mut_var=="n_mut_r" ~ "Mutation Number",
                                            TRUE ~ as.character(mut_var)))

p <- combo %>%
plot_forest(
    x = "mut_var2",
    fill ="study",
    col = "study",
   eb_w = 0,
      eb_s = 0.3,
      ps = 1.5,
      or_s = 2,
      nudge = -0.5,
      dodge_width = 0.5)+
   scale_fill_nejm() +
scale_color_nejm() +
ylab('HR') +
xlab("") +
facet_grid(group ~ ., scales = "free_y", space = "free_y") +
#  panel_theme +
theme(
    strip.placement = 'top',
      strip.text = element_blank(),
    axis.text.y = element_text(size = 7),
    axis.title = element_text(size = 7),
    axis.text.x = element_text(size = 7),
    legend.text = element_text(size=7),
      legend.title = element_text(size=7)
)

do_plot(p, "supp_figure_10.png", w = 5, h = 4, save_pdf = T)
```

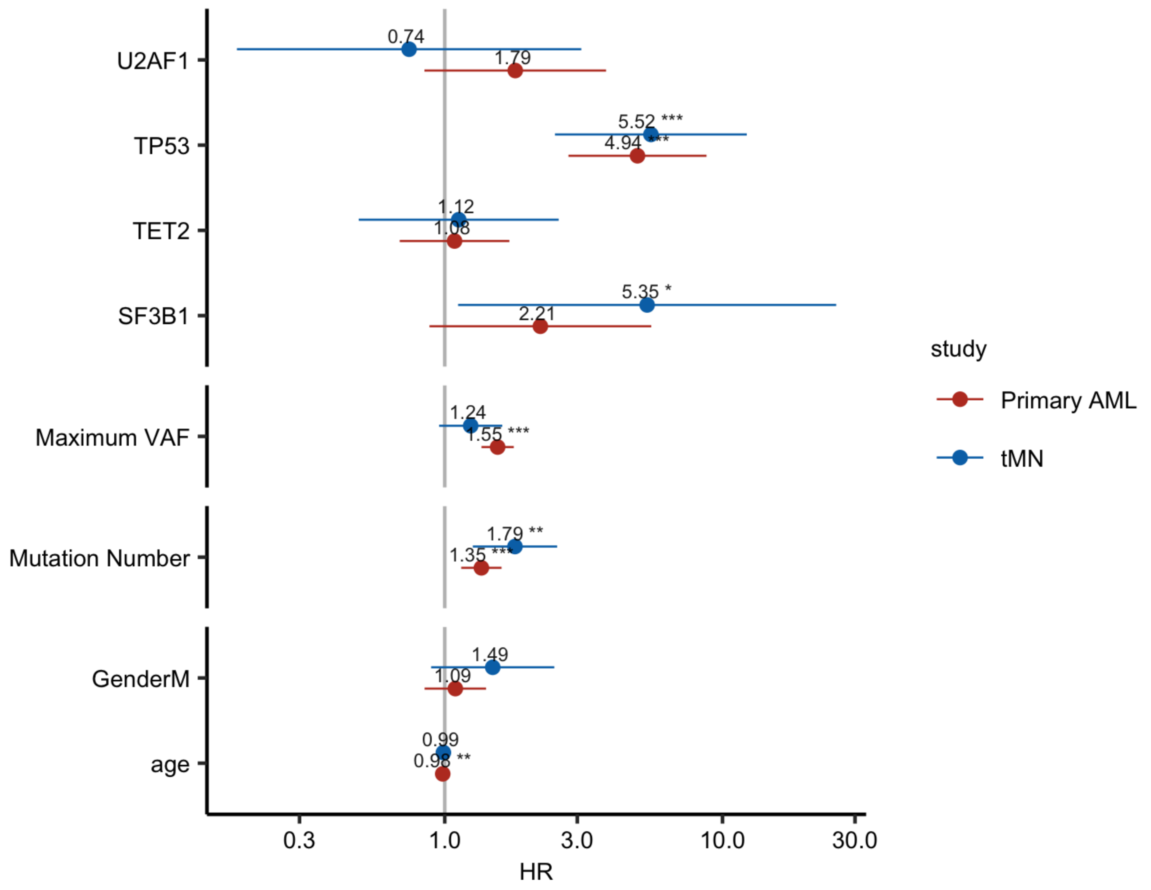
U2AF1
TP53
TET2
SF3B1
Maximum VAF
Mutation Number
GenderM
age
0.74
1.79
5.52 ***
4.94 ***
1.12
1.08
5.35 *
2.21
1.24
1.55 ***
1.79 **
1.35 ***
1.49
1.09
0.99
0.98 **
0.3
1.0
3.0
10.0
30.0
HR
study
Primary AML
tMN

### tMN heatmap

```
## gene set
mda_genes = read_tsv('../data/external/tmn/mda_panel.tsv', col_types = cols()) %>% pull(Gene)
mof_genes = read_tsv('../data/external/tmn/mof_panel.tsv', col_types = cols()) %>% pull(Gene)
msk_genes_v3 = read_tsv('../data/external/cv3_genelist.txt', col_types = cols()) %>% pull(Gene)
msk_genes_v5 = read_tsv('../data/external/cv5_genelist.txt', col_types = cols()) %>% pull(Gene)
msk_genes_v6 = read_tsv('../data/external/cv6_genelist.txt', col_types = cols()) %>% pull(Gene)
myeloid_genes = read_tsv("../data/external/iwg-pm-genes.txt", col_names = F, , col_types = cols()) %>%
    unlist() %>% unname() %>% as.character()

gene_set = Reduce(
    intersect,
    list(mda_genes, mof_genes, msk_genes_v3,
         msk_genes_v5, msk_genes_v6, myeloid_genes)
)

panel_to_genes = list(
    'v5' = msk_genes_v5, 'v3' = msk_genes_v3, 'v6' = msk_genes_v6,
    'IWG' = myeloid_genes, 'MDA' = mda_genes, 'MOF' = mof_genes)

gene_set = c(gene_set, 'SRSF2', 'NF1', 'PPM1D')

# cytogenetics data
CYTO = read_tsv('../data/external/tmn_cyto_merged.txt', col_types = cols())

CYTO = CYTO %>%
    mutate(notation = paste0(Chrom, ifelse(Alt == 'gain', '+', '-')))

## produce heatmap
D = rbind(
        # SNV data
        M_tmn_long %>%
            filter(Gene %in% gene_set) %>%
            filter(center %in% c('MSK', 'MOF', 'MDA')) %>%
            filter(post_tmn == 1) %>%
            select(center_id, center, Gene, VAF_1, VAF_2),
        # complex karyo and MLL
        M_tmn_wide %>% select(center_id, center, complex_karyotype, MLL_rearrangement) %>%
            melt(measure.vars = c('complex_karyotype', 'MLL_rearrangement'), variable.name = 'Gene') %>%
            filter(value == 1) %>%
            select(-value) %>%
            mutate(VAF_1 = 0, VAF_2 = 1),
        # other cytogenetics
        CYTO %>%
            filter(timepoint == 'tmn' & source != 'cnacs') %>%
            filter(str_detect(notation, '^5q-$|^7q-$|^5-$|^7-$|^8\\+|Y')) %>%
            mutate(VAF_1 = 0, VAF_2 = 1) %>%
            mutate(Gene = notation) %>%
            mutate(
                Gene = ifelse(Gene %in% c('5q-', '5-'), '-5/5q', Gene),
                Gene = ifelse(Gene %in% c('7q-', '7-'), '-7/7q', Gene)
            ) %>%
            select(center_id, center, Gene, VAF_1, VAF_2)
    ) %>%
    # only include patients that had serial
    filter(center_id %in% pull(filter(M_tmn_wide, has_serial == 1), center_id)) %>%
    rowwise() %>%
    mutate(
        # upper limit for alpha gradient
        VAF_1 = min(VAF_1, 0.5),
        VAF_2 = min(VAF_2, 0.5)
    ) %>%
    ungroup() %>%
    # order genes
    group_by(Gene) %>%
    mutate(gene_n = sum(VAF_2 != 0)) %>%
    ungroup() %>%
    arrange(gene_n) %>%
    mutate(
        Gene = factor(
            Gene,
            rev(unique(c(
                'TP53', 'complex_karyotype',
                'TET2', 'DNMT3A', 'ASXL1',
                'RUNX1',  'U2AF1', 'SRSF2',
                'STAG2','EZH2','KDM6A','ETV6','SUZ12','CREBBP','RAD21',
                'IDH1','IDH2','NRAS','KRAS','HRAS','FLT3','BRAF',
                rev(Gene[!Gene %in% c('NF1', 'PPM1D', 'MLL_rearrangement', '-5/5q', '-7/7q', '8+', 'Y-')]),
                'NF1', 'PPM1D', 'MLL_rearrangement', '-5/5q', '-7/7q', '8+', 'Y-'
            )))
```

```
    )
) %>%
melt(
    measure.vars = c('VAF_1', 'VAF_2'),
    variable.name = 'timepoint',
    value.name = 'VAF'
) %>%
mutate(timepoint = ifelse(timepoint == 'VAF_1', 'N01', 'tMN')) %>%
# add grey blocks to genes not covered
rbind(
    M_tmn_wide %>%
    filter(has_serial == 1) %>%
    select(center_id, serial_bait_1, center) %>%
    rowwise() %>%
    mutate(
        Gene = list(gene_set[!gene_set %in% panel_to_genes[[serial_bait_1]]])
    ) %>%
    tidyr::unnest() %>%
    mutate(VAF = -1, gene_n = 0) %>%
    select(-serial_bait_1) %>%
    mutate(timepoint = 'N01')
) %>%
rbind(
    M_tmn_wide %>%
    filter(has_serial == 1) %>%
    select(center_id, serial_bait_2, center) %>%
    rowwise() %>%
    mutate(
        Gene = list(gene_set[!gene_set %in% panel_to_genes[[serial_bait_2]]])
    ) %>%
    tidyr::unnest() %>%
    mutate(VAF = -1, gene_n = 0) %>%
    select(-serial_bait_2) %>%
    mutate(timepoint = 'tMN')
) %>%
# get DMP ID
left_join(
    M_tmn_wide[c('center_id', 'center', 'DMP_ID')],
    by = c('center_id', 'center')
) %>%
# order patients
{
    mutate(.,
        center_id = factor(
            center_id,
            group_by(., center_id) %>%
            filter(VAF != -1) %>%
            summarise(
                mut_n = n(),
                max_gene_n = max(gene_n),
                TP53 = any(Gene == 'TP53'),
                TET2 = any(Gene == 'TET2'),
                DNMT3A = any(Gene == 'DNMT3A'),
                ASXL1 = any(Gene == 'ASXL1'),
                RUNX1 = any(Gene == 'RUNX1'),
                MLL = any(Gene == 'MLL_rearrangement'),
                Complex = any(Gene == 'complex_karyotype'),
                SRSF2 = any(Gene == 'SRSF2'),
                U2AF1 = any(Gene == 'U2AF1'),
                `-5/5q` = any(Gene == '-5/5q'),
                `-7/7q` = any(Gene == '-7/7q'),
                `8+` = any(Gene == '8+'),
                `Y-` = any(Gene == 'Y-')
            ) %>%
            arrange(
                MLL,
                -TP53,
                -Complex,
                -TET2,
                -DNMT3A,
                -ASXL1,
                -RUNX1,
                -U2AF1,
                -SRSF2,
                -`-5/5q`,
                -`-7/7q`,
                -`8+`,
                -`Y-`,
                -mut_n
            ) %>% pull(center_id) %>% unique
```

```
            )
        )
    } %>%
    mutate(VAF = case_when(
        VAF > 0 ~ 'present',
        VAF == 0 ~ 'absent',
        VAF < 0 ~ 'unknown'
      )
    ) %>%
    # create patient labels
    mutate(patient_label = ifelse(center == 'MSK', DMP_ID, as.character(center_id))) %>%
    mutate(patient_label = paste0(patient_label, '-', timepoint)) %>%
    arrange(center_id, timepoint) %>%
    mutate(patient_label = factor(patient_label, unique(patient_label)))

p_main = ggplot(
        D,
        aes(x = patient_label, y = Gene, fill = VAF)
    ) +
    geom_tile(
        color = 'white',
        size = 0.4
    ) +
    scale_fill_manual(
      values = c(
        'present' = 'steelblue',
        'absent' = 'white',
        'unknown' = 'gainsboro'
      )
    ) +
    theme_bw() +
    theme(
        axis.ticks.x = element_blank(),
        axis.text.x = element_text(angle = 90),
        panel.grid = element_blank(),
        legend.position = 'left',
        plot.margin = margin(t = 0, r = 0, b = 0, l = 0),
        legend.key.size = unit(12, 'pt'),
        legend.text = element_text(size = 8),
        legend.title = element_blank(),
        legend.background = element_rect(fill = 'cornsilk3')
    ) +
    geom_vline(
        xintercept = seq(1, length(unique(D$center_id)) - 1) * 2 + 0.5,
        linetype = 'dashed',
        alpha = 0.5,
        size = 0.3
    ) +
    xlab('') +
    ylab('') +
    scale_x_discrete(expand = c(0,0)) +
    scale_y_discrete(expand = c(0,0))

p_right = ggplot(
        D %>%
        mutate(cohort_size = length(unique(center_id))) %>%
        group_by(Gene) %>%
        summarise(prop = sum(timepoint == 'tMN' & VAF == 'present')/unique(cohort_size)) %>%
        ungroup(),
        aes(x = Gene, y = prop)
    ) +
    geom_col(width = 0.925) +
    coord_flip() +
    theme_bw() +
    xlab('') +
    ylab('proportion of patients') +
    theme(
        axis.title.y = element_blank(),
        axis.line.y = element_blank(),
        axis.ticks.y = element_blank(),
        axis.text.y = element_blank(),
        panel.grid = element_blank(),
        legend.position = 'none',
        panel.border = element_blank(),
        plot.margin = margin(t = 0, r = 0, b = 0, l = 0)
    ) +
    scale_x_discrete(expand = c(0,0)) +
    scale_y_continuous(expand = c(0,0), limits = c(0, 0.5), position = 'left')

p_top = D %>%
```

```
    group_by(patient_label) %>%
    summarise(n = sum(VAF == 'present')) %>%
    ggplot(
        aes(x = patient_label, y = n)
    ) +
    geom_col(width = 0.925) +
    theme_bw() +
    xlab('') +
    ylab('mutation number') +
    theme(
        axis.title.x = element_blank(),
        axis.line.x = element_blank(),
        axis.ticks.x = element_blank(),
        axis.text.x = element_blank(),
        panel.grid = element_blank(),
        legend.position = 'none',
        panel.border = element_blank(),
        plot.margin = margin(t = 0, r = 0, b = 0, l = 0)
    ) +
    scale_x_discrete(expand = c(0,0)) +
    scale_y_continuous(expand = c(0,0), position = 'left', breaks = scales::pretty_breaks())

p = p_top +
  plot_spacer() +
  p_main + p_right +
  plot_layout(widths = c(10,1), heights = c(2,10))

do_plot(p, 'tmn_heatmap.png', w = 18, h = 8, save_pdf = T)
```

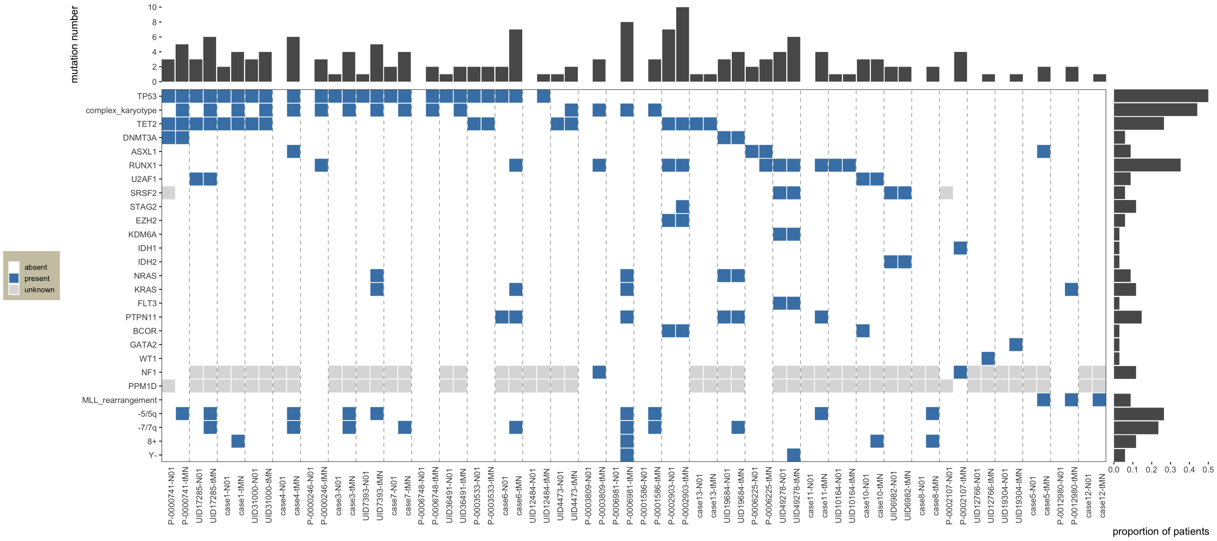


### tMN lineplot

```
source('../utils/plot_tmn.R')

all_ids = M_tmn_wide %>% filter(has_serial == 1) %>%
    pull(center_id)
plots = list()
for (patient in all_ids) {

    patient_id = M_tmn_wide %>%
        filter(center_id == patient) %>%
        pull(DMP_ID) %>%
        unique

    if (is.na(patient_id) | patient_id == '') {
        patient_id = patient
    }
    disease_names = c('acute_myeloid_leukemia' = 'AML', 'myelodysplastic_syndrome' = 'MDS', 'ctmn' = 'CTMN')
    age = M_tmn_wide %>% filter(center_id == patient) %>% pull(age) %>% unique %>% signif(2)
    disease = M_tmn_wide %>% filter(center_id == patient) %>% pull(postimp_heme_dx_type) %>% unique
    M_patient = M_tmn_long %>% filter(center_id == patient)
    title = paste0(patient_id, '\n', disease_names[disease], ', age ', age, '')

    if (sum(M_patient$VAF_1) > 0) {
        p = tmn_panel(
                M = M_patient,
                Tr = T_merged %>% filter(center_id == patient),
                title = title,
                font_size = 5
            )

        plots[[patient]] = p
    }
}
```

```
## Warning in melt.data.table(., measure.vars = list(date = c("date_1",
## "date_2"), : 'measure.vars' [date_1, date_2] are not all of the same
## type. By order of hierarchy, the molten data value column will be of type
## 'double'. All measure variables not of type 'double' will be coerced too.
## Check DETAILS in ?melt.data.table for more on coercion.

## Warning in melt.data.table(., measure.vars = list(date = c("date_1",
## "date_2"), : 'measure.vars' [date_1, date_2] are not all of the same
## type. By order of hierarchy, the molten data value column will be of type
## 'double'. All measure variables not of type 'double' will be coerced too.
## Check DETAILS in ?melt.data.table for more on coercion.

## Warning in melt.data.table(., measure.vars = list(date = c("date_1",
## "date_2"), : 'measure.vars' [date_1, date_2] are not all of the same
## type. By order of hierarchy, the molten data value column will be of type
## 'double'. All measure variables not of type 'double' will be coerced too.
## Check DETAILS in ?melt.data.table for more on coercion.

## Warning in melt.data.table(., measure.vars = list(date = c("date_1",
## "date_2"), : 'measure.vars' [date_1, date_2] are not all of the same
## type. By order of hierarchy, the molten data value column will be of type
## 'double'. All measure variables not of type 'double' will be coerced too.
## Check DETAILS in ?melt.data.table for more on coercion.

## Warning in melt.data.table(., measure.vars = list(date = c("date_1",
## "date_2"), : 'measure.vars' [date_1, date_2] are not all of the same
## type. By order of hierarchy, the molten data value column will be of type
## 'double'. All measure variables not of type 'double' will be coerced too.
## Check DETAILS in ?melt.data.table for more on coercion.

## Warning in melt.data.table(., measure.vars = list(date = c("date_1",
## "date_2"), : 'measure.vars' [date_1, date_2] are not all of the same
## type. By order of hierarchy, the molten data value column will be of type
## 'double'. All measure variables not of type 'double' will be coerced too.
## Check DETAILS in ?melt.data.table for more on coercion.

## Warning in melt.data.table(., measure.vars = list(date = c("date_1",
## "date_2"), : 'measure.vars' [date_1, date_2] are not all of the same
## type. By order of hierarchy, the molten data value column will be of type
## 'double'. All measure variables not of type 'double' will be coerced too.
## Check DETAILS in ?melt.data.table for more on coercion.

## Warning in melt.data.table(., measure.vars = list(date = c("date_1",
## "date_2"), : 'measure.vars' [date_1, date_2] are not all of the same
## type. By order of hierarchy, the molten data value column will be of type
## 'double'. All measure variables not of type 'double' will be coerced too.
## Check DETAILS in ?melt.data.table for more on coercion.

## Warning in melt.data.table(., measure.vars = list(date = c("date_1",
## "date_2"), : 'measure.vars' [date_1, date_2] are not all of the same
## type. By order of hierarchy, the molten data value column will be of type
## 'double'. All measure variables not of type 'double' will be coerced too.
## Check DETAILS in ?melt.data.table for more on coercion.

## Warning in melt.data.table(., measure.vars = list(date = c("date_1",
## "date_2"), : 'measure.vars' [date_1, date_2] are not all of the same
## type. By order of hierarchy, the molten data value column will be of type
## 'double'. All measure variables not of type 'double' will be coerced too.
## Check DETAILS in ?melt.data.table for more on coercion.

## Warning in melt.data.table(., measure.vars = list(date = c("date_1",
## "date_2"), : 'measure.vars' [date_1, date_2] are not all of the same
## type. By order of hierarchy, the molten data value column will be of type
## 'double'. All measure variables not of type 'double' will be coerced too.
## Check DETAILS in ?melt.data.table for more on coercion.

## Warning in melt.data.table(., measure.vars = list(date = c("date_1",
## "date_2"), : 'measure.vars' [date_1, date_2] are not all of the same
## type. By order of hierarchy, the molten data value column will be of type
## 'double'. All measure variables not of type 'double' will be coerced too.
## Check DETAILS in ?melt.data.table for more on coercion.

## Warning in melt.data.table(., measure.vars = list(date = c("date_1",
## "date_2"), : 'measure.vars' [date_1, date_2] are not all of the same
## type. By order of hierarchy, the molten data value column will be of type
## 'double'. All measure variables not of type 'double' will be coerced too.
## Check DETAILS in ?melt.data.table for more on coercion.

## Warning in melt.data.table(., measure.vars = list(date = c("date_1",
```

```
## "date_2"), : 'measure.vars' [date_1, date_2] are not all of the same
## type. By order of hierarchy, the molten data value column will be of type
## 'double'. All measure variables not of type 'double' will be coerced too.
## Check DETAILS in ?melt.data.table for more on coercion.

## Warning in melt.data.table(., measure.vars = list(date = c("date_1",
## "date_2"), : 'measure.vars' [date_1, date_2] are not all of the same
## type. By order of hierarchy, the molten data value column will be of type
## 'double'. All measure variables not of type 'double' will be coerced too.
## Check DETAILS in ?melt.data.table for more on coercion.

## Warning in melt.data.table(., measure.vars = list(date = c("date_1",
## "date_2"), : 'measure.vars' [date_1, date_2] are not all of the same
## type. By order of hierarchy, the molten data value column will be of type
## 'double'. All measure variables not of type 'double' will be coerced too.
## Check DETAILS in ?melt.data.table for more on coercion.

## Warning in melt.data.table(., measure.vars = list(date = c("date_1",
## "date_2"), : 'measure.vars' [date_1, date_2] are not all of the same
## type. By order of hierarchy, the molten data value column will be of type
## 'double'. All measure variables not of type 'double' will be coerced too.
## Check DETAILS in ?melt.data.table for more on coercion.

## Warning in melt.data.table(., measure.vars = list(date = c("date_1",
## "date_2"), : 'measure.vars' [date_1, date_2] are not all of the same
## type. By order of hierarchy, the molten data value column will be of type
## 'double'. All measure variables not of type 'double' will be coerced too.
## Check DETAILS in ?melt.data.table for more on coercion.

## Warning in melt.data.table(., measure.vars = list(date = c("date_1",
## "date_2"), : 'measure.vars' [date_1, date_2] are not all of the same
## type. By order of hierarchy, the molten data value column will be of type
## 'double'. All measure variables not of type 'double' will be coerced too.
## Check DETAILS in ?melt.data.table for more on coercion.
```

```
panel = ggarrange(
    plotlist = plots,
    ncol = 5,
    nrow = 4,
    align = 'hv'
)
do_plot(panel, 'tmn_lineplot.png', w = 10, h = 7, save_pdf = T)
```

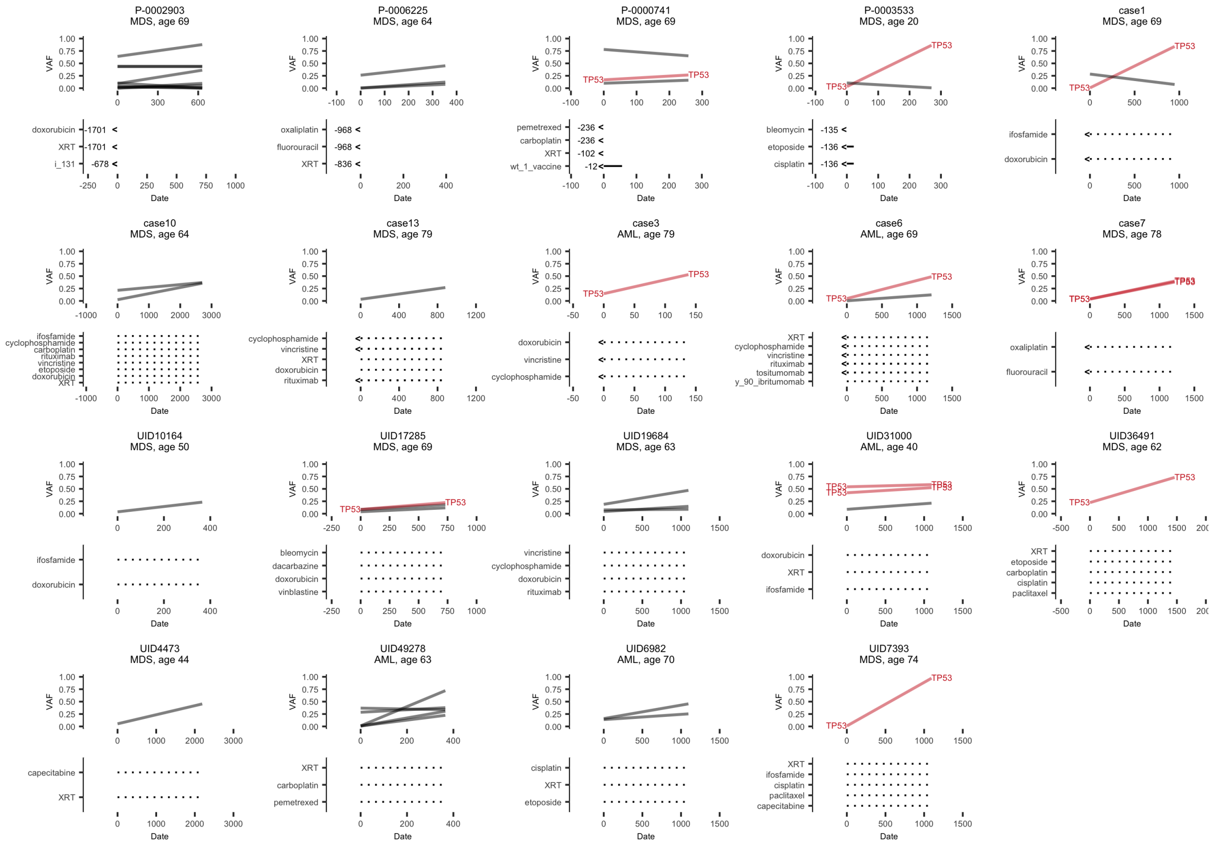
